# ADARs regulate cuticle collagen expression to promote survival to pathogen infection

**DOI:** 10.1101/2023.05.03.539277

**Authors:** Alfa Dhakal, Chinnu Salim, Mary Skelly, Yarden Amichan, Ayelet T. Lamm, Heather A. Hundley

**Affiliations:** Cell, Molecular and Cancer Biology Graduate Program, Indiana University School of Medicine-Bloomington, Bloomington, IN 47405, USA; Department of Biology, Indiana University, Bloomington IN 47405, USA; Faculty of Biology, Technion Institute of Technology, Haifa, Israel

**Keywords:** innate immunity, RNA editing, double-stranded RNA (dsRNA), *C. elegans*, *P. aeruginosa*, RNA modification, post-transcriptional regulation, RNA binding protein

## Abstract

**Background:** In all organisms, the innate immune system defends against pathogens through basal expression of molecules that provide critical barriers to invasion and inducible expression of effectors that combat infection. The adenosine deaminase that act on RNA (ADAR) family of RNA binding proteins has been reported to influence innate immunity in metazoans. However, studies on the susceptibility of ADAR mutant animals to infection are largely lacking.

**Results:** Here, by analyzing *adr-1* and *adr-2* null mutants in well-established slow-killing assays, we find that both *Caenorhabditis elegans* ADARs are important for organismal survival to gram-negative and gram-positive bacteria, all of which are pathogenic to humans. Furthermore, our high-throughput sequencing and genetic analysis reveal that ADR-1 and ADR-2 function in the same pathway to regulate collagen expression. Consistent with this finding, our scanning electron microscopy studies indicate *adr-1*;*adr-2* mutant animals also have altered cuticle morphology prior to pathogen exposure.

**Conclusions:** Our data uncover a critical role of the *C. elegans* ADAR family of RNA binding proteins in promoting cuticular collagen expression, which represents a new post-transcriptional regulatory node that influences the extracellular matrix. In addition, we provide the first evidence that ADAR mutant animals have altered susceptibility to infection with several opportunistic human pathogens, suggesting a broader role of ADARs in altering physical barriers to infection to influence innate immunity.

## Background

Pathogen infection is a major environmental threat that results in agricultural devastation and economic loss and serves as a major cause of human mortality/morbidity. To counter these attacks, plants and animals employ both physical barriers and physiological responses to resist and kill invading pathogens [1]. The most well studied innate immune responses are the evolutionary conserved signaling pathways, wherein the pathogenic “signal” is recognized by the host and triggers gene expression changes that produce cellular effectors capable of promoting organismal survival [2, 3]. Roles for RNA binding proteins (RBPs) in modulating pathogenic signal recognition have been examined [4, 5], particularly for viral infection as RNA can be the carrier of viral genomic information.

Members of the adenosine deaminase that act on RNA (ADAR) family of RBPs have well established roles during viral infection [6]. The initial focus on ADARs and virus infection was in large part because double-stranded RNA (dsRNA) is the substrate of ADARs, and dsRNA was initially thought to be unique to the genomes of some viruses and/or formed during the viral lifecycle. However, through studies of ADAR cellular targets, it has become clear that metazoan transcriptomes are ripe with dsRNA regions [7, 8]. ADARs bind dsRNA and can change the dsRNA sequence and structure via catalyzing deamination of adenosine (A) to inosine (I), a process commonly referred to as A-to-I RNA editing [9]. Editing of cellular dsRNAs is essential in mammals for both diversification of the nervous system proteome and to prevent the aberrant interaction of cellular transcripts with dsRNA sensors of the innate immune pathway [10]. This later function was uncovered after ADAR mutations were identified in patients suffering from autoimmune disorders [11], and additional studies demonstrated that loss of dsRNA sensors rescues lethality of ADAR mutations in mice [12-14]. Furthermore, as ADARs are conserved in metazoans, studies from several model organisms have explored these relationships and provide data that link ADAR loss with changes in immune gene expression [15-17]. However, studies on the susceptibility of ADAR mutant animals to infection are largely lacking.

In this work, we sought to determine the effect of loss of *Caenorhabditis elegans* ADARs on susceptibility to pathogen infection. The *C. elegans* genome encodes two ADAR family members, ADR-1 and ADR-2 [18]. While both genes contain the canonical ADAR domain structure, ADR-2 is the sole enzyme providing A-to-I editing activity in *C. elegans* [19]. However, loss of *adr-1* impacts both RNA editing and expression of edited genes during development [20, 21]. Recent studies have indicated that combined loss of both *adrs* and small RNA processing factors led to altered upregulation of antiviral genes and developmental defects, including vulva morphology defects and frequent bursting [22, 23]. However, neither study addressed sensitivity or resistance of the mutant animals to infection. Furthermore, the upregulated genes in the animals lacking *adrs* and small RNA processing factors overlapped not only with those regulated by viral infection, but also infection with intracellular pathogens and other general stress responses [23]. In fact, data from many recent studies, particularly in the model organism *C. elegans*, has indicated that innate immune responses are intertwined with different homeostatic mechanisms, such as the unfolded protein response as well as germline integrity [24].

To directly address the physiological role of ADARs in innate immunity, survival of *adr* mutant animals to pathogenic infection was assessed using well-established assays with several bacterial species, all of which are pathogenic to humans. Our data demonstrates that animals lacking ADARs exhibit enhanced susceptibility to pathogenic infection. To our knowledge, this is the first report to demonstrate a role for ADARs in antibacterial immunity. Furthermore, our gene regulatory analysis and scanning electron microscopy studies indicate that *adr* mutant animals have decreased collagen expression and altered cuticle morphology. As employment of physical barriers is also critical to resisting invading pathogens, these data suggest that the role of ADARs in innate immunity may not be limited to altering dsRNA structures to prevent aberrant activation of immune response.

## Results

*Loss of adr-1 or adr-2 increases sensitivity of C. elegans to Pseudomonas aeruginosa* To determine whether *C. elegans* ADR-1 and ADR-2 influence survival to pathogen infection, survival was assessed using a well-established assay with the gram-negative bacterium, *Pseudomonas aeruginosa. P. aeruginosa* is an opportunistic pathogen causing both acute and chronic infection in patients with cystic fibrosis, burn wounds and other diseases requiring ventilation, such as COVID-19 [25]. Similar to humans, *P. aeruginosa* can infect and kill *C. elegans* [26].

Using a standard slow-killing assay, survival of animals lacking *adr-1, adr-2* orboth genes was assessed on plates containing a small lawn of the PA14 clinical isolate of *P. aeruginosa* (Figure 1A). As expected, wildtype animals exposed to *P. aeruginosa* die over the course of several days (Figure 1A). Animals lacking *adr-1* or *adr-2* showed a reproducible and significant sensitivity to killing by *P. aeruginosa* (Figure 1A). Furthermore, animals lacking both *adr-1* and *adr-2* had a similar survival as the animals lacking the individual *adrs* (Figure 1A), suggesting the two *adrs* are functioning together to promote organismal resistance to *P. aeruginosa* infection.

**Figure 1:** ADAR mutant worms are susceptible to *Pseudomonas* infection. A-D) Representative survival curve (of three biological replicates) for the indicated animals subjected to the slow-killing assay and scored for survival in response to *P. aeruginosa* strain PA14. Statistical significance determined using OASIS. *p*<0.001 (***), *p*<0.001 (**) and *p*<0.05 (*).

To rigorously test the impact of ADR-1 and ADR-2, multiple, different deletion alleles were examined. For *adr-1*, survival was assessed for the established *adr-1(gv6)* animals [19] and a newly created CRISPR allele of *adr-1(tcn1)*, which has a complete deletion of the *adr-1* coding sequence (Figure 1B). For *adr-2*, survival was assessed for the established *adr-2(gv42)* [19] and *adr-2(uu28)* [22] animals (Figure 1C). Importantly, these four additional mutant strains all resulted in reproducible and significant sensitivity to killing by *P. aeruginosa* (Figures 1B, C). As a secondary approach, the pathogen susceptibility of an *adr-1(-)* strain carrying a transgene expressing *adr-1* under the control of the *adr-1* promoter was examined (Figure 1D). Re-introduction of *adr-1* into *adr-1(tm668)* animals significantly improved survival to *P. aeruginosa* (Figure 1D).

Transgenic rescue lines are not available for *adr-2*. However, consistent with the requirement for each *adr* in survival to *P. aeruginosa*, the presence of the *adr-1* transgene described above could not rescue the defect of *adr-1(-);adr-2(-)* animals (see Additional file 1: Fig.S1). Together, these data indicate that both ADR-1 and ADR-2 are important and function in the same pathway to promote *C. elegans* survival to *P. aeruginosa* infection. *adr mutant animals exhibit normal avoidance and feeding behavior to P. aeruginosa* Organismal survival to pathogen infection involves both critical gene regulatory programs as well as behavioral responses, such as movement away from the pathogen [27-29]. As *adr* mutant animals have altered chemotactic behavior [19], it is possible that the decreased survival is an indirect effect caused by the inability to sense *P. aeruginosa*. To directly test this possibility, occupancy of wildtype and *adr* mutant animals within the small lawn of *P. aeruginosa* was monitored at five different time-points during the first 30 hours of exposure. Importantly, there is no significant difference in survival of wildtype and *adr* mutant animals during these first hours of exposure (Figure 1A). Consistent with previous studies [30], in the first eight hours of exposure, most wildtype animals do not have a strong preference to avoid *P. aeruginosa*; however, between 12 and 30 hours after exposure, wildtype animals spend more time off the bacterial lawn than within the lawn (Figure 2A). There was no significant difference between the occupancy of wildtype and *adr* mutant animals at any point during the assay (Figure 2A). These data suggest that decreased survival of *adr* mutant animals exposed to *P. aeruginosa* is not caused by an inability to avoid pathogen.

**Figure 2:**
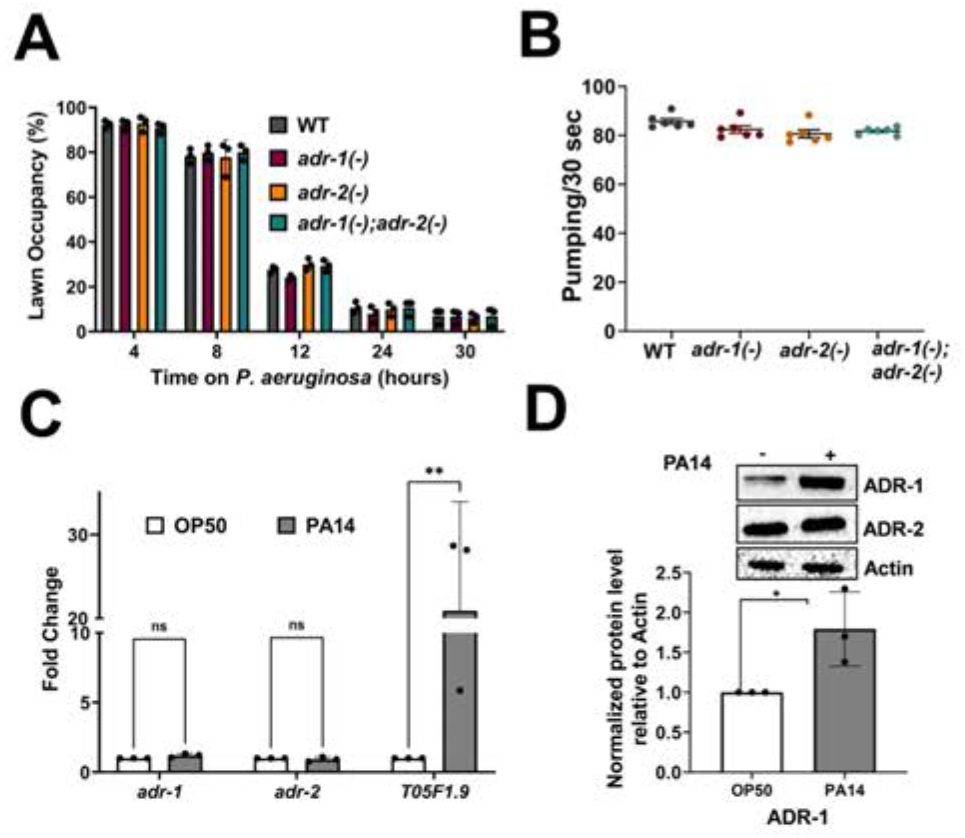
*Adr* mutant animals exhibit normal avoidance and feeding behavior to *P. aeruginosa*. A) Lawn occupancy percentage was calculated by counting the number of worms of the indicated strains in the lawn and outside the lawn, which was summed and then divided by the total number of worms. Each data point represents the average of three technical replicates performed at the indicated time and all experiments were performed in three biological replicates. Error bars represent the standard error of the mean (SEM). Statistical significance determined using two-way ANOVA Tukey’s multiple comparisons test. *p* value of the mutants relative to WT was not significant (*p*>0.05) for all timepoints. B) Each dot in the graph represents the average pumping rate for three technical replicates obtained in two independent experiments. Error bars represent SEM. Statistical significance determined using unpaired Mann Whitney test, with no significant differences observed (*p*>0.05). C) qRT-PCR quantification of the level of the indicated genes relative to *gpd-3* and normalized to the ratios obtained for OP50. The mean of three biological replicates was plotted. Error bars represent SEM. Statistical significance determined using a two-way ANOVA Sidak’s multiple comparisons test. \**p*≤0.05, \*\**p*≤0.005, ns indicates no significant difference (*p*>0.05). D) Equivalent amounts of lysate from animals with V5 and 3xFLAG epitope tags on ADR-1 and ADR-2, respectively, exposed to *P. aeruginosa* (PA14) (+) or the control *E. coli* (OP50) (-) for seven hours were subjected to SDS-PAGE and immunoblotting with V5 (ADR-1), FLAG (ADR-2) and Actin antibodies.

It is also possible that *adr* mutant animals are more susceptible due to increased *P. aeruginosa* intake. To monitor intake, pharyngeal pumping was measured for animals after twenty-four hours of exposure to *P. aeruginosa*. Pumping rates observed for wildtype animals were similar to those previously reported [31], and *adr* mutant animals did not have significantly different pharyngeal pumping rates when compared to wildtype animals (Figure 2B). This suggests a similar level of pathogen intake in all the strains and that the enhanced susceptibility of the *adr* mutant worms is likely not due to more intake of *P. aeruginosa*.

As *adr* mutant animals did not appear to have defects in pathogen avoidance or intake, it is possible that the susceptibility arises because, in wildtype animals, ADR-1 and ADR-2 are critical effectors that increase expression upon pathogen exposure, a feature lost in *adr* mutant animals. To examine this possibility, ADAR protein and mRNA levels were analyzed in response to *P. aeruginosa* infection. As activation of immune response genes occurs within four to eight hours after exposure to *P. aeruginosa* [32, 33], wildtype animals were subjected to a seven-hour exposure followed by RNA and protein isolation. To facilitate detection of protein levels, wildtype animals were CRISPR modified to express a V5 epitope at the N-terminus of ADR-1 and three copies of the FLAG epitope at the N-terminus of ADR-2. The epitope tags did not affect known behavioral consequences caused by lack of *adr* function or RNA editing (see Additional file 2: Fig. S2 A,B). To confirm activation of the immune response, expression of *T05F1*.*9*, a gene previously shown to be upregulated by *P. aeruginosa* exposure [33] was analyzed by quantitative real-time PCR (qRT-PCR) in three biological replicates of RNA isolated from animals exposed to *P. aeruginosa* and compared to RNA isolated from the same animals grown on plates with the standard *C. elegans* bacterial food source, *E. coli* strain OP50. In contrast to *T05F1*.*9*, both *adr-1* and *adr-2* mRNA levels did not change upon exposure to *P. aeruginosa* (Figure 2C). Consistent with the mRNA levels, ADR-2 expression did not change upon *P. aeruginosa* exposure (Figure 2D). In contrast, ADR-1 protein expression significantly increased upon *P. aeruginosa* exposure (Figure 2D), suggesting that ADR-1 expression may be post-transcriptionally controlled when animals encounter a bacterial pathogen.

### ADR-1 RNA binding is required for survival to P. aeruginosa infection

The upregulation of ADR-1 after *P. aeruginosa* exposure suggests ADR-1 may play a role in response to infection. While lacking deaminase activity [19], ADR-1 does possess double-stranded RNA (dsRNA) binding activity [34]. The impacts of loss of ADR-1 RNA binding on gene expression are not known. However, RNA binding by ADR-1 is known to both positively and negatively regulate ADR-2-mediated RNA editing, depending on the tissue, developmental timing and specific transcript [21, 35]. To investigate the role of ADR-1 RNA binding in survival of animals exposed to *P. aeruginosa*, the survival assay was performed with *adr-1(-)* animals containing an extrachromosomal array expressing an ADR-1 dsRBD1 mutant under the control of the *adr-1* promoter. The ADR-1 dsRBD1 mutant has EAxxA (E = glutamic acid, A = alanine and x = any amino acid) present in place of the conserved KKxxK (K = lysine) motif and previous studies have indicated that this ADR-1 dsRBD1 mutant lacks the ability to bind known ADR-1 mRNA targets *in vivo* [34]. Consistent with our earlier results (Figure 1A), survival of *adr-1(-)* animals exposed to *P. aeruginosa* infection was significantly shorter than wildtype animals (Figure 3A). However, in contrast to the ability of transgenic wildtype *adr-1* to restore survival to *adr-1(-)* animals (Figure 1D), survival of the ADR-1 dsRBD1 mutant animals was not significantly different from *adr-1(-)* animals (Figure 3A). These data suggest that ADR-1 binding to mRNA is important for survival to *P. aeruginosa* infection.

**Figure 3:**
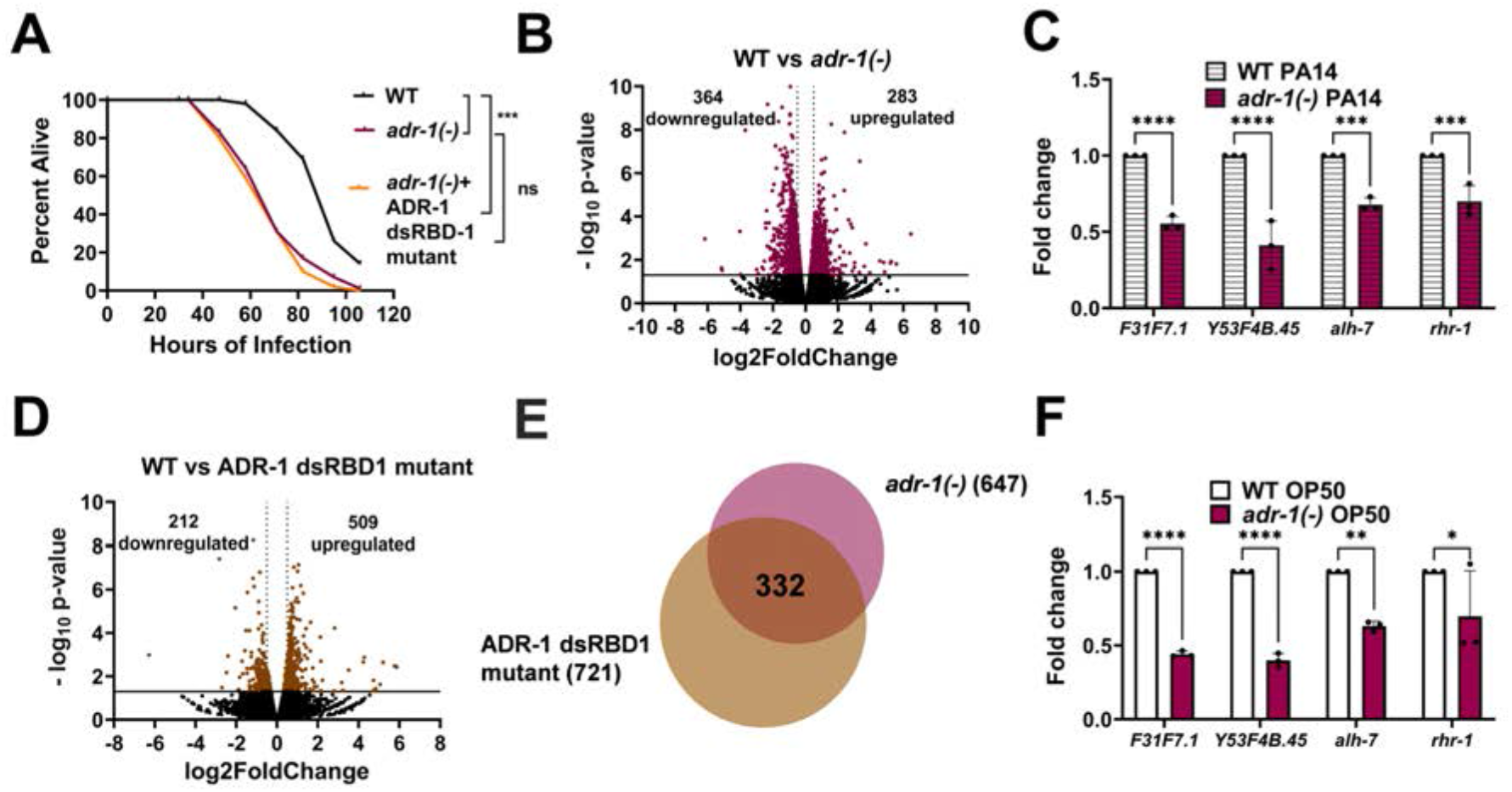
ADR-1 RNA binding is required for survival to *P. aeruginosa*. A) Representative survival curve (of three biological replicates) for the indicated animals subjected to the slow-killing assay and scored for survival in response to *P. aeruginosa* strain PA14. Statistical significance determined using OASIS, where *p*<0.001 (***), and ns indicates *p*>0.05. B and D) Dots represent individual genes that are down- or upregulated or not significantly differentially expressed (black) in RNA-seq data from WT animals compared to *adr-1(-)* (maroon) (B) or ADR-1 dsRBD1 mutant animals (brown). (D) Average log_2_ fold change (x-axis) is plotted against the negative log_10_ *p* values (y-axis). Genes considered significantly differentially expressed exhibited log_2_ fold change of |0.5| (light gray dotted vertical lines) and *p*<0.05 [log_10_ *p-*value of 1.3, solid black horizontal line]. C and F) qRT-PCR quantification of the level of the indicated genes relative to *gpd-3* and normalized to the ratios obtained for WT PA14 and WT OP50 in (C) and (F), respectively. The mean of three biological replicates was plotted. Error bars represent SEM. Statistical significance was determined using a two-way ANOVA Sidak’s multiple comparisons test. \**p*≤0.05, \*\**p*≤0.005, \*\*\**p*≤0.0005, \*\*\*\**p*≤0.0001, ns indicates no significant difference (*p* >0.05). E) Overlap of genes misregulated in (B) and (D).

To further investigate the role of ADR-1 in survival to *P. aeruginosa* infection, we sought to identify transcripts regulated by ADR-1 mRNA binding in response to *P. aeruginosa* infection. To this end, high-throughput sequencing was performed on polyadenylated RNA isolated from wildtype, *adr-1(-)* and the ADR-1 dsRBD1 mutant animals after seven hours of exposure to *P. aeruginosa*. When compared to wildtype animals, differential gene expression analysis of two biological replicates revealed 647 significantly differentially expressed genes (*p*<0.05, log_2_fold change > |0.5|) in the *adr-1(-)* RNA-sequencing (RNA-seq) data (Figure 3B, see Additional file 3: Table S3). Among the differentially expressed genes, there were 283 up- and 364 downregulated genes (Figure 3B). To independently validate the RNA-seq findings, four genes identified as differentially expressed were randomly chosen and analyzed by qRT-PCR in three independent biological replicates. Consistent with the RNA-seq data, all four genes (*F31F7*.*1, Y53F4B*.*45, alh-7* and *rhr-1*) were significantly downregulated in RNA isolated from *adr-1(-)* animals exposed to *P. aeruginosa* when compared to RNA isolated from wildtype animals exposed to *P. aeruginosa* (Figure 3C).

To determine how many differentially expressed genes are directly regulated by ADR-1 binding, the wildtype and ADR-1 dsRBD1 mutant RNA-seq datasets were compared and overlapped with the genes misregulated in the absence of *adr-1*. Differential gene expression analysis revealed 721 significantly differentially expressed (*p*<0.05, log_2_fold change > |0.5|) genes between the ADR-1 dsRBD1 mutant RNA-seq data and the wildtype RNA-seq data (Figure 3D, see Additional file 4: Table S4). Importantly, nearly half of these misregulated genes (332/721) are observed in our datasets of differentially regulated genes from *adr-1(-)* animals (Figure 3E, see Additional file 4: Table S4), suggesting that loss of ADR-1 binding to mRNA plays a major role in ADR-1-mediated control of gene expression. While human ADARs have been shown to have editing-independent, RNA binding-dependent gene regulatory functions on a handful of genes [36], our high-throughput sequencing analysis provides the first direct evidence that RNA binding by an ADAR family member significantly contributes to altered mRNA expression.

To assess the contribution of the newly identified genes controlled by ADR-1 RNA binding to the pathogen susceptibility phenotype, these targets were compared to known *C. elegans* pathogen response genes. Surprisingly, very few (21/332) of the ADR-1 regulated genes were also previously shown to be upregulated in wildtype worms exposed to *P. aeruginosa* [33] (see Additional file 4: Table S4). Furthermore, using the *C. elegans*-specific software, WormCat [37], an unbiased search of enriched gene sets was performed for the 332 transcripts regulated by loss of *adr-1* and loss of ADR-1 RNA binding and did not yield any categories related to immune function or pathogen response (data not shown). These data suggest that while ADR-1 RNA binding may be important for survival to *P. aeruginosa* infection, genes regulated by ADR-1 may be altered even prior to infection. Consistent with this, using qRT-PCR and comparing to RNA isolated from wildtype animals grown under the same feeding conditions, all four genes downregulated in RNA isolated from *adr-1(-)* animals exposed to *P. aeruginosa* (Figure 3C) were also significantly downregulated in *adr-1(-)* animals feeding on OP50 (Figure 3F). In sum, these data indicate that RNA binding by ADR-1 regulates hundreds of genes during infection, which may be interesting for future studies to understand the importance of ADR-1 upregulation during infection. However, these data also indicate that these genes, and perhaps others that are regulated by both ADR-1 and ADR-2 and are important for organismal survival to infection, are misregulated prior to *P. aeruginosa* exposure.

### Worms lacking adrs exhibit decreased collagen expression

To take an unbiased approach to understanding the role of ADARs in regulating basal expression of genes important for survival to *P. aeruginosa* infection, transcriptome-wide RNA-sequencing was performed on RNA isolated from wildtype, *adr-1(-), adr-2(-)* and *adr-1(-);adr-2(-)* animals that were grown similar to the slow-killing assay, but exposed to only OP50 at 25°C for seven hours. Polyadenylated RNA was isolated from three biological replicates of each genotype and subjected to high-throughput sequencing. Differential gene expression changes were analyzed in the wildtype RNA-seq dataset compared to the RNA-seq datasets of *adr-1(-), adr-2(-)* single mutant and the *adr-1(-);adr-2(-)* double mutant animals. The *adr-1(-)* and *adr-1(-);adr-2(-)* RNA-seq datasets had the largest number of significantly differentially expressed genes (*p*<0.05, log_2_fold change > |0.5|) with over 1800 (Figure 4A, see Additional file 5: Table S5) and nearly 1500 (Figure 4B, see Additional file 6: Table S6) misregulated genes identified, respectively, whereas approximately 350 differentially expressed genes were identified in the *adr-2(-)* RNA-seq dataset (Figure 4C, see Additional file 7: Table S7). It is unclear why the RNA from the *adr-2(-)* animals exhibited less overall gene expression changes but does suggest that more genes may be affected by loss of *adr-1* than the complete loss of editing, which is consistent with our previous developmental assessment of ADR-1 and ADR-2 function [20].

**Figure 4:**
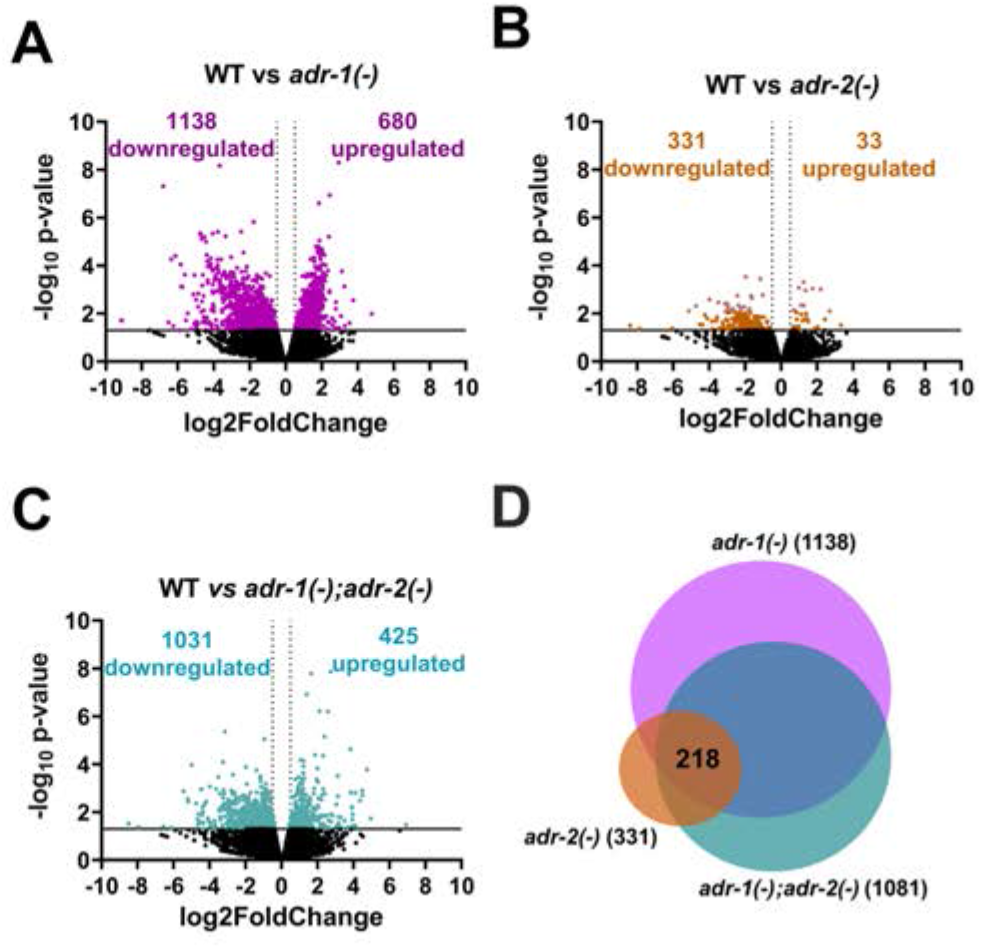
Altered gene expression in *adr* mutant animals. A-C) Dots represent individual genes down- or upregulated or not significantly differentially expressed (black) in RNA-seq data from WT animals compared to *adr-1(-)* (purple) (A), *adr-1(-);adr-2(-)* (green) (B) and *adr-2(-)* (yellow) grown on OP50. Average log_2_ fold change (x-axis) is plotted against the negative log_10_ *p* values (y-axis). Genes considered significantly differentially expressed exhibited log_2_ fold change of |0.5| (light gray dotted vertical lines) and *p* ≤ 0.05 [log_10_ *p*-value of 1.3, solid black horizontal line]. D) Overlap of genes downregulated in (A), (B) and (C).

To identify genes that might underlie the *adr* mutant animals’ enhanced susceptibility to *P. aeruginosa* infection, transcripts that were commonly misregulated across all three RNA-seq datasets were identified. Overlap of the upregulated transcripts in the *adr-1(-)* (680), *adr-2(-)* (33) and *adr-1(-);adr-2(-)* (425) RNA-seq datasets revealed only 4 commonly upregulated genes (data not shown). However, overlap of the downregulated transcripts in the *adr-1(-)* (1138), *adr-2(-)* (331) and *adr-1(-);adr-2(-)* (1081) RNA-seq datasets revealed nearly 220 commonly downregulated genes (Figure 4D, see Additional file 8: Table S8). Using the WormCat software, gene set enrichment analysis revealed only one significantly enriched category - extracellular material (*p* value=2.8*10^−07^). Further classification (category 2 output) of this enriched category revealed that 15 of the 17 misregulated genes associated with the extracellular material category were members of the collagen gene family. Collagens are the major component of cuticle which is the outer surface of *C. elegans* and acts as a barrier between the animal and the environment [38]. Collagen expression was observed to be altered in *C. elegans* during recovery from acute *P. aeruginosa* infection [39], and early genetic studies demonstrated that loss of the cuticular collagen gene, *col-179*, led to enhanced susceptibility to *P. aeruginosa* infection [32]. Interestingly, *col-179* is one of the collagen genes in the downregulated transcripts present in our *adr-1(-), adr-2(-)* and *adr-1(-);adr-2(-)* RNA-seq datasets from animals fed typical bacteria (*E. coli* OP50) (see Additional files 5-7). To independently validate the changes in collagen gene expression, RNA was isolated from three independent biological replicates of the *adr* mutant strains grown as in the slow-killing assay but exposed to only OP50 and qRT-PCR was performed for three collagen genes, *col-179, col-106* and *col-135*. Consistent with the RNA-seq results, loss of either *adr-1* or *adr-2* or loss of both *adrs* resulted in a significantly decreased expression of the collagens when grown on OP50 (see Additional file 9: Fig. S9 A,B,C). To further explore the altered common *adr*-regulated genes, a second, independent analysis was performed using the extracellular specific software, Matrisome Annotator [40], which indicated that 14 of the 15 collagen genes were in fact cuticular collagens (data not shown). Together, these data indicate that *adr* mutant animals have altered collagen expression and suggest that these molecular defects may impact cuticle structure and pathogen susceptibility.

### Worms lacking adrs exhibit altered cuticle structure and survival to several bacterial species

As the molecular data suggests that *adr* mutant animals have decreased expression of collagen genes, the cuticles of wildtype and *adr-1(-);adr-2(-)* animals were analyzed by scanning electron microscopy (SEM). Both strains of animals were grown as in the slow-killing assay and then fed either *E. coli* (OP50) or *P. aeruginosa* (PA14) for seven hours. The cuticle of wildtype animals changed from smooth to wrinkled after *P. aeruginosa* exposure (Figure 5A). Wrinkled cuticles are associated with the presence of a thinner hypodermis and/or alterations in the connections between the cuticle and hypodermis in aging animals [41]. This observation suggests that the cuticle structure changes in response to pathogen infection and has been previously observed in other SEM studies of wildtype *C. elegans* exposed to *P. aeruginosa* [42]. Interestingly, the cuticle of *adr-1(-);adr-2(-)* animals fed on *E. coli* appear to be more wrinkled compared to wildtype animals of same age and exposure conditions (Figure 5A). The cuticle of *adr-1(-);adr-2(-)* animals exposed to *P. aeruginosa* was further wrinkled (Figure 5A). However, the difference between the cuticles of wildtype and *adr-1(-);adr-2(-)* animals was less drastic in the *P. aeruginosa* exposure compared to the *E. coli* (OP50) exposure (Figure 5A). Together, these data suggest that ADARs regulate collagen levels, which in turn impacts cuticle structure and the ability of the animal to defend against pathogens.

**Figure 5:**
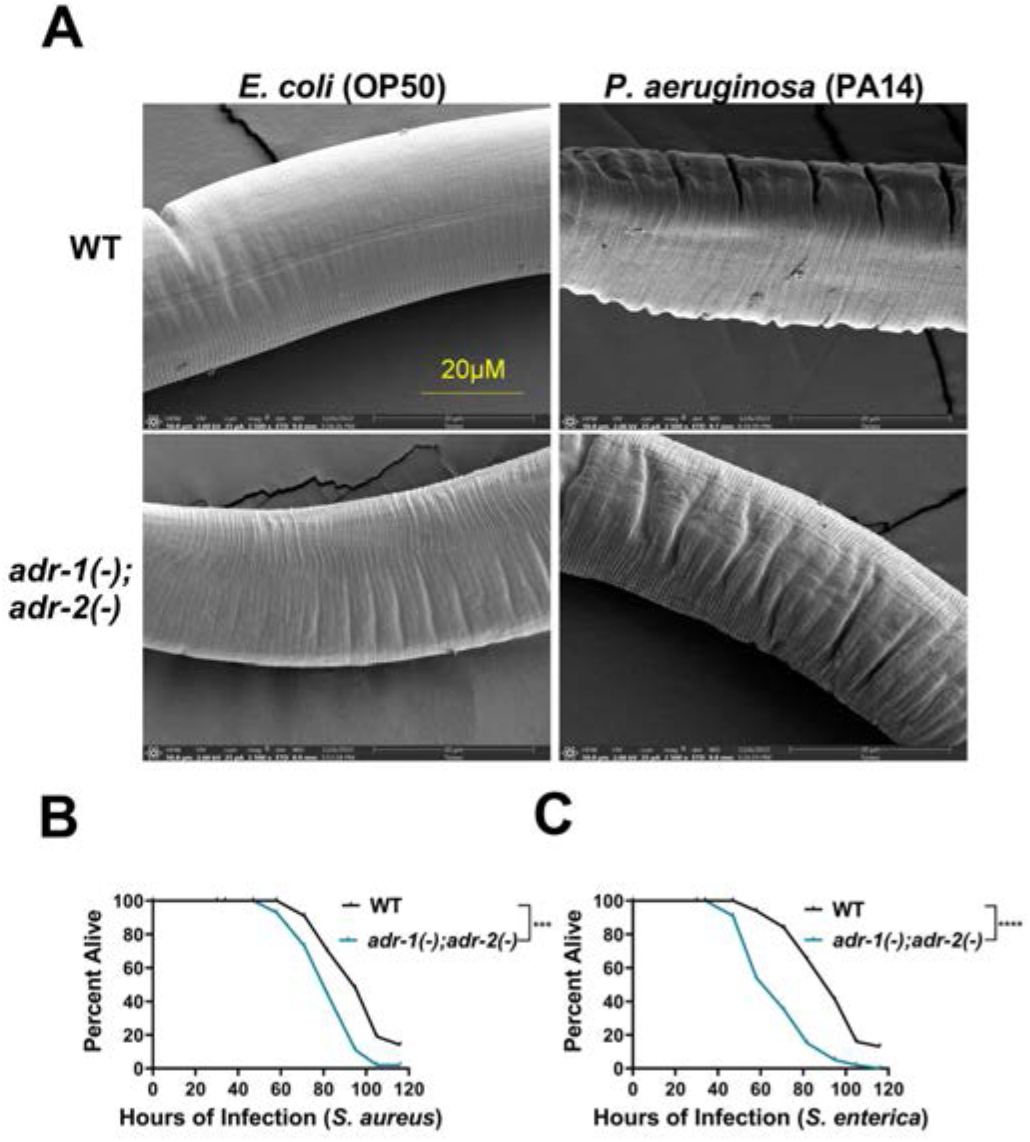
Loss of *adrs* results in altered cuticle structure and enhanced susceptibility to several pathogenic bacterial species. A) Representative (from two biological replicates, 5-10 images per replicate) SEM images of the indicated strains after exposure to the bacterial strains indicated. (B, C) Representative survival curve (of three biological replicates) for the indicated animals subjected to the slow-killing assay and scored for survival in response to *S. aureus* (B) and *S. enterica* (C). Statistical significance was determined using OASIS, \*\*\*\**p*<0.0001, \*\*\**p*<0.001.

As other mutant animals with altered cuticles have altered survival to a range of pathogens [42], we sought to examine the survival of *adr* mutant animals to two additional paradigmatic human pathogens: *Salmonella enterica* and *Staphylococcus aureus*. The gram-negative bacteria *S. enterica* can proliferate and establish infection in *C. elegans* [43, 44]. The gram-positive bacteria *S. aureus* has also previously been shown to both infect and kill *C. elegans* [45]. The standard slow-killing assay was performed with wildtype and *adr-1(*-);*adr-2(-)* mutant animals on small lawns of *S. enterica* (Figure 5B) and *S. aureus* (Figure 5C). As expected, wildtype animals exposed to either *S. enterica* or *S. aureus* die over the course of several days (Figures 5B,C). Survival of animals lacking both *adrs* was significantly shorter than wildtype animals when grown on either *S. enterica* (Figure 5B) or *S. aureus* (Figure 5C). Together, these data indicate the reproducible sensitivity of *adr* mutant animals to opportunistic human pathogens. Collectively, these data indicate that *C. elegans* ADARs can play important gene regulatory roles to contribute to the formation of physical barriers critical for promoting organismal survival to pathogen infection. Future research aimed at determining the susceptibility of ADAR mutants in other model systems as well as further mechanistic studies of how ADARs regulate collagen expression and the specific cuticular collagens that are key to organismal defense to infection are critical to improving our understanding of the complex relationship of ADARs and innate immunity.

## Discussion

In this study, we examined the contribution of *C. elegans* ADARs to survival from infection by opportunistic human pathogens. Specifically, we found that *adr* mutant animals are susceptible to both gram-negative (*Pseudomonas aeruginosa* and *Salmonella enterica*) and gram-positive (*Staphylococcus aureus*) bacteria. Using a combination of high-throughput sequencing, microscopy and functional genetics, we determined that ADR-1 and ADR-2 function together to regulate collagen expression, and the absence of these RNA binding proteins results in altered cuticle structure, which in turn may render these animals more susceptible to infection.

At present, it is unknown how ADR-1 and ADR-2 regulate collagen expression. The ADAR family of RBPs can regulate gene expression in both editing-dependent and independent manners [46]. Our data indicates a role for ADR-1 RNA binding in regulating survival to pathogen infection (Figure 3A) but does not eliminate the possibility of editing-dependent regulation, as ADR-1 binding to RNA has previously been shown to both promote and inhibit A-to-I editing by ADR-2 [21, 34]. Loss of *adr-1* leads to milder effects on editing compared to animals lacking *adr-2* or those lacking *adr-1* and *adr-2*, both of which completely lack editing. If survival to pathogen exposure was editing-dependent, the *adr-1(-)* animals would have an intermediate phenotype, similar to that observed for the chemotaxis defects of *adr* mutant animals [19]. Furthermore, we did not observe A-to-I editing events of any of the misregulated collagen genes in our study (data not shown). Interestingly, defects in RNA modification have been previously connected to altered collagen expression in *C. elegans* [47]. In the previous study, loss of methylation of cytosine (5mC) on ribosomal RNA (rRNA) resulted in decreased translation of cuticular collagen genes.

It is possible that ADARs are regulating collagen expression by directly binding to each of the misregulated collagen genes. In fact, two of the collagen genes, *col-179* and *col-106*, that exhibit decreased expression in the *adr* mutant animals were previously identified as ADR-1 bound mRNAs [20]. In addition, we also observed decreased expression of *col-179* and *col-106* in ADR-1 dsRBD1 mutant animals after exposure to

*P. aeruginosa* (Additional file 4, Table S4). However, previous studies did not observe ADR-2 binding to these transcripts [34]. An alternative possibility is that ADR-1 and ADR-2 impact signaling pathways that control collagen expression, including potentially binding to and directly regulating expression of key transcription factors and kinases in these pathways. In particular, some of the cuticular collagens misexpressed upon loss of *C. elegans adrs* are regulated by the TGF-β (6/15 genes overlap) and/or insulin signaling (4/15 genes overlap) pathways [48, 49]. We have not observed misregulation of any of the canonical TGF-β pathway genes (*daf-1, daf-4, daf-8, daf-14, daf-3, daf-5*) in *adr* mutant animals in this study (Additional files 5-7, Tables S5-7) or other tissue-specific studies [21]. However, for the insulin signaling pathway, we previously observed that the transcription factor which controls collagen expression, SKN-1, has reduced mRNA expression in the nervous system of *adr-2(-)* animals [21]. We did not observe altered *skn-1* expression in the RNA-seq analysis of young adult *adr* mutant animals presented in this work (Additional files 5-7, Tables S5-7). Future work should explore whether changes in collagen-regulating pathways, such as those driven by SKN-1, are misregulated in the nervous system of *adr* mutant animals and whether this could contribute to altered cuticle structure. Recently, it was demonstrated that a neural G-protein coupled receptor, NPR-8, dynamically regulates collagen expression and cuticle structure in response to temperature changes and infection [42, 50]. Furthermore, it has been proposed that although the epidermis plays a major role in synthesizing the cuticle, neurons can sense both the environment and tension to influence collagen dynamics [51].

It is also important to note that, in our study, animals were grown at 25ºC prior to isolating RNA for high-throughput sequencing or SEM imaging, which could influence cuticular structure. Previous studies have shown that the primary transcriptional regulator of cellular response to elevated temperature, HSF-1, is a major regulator of collagen gene expression both in the presence and absence of heat shock [52]. In total, comparing genes misregulated in *adr* mutant animals (this study) and animals lacking *hsf-1* [52], we observed 10 of the 15 collagen genes were commonly misregulated. Similar to *skn-1*, we do not observe altered *hsf-1* mRNA expression in the RNA isolated from *adr* mutant animals, but *hsf-1* expression was previously observed to be downregulated in the nervous system of *adr-2(-)* animals [21]. Future experiments should aim to dissect how temperature differentially impacts the cuticular structure of *adr* mutant animals and if HSF-1 is important for regulation of collagen gene expression by ADARs.

In addition to ADR-1 and ADR-2 functioning together to regulate collagen gene expression and organismal resistance to pathogen infection, our study revealed hundreds of transcripts that are regulated by ADR-1 binding upon exposure to pathogen. Interestingly, we also see an increase in ADR-1 protein expression upon pathogen exposure, which raises the possibility that ADR-1 could be binding to new targets in response to infection and potentially has additional functions beyond promoting survival to infection. Recently, roles for RNA binding proteins and small RNAs in promoting pathogenic memory and transgenerational inheritance have been identified [53, 54]. Future studies should explore changes in ADR-1 binding targets in response to infection and their effects on immunological memory.

## Conclusion

This study revealed a critical role of the *C. elegans* ADAR family of RNA binding proteins in promoting cuticular collagen expression and defense from pathogenic microbes. Previous studies of this RNA binding protein family have suggested a role in the antiviral response, but our data indicate a broader function of ADARs in innate immunity. This work sets the stage for future studies aimed at mechanistic dissection of how ADARs control collagen expression and the tissue-specific roles these proteins play in innate immunity. In addition, our study provides a list of targets regulated by ADR-1 RNA binding which could be critical for future research on ADAR function in immunity and development.

## Methods

### Worm Strains and Maintenance

All worms were maintained under standard laboratory conditions on nematode growth media (NGM) seeded with *Escherichia coli* OP50. Worm strains used in this study and previously published are wildtype (N2), BB19 *adr-1(tm668)*, BB20 *adr-2(ok735)*, BB21 *adr-1(tm668);adr-2(ok735)* [55], BB2 *adr-1(gv6)*, BB3 *adr-2(gv42)* [19], BB19 *adr-1(tm668)* + blmEx1[3XFLAG-*adr-1 genomic, rab3::gfp::unc-54*]) BB21 *adr-1(tm668);adr-2(ok735)* + blmEx1[3xFLAG-*adr-1* genomic, *rab3::gfp::unc-54*]) [35]. BB21 *adr-1(tm668)* + *blmEx11*[*3XFLAG-adr-1* genomic with mutations in *dsRBD1 (K223E, K224A, and K227A*), *rab3::gfp::unc-54 (3*′ *UTR)*] [34]. Additional strains used in this study were *adr-2(uu28)* (a kind gift from Brenda Bass) and the newly generated ALM63 *adr-1(tcn1)* strain and the HAH36 V5-ADR-1; 3xFLAG-ADR-2 strain, which were created by CRISPR using the large deletion protocol [56] and [57], respectively. Guides and repair templates are listed in Additional file 10 (IDT). For ALM63, the injection mix contained 25 μM KCl, 7.5 mM HEPES, pH 7.4, 4.9 μM Cas9 (Invitrogen, TrueCut), 5 μM tracrRNA (IDT), 2 μM *dpy-10* crRNA, 25 μM each of two crRNAs to *adr-1*, 2 μM *dpy-10* single-stranded oligo nucleotide (ssODN) repair sequence and 5 μM of a target ssODN to *adr-1*. To avoid compounding effects from off-target mutations, the generated ALM63 strain was crossed twice with the wildtype strain. For HAH36, the V5 epitope at the N-terminus of ADR-1 and 3 copies of the FLAG epitope at the N-terminus of ADR-2 were constructed in wildtype worms individually, back-crossed to wildtype worms and then crossed to generate HAH36. Injection mix for the V5-ADR-1 and 3xFLAG-ADR-2 strains included 1.5 μM Cas9 (IDT, Alt-R Cas9 nucleas V3), 4 μM tracrRNA (IDT), 4 μM of crRNA (IDT), 37 ng/μl *rol-6* plasmid (HAH293) and 1 μM target ssODN (Additional file 10: Table S10). Genomic modifications were verified using PCR (primers listed in Additional file 10: Table S10) and Sanger sequencing. Western blotting was also performed to verify the V5 and 3xFLAG insertions.

### Pathogenic bacterial growth

Three pathogenic bacterial strains were used: *P. aeruginosa* PA14 (a kind gift of Read Pukkila-Worley), *S*.*enterica* SL1344 and *S. aureus* MSSA476 from (kind gifts of Jingru Sun, Washington State University). Bacterial strains were freshly streaked on LB plates and grown as 5 ml cultures at 37°C overnight. The next day, 20 μl of culture was spotted onto 6 cm NGM agar plates. Plates were incubated at 37°C overnight (not exceeding 16 hours) and then moved to 25°C for at least 24 hours before starting the slow-killing assay.

### Slow-killing assay

Slow-killing assays were performed as previously described [26, 42] with slight modifications. For each assay, 45 worms of each strain were plated on each of three NGM plates containing 0.05 mg/ml 5-Fluoro-2’-deoxyuridine (MP Biomedical) spotted with 20 μl of a given bacterial strain (grown as described above). Plates were incubated at 25°C and after 24 hours, animals were scored as dead or alive at least once every 11-13 hours over the course of 120 hours.

### P. aeruginosa exposure assay for gene expression

Gravid adult worms were collected in 1x M9 buffer (3 g KH_2_PO_4_, 6 g Na_2_HPO_4_, 5 g NaCl, 1 ml 1 M MgSO_4_, H_2_O to 1 L) and incubated with 0.5 M NaOH in 1.2% NaClO (Fisher) to release eggs. Eggs were washed thoroughly with 1x M9 buffer and hatched overnight at 20°C to obtain synchronized first larval stage (L1) animals. Hatched L1 animals were washed with 1x M9 and grown at 20°C on standard NGM plates with OP50 for 42 hours. For exposures of each strain, three 10 cm NGM plates were seeded with 40 μl of OP50 or PA14. After exposure for 7 hours at 25°C, worms were washed with 1x M9 buffer and collected in TRIzol (Invitrogen).

### Pharyngeal pumping rate assay

These experiments were performed as previously described [58]. Briefly, 6cm NGM plates were seeded with 30 μl of an overnight culture of *P. aeruginosa* (PA14) and incubated as described earlier. Fifteen synchronized young adult worms were transferred to the seeded plates and incubated 24 hours at 25°C. Individual worms were tracked under Carl Zeiss Stemi 305 microscope, and the number of contractions of the pharyngeal bulb was counted over thirty seconds.

### RNA isolation and quantitative real-time PCR (qRT-PCR)

Total RNA was isolated using TRIzol (Invitrogen) and purified using TURBO DNase (Ambion) followed by the RNeasy Extraction kit (Qiagen). For qRT-PCR, 2 μg of DNase-treated RNA was subjected to cDNA synthesis using Superscript III (Invitrogen) reverse transcriptase and a combination of both random hexamers and oligo dT primers (Fisher Scientific). After reverse transcription, 20 μl of water was added to each cDNA sample. Gene expression was determined by qRT-PCR using SybrFast Master Mix (KAPA) and gene-specific primers using a Thermofisher Quantstudio 3 instrument. Primers for qRT-PCR (see Additional file 10) were designed to span an exon-exon boundary. For each gene analyzed, a standard curve was generated using ten-fold serial dilutions of the amplicon to test the relative concentration versus the threshold of amplification. Standard curves were plotted on a logarithmic scale in relation to concentration and fit of the line (r^2^ value). The r^2^ value was typically 0.99, and all data points fell within the standard curve. For each sample, cDNA concentration was measured in triplicate and three biological replicates were performed for each experiment.

### Western analysis

Synchronized L4 animals after exposure to either *P. aeruginosa* (PA14) or *E. coli* (OP50) at 25°C for seven hours were collected in 1x M9 buffer and washed 3 times. Collected animals were rocked for 20 minutes at room temperature. After a brief centrifugation step, the animals were pelleted, resuspended in 1x SDS buffer (2% SDS, 50 mM Tris HCl and 10% Glycerol) and snap-frozen using liquid nitrogen. Lysates were prepared by boiling the pellet for 15 minutes and vortexing every 7-8 minutes. Protein concentration was measured using Bradford reagent (Sigma) and then 100 mM DTT and bromophenol blue (0.1%) were added to the lysate before boiling for 5 minutes. An equivalent amount of protein lysate was subjected to SDS-PAGE and immunoblotting with antibodies against FLAG (Sigma), V5 (Cell Signaling) and β-actin (Cell Signaling).

### Library preparation, RNA sequencing and analysis

DNase-treated RNA was incubated with oligo(dT) beads (Invitrogen) and followed by library preparation using a stranded RNA-seq library preparation kit (KAPA) as per the manufacturer’s instructions. Libraries were sequenced for SE75 cycles on an Illumina NextSeq 500 instrument at the Center for Genomics and Bioinformatics at Indiana University. Single stranded sequencing reads were aligned to *C. elegans* genome (WS275) using STAR (v2.7.6a) using the following parameters: outFilterMultimapNmax 1 \ outFilterScoreMinOverLread .66 \ outFilterMismatchNmax 10 \ outFilterMismatchNoverLmax .3. Uniquely mapped reads were then used as an input for running featureCounts (v2.0.1). The raw read counts obtained from featureCounts were used for differential gene expression analysis using DeSeq2. R studio version 4.1.1 was used to install Bioconductor package (v3.10) to load DeSeq2.

### Scanning Electron Microscopy

Scanning Electron Microscopy was performed as previously described [42]. Briefly, synchronized young adult animals were exposed to *P. aeruginosa* (PA14) or *E. coli* (OP50) for seven hours at 25°C. Animals were removed from plates with 1x M9 buffer, washed five times wherein the animals were allowed to settle by gravity. After washing, genotypes and exposures were blinded. Animals were incubated overnight in fixation buffer (2.5% glutaraldehyde, 1.0% paraformaldehyde, and 0.1 M sodium phosphate (Electron Microscopy Sciences)) at 4°C. Samples were then washed with 0.1 M sodium phosphate, and the sample suspension was placed in BEEM capsules (size 00) (Ted Pella, Inc.). Samples were dehydrated at room temperature in a graded series of ethanol (30%, 50%, 75%, 90%, 95%, and 100%) with incubation for 10 minutes at each step. Dried samples were placed in aluminum SEM stubs (Electron Microscopy Sciences); which were sputter coated at 45 nm with a Safematic CCU-010. The sputter coated target was composed of gold:palladium (80:20). SEM imaging was done using a ThermoFisher Teneo instrument set to 2.0 kV.

## List of Abbreviations

ADAR: adenosine deaminase that act on RNA
dsRBD1: double stranded RNA Binding Domain 1
qRT-PCR: quantitative Real-Time PCR
CRISPR: Clustered regularly interspaced short palindromic repeats
STAR: Spliced Transcripts Alignment to a Reference
GEO: Gene Expression Omnibus
BEEM: Ballistic Electron Emission Microscopy
OASIS: Online Application for Survival Analysis
rRNA: ribosomal RNA

## Declarations

### Availability of data and materials

All bacterial and nematode strains can be provided upon request. Raw high-throughput sequencing reads have been uploaded to GEO under GSE223919.

### Author Contributions

Designed the experiments: A.D., C.S., H.A.H.

Created reagents: Y.A., C.S., H.A.H.

Performed the experiments in Figure 1 (A.D., C.S.), Additional file 1 (A.D.), Figure 2 (A.D., C.S.), Additional file 2 (C.S., M.S.), Figure 3 (A.D.), Figure 4 (A.D., C.S.), Additional file 9 (A.D., C.S.), Figure 5 (A.D., C.S.)

Performed the bioinformatics analysis: H.A.H and A.D.

Wrote the manuscript: H.A.H., A.D., C.S.

Edited the manuscript: H.A.H., A.T.L., A.D., C.S.

## Acknowledgements

We would like to thank current members of the Hundley lab, Ananya Mahapatra, Boyoon Yang, Emily Erdmann and Priyanka Mukherjee, for careful reading of the manuscript. We would like to thank former lab members Dr. Sarah Deffit for providing the initial data about pathogen exposure that led to this work, Pranathi Vadlamani for developing the pathogen assay protocol, and Halle Stump for providing the initial data collection. We thank Dr. Barry Stein at the IU Electron Microscopy Center for image preparation and capture.

## Supplementary Materials and Methods

### Chemotaxis assay

A chemotaxis assay was performed with wildtype animals and those with a V5 epitope at the N-terminus of ADR-1 and three copies of the FLAG epitope at the N-terminus of ADR-2 (HAH36) as previously described (Tonkin et al. 2002). Briefly, chemotaxis plates (10 cm) were spotted with 1 ul of butanone (1:1000 dilution in ethanol) and 1 ul of ethanol (control) equidistant from the midpoint of the chemotaxis plate. To anesthetize animals reaching these regions, 1 μl of sodium azide (1 M) was added to the attractant and control spots. Between 100 and 150 young adult animals were placed in a circle at the center of the plate. After one hour, animals were counted to calculate a chemotaxis index. Chemotaxis index = animals at the attractant-animals at control)/total number of animals on the plate.

### RNA editing assay

Mixed stage worms were stored in TRIzol (Invitrogen) and RNA was extracted according to the methods described in the main manuscript. After DNase treatment and clean-up, the RNA was reverse transcribed with a gene specific primer (Supplementary Table 7) using Superscript III (Invitrogen). The resulting cDNA was amplified by of PCR with Phusion DNA polymerase (NEB) (primers listed in Supplementary Table 7). To confirm the resulting products were amplified from the RNA, negative controls were performed wherein the reverse transcription reaction was followed but without the addition of Superscript III. The resulting cDNA for *lam-2* gene was purified with gel electrophoresis and sent for Sanger sequencing (QuintaraBio).

## References

1. Wibisono P, Sun J: Neuro-immune communication in C. elegans defense against pathogen infection. Curr Res Immunol 2021, 2:60–65.

2. Kim DH, Ewbank JJ: Signaling in the innate immune response. WormBook 2018, 2018:1–35.

3. Hsu AP, Holland SM: Host genetics of innate immune system in infection. Current Opinion in Immunology 2022, 74:140–149.

4. Lisy S, Rothamel K, Ascano M: RNA Binding Proteins as Pioneer Determinants of Infection: Protective, Proviral, or Both? Viruses 2021, 13(11).

5. Girardi E, Pfeffer S, Baumert TF, Majzoub K: Roadblocks and fast tracks: How RNA binding proteins affect the viral RNA journey in the cell. Semin Cell Dev Biol 2021, 111:86–100.

6. Pfaller CK, George CX, Samuel CE: Adenosine Deaminases Acting on RNA (ADARs) and Viral Infections. Annu Rev Virol 2021, 8(1):239–264.

7. Reich DP, Bass BL: Mapping the dsRNA World. Cold Spring Harb Perspect Biol 2019, 11(3).

8. Chen YG, Hur S: Cellular origins of dsRNA, their recognition and consequences. Nat Rev Mol Cell Biol 2022, 23(4):286–301.

9. Vesely C, Jantsch MF: An I for an A: Dynamic Regulation of Adenosine Deamination-Mediated RNA Editing. Genes-Basel 2021, 12(7).

10. Quin J, Sedmik J, Vukic D, Khan A, Keegan LP, O’Connell MA: ADAR RNA Modifications, the Epitranscriptome and Innate Immunity. Trends Biochem Sci 2021, 46(9):758–771.

11. Rice GI, Kasher PR, Forte GM, Mannion NM, Greenwood SM, Szynkiewicz M, Dickerson JE, Bhaskar SS, Zampini M, Briggs TA et al: Mutations in ADAR1 cause Aicardi-Goutieres syndrome associated with a type I interferon signature. Nat Genet 2012, 44(11):1243–1248.

12. Mannion NM, Greenwood SM, Young R, Cox S, Brindle J, Read D, Nellaker C, Vesely C, Ponting CP, McLaughlin PJ et al: The RNA-editing enzyme ADAR1 controls innate immune responses to RNA. Cell Rep 2014, 9(4):1482–1494.

13. Liddicoat BJ, Piskol R, Chalk AM, Ramaswami G, Higuchi M, Hartner JC, Li JB, Seeburg PH, Walkley CR: RNA editing by ADAR1 prevents MDA5 sensing of endogenous dsRNA as nonself. Science 2015, 349(6252):1115–1120.

14. Pestal K, Funk CC, Snyder JM, Price ND, Treuting PM, Stetson DB: Isoforms of RNA-Editing Enzyme ADAR1 Independently Control Nucleic Acid Sensor MDA5-Driven Autoimmunity and Multi-organ Development. Immunity 2015, 43(5):933–944.

15. Deng P, Khan A, Jacobson D, Sambrani N, McGurk L, Li X, Jayasree A, Hejatko J, Shohat-Ophir G, O’Connell MA et al: Adar RNA editing-dependent and - independent effects are required for brain and innate immune functions in Drosophila. Nat Commun 2020, 11(1):1580.

16. Bar Yaacov D: Functional analysis of ADARs in planarians supports a bilaterian ancestral role in suppressing double-stranded RNA-response. PLoS Pathog 2022, 18(1):e1010250.

17. Niescierowicz K, Pryszcz L, Navarrete C, Tralle E, Sulej A, Abu Nahia K, Kasprzyk ME, Misztal K, Pateria A, Pakula A et al: Adar-mediated A-to-I editing is required for embryonic patterning and innate immune response regulation in zebrafish. Nat Commun 2022, 13(1):5520.

18. Arribere JA, Kuroyanagi H, Hundley HA: mRNA Editing, Processing and Quality Control in Caenorhabditis elegans. Genetics 2020, 215(3):531–568.

19. Tonkin LA, Saccomanno L, Morse DP, Brodigan T, Krause M, Bass BL: RNA editing by ADARs is important for normal behavior in Caenorhabditis elegans. EMBO J 2002, 21(22):6025–6035.

20. Ganem NS, Ben-Asher N, Manning AC, Deffit SN, Washburn MC, Wheeler EC, Yeo GW, Zgayer OB, Mantsur E, Hundley HA et al: Disruption in A-to-I Editing Levels Affects C. elegans Development More Than a Complete Lack of Editing. Cell Rep 2019, 27(4):1244–1253 e1244.

21. Rajendren S, Dhakal A, Vadlamani P, Townsend J, Deffit SN, Hundley HA: Profiling neural editomes reveals a molecular mechanism to regulate RNA editing during development. Genome Res 2021, 31(1):27–39.

22. Reich DP, Tyc KM, Bass BL: C. elegans ADARs antagonize silencing of cellular dsRNAs by the antiviral RNAi pathway. Genes Dev 2018, 32(3-4):271–282.

23. Fischer SEJ, Ruvkun G: Caenorhabditis elegans ADAR editing and the ERI-6/7/MOV10 RNAi pathway silence endogenous viral elements and LTR retrotransposons. Proc Natl Acad Sci U S A 2020, 117(11):5987–5996.

24. Harding BW, Ewbank JJ: An integrated view of innate immune mechanisms in C. elegans. Biochem Soc Trans 2021, 49(5):2307–2317.

25. Qin S, Xiao W, Zhou C, Pu Q, Deng X, Lan L, Liang H, Song X, Wu M: Pseudomonas aeruginosa: pathogenesis, virulence factors, antibiotic resistance, interaction with host, technology advances and emerging therapeutics. Signal Transduct Target Ther 2022, 7(1):199.

26. Tan MW, Mahajan-Miklos S, Ausubel FM: Killing of Caenorhabditis elegans by Pseudomonas aeruginosa used to model mammalian bacterial pathogenesis. Proc Natl Acad Sci U S A 1999, 96(2):715–720.

27. Singh J, Aballay A: Neural control of behavioral and molecular defenses in C. elegans. Curr Opin Neurobiol 2020, 62:34–40.

28. Pukkila-Worley R: Surveillance Immunity: An Emerging Paradigm of Innate Defense Activation in Caenorhabditis elegans. PLoS Pathog 2016, 12(9):e1005795.

29. Meisel JD, Kim DH: Behavioral avoidance of pathogenic bacteria by Caenorhabditis elegans. Trends Immunol 2014, 35(10):465–470.

30. Sun J, Singh V, Kajino-Sakamoto R, Aballay A: Neuronal GPCR controls innate immunity by regulating noncanonical unfolded protein response genes. Science 2011, 332(6030):729–732.

31. Sellegounder D, Yuan CH, Wibisono P, Liu Y, Sun J: Octopaminergic Signaling Mediates Neural Regulation of Innate Immunity in Caenorhabditis elegans. mBio 2018, 9(5).

32. Shapira M, Hamlin BJ, Rong J, Chen K, Ronen M, Tan MW: A conserved role for a GATA transcription factor in regulating epithelial innate immune responses. Proc Natl Acad Sci U S A 2006, 103(38):14086–14091.

33. Troemel ER, Chu SW, Reinke V, Lee SS, Ausubel FM, Kim DH: p38 MAPK regulates expression of immune response genes and contributes to longevity in C. elegans. PLoS Genet 2006, 2(11):e183.

34. Rajendren S, Manning AC, Al-Awadi H, Yamada K, Takagi Y, Hundley HA: A protein-protein interaction underlies the molecular basis for substrate recognition by an adenosine-to-inosine RNA-editing enzyme. Nucleic Acids Res 2018, 46(18):9647–9659.

35. Washburn MC, Kakaradov B, Sundararaman B, Wheeler E, Hoon S, Yeo GW, Hundley HA: The dsRBP and inactive editor ADR-1 utilizes dsRNA binding to regulate A-to-I RNA editing across the C. elegans transcriptome. Cell Rep 2014, 6(4):599–607.

36. Erdmann EA, Mahapatra A, Mukherjee P, Yang B, Hundley HA: To protect and modify double-stranded RNA -the critical roles of ADARs in development, immunity and oncogenesis. Crit Rev Biochem Mol Biol 2021, 56(1):54–87.

37. Holdorf AD, Higgins DP, Hart AC, Boag PR, Pazour GJ, Walhout AJM, Walker AK: WormCat: An Online Tool for Annotation and Visualization of Caenorhabditis elegans Genome-Scale Data. Genetics 2020, 214(2):279–294.

38. Page AP, Johnstone IL: The cuticle. WormBook 2007:1–15.

39. Head BP, Olaitan AO, Aballay A: Role of GATA transcription factor ELT-2 and p38 MAPK PMK-1 in recovery from acute P. aeruginosa infection in C. elegans. Virulence 2017, 8(3):261–274.

40. Teuscher AC, Jongsma E, Davis MN, Statzer C, Gebauer JM, Naba A, Ewald CY: The in-silico characterization of the Caenorhabditis elegans matrisome and proposal of a novel collagen classification. Matrix Biol Plus 2019, 1:100001.

41. Herndon LA, Wolkow CA, Driscoll M, Hall DH: Effects of Ageing on the Basic Biology and Anatomy of C. elegans. Healthy Ageing Long 2017, 5:9–39.

42. Sellegounder D, Liu Y, Wibisono P, Chen CH, Leap D, Sun J: Neuronal GPCR NPR-8 regulates C. elegans defense against pathogen infection. Sci Adv 2019, 5(11):eaaw4717.

43. Aballay A, Yorgey P, Ausubel FM: Salmonella typhimurium proliferates and establishes a persistent infection in the intestine of Caenorhabditis elegans. Curr Biol 2000, 10(23):1539–1542.

44. Labrousse A, Chauvet S, Couillault C, Kurz CL, Ewbank JJ: Caenorhabditis elegans is a model host for Salmonella typhimurium. Curr Biol 2000, 10(23):1543–1545.

45. Sifri CD, Begun J, Ausubel FM, Calderwood SB: Caenorhabditis elegans as a model host for Staphylococcus aureus pathogenesis. Infect Immun 2003, 71(4):2208–2217.

46. Shevchenko G, Morris KV: All I’s on the RADAR: role of ADAR in gene regulation. FEBS Lett 2018, 592(17):2860–2873.

47. Heissenberger C, Rollins JA, Krammer TL, Nagelreiter F, Stocker I, Wacheul L, Shpylovyi A, Tav K, Snow S, Grillari J et al: The ribosomal RNA m(5)C methyltransferase NSUN-1 modulates healthspan and oogenesis in Caenorhabditis elegans. Elife 2020, 9.

48. Ewald CY, Landis JN, Porter Abate J, Murphy CT, Blackwell TK: Dauer-independent insulin/IGF-1-signalling implicates collagen remodelling in longevity. Nature 2015, 519(7541):97–101.

49. Shaw WM, Luo S, Landis J, Ashraf J, Murphy CT: The C. elegans TGF-beta Dauer pathway regulates longevity via insulin signaling. Curr Biol 2007, 17(19):1635–1645.

50. Palani SN, Sellegounder D, Wibisono P, Liu Y: The longevity response to warm temperature is neurally controlled via the regulation of collagen genes. Aging Cell 2023:e13815.

51. Goodman MB, Savage-Dunn C: Reciprocal interactions between transforming growth factor beta signaling and collagens: Insights from Caenorhabditis elegans. Dev Dyn 2022, 251(1):47–60.

52. Brunquell J, Morris S, Lu Y, Cheng F, Westerheide SD: The genome-wide role of HSF-1 in the regulation of gene expression in Caenorhabditis elegans. BMC Genomics 2016, 17:559.

53. Pereira AG, Gracida X, Kagias K, Zhang Y: C. elegans aversive olfactory learning generates diverse intergenerational effects. J Neurogenet 2020, 34(3-4):378–388.

54. Kaletsky R, Moore RS, Vrla GD, Parsons LR, Gitai Z, Murphy CT: C. elegans interprets bacterial non-coding RNAs to learn pathogenic avoidance. Nature 2020, 586(7829):445–451.

55. Hundley HA, Krauchuk AA, Bass BL: C. elegans and H. sapiens mRNAs with edited 3’ UTRs are present on polysomes. RNA 2008, 14(10):2050–2060.

56. Paix A, Folkmann A, Seydoux G: Precision genome editing using CRISPR-Cas9 and linear repair templates in C. elegans. Methods 2017, 121-122:86–93.

57. Ghanta KS, Ishidate T, Mello CC: Microinjection for precision genome editing in Caenorhabditis elegans. STAR Protoc 2021, 2(3):100748.

58. Wibisono P, Sun J: Protocol to measure bacterial intake and gut clearance of Caenorhabditis elegans. STAR Protoc 2022, 3(3):101558.

