## Additional File 1 for "ADARs regulate cuticle collagen expression to promote survival to pathogen infection": Additional File 1.pdf

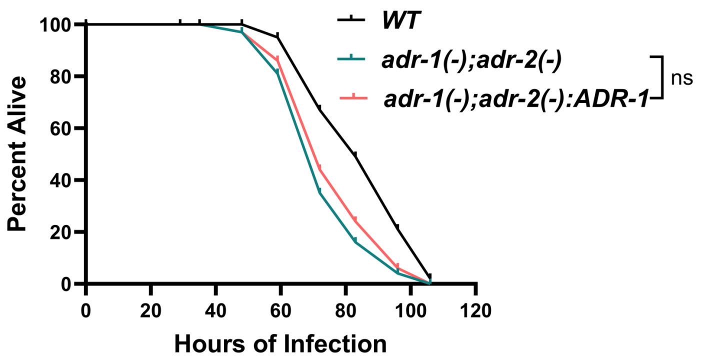


**Additional File 1: Fig. S1. ADR-1 alone is not sufficient to rescue the susceptibility phenotype of the *adr-1(-);adr-2(-)* animals.** Representative survival curve (of three independent biological replicates) for the indicated strains subjected to the slow-killing assay and scored for survival in response to *Pseudomonas aeruginosa* strain PA14. Statistical significance was determined using OASIS, ns indicates no significant difference (*p* >0.05)
