## Additional File 2 for "ADARs regulate cuticle collagen expression to promote survival to pathogen infection": Additional File 2.pdf

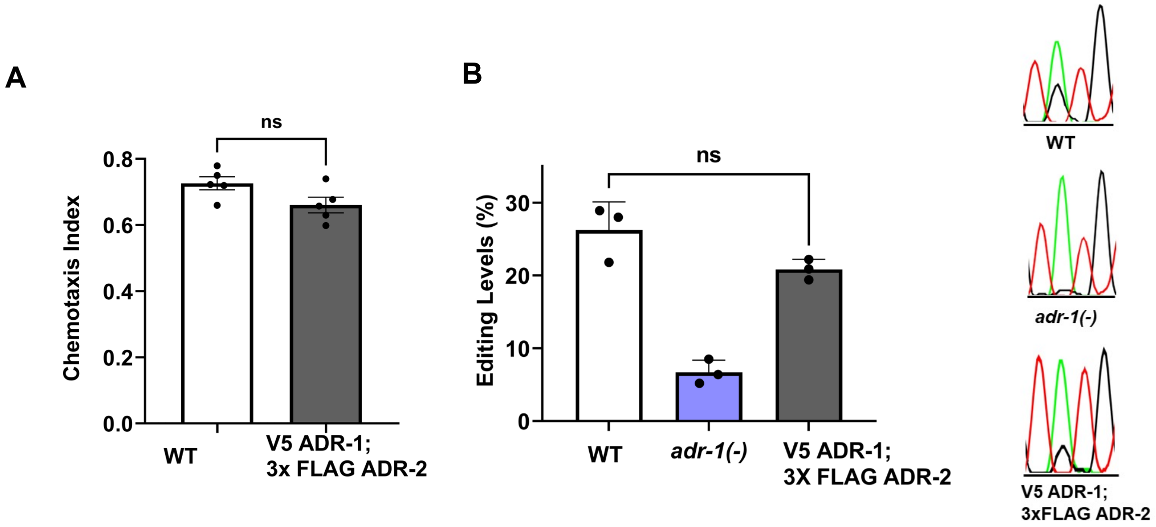


**Additional File 2: Fig. S2. Presence of epitope tags does not significantly alter chemotaxis behavior or RNA editing.** A) Chemotaxis index was calculated for the indicated strains. The graph represents the mean of three technical replicates for each strain. Error bars represent SEM. Statistical significance was measured using unpaired Mann Whitney test. *p* value was not significant (ns) ($p>0.05$). B) The bar graph represents the average editing levels quantified from the Sanger sequencing traces (representative traces on right) from three biological replicates. Editing at site 164 within *lam-2*, which is a known ADR-1 regulated site (Washburn et al. 2014). Error bars represent the standard error of the mean (SEM). Statistical significance was determined using a one-way ANOVA, *p* value was not significant (ns) ($p>0.05$). The nucleotides in the Sanger sequencing chromatogram are characterized by different colors (green, adenosine; black, guanosine; red, thymidine).
