## Additional File 6 for "ADARs regulate cuticle collagen expression to promote survival to pathogen infection": Additional File 6.txt

"Additional file 6: Table S6. Differentially expressed genes in adr-1(-);adr-2(-) animals grown on OP50 for 7 hours compared to wildtype animals grown in the same conditions (Related to Figure 4). Differentially expressed genes are listed as Wormbase annotations in Column A and Gene name in Column B. The log2fold change values, raw p values and the adjusted p values as obtained from the DESeq2 analysis are shown in columns C, D, and E, respectively. Read counts from the DESeq2 analysis for the three biological replicates of wildtype are listed in columns F, G and H. Read counts from the DESeq2 analysis for the three biological replicates of adr-1(-);adr-2(-) animals  are listed in columns I, J and K . "																																																																																																																																																																																																																																																																																																																																																																																																																																																																																																																																																																																																																																																																																																																																																																																																																																																																																																																																																																																																																																																																																																																																																																																																																																																																					"Supplemental Table 4: Differentially expressed genes in adr-1(-);adr-2(-) animals grown on OP40 for 7 hours compared to wildtype animals grown on OP50 for 7 hours (Related to Figure 4). DESeq2 was used to identify differentially expressed genes in adr-1(-);adr-2(-) double mutant animals compared to wildtype animals. p-value<0.05 were taken as targets misregulated in absence of ad-1 and adr-2. Differentially expressed genes are listed as Wormbase annotations in Column A, Gene name in Column B. The log2fold change values, raw p values and the adjusted p values as obtained from the DESeq2 analysis are shown in columns C, D, and E, respectively. Read counts from the DESeq2 analysis for the three biological replicates of wildtype are listed in columns F, G and H. Read counts from the DESeq2 analysis for the three biological replicates of adr-1(-);adr-2(-) animals are listed in columns I, J and K . "	"Supplemental Table 4: Differentially expressed genes in adr-1(-);adr-2(-) animals grown on OP40 for 7 hours compared to wildtype animals grown on OP50 for 7 hours (Related to Figure 4). DESeq2 was used to identify differentially expressed genes in adr-1(-);adr-2(-) double mutant animals compared to wildtype animals. p-value<0.05 were taken as targets misregulated in absence of ad-1 and adr-2. Differentially expressed genes are listed as Wormbase annotations in Column A, Gene name in Column B. The log2fold change values, raw p values and the adjusted p values as obtained from the DESeq2 analysis are shown in columns C, D, and E, respectively. Read counts from the DESeq2 analysis for the three biological replicates of wildtype are listed in columns F, G and H. Read counts from the DESeq2 analysis for the three biological replicates of adr-1(-);adr-2(-) animals are listed in columns I, J and K . "	"Supplemental Table 4: Differentially expressed genes in adr-1(-);adr-2(-) animals grown on OP40 for 7 hours compared to wildtype animals grown on OP50 for 7 hours (Related to Figure 4). DESeq2 was used to identify differentially expressed genes in adr-1(-);adr-2(-) double mutant animals compared to wildtype animals. p-value<0.05 were taken as targets misregulated in absence of ad-1 and adr-2. Differentially expressed genes are listed as Wormbase annotations in Column A, Gene name in Column B. The log2fold change values, raw p values and the adjusted p values as obtained from the DESeq2 analysis are shown in columns C, D, and E, respectively. Read counts from the DESeq2 analysis for the three biological replicates of wildtype are listed in columns F, G and H. Read counts from the DESeq2 analysis for the three biological replicates of adr-1(-);adr-2(-) animals are listed in columns I, J and K . "	"Supplemental Table 4: Differentially expressed genes in adr-1(-);adr-2(-) animals grown on OP40 for 7 hours compared to wildtype animals grown on OP50 for 7 hours (Related to Figure 4). DESeq2 was used to identify differentially expressed genes in adr-1(-);adr-2(-) double mutant animals compared to wildtype animals. p-value<0.05 were taken as targets misregulated in absence of ad-1 and adr-2. Differentially expressed genes are listed as Wormbase annotations in Column A, Gene name in Column B. The log2fold change values, raw p values and the adjusted p values as obtained from the DESeq2 analysis are shown in columns C, D, and E, respectively. Read counts from the DESeq2 analysis for the three biological replicates of wildtype are listed in columns F, G and H. Read counts from the DESeq2 analysis for the three biological replicates of adr-1(-);adr-2(-) animals are listed in columns I, J and K . "	"Supplemental Table 4: Differentially expressed genes in adr-1(-);adr-2(-) animals grown on OP40 for 7 hours compared to wildtype animals grown on OP50 for 7 hours (Related to Figure 4). DESeq2 was used to identify differentially expressed genes in adr-1(-);adr-2(-) double mutant animals compared to wildtype animals. p-value<0.05 were taken as targets misregulated in absence of ad-1 and adr-2. Differentially expressed genes are listed as Wormbase annotations in Column A, Gene name in Column B. The log2fold change values, raw p values and the adjusted p values as obtained from the DESeq2 analysis are shown in columns C, D, and E, respectively. Read counts from the DESeq2 analysis for the three biological replicates of wildtype are listed in columns F, G and H. Read counts from the DESeq2 analysis for the three biological replicates of adr-1(-);adr-2(-) animals are listed in columns I, J and K . "	"Supplemental Table 4: Differentially expressed genes in adr-1(-);adr-2(-) animals grown on OP40 for 7 hours compared to wildtype animals grown on OP50 for 7 hours (Related to Figure 4). DESeq2 was used to identify differentially expressed genes in adr-1(-);adr-2(-) double mutant animals compared to wildtype animals. p-value<0.05 were taken as targets misregulated in absence of ad-1 and adr-2. Differentially expressed genes are listed as Wormbase annotations in Column A, Gene name in Column B. The log2fold change values, raw p values and the adjusted p values as obtained from the DESeq2 analysis are shown in columns C, D, and E, respectively. Read counts from the DESeq2 analysis for the three biological replicates of wildtype are listed in columns F, G and H. Read counts from the DESeq2 analysis for the three biological replicates of adr-1(-);adr-2(-) animals are listed in columns I, J and K . "	"Supplemental Table 4: Differentially expressed genes in adr-1(-);adr-2(-) animals grown on OP40 for 7 hours compared to wildtype animals grown on OP50 for 7 hours (Related to Figure 4). DESeq2 was used to identify differentially expressed genes in adr-1(-);adr-2(-) double mutant animals compared to wildtype animals. p-value<0.05 were taken as targets misregulated in absence of ad-1 and adr-2. Differentially expressed genes are listed as Wormbase annotations in Column A, Gene name in Column B. The log2fold change values, raw p values and the adjusted p values as obtained from the DESeq2 analysis are shown in columns C, D, and E, respectively. Read counts from the DESeq2 analysis for the three biological replicates of wildtype are listed in columns F, G and H. Read counts from the DESeq2 analysis for the three biological replicates of adr-1(-);adr-2(-) animals are listed in columns I, J and K . "	"Supplemental Table 4: Differentially expressed genes in adr-1(-);adr-2(-) animals grown on OP40 for 7 hours compared to wildtype animals grown on OP50 for 7 hours (Related to Figure 4). DESeq2 was used to identify differentially expressed genes in adr-1(-);adr-2(-) double mutant animals compared to wildtype animals. p-value<0.05 were taken as targets misregulated in absence of ad-1 and adr-2. Differentially expressed genes are listed as Wormbase annotations in Column A, Gene name in Column B. The log2fold change values, raw p values and the adjusted p values as obtained from the DESeq2 analysis are shown in columns C, D, and E, respectively. Read counts from the DESeq2 analysis for the three biological replicates of wildtype are listed in columns F, G and H. Read counts from the DESeq2 analysis for the three biological replicates of adr-1(-);adr-2(-) animals are listed in columns I, J and K . "	"Supplemental Table 4: Differentially expressed genes in adr-1(-);adr-2(-) animals grown on OP40 for 7 hours compared to wildtype animals grown on OP50 for 7 hours (Related to Figure 4). DESeq2 was used to identify differentially expressed genes in adr-1(-);adr-2(-) double mutant animals compared to wildtype animals. p-value<0.05 were taken as targets misregulated in absence of ad-1 and adr-2. Differentially expressed genes are listed as Wormbase annotations in Column A, Gene name in Column B. The log2fold change values, raw p values and the adjusted p values as obtained from the DESeq2 analysis are shown in columns C, D, and E, respectively. Read counts from the DESeq2 analysis for the three biological replicates of wildtype are listed in columns F, G and H. Read counts from the DESeq2 analysis for the three biological replicates of adr-1(-);adr-2(-) animals are listed in columns I, J and K . "	"Supplemental Table 4: Differentially expressed genes in adr-1(-);adr-2(-) animals grown on OP40 for 7 hours compared to wildtype animals grown on OP50 for 7 hours (Related to Figure 4). DESeq2 was used to identify differentially expressed genes in adr-1(-);adr-2(-) double mutant animals compared to wildtype animals. p-value<0.05 were taken as targets misregulated in absence of ad-1 and adr-2. Differentially expressed genes are listed as Wormbase annotations in Column A, Gene name in Column B. The log2fold change values, raw p values and the adjusted p values as obtained from the DESeq2 analysis are shown in columns C, D, and E, respectively. Read counts from the DESeq2 analysis for the three biological replicates of wildtype are listed in columns F, G and H. Read counts from the DESeq2 analysis for the three biological replicates of adr-1(-);adr-2(-) animals are listed in columns I, J and K . "	"Supplemental Table 4: Differentially expressed genes in adr-1(-);adr-2(-) animals grown on OP40 for 7 hours compared to wildtype animals grown on OP50 for 7 hours (Related to Figure 4). DESeq2 was used to identify differentially expressed genes in adr-1(-);adr-2(-) double mutant animals compared to wildtype animals. p-value<0.05 were taken as targets misregulated in absence of ad-1 and adr-2. Differentially expressed genes are listed as Wormbase annotations in Column A, Gene name in Column B. The log2fold change values, raw p values and the adjusted p values as obtained from the DESeq2 analysis are shown in columns C, D, and E, respectively. Read counts from the DESeq2 analysis for the three biological replicates of wildtype are listed in columns F, G and H. Read counts from the DESeq2 analysis for the three biological replicates of adr-1(-);adr-2(-) animals are listed in columns I, J and K . "	"Supplemental Table 4: Differentially expressed genes in adr-1(-);adr-2(-) animals grown on OP40 for 7 hours compared to wildtype animals grown on OP50 for 7 hours (Related to Figure 4). DESeq2 was used to identify differentially expressed genes in adr-1(-);adr-2(-) double mutant animals compared to wildtype animals. p-value<0.05 were taken as targets misregulated in absence of ad-1 and adr-2. Differentially expressed genes are listed as Wormbase annotations in Column A, Gene name in Column B. The log2fold change values, raw p values and the adjusted p values as obtained from the DESeq2 analysis are shown in columns C, D, and E, respectively. Read counts from the DESeq2 analysis for the three biological replicates of wildtype are listed in columns F, G and H. Read counts from the DESeq2 analysis for the three biological replicates of adr-1(-);adr-2(-) animals are listed in columns I, J and K . "	"Supplemental Table 4: Differentially expressed genes in adr-1(-);adr-2(-) animals grown on OP40 for 7 hours compared to wildtype animals grown on OP50 for 7 hours (Related to Figure 4). DESeq2 was used to identify differentially expressed genes in adr-1(-);adr-2(-) double mutant animals compared to wildtype animals. p-value<0.05 were taken as targets misregulated in absence of ad-1 and adr-2. Differentially expressed genes are listed as Wormbase annotations in Column A, Gene name in Column B. The log2fold change values, raw p values and the adjusted p values as obtained from the DESeq2 analysis are shown in columns C, D, and E, respectively. Read counts from the DESeq2 analysis for the three biological replicates of wildtype are listed in columns F, G and H. Read counts from the DESeq2 analysis for the three biological replicates of adr-1(-);adr-2(-) animals are listed in columns I, J and K . "	"Supplemental Table 4: Differentially expressed genes in adr-1(-);adr-2(-) animals grown on OP40 for 7 hours compared to wildtype animals grown on OP50 for 7 hours (Related to Figure 4). DESeq2 was used to identify differentially expressed genes in adr-1(-);adr-2(-) double mutant animals compared to wildtype animals. p-value<0.05 were taken as targets misregulated in absence of ad-1 and adr-2. Differentially expressed genes are listed as Wormbase annotations in Column A, Gene name in Column B. The log2fold change values, raw p values and the adjusted p values as obtained from the DESeq2 analysis are shown in columns C, D, and E, respectively. Read counts from the DESeq2 analysis for the three biological replicates of wildtype are listed in columns F, G and H. Read counts from the DESeq2 analysis for the three biological replicates of adr-1(-);adr-2(-) animals are listed in columns I, J and K . "	"Supplemental Table 4: Differentially expressed genes in adr-1(-);adr-2(-) animals grown on OP40 for 7 hours compared to wildtype animals grown on OP50 for 7 hours (Related to Figure 4). DESeq2 was used to identify differentially expressed genes in adr-1(-);adr-2(-) double mutant animals compared to wildtype animals. p-value<0.05 were taken as targets misregulated in absence of ad-1 and adr-2. Differentially expressed genes are listed as Wormbase annotations in Column A, Gene name in Column B. The log2fold change values, raw p values and the adjusted p values as obtained from the DESeq2 analysis are shown in columns C, D, and E, respectively. Read counts from the DESeq2 analysis for the three biological replicates of wildtype are listed in columns F, G and H. Read counts from the DESeq2 analysis for the three biological replicates of adr-1(-);adr-2(-) animals are listed in columns I, J and K . "	"Supplemental Table 4: Differentially expressed genes in adr-1(-);adr-2(-) animals grown on OP40 for 7 hours compared to wildtype animals grown on OP50 for 7 hours (Related to Figure 4). DESeq2 was used to identify differentially expressed genes in adr-1(-);adr-2(-) double mutant animals compared to wildtype animals. p-value<0.05 were taken as targets misregulated in absence of ad-1 and adr-2. Differentially expressed genes are listed as Wormbase annotations in Column A, Gene name in Column B. The log2fold change values, raw p values and the adjusted p values as obtained from the DESeq2 analysis are shown in columns C, D, and E, respectively. Read counts from the DESeq2 analysis for the three biological replicates of wildtype are listed in columns F, G and H. Read counts from the DESeq2 analysis for the three biological replicates of adr-1(-);adr-2(-) animals are listed in columns I, J and K . "	"Supplemental Table 4: Differentially expressed genes in adr-1(-);adr-2(-) animals grown on OP40 for 7 hours compared to wildtype animals grown on OP50 for 7 hours (Related to Figure 4). DESeq2 was used to identify differentially expressed genes in adr-1(-);adr-2(-) double mutant animals compared to wildtype animals. p-value<0.05 were taken as targets misregulated in absence of ad-1 and adr-2. Differentially expressed genes are listed as Wormbase annotations in Column A, Gene name in Column B. The log2fold change values, raw p values and the adjusted p values as obtained from the DESeq2 analysis are shown in columns C, D, and E, respectively. Read counts from the DESeq2 analysis for the three biological replicates of wildtype are listed in columns F, G and H. Read counts from the DESeq2 analysis for the three biological replicates of adr-1(-);adr-2(-) animals are listed in columns I, J and K . "	"Supplemental Table 4: Differentially expressed genes in adr-1(-);adr-2(-) animals grown on OP40 for 7 hours compared to wildtype animals grown on OP50 for 7 hours (Related to Figure 4). DESeq2 was used to identify differentially expressed genes in adr-1(-);adr-2(-) double mutant animals compared to wildtype animals. p-value<0.05 were taken as targets misregulated in absence of ad-1 and adr-2. Differentially expressed genes are listed as Wormbase annotations in Column A, Gene name in Column B. The log2fold change values, raw p values and the adjusted p values as obtained from the DESeq2 analysis are shown in columns C, D, and E, respectively. Read counts from the DESeq2 analysis for the three biological replicates of wildtype are listed in columns F, G and H. Read counts from the DESeq2 analysis for the three biological replicates of adr-1(-);adr-2(-) animals are listed in columns I, J and K . "	"Supplemental Table 4: Differentially expressed genes in adr-1(-);adr-2(-) animals grown on OP40 for 7 hours compared to wildtype animals grown on OP50 for 7 hours (Related to Figure 4). DESeq2 was used to identify differentially expressed genes in adr-1(-);adr-2(-) double mutant animals compared to wildtype animals. p-value<0.05 were taken as targets misregulated in absence of ad-1 and adr-2. Differentially expressed genes are listed as Wormbase annotations in Column A, Gene name in Column B. The log2fold change values, raw p values and the adjusted p values as obtained from the DESeq2 analysis are shown in columns C, D, and E, respectively. Read counts from the DESeq2 analysis for the three biological replicates of wildtype are listed in columns F, G and H. Read counts from the DESeq2 analysis for the three biological replicates of adr-1(-);adr-2(-) animals are listed in columns I, J and K . "	"Supplemental Table 4: Differentially expressed genes in adr-1(-);adr-2(-) animals grown on OP40 for 7 hours compared to wildtype animals grown on OP50 for 7 hours (Related to Figure 4). DESeq2 was used to identify differentially expressed genes in adr-1(-);adr-2(-) double mutant animals compared to wildtype animals. p-value<0.05 were taken as targets misregulated in absence of ad-1 and adr-2. Differentially expressed genes are listed as Wormbase annotations in Column A, Gene name in Column B. The log2fold change values, raw p values and the adjusted p values as obtained from the DESeq2 analysis are shown in columns C, D, and E, respectively. Read counts from the DESeq2 analysis for the three biological replicates of wildtype are listed in columns F, G and H. Read counts from the DESeq2 analysis for the three biological replicates of adr-1(-);adr-2(-) animals are listed in columns I, J and K . "	"Supplemental Table 4: Differentially expressed genes in adr-1(-);adr-2(-) animals grown on OP40 for 7 hours compared to wildtype animals grown on OP50 for 7 hours (Related to Figure 4). DESeq2 was used to identify differentially expressed genes in adr-1(-);adr-2(-) double mutant animals compared to wildtype animals. p-value<0.05 were taken as targets misregulated in absence of ad-1 and adr-2. Differentially expressed genes are listed as Wormbase annotations in Column A, Gene name in Column B. The log2fold change values, raw p values and the adjusted p values as obtained from the DESeq2 analysis are shown in columns C, D, and E, respectively. Read counts from the DESeq2 analysis for the three biological replicates of wildtype are listed in columns F, G and H. Read counts from the DESeq2 analysis for the three biological replicates of adr-1(-);adr-2(-) animals are listed in columns I, J and K . "	"Supplemental Table 4: Differentially expressed genes in adr-1(-);adr-2(-) animals grown on OP40 for 7 hours compared to wildtype animals grown on OP50 for 7 hours (Related to Figure 4). DESeq2 was used to identify differentially expressed genes in adr-1(-);adr-2(-) double mutant animals compared to wildtype animals. p-value<0.05 were taken as targets misregulated in absence of ad-1 and adr-2. Differentially expressed genes are listed as Wormbase annotations in Column A, Gene name in Column B. The log2fold change values, raw p values and the adjusted p values as obtained from the DESeq2 analysis are shown in columns C, D, and E, respectively. Read counts from the DESeq2 analysis for the three biological replicates of wildtype are listed in columns F, G and H. Read counts from the DESeq2 analysis for the three biological replicates of adr-1(-);adr-2(-) animals are listed in columns I, J and K . "	"Supplemental Table 4: Differentially expressed genes in adr-1(-);adr-2(-) animals grown on OP40 for 7 hours compared to wildtype animals grown on OP50 for 7 hours (Related to Figure 4). DESeq2 was used to identify differentially expressed genes in adr-1(-);adr-2(-) double mutant animals compared to wildtype animals. p-value<0.05 were taken as targets misregulated in absence of ad-1 and adr-2. Differentially expressed genes are listed as Wormbase annotations in Column A, Gene name in Column B. The log2fold change values, raw p values and the adjusted p values as obtained from the DESeq2 analysis are shown in columns C, D, and E, respectively. Read counts from the DESeq2 analysis for the three biological replicates of wildtype are listed in columns F, G and H. Read counts from the DESeq2 analysis for the three biological replicates of adr-1(-);adr-2(-) animals are listed in columns I, J and K . "	"Supplemental Table 4: Differentially expressed genes in adr-1(-);adr-2(-) animals grown on OP40 for 7 hours compared to wildtype animals grown on OP50 for 7 hours (Related to Figure 4). DESeq2 was used to identify differentially expressed genes in adr-1(-);adr-2(-) double mutant animals compared to wildtype animals. p-value<0.05 were taken as targets misregulated in absence of ad-1 and adr-2. Differentially expressed genes are listed as Wormbase annotations in Column A, Gene name in Column B. The log2fold change values, raw p values and the adjusted p values as obtained from the DESeq2 analysis are shown in columns C, D, and E, respectively. Read counts from the DESeq2 analysis for the three biological replicates of wildtype are listed in columns F, G and H. Read counts from the DESeq2 analysis for the three biological replicates of adr-1(-);adr-2(-) animals are listed in columns I, J and K . "	"Supplemental Table 4: Differentially expressed genes in adr-1(-);adr-2(-) animals grown on OP40 for 7 hours compared to wildtype animals grown on OP50 for 7 hours (Related to Figure 4). DESeq2 was used to identify differentially expressed genes in adr-1(-);adr-2(-) double mutant animals compared to wildtype animals. p-value<0.05 were taken as targets misregulated in absence of ad-1 and adr-2. Differentially expressed genes are listed as Wormbase annotations in Column A, Gene name in Column B. The log2fold change values, raw p values and the adjusted p values as obtained from the DESeq2 analysis are shown in columns C, D, and E, respectively. Read counts from the DESeq2 analysis for the three biological replicates of wildtype are listed in columns F, G and H. Read counts from the DESeq2 analysis for the three biological replicates of adr-1(-);adr-2(-) animals are listed in columns I, J and K . "	"Supplemental Table 4: Differentially expressed genes in adr-1(-);adr-2(-) animals grown on OP40 for 7 hours compared to wildtype animals grown on OP50 for 7 hours (Related to Figure 4). DESeq2 was used to identify differentially expressed genes in adr-1(-);adr-2(-) double mutant animals compared to wildtype animals. p-value<0.05 were taken as targets misregulated in absence of ad-1 and adr-2. Differentially expressed genes are listed as Wormbase annotations in Column A, Gene name in Column B. The log2fold change values, raw p values and the adjusted p values as obtained from the DESeq2 analysis are shown in columns C, D, and E, respectively. Read counts from the DESeq2 analysis for the three biological replicates of wildtype are listed in columns F, G and H. Read counts from the DESeq2 analysis for the three biological replicates of adr-1(-);adr-2(-) animals are listed in columns I, J and K . "	"Supplemental Table 4: Differentially expressed genes in adr-1(-);adr-2(-) animals grown on OP40 for 7 hours compared to wildtype animals grown on OP50 for 7 hours (Related to Figure 4). DESeq2 was used to identify differentially expressed genes in adr-1(-);adr-2(-) double mutant animals compared to wildtype animals. p-value<0.05 were taken as targets misregulated in absence of ad-1 and adr-2. Differentially expressed genes are listed as Wormbase annotations in Column A, Gene name in Column B. The log2fold change values, raw p values and the adjusted p values as obtained from the DESeq2 analysis are shown in columns C, D, and E, respectively. Read counts from the DESeq2 analysis for the three biological replicates of wildtype are listed in columns F, G and H. Read counts from the DESeq2 analysis for the three biological replicates of adr-1(-);adr-2(-) animals are listed in columns I, J and K . "	"Supplemental Table 4: Differentially expressed genes in adr-1(-);adr-2(-) animals grown on OP40 for 7 hours compared to wildtype animals grown on OP50 for 7 hours (Related to Figure 4). DESeq2 was used to identify differentially expressed genes in adr-1(-);adr-2(-) double mutant animals compared to wildtype animals. p-value<0.05 were taken as targets misregulated in absence of ad-1 and adr-2. Differentially expressed genes are listed as Wormbase annotations in Column A, Gene name in Column B. The log2fold change values, raw p values and the adjusted p values as obtained from the DESeq2 analysis are shown in columns C, D, and E, respectively. Read counts from the DESeq2 analysis for the three biological replicates of wildtype are listed in columns F, G and H. Read counts from the DESeq2 analysis for the three biological replicates of adr-1(-);adr-2(-) animals are listed in columns I, J and K . "	"Supplemental Table 4: Differentially expressed genes in adr-1(-);adr-2(-) animals grown on OP40 for 7 hours compared to wildtype animals grown on OP50 for 7 hours (Related to Figure 4). DESeq2 was used to identify differentially expressed genes in adr-1(-);adr-2(-) double mutant animals compared to wildtype animals. p-value<0.05 were taken as targets misregulated in absence of ad-1 and adr-2. Differentially expressed genes are listed as Wormbase annotations in Column A, Gene name in Column B. The log2fold change values, raw p values and the adjusted p values as obtained from the DESeq2 analysis are shown in columns C, D, and E, respectively. Read counts from the DESeq2 analysis for the three biological replicates of wildtype are listed in columns F, G and H. Read counts from the DESeq2 analysis for the three biological replicates of adr-1(-);adr-2(-) animals are listed in columns I, J and K . "	"Supplemental Table 4: Differentially expressed genes in adr-1(-);adr-2(-) animals grown on OP40 for 7 hours compared to wildtype animals grown on OP50 for 7 hours (Related to Figure 4). DESeq2 was used to identify differentially expressed genes in adr-1(-);adr-2(-) double mutant animals compared to wildtype animals. p-value<0.05 were taken as targets misregulated in absence of ad-1 and adr-2. Differentially expressed genes are listed as Wormbase annotations in Column A, Gene name in Column B. The log2fold change values, raw p values and the adjusted p values as obtained from the DESeq2 analysis are shown in columns C, D, and E, respectively. Read counts from the DESeq2 analysis for the three biological replicates of wildtype are listed in columns F, G and H. Read counts from the DESeq2 analysis for the three biological replicates of adr-1(-);adr-2(-) animals are listed in columns I, J and K . "	"Supplemental Table 4: Differentially expressed genes in adr-1(-);adr-2(-) animals grown on OP40 for 7 hours compared to wildtype animals grown on OP50 for 7 hours (Related to Figure 4). DESeq2 was used to identify differentially expressed genes in adr-1(-);adr-2(-) double mutant animals compared to wildtype animals. p-value<0.05 were taken as targets misregulated in absence of ad-1 and adr-2. Differentially expressed genes are listed as Wormbase annotations in Column A, Gene name in Column B. The log2fold change values, raw p values and the adjusted p values as obtained from the DESeq2 analysis are shown in columns C, D, and E, respectively. Read counts from the DESeq2 analysis for the three biological replicates of wildtype are listed in columns F, G and H. Read counts from the DESeq2 analysis for the three biological replicates of adr-1(-);adr-2(-) animals are listed in columns I, J and K . "	"Supplemental Table 4: Differentially expressed genes in adr-1(-);adr-2(-) animals grown on OP40 for 7 hours compared to wildtype animals grown on OP50 for 7 hours (Related to Figure 4). DESeq2 was used to identify differentially expressed genes in adr-1(-);adr-2(-) double mutant animals compared to wildtype animals. p-value<0.05 were taken as targets misregulated in absence of ad-1 and adr-2. Differentially expressed genes are listed as Wormbase annotations in Column A, Gene name in Column B. The log2fold change values, raw p values and the adjusted p values as obtained from the DESeq2 analysis are shown in columns C, D, and E, respectively. Read counts from the DESeq2 analysis for the three biological replicates of wildtype are listed in columns F, G and H. Read counts from the DESeq2 analysis for the three biological replicates of adr-1(-);adr-2(-) animals are listed in columns I, J and K . "	"Supplemental Table 4: Differentially expressed genes in adr-1(-);adr-2(-) animals grown on OP40 for 7 hours compared to wildtype animals grown on OP50 for 7 hours (Related to Figure 4). DESeq2 was used to identify differentially expressed genes in adr-1(-);adr-2(-) double mutant animals compared to wildtype animals. p-value<0.05 were taken as targets misregulated in absence of ad-1 and adr-2. Differentially expressed genes are listed as Wormbase annotations in Column A, Gene name in Column B. The log2fold change values, raw p values and the adjusted p values as obtained from the DESeq2 analysis are shown in columns C, D, and E, respectively. Read counts from the DESeq2 analysis for the three biological replicates of wildtype are listed in columns F, G and H. Read counts from the DESeq2 analysis for the three biological replicates of adr-1(-);adr-2(-) animals are listed in columns I, J and K . "	"Supplemental Table 4: Differentially expressed genes in adr-1(-);adr-2(-) animals grown on OP40 for 7 hours compared to wildtype animals grown on OP50 for 7 hours (Related to Figure 4). DESeq2 was used to identify differentially expressed genes in adr-1(-);adr-2(-) double mutant animals compared to wildtype animals. p-value<0.05 were taken as targets misregulated in absence of ad-1 and adr-2. Differentially expressed genes are listed as Wormbase annotations in Column A, Gene name in Column B. The log2fold change values, raw p values and the adjusted p values as obtained from the DESeq2 analysis are shown in columns C, D, and E, respectively. Read counts from the DESeq2 analysis for the three biological replicates of wildtype are listed in columns F, G and H. Read counts from the DESeq2 analysis for the three biological replicates of adr-1(-);adr-2(-) animals are listed in columns I, J and K . "	"Supplemental Table 4: Differentially expressed genes in adr-1(-);adr-2(-) animals grown on OP40 for 7 hours compared to wildtype animals grown on OP50 for 7 hours (Related to Figure 4). DESeq2 was used to identify differentially expressed genes in adr-1(-);adr-2(-) double mutant animals compared to wildtype animals. p-value<0.05 were taken as targets misregulated in absence of ad-1 and adr-2. Differentially expressed genes are listed as Wormbase annotations in Column A, Gene name in Column B. The log2fold change values, raw p values and the adjusted p values as obtained from the DESeq2 analysis are shown in columns C, D, and E, respectively. Read counts from the DESeq2 analysis for the three biological replicates of wildtype are listed in columns F, G and H. Read counts from the DESeq2 analysis for the three biological replicates of adr-1(-);adr-2(-) animals are listed in columns I, J and K . "	"Supplemental Table 4: Differentially expressed genes in adr-1(-);adr-2(-) animals grown on OP40 for 7 hours compared to wildtype animals grown on OP50 for 7 hours (Related to Figure 4). DESeq2 was used to identify differentially expressed genes in adr-1(-);adr-2(-) double mutant animals compared to wildtype animals. p-value<0.05 were taken as targets misregulated in absence of ad-1 and adr-2. Differentially expressed genes are listed as Wormbase annotations in Column A, Gene name in Column B. The log2fold change values, raw p values and the adjusted p values as obtained from the DESeq2 analysis are shown in columns C, D, and E, respectively. Read counts from the DESeq2 analysis for the three biological replicates of wildtype are listed in columns F, G and H. Read counts from the DESeq2 analysis for the three biological replicates of adr-1(-);adr-2(-) animals are listed in columns I, J and K . "	"Supplemental Table 4: Differentially expressed genes in adr-1(-);adr-2(-) animals grown on OP40 for 7 hours compared to wildtype animals grown on OP50 for 7 hours (Related to Figure 4). DESeq2 was used to identify differentially expressed genes in adr-1(-);adr-2(-) double mutant animals compared to wildtype animals. p-value<0.05 were taken as targets misregulated in absence of ad-1 and adr-2. Differentially expressed genes are listed as Wormbase annotations in Column A, Gene name in Column B. The log2fold change values, raw p values and the adjusted p values as obtained from the DESeq2 analysis are shown in columns C, D, and E, respectively. Read counts from the DESeq2 analysis for the three biological replicates of wildtype are listed in columns F, G and H. Read counts from the DESeq2 analysis for the three biological replicates of adr-1(-);adr-2(-) animals are listed in columns I, J and K . "	"Supplemental Table 4: Differentially expressed genes in adr-1(-);adr-2(-) animals grown on OP40 for 7 hours compared to wildtype animals grown on OP50 for 7 hours (Related to Figure 4). DESeq2 was used to identify differentially expressed genes in adr-1(-);adr-2(-) double mutant animals compared to wildtype animals. p-value<0.05 were taken as targets misregulated in absence of ad-1 and adr-2. Differentially expressed genes are listed as Wormbase annotations in Column A, Gene name in Column B. The log2fold change values, raw p values and the adjusted p values as obtained from the DESeq2 analysis are shown in columns C, D, and E, respectively. Read counts from the DESeq2 analysis for the three biological replicates of wildtype are listed in columns F, G and H. Read counts from the DESeq2 analysis for the three biological replicates of adr-1(-);adr-2(-) animals are listed in columns I, J and K . "	"Supplemental Table 4: Differentially expressed genes in adr-1(-);adr-2(-) animals grown on OP40 for 7 hours compared to wildtype animals grown on OP50 for 7 hours (Related to Figure 4). DESeq2 was used to identify differentially expressed genes in adr-1(-);adr-2(-) double mutant animals compared to wildtype animals. p-value<0.05 were taken as targets misregulated in absence of ad-1 and adr-2. Differentially expressed genes are listed as Wormbase annotations in Column A, Gene name in Column B. The log2fold change values, raw p values and the adjusted p values as obtained from the DESeq2 analysis are shown in columns C, D, and E, respectively. Read counts from the DESeq2 analysis for the three biological replicates of wildtype are listed in columns F, G and H. Read counts from the DESeq2 analysis for the three biological replicates of adr-1(-);adr-2(-) animals are listed in columns I, J and K . "	"Supplemental Table 4: Differentially expressed genes in adr-1(-);adr-2(-) animals grown on OP40 for 7 hours compared to wildtype animals grown on OP50 for 7 hours (Related to Figure 4). DESeq2 was used to identify differentially expressed genes in adr-1(-);adr-2(-) double mutant animals compared to wildtype animals. p-value<0.05 were taken as targets misregulated in absence of ad-1 and adr-2. Differentially expressed genes are listed as Wormbase annotations in Column A, Gene name in Column B. The log2fold change values, raw p values and the adjusted p values as obtained from the DESeq2 analysis are shown in columns C, D, and E, respectively. Read counts from the DESeq2 analysis for the three biological replicates of wildtype are listed in columns F, G and H. Read counts from the DESeq2 analysis for the three biological replicates of adr-1(-);adr-2(-) animals are listed in columns I, J and K . "	"Supplemental Table 4: Differentially expressed genes in adr-1(-);adr-2(-) animals grown on OP40 for 7 hours compared to wildtype animals grown on OP50 for 7 hours (Related to Figure 4). DESeq2 was used to identify differentially expressed genes in adr-1(-);adr-2(-) double mutant animals compared to wildtype animals. p-value<0.05 were taken as targets misregulated in absence of ad-1 and adr-2. Differentially expressed genes are listed as Wormbase annotations in Column A, Gene name in Column B. The log2fold change values, raw p values and the adjusted p values as obtained from the DESeq2 analysis are shown in columns C, D, and E, respectively. Read counts from the DESeq2 analysis for the three biological replicates of wildtype are listed in columns F, G and H. Read counts from the DESeq2 analysis for the three biological replicates of adr-1(-);adr-2(-) animals are listed in columns I, J and K . "	"Supplemental Table 4: Differentially expressed genes in adr-1(-);adr-2(-) animals grown on OP40 for 7 hours compared to wildtype animals grown on OP50 for 7 hours (Related to Figure 4). DESeq2 was used to identify differentially expressed genes in adr-1(-);adr-2(-) double mutant animals compared to wildtype animals. p-value<0.05 were taken as targets misregulated in absence of ad-1 and adr-2. Differentially expressed genes are listed as Wormbase annotations in Column A, Gene name in Column B. The log2fold change values, raw p values and the adjusted p values as obtained from the DESeq2 analysis are shown in columns C, D, and E, respectively. Read counts from the DESeq2 analysis for the three biological replicates of wildtype are listed in columns F, G and H. Read counts from the DESeq2 analysis for the three biological replicates of adr-1(-);adr-2(-) animals are listed in columns I, J and K . "	"Supplemental Table 4: Differentially expressed genes in adr-1(-);adr-2(-) animals grown on OP40 for 7 hours compared to wildtype animals grown on OP50 for 7 hours (Related to Figure 4). DESeq2 was used to identify differentially expressed genes in adr-1(-);adr-2(-) double mutant animals compared to wildtype animals. p-value<0.05 were taken as targets misregulated in absence of ad-1 and adr-2. Differentially expressed genes are listed as Wormbase annotations in Column A, Gene name in Column B. The log2fold change values, raw p values and the adjusted p values as obtained from the DESeq2 analysis are shown in columns C, D, and E, respectively. Read counts from the DESeq2 analysis for the three biological replicates of wildtype are listed in columns F, G and H. Read counts from the DESeq2 analysis for the three biological replicates of adr-1(-);adr-2(-) animals are listed in columns I, J and K . "	"Supplemental Table 4: Differentially expressed genes in adr-1(-);adr-2(-) animals grown on OP40 for 7 hours compared to wildtype animals grown on OP50 for 7 hours (Related to Figure 4). DESeq2 was used to identify differentially expressed genes in adr-1(-);adr-2(-) double mutant animals compared to wildtype animals. p-value<0.05 were taken as targets misregulated in absence of ad-1 and adr-2. Differentially expressed genes are listed as Wormbase annotations in Column A, Gene name in Column B. The log2fold change values, raw p values and the adjusted p values as obtained from the DESeq2 analysis are shown in columns C, D, and E, respectively. Read counts from the DESeq2 analysis for the three biological replicates of wildtype are listed in columns F, G and H. Read counts from the DESeq2 analysis for the three biological replicates of adr-1(-);adr-2(-) animals are listed in columns I, J and K . "	"Supplemental Table 4: Differentially expressed genes in adr-1(-);adr-2(-) animals grown on OP40 for 7 hours compared to wildtype animals grown on OP50 for 7 hours (Related to Figure 4). DESeq2 was used to identify differentially expressed genes in adr-1(-);adr-2(-) double mutant animals compared to wildtype animals. p-value<0.05 were taken as targets misregulated in absence of ad-1 and adr-2. Differentially expressed genes are listed as Wormbase annotations in Column A, Gene name in Column B. The log2fold change values, raw p values and the adjusted p values as obtained from the DESeq2 analysis are shown in columns C, D, and E, respectively. Read counts from the DESeq2 analysis for the three biological replicates of wildtype are listed in columns F, G and H. Read counts from the DESeq2 analysis for the three biological replicates of adr-1(-);adr-2(-) animals are listed in columns I, J and K . "	"Supplemental Table 4: Differentially expressed genes in adr-1(-);adr-2(-) animals grown on OP40 for 7 hours compared to wildtype animals grown on OP50 for 7 hours (Related to Figure 4). DESeq2 was used to identify differentially expressed genes in adr-1(-);adr-2(-) double mutant animals compared to wildtype animals. p-value<0.05 were taken as targets misregulated in absence of ad-1 and adr-2. Differentially expressed genes are listed as Wormbase annotations in Column A, Gene name in Column B. The log2fold change values, raw p values and the adjusted p values as obtained from the DESeq2 analysis are shown in columns C, D, and E, respectively. Read counts from the DESeq2 analysis for the three biological replicates of wildtype are listed in columns F, G and H. Read counts from the DESeq2 analysis for the three biological replicates of adr-1(-);adr-2(-) animals are listed in columns I, J and K . "	"Supplemental Table 4: Differentially expressed genes in adr-1(-);adr-2(-) animals grown on OP40 for 7 hours compared to wildtype animals grown on OP50 for 7 hours (Related to Figure 4). DESeq2 was used to identify differentially expressed genes in adr-1(-);adr-2(-) double mutant animals compared to wildtype animals. p-value<0.05 were taken as targets misregulated in absence of ad-1 and adr-2. Differentially expressed genes are listed as Wormbase annotations in Column A, Gene name in Column B. The log2fold change values, raw p values and the adjusted p values as obtained from the DESeq2 analysis are shown in columns C, D, and E, respectively. Read counts from the DESeq2 analysis for the three biological replicates of wildtype are listed in columns F, G and H. Read counts from the DESeq2 analysis for the three biological replicates of adr-1(-);adr-2(-) animals are listed in columns I, J and K . "	"Supplemental Table 4: Differentially expressed genes in adr-1(-);adr-2(-) animals grown on OP40 for 7 hours compared to wildtype animals grown on OP50 for 7 hours (Related to Figure 4). DESeq2 was used to identify differentially expressed genes in adr-1(-);adr-2(-) double mutant animals compared to wildtype animals. p-value<0.05 were taken as targets misregulated in absence of ad-1 and adr-2. Differentially expressed genes are listed as Wormbase annotations in Column A, Gene name in Column B. The log2fold change values, raw p values and the adjusted p values as obtained from the DESeq2 analysis are shown in columns C, D, and E, respectively. Read counts from the DESeq2 analysis for the three biological replicates of wildtype are listed in columns F, G and H. Read counts from the DESeq2 analysis for the three biological replicates of adr-1(-);adr-2(-) animals are listed in columns I, J and K . "	"Supplemental Table 4: Differentially expressed genes in adr-1(-);adr-2(-) animals grown on OP40 for 7 hours compared to wildtype animals grown on OP50 for 7 hours (Related to Figure 4). DESeq2 was used to identify differentially expressed genes in adr-1(-);adr-2(-) double mutant animals compared to wildtype animals. p-value<0.05 were taken as targets misregulated in absence of ad-1 and adr-2. Differentially expressed genes are listed as Wormbase annotations in Column A, Gene name in Column B. The log2fold change values, raw p values and the adjusted p values as obtained from the DESeq2 analysis are shown in columns C, D, and E, respectively. Read counts from the DESeq2 analysis for the three biological replicates of wildtype are listed in columns F, G and H. Read counts from the DESeq2 analysis for the three biological replicates of adr-1(-);adr-2(-) animals are listed in columns I, J and K . "	"Supplemental Table 4: Differentially expressed genes in adr-1(-);adr-2(-) animals grown on OP40 for 7 hours compared to wildtype animals grown on OP50 for 7 hours (Related to Figure 4). DESeq2 was used to identify differentially expressed genes in adr-1(-);adr-2(-) double mutant animals compared to wildtype animals. p-value<0.05 were taken as targets misregulated in absence of ad-1 and adr-2. Differentially expressed genes are listed as Wormbase annotations in Column A, Gene name in Column B. The log2fold change values, raw p values and the adjusted p values as obtained from the DESeq2 analysis are shown in columns C, D, and E, respectively. Read counts from the DESeq2 analysis for the three biological replicates of wildtype are listed in columns F, G and H. Read counts from the DESeq2 analysis for the three biological replicates of adr-1(-);adr-2(-) animals are listed in columns I, J and K . "	"Supplemental Table 4: Differentially expressed genes in adr-1(-);adr-2(-) animals grown on OP40 for 7 hours compared to wildtype animals grown on OP50 for 7 hours (Related to Figure 4). DESeq2 was used to identify differentially expressed genes in adr-1(-);adr-2(-) double mutant animals compared to wildtype animals. p-value<0.05 were taken as targets misregulated in absence of ad-1 and adr-2. Differentially expressed genes are listed as Wormbase annotations in Column A, Gene name in Column B. The log2fold change values, raw p values and the adjusted p values as obtained from the DESeq2 analysis are shown in columns C, D, and E, respectively. Read counts from the DESeq2 analysis for the three biological replicates of wildtype are listed in columns F, G and H. Read counts from the DESeq2 analysis for the three biological replicates of adr-1(-);adr-2(-) animals are listed in columns I, J and K . "	"Supplemental Table 4: Differentially expressed genes in adr-1(-);adr-2(-) animals grown on OP40 for 7 hours compared to wildtype animals grown on OP50 for 7 hours (Related to Figure 4). DESeq2 was used to identify differentially expressed genes in adr-1(-);adr-2(-) double mutant animals compared to wildtype animals. p-value<0.05 were taken as targets misregulated in absence of ad-1 and adr-2. Differentially expressed genes are listed as Wormbase annotations in Column A, Gene name in Column B. The log2fold change values, raw p values and the adjusted p values as obtained from the DESeq2 analysis are shown in columns C, D, and E, respectively. Read counts from the DESeq2 analysis for the three biological replicates of wildtype are listed in columns F, G and H. Read counts from the DESeq2 analysis for the three biological replicates of adr-1(-);adr-2(-) animals are listed in columns I, J and K . "	"Supplemental Table 4: Differentially expressed genes in adr-1(-);adr-2(-) animals grown on OP40 for 7 hours compared to wildtype animals grown on OP50 for 7 hours (Related to Figure 4). DESeq2 was used to identify differentially expressed genes in adr-1(-);adr-2(-) double mutant animals compared to wildtype animals. p-value<0.05 were taken as targets misregulated in absence of ad-1 and adr-2. Differentially expressed genes are listed as Wormbase annotations in Column A, Gene name in Column B. The log2fold change values, raw p values and the adjusted p values as obtained from the DESeq2 analysis are shown in columns C, D, and E, respectively. Read counts from the DESeq2 analysis for the three biological replicates of wildtype are listed in columns F, G and H. Read counts from the DESeq2 analysis for the three biological replicates of adr-1(-);adr-2(-) animals are listed in columns I, J and K . "	"Supplemental Table 4: Differentially expressed genes in adr-1(-);adr-2(-) animals grown on OP40 for 7 hours compared to wildtype animals grown on OP50 for 7 hours (Related to Figure 4). DESeq2 was used to identify differentially expressed genes in adr-1(-);adr-2(-) double mutant animals compared to wildtype animals. p-value<0.05 were taken as targets misregulated in absence of ad-1 and adr-2. Differentially expressed genes are listed as Wormbase annotations in Column A, Gene name in Column B. The log2fold change values, raw p values and the adjusted p values as obtained from the DESeq2 analysis are shown in columns C, D, and E, respectively. Read counts from the DESeq2 analysis for the three biological replicates of wildtype are listed in columns F, G and H. Read counts from the DESeq2 analysis for the three biological replicates of adr-1(-);adr-2(-) animals are listed in columns I, J and K . "	"Supplemental Table 4: Differentially expressed genes in adr-1(-);adr-2(-) animals grown on OP40 for 7 hours compared to wildtype animals grown on OP50 for 7 hours (Related to Figure 4). DESeq2 was used to identify differentially expressed genes in adr-1(-);adr-2(-) double mutant animals compared to wildtype animals. p-value<0.05 were taken as targets misregulated in absence of ad-1 and adr-2. Differentially expressed genes are listed as Wormbase annotations in Column A, Gene name in Column B. The log2fold change values, raw p values and the adjusted p values as obtained from the DESeq2 analysis are shown in columns C, D, and E, respectively. Read counts from the DESeq2 analysis for the three biological replicates of wildtype are listed in columns F, G and H. Read counts from the DESeq2 analysis for the three biological replicates of adr-1(-);adr-2(-) animals are listed in columns I, J and K . "	"Supplemental Table 4: Differentially expressed genes in adr-1(-);adr-2(-) animals grown on OP40 for 7 hours compared to wildtype animals grown on OP50 for 7 hours (Related to Figure 4). DESeq2 was used to identify differentially expressed genes in adr-1(-);adr-2(-) double mutant animals compared to wildtype animals. p-value<0.05 were taken as targets misregulated in absence of ad-1 and adr-2. Differentially expressed genes are listed as Wormbase annotations in Column A, Gene name in Column B. The log2fold change values, raw p values and the adjusted p values as obtained from the DESeq2 analysis are shown in columns C, D, and E, respectively. Read counts from the DESeq2 analysis for the three biological replicates of wildtype are listed in columns F, G and H. Read counts from the DESeq2 analysis for the three biological replicates of adr-1(-);adr-2(-) animals are listed in columns I, J and K . "	"Supplemental Table 4: Differentially expressed genes in adr-1(-);adr-2(-) animals grown on OP40 for 7 hours compared to wildtype animals grown on OP50 for 7 hours (Related to Figure 4). DESeq2 was used to identify differentially expressed genes in adr-1(-);adr-2(-) double mutant animals compared to wildtype animals. p-value<0.05 were taken as targets misregulated in absence of ad-1 and adr-2. Differentially expressed genes are listed as Wormbase annotations in Column A, Gene name in Column B. The log2fold change values, raw p values and the adjusted p values as obtained from the DESeq2 analysis are shown in columns C, D, and E, respectively. Read counts from the DESeq2 analysis for the three biological replicates of wildtype are listed in columns F, G and H. Read counts from the DESeq2 analysis for the three biological replicates of adr-1(-);adr-2(-) animals are listed in columns I, J and K . "	"Supplemental Table 4: Differentially expressed genes in adr-1(-);adr-2(-) animals grown on OP40 for 7 hours compared to wildtype animals grown on OP50 for 7 hours (Related to Figure 4). DESeq2 was used to identify differentially expressed genes in adr-1(-);adr-2(-) double mutant animals compared to wildtype animals. p-value<0.05 were taken as targets misregulated in absence of ad-1 and adr-2. Differentially expressed genes are listed as Wormbase annotations in Column A, Gene name in Column B. The log2fold change values, raw p values and the adjusted p values as obtained from the DESeq2 analysis are shown in columns C, D, and E, respectively. Read counts from the DESeq2 analysis for the three biological replicates of wildtype are listed in columns F, G and H. Read counts from the DESeq2 analysis for the three biological replicates of adr-1(-);adr-2(-) animals are listed in columns I, J and K . "	"Supplemental Table 4: Differentially expressed genes in adr-1(-);adr-2(-) animals grown on OP40 for 7 hours compared to wildtype animals grown on OP50 for 7 hours (Related to Figure 4). DESeq2 was used to identify differentially expressed genes in adr-1(-);adr-2(-) double mutant animals compared to wildtype animals. p-value<0.05 were taken as targets misregulated in absence of ad-1 and adr-2. Differentially expressed genes are listed as Wormbase annotations in Column A, Gene name in Column B. The log2fold change values, raw p values and the adjusted p values as obtained from the DESeq2 analysis are shown in columns C, D, and E, respectively. Read counts from the DESeq2 analysis for the three biological replicates of wildtype are listed in columns F, G and H. Read counts from the DESeq2 analysis for the three biological replicates of adr-1(-);adr-2(-) animals are listed in columns I, J and K . "	"Supplemental Table 4: Differentially expressed genes in adr-1(-);adr-2(-) animals grown on OP40 for 7 hours compared to wildtype animals grown on OP50 for 7 hours (Related to Figure 4). DESeq2 was used to identify differentially expressed genes in adr-1(-);adr-2(-) double mutant animals compared to wildtype animals. p-value<0.05 were taken as targets misregulated in absence of ad-1 and adr-2. Differentially expressed genes are listed as Wormbase annotations in Column A, Gene name in Column B. The log2fold change values, raw p values and the adjusted p values as obtained from the DESeq2 analysis are shown in columns C, D, and E, respectively. Read counts from the DESeq2 analysis for the three biological replicates of wildtype are listed in columns F, G and H. Read counts from the DESeq2 analysis for the three biological replicates of adr-1(-);adr-2(-) animals are listed in columns I, J and K . "	"Supplemental Table 4: Differentially expressed genes in adr-1(-);adr-2(-) animals grown on OP40 for 7 hours compared to wildtype animals grown on OP50 for 7 hours (Related to Figure 4). DESeq2 was used to identify differentially expressed genes in adr-1(-);adr-2(-) double mutant animals compared to wildtype animals. p-value<0.05 were taken as targets misregulated in absence of ad-1 and adr-2. Differentially expressed genes are listed as Wormbase annotations in Column A, Gene name in Column B. The log2fold change values, raw p values and the adjusted p values as obtained from the DESeq2 analysis are shown in columns C, D, and E, respectively. Read counts from the DESeq2 analysis for the three biological replicates of wildtype are listed in columns F, G and H. Read counts from the DESeq2 analysis for the three biological replicates of adr-1(-);adr-2(-) animals are listed in columns I, J and K . "	"Supplemental Table 4: Differentially expressed genes in adr-1(-);adr-2(-) animals grown on OP40 for 7 hours compared to wildtype animals grown on OP50 for 7 hours (Related to Figure 4). DESeq2 was used to identify differentially expressed genes in adr-1(-);adr-2(-) double mutant animals compared to wildtype animals. p-value<0.05 were taken as targets misregulated in absence of ad-1 and adr-2. Differentially expressed genes are listed as Wormbase annotations in Column A, Gene name in Column B. The log2fold change values, raw p values and the adjusted p values as obtained from the DESeq2 analysis are shown in columns C, D, and E, respectively. Read counts from the DESeq2 analysis for the three biological replicates of wildtype are listed in columns F, G and H. Read counts from the DESeq2 analysis for the three biological replicates of adr-1(-);adr-2(-) animals are listed in columns I, J and K . "	"Supplemental Table 4: Differentially expressed genes in adr-1(-);adr-2(-) animals grown on OP40 for 7 hours compared to wildtype animals grown on OP50 for 7 hours (Related to Figure 4). DESeq2 was used to identify differentially expressed genes in adr-1(-);adr-2(-) double mutant animals compared to wildtype animals. p-value<0.05 were taken as targets misregulated in absence of ad-1 and adr-2. Differentially expressed genes are listed as Wormbase annotations in Column A, Gene name in Column B. The log2fold change values, raw p values and the adjusted p values as obtained from the DESeq2 analysis are shown in columns C, D, and E, respectively. Read counts from the DESeq2 analysis for the three biological replicates of wildtype are listed in columns F, G and H. Read counts from the DESeq2 analysis for the three biological replicates of adr-1(-);adr-2(-) animals are listed in columns I, J and K . "	"Supplemental Table 4: Differentially expressed genes in adr-1(-);adr-2(-) animals grown on OP40 for 7 hours compared to wildtype animals grown on OP50 for 7 hours (Related to Figure 4). DESeq2 was used to identify differentially expressed genes in adr-1(-);adr-2(-) double mutant animals compared to wildtype animals. p-value<0.05 were taken as targets misregulated in absence of ad-1 and adr-2. Differentially expressed genes are listed as Wormbase annotations in Column A, Gene name in Column B. The log2fold change values, raw p values and the adjusted p values as obtained from the DESeq2 analysis are shown in columns C, D, and E, respectively. Read counts from the DESeq2 analysis for the three biological replicates of wildtype are listed in columns F, G and H. Read counts from the DESeq2 analysis for the three biological replicates of adr-1(-);adr-2(-) animals are listed in columns I, J and K . "	"Supplemental Table 4: Differentially expressed genes in adr-1(-);adr-2(-) animals grown on OP40 for 7 hours compared to wildtype animals grown on OP50 for 7 hours (Related to Figure 4). DESeq2 was used to identify differentially expressed genes in adr-1(-);adr-2(-) double mutant animals compared to wildtype animals. p-value<0.05 were taken as targets misregulated in absence of ad-1 and adr-2. Differentially expressed genes are listed as Wormbase annotations in Column A, Gene name in Column B. The log2fold change values, raw p values and the adjusted p values as obtained from the DESeq2 analysis are shown in columns C, D, and E, respectively. Read counts from the DESeq2 analysis for the three biological replicates of wildtype are listed in columns F, G and H. Read counts from the DESeq2 analysis for the three biological replicates of adr-1(-);adr-2(-) animals are listed in columns I, J and K . "	"Supplemental Table 4: Differentially expressed genes in adr-1(-);adr-2(-) animals grown on OP40 for 7 hours compared to wildtype animals grown on OP50 for 7 hours (Related to Figure 4). DESeq2 was used to identify differentially expressed genes in adr-1(-);adr-2(-) double mutant animals compared to wildtype animals. p-value<0.05 were taken as targets misregulated in absence of ad-1 and adr-2. Differentially expressed genes are listed as Wormbase annotations in Column A, Gene name in Column B. The log2fold change values, raw p values and the adjusted p values as obtained from the DESeq2 analysis are shown in columns C, D, and E, respectively. Read counts from the DESeq2 analysis for the three biological replicates of wildtype are listed in columns F, G and H. Read counts from the DESeq2 analysis for the three biological replicates of adr-1(-);adr-2(-) animals are listed in columns I, J and K . "	"Supplemental Table 4: Differentially expressed genes in adr-1(-);adr-2(-) animals grown on OP40 for 7 hours compared to wildtype animals grown on OP50 for 7 hours (Related to Figure 4). DESeq2 was used to identify differentially expressed genes in adr-1(-);adr-2(-) double mutant animals compared to wildtype animals. p-value<0.05 were taken as targets misregulated in absence of ad-1 and adr-2. Differentially expressed genes are listed as Wormbase annotations in Column A, Gene name in Column B. The log2fold change values, raw p values and the adjusted p values as obtained from the DESeq2 analysis are shown in columns C, D, and E, respectively. Read counts from the DESeq2 analysis for the three biological replicates of wildtype are listed in columns F, G and H. Read counts from the DESeq2 analysis for the three biological replicates of adr-1(-);adr-2(-) animals are listed in columns I, J and K . "	"Supplemental Table 4: Differentially expressed genes in adr-1(-);adr-2(-) animals grown on OP40 for 7 hours compared to wildtype animals grown on OP50 for 7 hours (Related to Figure 4). DESeq2 was used to identify differentially expressed genes in adr-1(-);adr-2(-) double mutant animals compared to wildtype animals. p-value<0.05 were taken as targets misregulated in absence of ad-1 and adr-2. Differentially expressed genes are listed as Wormbase annotations in Column A, Gene name in Column B. The log2fold change values, raw p values and the adjusted p values as obtained from the DESeq2 analysis are shown in columns C, D, and E, respectively. Read counts from the DESeq2 analysis for the three biological replicates of wildtype are listed in columns F, G and H. Read counts from the DESeq2 analysis for the three biological replicates of adr-1(-);adr-2(-) animals are listed in columns I, J and K . "	"Supplemental Table 4: Differentially expressed genes in adr-1(-);adr-2(-) animals grown on OP40 for 7 hours compared to wildtype animals grown on OP50 for 7 hours (Related to Figure 4). DESeq2 was used to identify differentially expressed genes in adr-1(-);adr-2(-) double mutant animals compared to wildtype animals. p-value<0.05 were taken as targets misregulated in absence of ad-1 and adr-2. Differentially expressed genes are listed as Wormbase annotations in Column A, Gene name in Column B. The log2fold change values, raw p values and the adjusted p values as obtained from the DESeq2 analysis are shown in columns C, D, and E, respectively. Read counts from the DESeq2 analysis for the three biological replicates of wildtype are listed in columns F, G and H. Read counts from the DESeq2 analysis for the three biological replicates of adr-1(-);adr-2(-) animals are listed in columns I, J and K . "	"Supplemental Table 4: Differentially expressed genes in adr-1(-);adr-2(-) animals grown on OP40 for 7 hours compared to wildtype animals grown on OP50 for 7 hours (Related to Figure 4). DESeq2 was used to identify differentially expressed genes in adr-1(-);adr-2(-) double mutant animals compared to wildtype animals. p-value<0.05 were taken as targets misregulated in absence of ad-1 and adr-2. Differentially expressed genes are listed as Wormbase annotations in Column A, Gene name in Column B. The log2fold change values, raw p values and the adjusted p values as obtained from the DESeq2 analysis are shown in columns C, D, and E, respectively. Read counts from the DESeq2 analysis for the three biological replicates of wildtype are listed in columns F, G and H. Read counts from the DESeq2 analysis for the three biological replicates of adr-1(-);adr-2(-) animals are listed in columns I, J and K . "	"Supplemental Table 4: Differentially expressed genes in adr-1(-);adr-2(-) animals grown on OP40 for 7 hours compared to wildtype animals grown on OP50 for 7 hours (Related to Figure 4). DESeq2 was used to identify differentially expressed genes in adr-1(-);adr-2(-) double mutant animals compared to wildtype animals. p-value<0.05 were taken as targets misregulated in absence of ad-1 and adr-2. Differentially expressed genes are listed as Wormbase annotations in Column A, Gene name in Column B. The log2fold change values, raw p values and the adjusted p values as obtained from the DESeq2 analysis are shown in columns C, D, and E, respectively. Read counts from the DESeq2 analysis for the three biological replicates of wildtype are listed in columns F, G and H. Read counts from the DESeq2 analysis for the three biological replicates of adr-1(-);adr-2(-) animals are listed in columns I, J and K . "	"Supplemental Table 4: Differentially expressed genes in adr-1(-);adr-2(-) animals grown on OP40 for 7 hours compared to wildtype animals grown on OP50 for 7 hours (Related to Figure 4). DESeq2 was used to identify differentially expressed genes in adr-1(-);adr-2(-) double mutant animals compared to wildtype animals. p-value<0.05 were taken as targets misregulated in absence of ad-1 and adr-2. Differentially expressed genes are listed as Wormbase annotations in Column A, Gene name in Column B. The log2fold change values, raw p values and the adjusted p values as obtained from the DESeq2 analysis are shown in columns C, D, and E, respectively. Read counts from the DESeq2 analysis for the three biological replicates of wildtype are listed in columns F, G and H. Read counts from the DESeq2 analysis for the three biological replicates of adr-1(-);adr-2(-) animals are listed in columns I, J and K . "	"Supplemental Table 4: Differentially expressed genes in adr-1(-);adr-2(-) animals grown on OP40 for 7 hours compared to wildtype animals grown on OP50 for 7 hours (Related to Figure 4). DESeq2 was used to identify differentially expressed genes in adr-1(-);adr-2(-) double mutant animals compared to wildtype animals. p-value<0.05 were taken as targets misregulated in absence of ad-1 and adr-2. Differentially expressed genes are listed as Wormbase annotations in Column A, Gene name in Column B. The log2fold change values, raw p values and the adjusted p values as obtained from the DESeq2 analysis are shown in columns C, D, and E, respectively. Read counts from the DESeq2 analysis for the three biological replicates of wildtype are listed in columns F, G and H. Read counts from the DESeq2 analysis for the three biological replicates of adr-1(-);adr-2(-) animals are listed in columns I, J and K . "	"Supplemental Table 4: Differentially expressed genes in adr-1(-);adr-2(-) animals grown on OP40 for 7 hours compared to wildtype animals grown on OP50 for 7 hours (Related to Figure 4). DESeq2 was used to identify differentially expressed genes in adr-1(-);adr-2(-) double mutant animals compared to wildtype animals. p-value<0.05 were taken as targets misregulated in absence of ad-1 and adr-2. Differentially expressed genes are listed as Wormbase annotations in Column A, Gene name in Column B. The log2fold change values, raw p values and the adjusted p values as obtained from the DESeq2 analysis are shown in columns C, D, and E, respectively. Read counts from the DESeq2 analysis for the three biological replicates of wildtype are listed in columns F, G and H. Read counts from the DESeq2 analysis for the three biological replicates of adr-1(-);adr-2(-) animals are listed in columns I, J and K . "	"Supplemental Table 4: Differentially expressed genes in adr-1(-);adr-2(-) animals grown on OP40 for 7 hours compared to wildtype animals grown on OP50 for 7 hours (Related to Figure 4). DESeq2 was used to identify differentially expressed genes in adr-1(-);adr-2(-) double mutant animals compared to wildtype animals. p-value<0.05 were taken as targets misregulated in absence of ad-1 and adr-2. Differentially expressed genes are listed as Wormbase annotations in Column A, Gene name in Column B. The log2fold change values, raw p values and the adjusted p values as obtained from the DESeq2 analysis are shown in columns C, D, and E, respectively. Read counts from the DESeq2 analysis for the three biological replicates of wildtype are listed in columns F, G and H. Read counts from the DESeq2 analysis for the three biological replicates of adr-1(-);adr-2(-) animals are listed in columns I, J and K . "	"Supplemental Table 4: Differentially expressed genes in adr-1(-);adr-2(-) animals grown on OP40 for 7 hours compared to wildtype animals grown on OP50 for 7 hours (Related to Figure 4). DESeq2 was used to identify differentially expressed genes in adr-1(-);adr-2(-) double mutant animals compared to wildtype animals. p-value<0.05 were taken as targets misregulated in absence of ad-1 and adr-2. Differentially expressed genes are listed as Wormbase annotations in Column A, Gene name in Column B. The log2fold change values, raw p values and the adjusted p values as obtained from the DESeq2 analysis are shown in columns C, D, and E, respectively. Read counts from the DESeq2 analysis for the three biological replicates of wildtype are listed in columns F, G and H. Read counts from the DESeq2 analysis for the three biological replicates of adr-1(-);adr-2(-) animals are listed in columns I, J and K . "	"Supplemental Table 4: Differentially expressed genes in adr-1(-);adr-2(-) animals grown on OP40 for 7 hours compared to wildtype animals grown on OP50 for 7 hours (Related to Figure 4). DESeq2 was used to identify differentially expressed genes in adr-1(-);adr-2(-) double mutant animals compared to wildtype animals. p-value<0.05 were taken as targets misregulated in absence of ad-1 and adr-2. Differentially expressed genes are listed as Wormbase annotations in Column A, Gene name in Column B. The log2fold change values, raw p values and the adjusted p values as obtained from the DESeq2 analysis are shown in columns C, D, and E, respectively. Read counts from the DESeq2 analysis for the three biological replicates of wildtype are listed in columns F, G and H. Read counts from the DESeq2 analysis for the three biological replicates of adr-1(-);adr-2(-) animals are listed in columns I, J and K . "	"Supplemental Table 4: Differentially expressed genes in adr-1(-);adr-2(-) animals grown on OP40 for 7 hours compared to wildtype animals grown on OP50 for 7 hours (Related to Figure 4). DESeq2 was used to identify differentially expressed genes in adr-1(-);adr-2(-) double mutant animals compared to wildtype animals. p-value<0.05 were taken as targets misregulated in absence of ad-1 and adr-2. Differentially expressed genes are listed as Wormbase annotations in Column A, Gene name in Column B. The log2fold change values, raw p values and the adjusted p values as obtained from the DESeq2 analysis are shown in columns C, D, and E, respectively. Read counts from the DESeq2 analysis for the three biological replicates of wildtype are listed in columns F, G and H. Read counts from the DESeq2 analysis for the three biological replicates of adr-1(-);adr-2(-) animals are listed in columns I, J and K . "	"Supplemental Table 4: Differentially expressed genes in adr-1(-);adr-2(-) animals grown on OP40 for 7 hours compared to wildtype animals grown on OP50 for 7 hours (Related to Figure 4). DESeq2 was used to identify differentially expressed genes in adr-1(-);adr-2(-) double mutant animals compared to wildtype animals. p-value<0.05 were taken as targets misregulated in absence of ad-1 and adr-2. Differentially expressed genes are listed as Wormbase annotations in Column A, Gene name in Column B. The log2fold change values, raw p values and the adjusted p values as obtained from the DESeq2 analysis are shown in columns C, D, and E, respectively. Read counts from the DESeq2 analysis for the three biological replicates of wildtype are listed in columns F, G and H. Read counts from the DESeq2 analysis for the three biological replicates of adr-1(-);adr-2(-) animals are listed in columns I, J and K . "	"Supplemental Table 4: Differentially expressed genes in adr-1(-);adr-2(-) animals grown on OP40 for 7 hours compared to wildtype animals grown on OP50 for 7 hours (Related to Figure 4). DESeq2 was used to identify differentially expressed genes in adr-1(-);adr-2(-) double mutant animals compared to wildtype animals. p-value<0.05 were taken as targets misregulated in absence of ad-1 and adr-2. Differentially expressed genes are listed as Wormbase annotations in Column A, Gene name in Column B. The log2fold change values, raw p values and the adjusted p values as obtained from the DESeq2 analysis are shown in columns C, D, and E, respectively. Read counts from the DESeq2 analysis for the three biological replicates of wildtype are listed in columns F, G and H. Read counts from the DESeq2 analysis for the three biological replicates of adr-1(-);adr-2(-) animals are listed in columns I, J and K . "	"Supplemental Table 4: Differentially expressed genes in adr-1(-);adr-2(-) animals grown on OP40 for 7 hours compared to wildtype animals grown on OP50 for 7 hours (Related to Figure 4). DESeq2 was used to identify differentially expressed genes in adr-1(-);adr-2(-) double mutant animals compared to wildtype animals. p-value<0.05 were taken as targets misregulated in absence of ad-1 and adr-2. Differentially expressed genes are listed as Wormbase annotations in Column A, Gene name in Column B. The log2fold change values, raw p values and the adjusted p values as obtained from the DESeq2 analysis are shown in columns C, D, and E, respectively. Read counts from the DESeq2 analysis for the three biological replicates of wildtype are listed in columns F, G and H. Read counts from the DESeq2 analysis for the three biological replicates of adr-1(-);adr-2(-) animals are listed in columns I, J and K . "	"Supplemental Table 4: Differentially expressed genes in adr-1(-);adr-2(-) animals grown on OP40 for 7 hours compared to wildtype animals grown on OP50 for 7 hours (Related to Figure 4). DESeq2 was used to identify differentially expressed genes in adr-1(-);adr-2(-) double mutant animals compared to wildtype animals. p-value<0.05 were taken as targets misregulated in absence of ad-1 and adr-2. Differentially expressed genes are listed as Wormbase annotations in Column A, Gene name in Column B. The log2fold change values, raw p values and the adjusted p values as obtained from the DESeq2 analysis are shown in columns C, D, and E, respectively. Read counts from the DESeq2 analysis for the three biological replicates of wildtype are listed in columns F, G and H. Read counts from the DESeq2 analysis for the three biological replicates of adr-1(-);adr-2(-) animals are listed in columns I, J and K . "	"Supplemental Table 4: Differentially expressed genes in adr-1(-);adr-2(-) animals grown on OP40 for 7 hours compared to wildtype animals grown on OP50 for 7 hours (Related to Figure 4). DESeq2 was used to identify differentially expressed genes in adr-1(-);adr-2(-) double mutant animals compared to wildtype animals. p-value<0.05 were taken as targets misregulated in absence of ad-1 and adr-2. Differentially expressed genes are listed as Wormbase annotations in Column A, Gene name in Column B. The log2fold change values, raw p values and the adjusted p values as obtained from the DESeq2 analysis are shown in columns C, D, and E, respectively. Read counts from the DESeq2 analysis for the three biological replicates of wildtype are listed in columns F, G and H. Read counts from the DESeq2 analysis for the three biological replicates of adr-1(-);adr-2(-) animals are listed in columns I, J and K . "	"Supplemental Table 4: Differentially expressed genes in adr-1(-);adr-2(-) animals grown on OP40 for 7 hours compared to wildtype animals grown on OP50 for 7 hours (Related to Figure 4). DESeq2 was used to identify differentially expressed genes in adr-1(-);adr-2(-) double mutant animals compared to wildtype animals. p-value<0.05 were taken as targets misregulated in absence of ad-1 and adr-2. Differentially expressed genes are listed as Wormbase annotations in Column A, Gene name in Column B. The log2fold change values, raw p values and the adjusted p values as obtained from the DESeq2 analysis are shown in columns C, D, and E, respectively. Read counts from the DESeq2 analysis for the three biological replicates of wildtype are listed in columns F, G and H. Read counts from the DESeq2 analysis for the three biological replicates of adr-1(-);adr-2(-) animals are listed in columns I, J and K . "	"Supplemental Table 4: Differentially expressed genes in adr-1(-);adr-2(-) animals grown on OP40 for 7 hours compared to wildtype animals grown on OP50 for 7 hours (Related to Figure 4). DESeq2 was used to identify differentially expressed genes in adr-1(-);adr-2(-) double mutant animals compared to wildtype animals. p-value<0.05 were taken as targets misregulated in absence of ad-1 and adr-2. Differentially expressed genes are listed as Wormbase annotations in Column A, Gene name in Column B. The log2fold change values, raw p values and the adjusted p values as obtained from the DESeq2 analysis are shown in columns C, D, and E, respectively. Read counts from the DESeq2 analysis for the three biological replicates of wildtype are listed in columns F, G and H. Read counts from the DESeq2 analysis for the three biological replicates of adr-1(-);adr-2(-) animals are listed in columns I, J and K . "	"Supplemental Table 4: Differentially expressed genes in adr-1(-);adr-2(-) animals grown on OP40 for 7 hours compared to wildtype animals grown on OP50 for 7 hours (Related to Figure 4). DESeq2 was used to identify differentially expressed genes in adr-1(-);adr-2(-) double mutant animals compared to wildtype animals. p-value<0.05 were taken as targets misregulated in absence of ad-1 and adr-2. Differentially expressed genes are listed as Wormbase annotations in Column A, Gene name in Column B. The log2fold change values, raw p values and the adjusted p values as obtained from the DESeq2 analysis are shown in columns C, D, and E, respectively. Read counts from the DESeq2 analysis for the three biological replicates of wildtype are listed in columns F, G and H. Read counts from the DESeq2 analysis for the three biological replicates of adr-1(-);adr-2(-) animals are listed in columns I, J and K . "	"Supplemental Table 4: Differentially expressed genes in adr-1(-);adr-2(-) animals grown on OP40 for 7 hours compared to wildtype animals grown on OP50 for 7 hours (Related to Figure 4). DESeq2 was used to identify differentially expressed genes in adr-1(-);adr-2(-) double mutant animals compared to wildtype animals. p-value<0.05 were taken as targets misregulated in absence of ad-1 and adr-2. Differentially expressed genes are listed as Wormbase annotations in Column A, Gene name in Column B. The log2fold change values, raw p values and the adjusted p values as obtained from the DESeq2 analysis are shown in columns C, D, and E, respectively. Read counts from the DESeq2 analysis for the three biological replicates of wildtype are listed in columns F, G and H. Read counts from the DESeq2 analysis for the three biological replicates of adr-1(-);adr-2(-) animals are listed in columns I, J and K . "	"Supplemental Table 4: Differentially expressed genes in adr-1(-);adr-2(-) animals grown on OP40 for 7 hours compared to wildtype animals grown on OP50 for 7 hours (Related to Figure 4). DESeq2 was used to identify differentially expressed genes in adr-1(-);adr-2(-) double mutant animals compared to wildtype animals. p-value<0.05 were taken as targets misregulated in absence of ad-1 and adr-2. Differentially expressed genes are listed as Wormbase annotations in Column A, Gene name in Column B. The log2fold change values, raw p values and the adjusted p values as obtained from the DESeq2 analysis are shown in columns C, D, and E, respectively. Read counts from the DESeq2 analysis for the three biological replicates of wildtype are listed in columns F, G and H. Read counts from the DESeq2 analysis for the three biological replicates of adr-1(-);adr-2(-) animals are listed in columns I, J and K . "	"Supplemental Table 4: Differentially expressed genes in adr-1(-);adr-2(-) animals grown on OP40 for 7 hours compared to wildtype animals grown on OP50 for 7 hours (Related to Figure 4). DESeq2 was used to identify differentially expressed genes in adr-1(-);adr-2(-) double mutant animals compared to wildtype animals. p-value<0.05 were taken as targets misregulated in absence of ad-1 and adr-2. Differentially expressed genes are listed as Wormbase annotations in Column A, Gene name in Column B. The log2fold change values, raw p values and the adjusted p values as obtained from the DESeq2 analysis are shown in columns C, D, and E, respectively. Read counts from the DESeq2 analysis for the three biological replicates of wildtype are listed in columns F, G and H. Read counts from the DESeq2 analysis for the three biological replicates of adr-1(-);adr-2(-) animals are listed in columns I, J and K . "	"Supplemental Table 4: Differentially expressed genes in adr-1(-);adr-2(-) animals grown on OP40 for 7 hours compared to wildtype animals grown on OP50 for 7 hours (Related to Figure 4). DESeq2 was used to identify differentially expressed genes in adr-1(-);adr-2(-) double mutant animals compared to wildtype animals. p-value<0.05 were taken as targets misregulated in absence of ad-1 and adr-2. Differentially expressed genes are listed as Wormbase annotations in Column A, Gene name in Column B. The log2fold change values, raw p values and the adjusted p values as obtained from the DESeq2 analysis are shown in columns C, D, and E, respectively. Read counts from the DESeq2 analysis for the three biological replicates of wildtype are listed in columns F, G and H. Read counts from the DESeq2 analysis for the three biological replicates of adr-1(-);adr-2(-) animals are listed in columns I, J and K . "	"Supplemental Table 4: Differentially expressed genes in adr-1(-);adr-2(-) animals grown on OP40 for 7 hours compared to wildtype animals grown on OP50 for 7 hours (Related to Figure 4). DESeq2 was used to identify differentially expressed genes in adr-1(-);adr-2(-) double mutant animals compared to wildtype animals. p-value<0.05 were taken as targets misregulated in absence of ad-1 and adr-2. Differentially expressed genes are listed as Wormbase annotations in Column A, Gene name in Column B. The log2fold change values, raw p values and the adjusted p values as obtained from the DESeq2 analysis are shown in columns C, D, and E, respectively. Read counts from the DESeq2 analysis for the three biological replicates of wildtype are listed in columns F, G and H. Read counts from the DESeq2 analysis for the three biological replicates of adr-1(-);adr-2(-) animals are listed in columns I, J and K . "	"Supplemental Table 4: Differentially expressed genes in adr-1(-);adr-2(-) animals grown on OP40 for 7 hours compared to wildtype animals grown on OP50 for 7 hours (Related to Figure 4). DESeq2 was used to identify differentially expressed genes in adr-1(-);adr-2(-) double mutant animals compared to wildtype animals. p-value<0.05 were taken as targets misregulated in absence of ad-1 and adr-2. Differentially expressed genes are listed as Wormbase annotations in Column A, Gene name in Column B. The log2fold change values, raw p values and the adjusted p values as obtained from the DESeq2 analysis are shown in columns C, D, and E, respectively. Read counts from the DESeq2 analysis for the three biological replicates of wildtype are listed in columns F, G and H. Read counts from the DESeq2 analysis for the three biological replicates of adr-1(-);adr-2(-) animals are listed in columns I, J and K . "	"Supplemental Table 4: Differentially expressed genes in adr-1(-);adr-2(-) animals grown on OP40 for 7 hours compared to wildtype animals grown on OP50 for 7 hours (Related to Figure 4). DESeq2 was used to identify differentially expressed genes in adr-1(-);adr-2(-) double mutant animals compared to wildtype animals. p-value<0.05 were taken as targets misregulated in absence of ad-1 and adr-2. Differentially expressed genes are listed as Wormbase annotations in Column A, Gene name in Column B. The log2fold change values, raw p values and the adjusted p values as obtained from the DESeq2 analysis are shown in columns C, D, and E, respectively. Read counts from the DESeq2 analysis for the three biological replicates of wildtype are listed in columns F, G and H. Read counts from the DESeq2 analysis for the three biological replicates of adr-1(-);adr-2(-) animals are listed in columns I, J and K . "	"Supplemental Table 4: Differentially expressed genes in adr-1(-);adr-2(-) animals grown on OP40 for 7 hours compared to wildtype animals grown on OP50 for 7 hours (Related to Figure 4). DESeq2 was used to identify differentially expressed genes in adr-1(-);adr-2(-) double mutant animals compared to wildtype animals. p-value<0.05 were taken as targets misregulated in absence of ad-1 and adr-2. Differentially expressed genes are listed as Wormbase annotations in Column A, Gene name in Column B. The log2fold change values, raw p values and the adjusted p values as obtained from the DESeq2 analysis are shown in columns C, D, and E, respectively. Read counts from the DESeq2 analysis for the three biological replicates of wildtype are listed in columns F, G and H. Read counts from the DESeq2 analysis for the three biological replicates of adr-1(-);adr-2(-) animals are listed in columns I, J and K . "	"Supplemental Table 4: Differentially expressed genes in adr-1(-);adr-2(-) animals grown on OP40 for 7 hours compared to wildtype animals grown on OP50 for 7 hours (Related to Figure 4). DESeq2 was used to identify differentially expressed genes in adr-1(-);adr-2(-) double mutant animals compared to wildtype animals. p-value<0.05 were taken as targets misregulated in absence of ad-1 and adr-2. Differentially expressed genes are listed as Wormbase annotations in Column A, Gene name in Column B. The log2fold change values, raw p values and the adjusted p values as obtained from the DESeq2 analysis are shown in columns C, D, and E, respectively. Read counts from the DESeq2 analysis for the three biological replicates of wildtype are listed in columns F, G and H. Read counts from the DESeq2 analysis for the three biological replicates of adr-1(-);adr-2(-) animals are listed in columns I, J and K . "	"Supplemental Table 4: Differentially expressed genes in adr-1(-);adr-2(-) animals grown on OP40 for 7 hours compared to wildtype animals grown on OP50 for 7 hours (Related to Figure 4). DESeq2 was used to identify differentially expressed genes in adr-1(-);adr-2(-) double mutant animals compared to wildtype animals. p-value<0.05 were taken as targets misregulated in absence of ad-1 and adr-2. Differentially expressed genes are listed as Wormbase annotations in Column A, Gene name in Column B. The log2fold change values, raw p values and the adjusted p values as obtained from the DESeq2 analysis are shown in columns C, D, and E, respectively. Read counts from the DESeq2 analysis for the three biological replicates of wildtype are listed in columns F, G and H. Read counts from the DESeq2 analysis for the three biological replicates of adr-1(-);adr-2(-) animals are listed in columns I, J and K . "	"Supplemental Table 4: Differentially expressed genes in adr-1(-);adr-2(-) animals grown on OP40 for 7 hours compared to wildtype animals grown on OP50 for 7 hours (Related to Figure 4). DESeq2 was used to identify differentially expressed genes in adr-1(-);adr-2(-) double mutant animals compared to wildtype animals. p-value<0.05 were taken as targets misregulated in absence of ad-1 and adr-2. Differentially expressed genes are listed as Wormbase annotations in Column A, Gene name in Column B. The log2fold change values, raw p values and the adjusted p values as obtained from the DESeq2 analysis are shown in columns C, D, and E, respectively. Read counts from the DESeq2 analysis for the three biological replicates of wildtype are listed in columns F, G and H. Read counts from the DESeq2 analysis for the three biological replicates of adr-1(-);adr-2(-) animals are listed in columns I, J and K . "	"Supplemental Table 4: Differentially expressed genes in adr-1(-);adr-2(-) animals grown on OP40 for 7 hours compared to wildtype animals grown on OP50 for 7 hours (Related to Figure 4). DESeq2 was used to identify differentially expressed genes in adr-1(-);adr-2(-) double mutant animals compared to wildtype animals. p-value<0.05 were taken as targets misregulated in absence of ad-1 and adr-2. Differentially expressed genes are listed as Wormbase annotations in Column A, Gene name in Column B. The log2fold change values, raw p values and the adjusted p values as obtained from the DESeq2 analysis are shown in columns C, D, and E, respectively. Read counts from the DESeq2 analysis for the three biological replicates of wildtype are listed in columns F, G and H. Read counts from the DESeq2 analysis for the three biological replicates of adr-1(-);adr-2(-) animals are listed in columns I, J and K . "	"Supplemental Table 4: Differentially expressed genes in adr-1(-);adr-2(-) animals grown on OP40 for 7 hours compared to wildtype animals grown on OP50 for 7 hours (Related to Figure 4). DESeq2 was used to identify differentially expressed genes in adr-1(-);adr-2(-) double mutant animals compared to wildtype animals. p-value<0.05 were taken as targets misregulated in absence of ad-1 and adr-2. Differentially expressed genes are listed as Wormbase annotations in Column A, Gene name in Column B. The log2fold change values, raw p values and the adjusted p values as obtained from the DESeq2 analysis are shown in columns C, D, and E, respectively. Read counts from the DESeq2 analysis for the three biological replicates of wildtype are listed in columns F, G and H. Read counts from the DESeq2 analysis for the three biological replicates of adr-1(-);adr-2(-) animals are listed in columns I, J and K . "	"Supplemental Table 4: Differentially expressed genes in adr-1(-);adr-2(-) animals grown on OP40 for 7 hours compared to wildtype animals grown on OP50 for 7 hours (Related to Figure 4). DESeq2 was used to identify differentially expressed genes in adr-1(-);adr-2(-) double mutant animals compared to wildtype animals. p-value<0.05 were taken as targets misregulated in absence of ad-1 and adr-2. Differentially expressed genes are listed as Wormbase annotations in Column A, Gene name in Column B. The log2fold change values, raw p values and the adjusted p values as obtained from the DESeq2 analysis are shown in columns C, D, and E, respectively. Read counts from the DESeq2 analysis for the three biological replicates of wildtype are listed in columns F, G and H. Read counts from the DESeq2 analysis for the three biological replicates of adr-1(-);adr-2(-) animals are listed in columns I, J and K . "	"Supplemental Table 4: Differentially expressed genes in adr-1(-);adr-2(-) animals grown on OP40 for 7 hours compared to wildtype animals grown on OP50 for 7 hours (Related to Figure 4). DESeq2 was used to identify differentially expressed genes in adr-1(-);adr-2(-) double mutant animals compared to wildtype animals. p-value<0.05 were taken as targets misregulated in absence of ad-1 and adr-2. Differentially expressed genes are listed as Wormbase annotations in Column A, Gene name in Column B. The log2fold change values, raw p values and the adjusted p values as obtained from the DESeq2 analysis are shown in columns C, D, and E, respectively. Read counts from the DESeq2 analysis for the three biological replicates of wildtype are listed in columns F, G and H. Read counts from the DESeq2 analysis for the three biological replicates of adr-1(-);adr-2(-) animals are listed in columns I, J and K . "	"Supplemental Table 4: Differentially expressed genes in adr-1(-);adr-2(-) animals grown on OP40 for 7 hours compared to wildtype animals grown on OP50 for 7 hours (Related to Figure 4). DESeq2 was used to identify differentially expressed genes in adr-1(-);adr-2(-) double mutant animals compared to wildtype animals. p-value<0.05 were taken as targets misregulated in absence of ad-1 and adr-2. Differentially expressed genes are listed as Wormbase annotations in Column A, Gene name in Column B. The log2fold change values, raw p values and the adjusted p values as obtained from the DESeq2 analysis are shown in columns C, D, and E, respectively. Read counts from the DESeq2 analysis for the three biological replicates of wildtype are listed in columns F, G and H. Read counts from the DESeq2 analysis for the three biological replicates of adr-1(-);adr-2(-) animals are listed in columns I, J and K . "	"Supplemental Table 4: Differentially expressed genes in adr-1(-);adr-2(-) animals grown on OP40 for 7 hours compared to wildtype animals grown on OP50 for 7 hours (Related to Figure 4). DESeq2 was used to identify differentially expressed genes in adr-1(-);adr-2(-) double mutant animals compared to wildtype animals. p-value<0.05 were taken as targets misregulated in absence of ad-1 and adr-2. Differentially expressed genes are listed as Wormbase annotations in Column A, Gene name in Column B. The log2fold change values, raw p values and the adjusted p values as obtained from the DESeq2 analysis are shown in columns C, D, and E, respectively. Read counts from the DESeq2 analysis for the three biological replicates of wildtype are listed in columns F, G and H. Read counts from the DESeq2 analysis for the three biological replicates of adr-1(-);adr-2(-) animals are listed in columns I, J and K . "	"Supplemental Table 4: Differentially expressed genes in adr-1(-);adr-2(-) animals grown on OP40 for 7 hours compared to wildtype animals grown on OP50 for 7 hours (Related to Figure 4). DESeq2 was used to identify differentially expressed genes in adr-1(-);adr-2(-) double mutant animals compared to wildtype animals. p-value<0.05 were taken as targets misregulated in absence of ad-1 and adr-2. Differentially expressed genes are listed as Wormbase annotations in Column A, Gene name in Column B. The log2fold change values, raw p values and the adjusted p values as obtained from the DESeq2 analysis are shown in columns C, D, and E, respectively. Read counts from the DESeq2 analysis for the three biological replicates of wildtype are listed in columns F, G and H. Read counts from the DESeq2 analysis for the three biological replicates of adr-1(-);adr-2(-) animals are listed in columns I, J and K . "	"Supplemental Table 4: Differentially expressed genes in adr-1(-);adr-2(-) animals grown on OP40 for 7 hours compared to wildtype animals grown on OP50 for 7 hours (Related to Figure 4). DESeq2 was used to identify differentially expressed genes in adr-1(-);adr-2(-) double mutant animals compared to wildtype animals. p-value<0.05 were taken as targets misregulated in absence of ad-1 and adr-2. Differentially expressed genes are listed as Wormbase annotations in Column A, Gene name in Column B. The log2fold change values, raw p values and the adjusted p values as obtained from the DESeq2 analysis are shown in columns C, D, and E, respectively. Read counts from the DESeq2 analysis for the three biological replicates of wildtype are listed in columns F, G and H. Read counts from the DESeq2 analysis for the three biological replicates of adr-1(-);adr-2(-) animals are listed in columns I, J and K . "	"Supplemental Table 4: Differentially expressed genes in adr-1(-);adr-2(-) animals grown on OP40 for 7 hours compared to wildtype animals grown on OP50 for 7 hours (Related to Figure 4). DESeq2 was used to identify differentially expressed genes in adr-1(-);adr-2(-) double mutant animals compared to wildtype animals. p-value<0.05 were taken as targets misregulated in absence of ad-1 and adr-2. Differentially expressed genes are listed as Wormbase annotations in Column A, Gene name in Column B. The log2fold change values, raw p values and the adjusted p values as obtained from the DESeq2 analysis are shown in columns C, D, and E, respectively. Read counts from the DESeq2 analysis for the three biological replicates of wildtype are listed in columns F, G and H. Read counts from the DESeq2 analysis for the three biological replicates of adr-1(-);adr-2(-) animals are listed in columns I, J and K . "	"Supplemental Table 4: Differentially expressed genes in adr-1(-);adr-2(-) animals grown on OP40 for 7 hours compared to wildtype animals grown on OP50 for 7 hours (Related to Figure 4). DESeq2 was used to identify differentially expressed genes in adr-1(-);adr-2(-) double mutant animals compared to wildtype animals. p-value<0.05 were taken as targets misregulated in absence of ad-1 and adr-2. Differentially expressed genes are listed as Wormbase annotations in Column A, Gene name in Column B. The log2fold change values, raw p values and the adjusted p values as obtained from the DESeq2 analysis are shown in columns C, D, and E, respectively. Read counts from the DESeq2 analysis for the three biological replicates of wildtype are listed in columns F, G and H. Read counts from the DESeq2 analysis for the three biological replicates of adr-1(-);adr-2(-) animals are listed in columns I, J and K . "	"Supplemental Table 4: Differentially expressed genes in adr-1(-);adr-2(-) animals grown on OP40 for 7 hours compared to wildtype animals grown on OP50 for 7 hours (Related to Figure 4). DESeq2 was used to identify differentially expressed genes in adr-1(-);adr-2(-) double mutant animals compared to wildtype animals. p-value<0.05 were taken as targets misregulated in absence of ad-1 and adr-2. Differentially expressed genes are listed as Wormbase annotations in Column A, Gene name in Column B. The log2fold change values, raw p values and the adjusted p values as obtained from the DESeq2 analysis are shown in columns C, D, and E, respectively. Read counts from the DESeq2 analysis for the three biological replicates of wildtype are listed in columns F, G and H. Read counts from the DESeq2 analysis for the three biological replicates of adr-1(-);adr-2(-) animals are listed in columns I, J and K . "	"Supplemental Table 4: Differentially expressed genes in adr-1(-);adr-2(-) animals grown on OP40 for 7 hours compared to wildtype animals grown on OP50 for 7 hours (Related to Figure 4). DESeq2 was used to identify differentially expressed genes in adr-1(-);adr-2(-) double mutant animals compared to wildtype animals. p-value<0.05 were taken as targets misregulated in absence of ad-1 and adr-2. Differentially expressed genes are listed as Wormbase annotations in Column A, Gene name in Column B. The log2fold change values, raw p values and the adjusted p values as obtained from the DESeq2 analysis are shown in columns C, D, and E, respectively. Read counts from the DESeq2 analysis for the three biological replicates of wildtype are listed in columns F, G and H. Read counts from the DESeq2 analysis for the three biological replicates of adr-1(-);adr-2(-) animals are listed in columns I, J and K . "	"Supplemental Table 4: Differentially expressed genes in adr-1(-);adr-2(-) animals grown on OP40 for 7 hours compared to wildtype animals grown on OP50 for 7 hours (Related to Figure 4). DESeq2 was used to identify differentially expressed genes in adr-1(-);adr-2(-) double mutant animals compared to wildtype animals. p-value<0.05 were taken as targets misregulated in absence of ad-1 and adr-2. Differentially expressed genes are listed as Wormbase annotations in Column A, Gene name in Column B. The log2fold change values, raw p values and the adjusted p values as obtained from the DESeq2 analysis are shown in columns C, D, and E, respectively. Read counts from the DESeq2 analysis for the three biological replicates of wildtype are listed in columns F, G and H. Read counts from the DESeq2 analysis for the three biological replicates of adr-1(-);adr-2(-) animals are listed in columns I, J and K . "	"Supplemental Table 4: Differentially expressed genes in adr-1(-);adr-2(-) animals grown on OP40 for 7 hours compared to wildtype animals grown on OP50 for 7 hours (Related to Figure 4). DESeq2 was used to identify differentially expressed genes in adr-1(-);adr-2(-) double mutant animals compared to wildtype animals. p-value<0.05 were taken as targets misregulated in absence of ad-1 and adr-2. Differentially expressed genes are listed as Wormbase annotations in Column A, Gene name in Column B. The log2fold change values, raw p values and the adjusted p values as obtained from the DESeq2 analysis are shown in columns C, D, and E, respectively. Read counts from the DESeq2 analysis for the three biological replicates of wildtype are listed in columns F, G and H. Read counts from the DESeq2 analysis for the three biological replicates of adr-1(-);adr-2(-) animals are listed in columns I, J and K . "	"Supplemental Table 4: Differentially expressed genes in adr-1(-);adr-2(-) animals grown on OP40 for 7 hours compared to wildtype animals grown on OP50 for 7 hours (Related to Figure 4). DESeq2 was used to identify differentially expressed genes in adr-1(-);adr-2(-) double mutant animals compared to wildtype animals. p-value<0.05 were taken as targets misregulated in absence of ad-1 and adr-2. Differentially expressed genes are listed as Wormbase annotations in Column A, Gene name in Column B. The log2fold change values, raw p values and the adjusted p values as obtained from the DESeq2 analysis are shown in columns C, D, and E, respectively. Read counts from the DESeq2 analysis for the three biological replicates of wildtype are listed in columns F, G and H. Read counts from the DESeq2 analysis for the three biological replicates of adr-1(-);adr-2(-) animals are listed in columns I, J and K . "	"Supplemental Table 4: Differentially expressed genes in adr-1(-);adr-2(-) animals grown on OP40 for 7 hours compared to wildtype animals grown on OP50 for 7 hours (Related to Figure 4). DESeq2 was used to identify differentially expressed genes in adr-1(-);adr-2(-) double mutant animals compared to wildtype animals. p-value<0.05 were taken as targets misregulated in absence of ad-1 and adr-2. Differentially expressed genes are listed as Wormbase annotations in Column A, Gene name in Column B. The log2fold change values, raw p values and the adjusted p values as obtained from the DESeq2 analysis are shown in columns C, D, and E, respectively. Read counts from the DESeq2 analysis for the three biological replicates of wildtype are listed in columns F, G and H. Read counts from the DESeq2 analysis for the three biological replicates of adr-1(-);adr-2(-) animals are listed in columns I, J and K . "	"Supplemental Table 4: Differentially expressed genes in adr-1(-);adr-2(-) animals grown on OP40 for 7 hours compared to wildtype animals grown on OP50 for 7 hours (Related to Figure 4). DESeq2 was used to identify differentially expressed genes in adr-1(-);adr-2(-) double mutant animals compared to wildtype animals. p-value<0.05 were taken as targets misregulated in absence of ad-1 and adr-2. Differentially expressed genes are listed as Wormbase annotations in Column A, Gene name in Column B. The log2fold change values, raw p values and the adjusted p values as obtained from the DESeq2 analysis are shown in columns C, D, and E, respectively. Read counts from the DESeq2 analysis for the three biological replicates of wildtype are listed in columns F, G and H. Read counts from the DESeq2 analysis for the three biological replicates of adr-1(-);adr-2(-) animals are listed in columns I, J and K . "	"Supplemental Table 4: Differentially expressed genes in adr-1(-);adr-2(-) animals grown on OP40 for 7 hours compared to wildtype animals grown on OP50 for 7 hours (Related to Figure 4). DESeq2 was used to identify differentially expressed genes in adr-1(-);adr-2(-) double mutant animals compared to wildtype animals. p-value<0.05 were taken as targets misregulated in absence of ad-1 and adr-2. Differentially expressed genes are listed as Wormbase annotations in Column A, Gene name in Column B. The log2fold change values, raw p values and the adjusted p values as obtained from the DESeq2 analysis are shown in columns C, D, and E, respectively. Read counts from the DESeq2 analysis for the three biological replicates of wildtype are listed in columns F, G and H. Read counts from the DESeq2 analysis for the three biological replicates of adr-1(-);adr-2(-) animals are listed in columns I, J and K . "	"Supplemental Table 4: Differentially expressed genes in adr-1(-);adr-2(-) animals grown on OP40 for 7 hours compared to wildtype animals grown on OP50 for 7 hours (Related to Figure 4). DESeq2 was used to identify differentially expressed genes in adr-1(-);adr-2(-) double mutant animals compared to wildtype animals. p-value<0.05 were taken as targets misregulated in absence of ad-1 and adr-2. Differentially expressed genes are listed as Wormbase annotations in Column A, Gene name in Column B. The log2fold change values, raw p values and the adjusted p values as obtained from the DESeq2 analysis are shown in columns C, D, and E, respectively. Read counts from the DESeq2 analysis for the three biological replicates of wildtype are listed in columns F, G and H. Read counts from the DESeq2 analysis for the three biological replicates of adr-1(-);adr-2(-) animals are listed in columns I, J and K . "	"Supplemental Table 4: Differentially expressed genes in adr-1(-);adr-2(-) animals grown on OP40 for 7 hours compared to wildtype animals grown on OP50 for 7 hours (Related to Figure 4). DESeq2 was used to identify differentially expressed genes in adr-1(-);adr-2(-) double mutant animals compared to wildtype animals. p-value<0.05 were taken as targets misregulated in absence of ad-1 and adr-2. Differentially expressed genes are listed as Wormbase annotations in Column A, Gene name in Column B. The log2fold change values, raw p values and the adjusted p values as obtained from the DESeq2 analysis are shown in columns C, D, and E, respectively. Read counts from the DESeq2 analysis for the three biological replicates of wildtype are listed in columns F, G and H. Read counts from the DESeq2 analysis for the three biological replicates of adr-1(-);adr-2(-) animals are listed in columns I, J and K . "	"Supplemental Table 4: Differentially expressed genes in adr-1(-);adr-2(-) animals grown on OP40 for 7 hours compared to wildtype animals grown on OP50 for 7 hours (Related to Figure 4). DESeq2 was used to identify differentially expressed genes in adr-1(-);adr-2(-) double mutant animals compared to wildtype animals. p-value<0.05 were taken as targets misregulated in absence of ad-1 and adr-2. Differentially expressed genes are listed as Wormbase annotations in Column A, Gene name in Column B. The log2fold change values, raw p values and the adjusted p values as obtained from the DESeq2 analysis are shown in columns C, D, and E, respectively. Read counts from the DESeq2 analysis for the three biological replicates of wildtype are listed in columns F, G and H. Read counts from the DESeq2 analysis for the three biological replicates of adr-1(-);adr-2(-) animals are listed in columns I, J and K . "	"Supplemental Table 4: Differentially expressed genes in adr-1(-);adr-2(-) animals grown on OP40 for 7 hours compared to wildtype animals grown on OP50 for 7 hours (Related to Figure 4). DESeq2 was used to identify differentially expressed genes in adr-1(-);adr-2(-) double mutant animals compared to wildtype animals. p-value<0.05 were taken as targets misregulated in absence of ad-1 and adr-2. Differentially expressed genes are listed as Wormbase annotations in Column A, Gene name in Column B. The log2fold change values, raw p values and the adjusted p values as obtained from the DESeq2 analysis are shown in columns C, D, and E, respectively. Read counts from the DESeq2 analysis for the three biological replicates of wildtype are listed in columns F, G and H. Read counts from the DESeq2 analysis for the three biological replicates of adr-1(-);adr-2(-) animals are listed in columns I, J and K . "	"Supplemental Table 4: Differentially expressed genes in adr-1(-);adr-2(-) animals grown on OP40 for 7 hours compared to wildtype animals grown on OP50 for 7 hours (Related to Figure 4). DESeq2 was used to identify differentially expressed genes in adr-1(-);adr-2(-) double mutant animals compared to wildtype animals. p-value<0.05 were taken as targets misregulated in absence of ad-1 and adr-2. Differentially expressed genes are listed as Wormbase annotations in Column A, Gene name in Column B. The log2fold change values, raw p values and the adjusted p values as obtained from the DESeq2 analysis are shown in columns C, D, and E, respectively. Read counts from the DESeq2 analysis for the three biological replicates of wildtype are listed in columns F, G and H. Read counts from the DESeq2 analysis for the three biological replicates of adr-1(-);adr-2(-) animals are listed in columns I, J and K . "	"Supplemental Table 4: Differentially expressed genes in adr-1(-);adr-2(-) animals grown on OP40 for 7 hours compared to wildtype animals grown on OP50 for 7 hours (Related to Figure 4). DESeq2 was used to identify differentially expressed genes in adr-1(-);adr-2(-) double mutant animals compared to wildtype animals. p-value<0.05 were taken as targets misregulated in absence of ad-1 and adr-2. Differentially expressed genes are listed as Wormbase annotations in Column A, Gene name in Column B. The log2fold change values, raw p values and the adjusted p values as obtained from the DESeq2 analysis are shown in columns C, D, and E, respectively. Read counts from the DESeq2 analysis for the three biological replicates of wildtype are listed in columns F, G and H. Read counts from the DESeq2 analysis for the three biological replicates of adr-1(-);adr-2(-) animals are listed in columns I, J and K . "	"Supplemental Table 4: Differentially expressed genes in adr-1(-);adr-2(-) animals grown on OP40 for 7 hours compared to wildtype animals grown on OP50 for 7 hours (Related to Figure 4). DESeq2 was used to identify differentially expressed genes in adr-1(-);adr-2(-) double mutant animals compared to wildtype animals. p-value<0.05 were taken as targets misregulated in absence of ad-1 and adr-2. Differentially expressed genes are listed as Wormbase annotations in Column A, Gene name in Column B. The log2fold change values, raw p values and the adjusted p values as obtained from the DESeq2 analysis are shown in columns C, D, and E, respectively. Read counts from the DESeq2 analysis for the three biological replicates of wildtype are listed in columns F, G and H. Read counts from the DESeq2 analysis for the three biological replicates of adr-1(-);adr-2(-) animals are listed in columns I, J and K . "	"Supplemental Table 4: Differentially expressed genes in adr-1(-);adr-2(-) animals grown on OP40 for 7 hours compared to wildtype animals grown on OP50 for 7 hours (Related to Figure 4). DESeq2 was used to identify differentially expressed genes in adr-1(-);adr-2(-) double mutant animals compared to wildtype animals. p-value<0.05 were taken as targets misregulated in absence of ad-1 and adr-2. Differentially expressed genes are listed as Wormbase annotations in Column A, Gene name in Column B. The log2fold change values, raw p values and the adjusted p values as obtained from the DESeq2 analysis are shown in columns C, D, and E, respectively. Read counts from the DESeq2 analysis for the three biological replicates of wildtype are listed in columns F, G and H. Read counts from the DESeq2 analysis for the three biological replicates of adr-1(-);adr-2(-) animals are listed in columns I, J and K . "	"Supplemental Table 4: Differentially expressed genes in adr-1(-);adr-2(-) animals grown on OP40 for 7 hours compared to wildtype animals grown on OP50 for 7 hours (Related to Figure 4). DESeq2 was used to identify differentially expressed genes in adr-1(-);adr-2(-) double mutant animals compared to wildtype animals. p-value<0.05 were taken as targets misregulated in absence of ad-1 and adr-2. Differentially expressed genes are listed as Wormbase annotations in Column A, Gene name in Column B. The log2fold change values, raw p values and the adjusted p values as obtained from the DESeq2 analysis are shown in columns C, D, and E, respectively. Read counts from the DESeq2 analysis for the three biological replicates of wildtype are listed in columns F, G and H. Read counts from the DESeq2 analysis for the three biological replicates of adr-1(-);adr-2(-) animals are listed in columns I, J and K . "	"Supplemental Table 4: Differentially expressed genes in adr-1(-);adr-2(-) animals grown on OP40 for 7 hours compared to wildtype animals grown on OP50 for 7 hours (Related to Figure 4). DESeq2 was used to identify differentially expressed genes in adr-1(-);adr-2(-) double mutant animals compared to wildtype animals. p-value<0.05 were taken as targets misregulated in absence of ad-1 and adr-2. Differentially expressed genes are listed as Wormbase annotations in Column A, Gene name in Column B. The log2fold change values, raw p values and the adjusted p values as obtained from the DESeq2 analysis are shown in columns C, D, and E, respectively. Read counts from the DESeq2 analysis for the three biological replicates of wildtype are listed in columns F, G and H. Read counts from the DESeq2 analysis for the three biological replicates of adr-1(-);adr-2(-) animals are listed in columns I, J and K . "	"Supplemental Table 4: Differentially expressed genes in adr-1(-);adr-2(-) animals grown on OP40 for 7 hours compared to wildtype animals grown on OP50 for 7 hours (Related to Figure 4). DESeq2 was used to identify differentially expressed genes in adr-1(-);adr-2(-) double mutant animals compared to wildtype animals. p-value<0.05 were taken as targets misregulated in absence of ad-1 and adr-2. Differentially expressed genes are listed as Wormbase annotations in Column A, Gene name in Column B. The log2fold change values, raw p values and the adjusted p values as obtained from the DESeq2 analysis are shown in columns C, D, and E, respectively. Read counts from the DESeq2 analysis for the three biological replicates of wildtype are listed in columns F, G and H. Read counts from the DESeq2 analysis for the three biological replicates of adr-1(-);adr-2(-) animals are listed in columns I, J and K . "	"Supplemental Table 4: Differentially expressed genes in adr-1(-);adr-2(-) animals grown on OP40 for 7 hours compared to wildtype animals grown on OP50 for 7 hours (Related to Figure 4). DESeq2 was used to identify differentially expressed genes in adr-1(-);adr-2(-) double mutant animals compared to wildtype animals. p-value<0.05 were taken as targets misregulated in absence of ad-1 and adr-2. Differentially expressed genes are listed as Wormbase annotations in Column A, Gene name in Column B. The log2fold change values, raw p values and the adjusted p values as obtained from the DESeq2 analysis are shown in columns C, D, and E, respectively. Read counts from the DESeq2 analysis for the three biological replicates of wildtype are listed in columns F, G and H. Read counts from the DESeq2 analysis for the three biological replicates of adr-1(-);adr-2(-) animals are listed in columns I, J and K . "	"Supplemental Table 4: Differentially expressed genes in adr-1(-);adr-2(-) animals grown on OP40 for 7 hours compared to wildtype animals grown on OP50 for 7 hours (Related to Figure 4). DESeq2 was used to identify differentially expressed genes in adr-1(-);adr-2(-) double mutant animals compared to wildtype animals. p-value<0.05 were taken as targets misregulated in absence of ad-1 and adr-2. Differentially expressed genes are listed as Wormbase annotations in Column A, Gene name in Column B. The log2fold change values, raw p values and the adjusted p values as obtained from the DESeq2 analysis are shown in columns C, D, and E, respectively. Read counts from the DESeq2 analysis for the three biological replicates of wildtype are listed in columns F, G and H. Read counts from the DESeq2 analysis for the three biological replicates of adr-1(-);adr-2(-) animals are listed in columns I, J and K . "	"Supplemental Table 4: Differentially expressed genes in adr-1(-);adr-2(-) animals grown on OP40 for 7 hours compared to wildtype animals grown on OP50 for 7 hours (Related to Figure 4). DESeq2 was used to identify differentially expressed genes in adr-1(-);adr-2(-) double mutant animals compared to wildtype animals. p-value<0.05 were taken as targets misregulated in absence of ad-1 and adr-2. Differentially expressed genes are listed as Wormbase annotations in Column A, Gene name in Column B. The log2fold change values, raw p values and the adjusted p values as obtained from the DESeq2 analysis are shown in columns C, D, and E, respectively. Read counts from the DESeq2 analysis for the three biological replicates of wildtype are listed in columns F, G and H. Read counts from the DESeq2 analysis for the three biological replicates of adr-1(-);adr-2(-) animals are listed in columns I, J and K . "	"Supplemental Table 4: Differentially expressed genes in adr-1(-);adr-2(-) animals grown on OP40 for 7 hours compared to wildtype animals grown on OP50 for 7 hours (Related to Figure 4). DESeq2 was used to identify differentially expressed genes in adr-1(-);adr-2(-) double mutant animals compared to wildtype animals. p-value<0.05 were taken as targets misregulated in absence of ad-1 and adr-2. Differentially expressed genes are listed as Wormbase annotations in Column A, Gene name in Column B. The log2fold change values, raw p values and the adjusted p values as obtained from the DESeq2 analysis are shown in columns C, D, and E, respectively. Read counts from the DESeq2 analysis for the three biological replicates of wildtype are listed in columns F, G and H. Read counts from the DESeq2 analysis for the three biological replicates of adr-1(-);adr-2(-) animals are listed in columns I, J and K . "	"Supplemental Table 4: Differentially expressed genes in adr-1(-);adr-2(-) animals grown on OP40 for 7 hours compared to wildtype animals grown on OP50 for 7 hours (Related to Figure 4). DESeq2 was used to identify differentially expressed genes in adr-1(-);adr-2(-) double mutant animals compared to wildtype animals. p-value<0.05 were taken as targets misregulated in absence of ad-1 and adr-2. Differentially expressed genes are listed as Wormbase annotations in Column A, Gene name in Column B. The log2fold change values, raw p values and the adjusted p values as obtained from the DESeq2 analysis are shown in columns C, D, and E, respectively. Read counts from the DESeq2 analysis for the three biological replicates of wildtype are listed in columns F, G and H. Read counts from the DESeq2 analysis for the three biological replicates of adr-1(-);adr-2(-) animals are listed in columns I, J and K . "	"Supplemental Table 4: Differentially expressed genes in adr-1(-);adr-2(-) animals grown on OP40 for 7 hours compared to wildtype animals grown on OP50 for 7 hours (Related to Figure 4). DESeq2 was used to identify differentially expressed genes in adr-1(-);adr-2(-) double mutant animals compared to wildtype animals. p-value<0.05 were taken as targets misregulated in absence of ad-1 and adr-2. Differentially expressed genes are listed as Wormbase annotations in Column A, Gene name in Column B. The log2fold change values, raw p values and the adjusted p values as obtained from the DESeq2 analysis are shown in columns C, D, and E, respectively. Read counts from the DESeq2 analysis for the three biological replicates of wildtype are listed in columns F, G and H. Read counts from the DESeq2 analysis for the three biological replicates of adr-1(-);adr-2(-) animals are listed in columns I, J and K . "	"Supplemental Table 4: Differentially expressed genes in adr-1(-);adr-2(-) animals grown on OP40 for 7 hours compared to wildtype animals grown on OP50 for 7 hours (Related to Figure 4). DESeq2 was used to identify differentially expressed genes in adr-1(-);adr-2(-) double mutant animals compared to wildtype animals. p-value<0.05 were taken as targets misregulated in absence of ad-1 and adr-2. Differentially expressed genes are listed as Wormbase annotations in Column A, Gene name in Column B. The log2fold change values, raw p values and the adjusted p values as obtained from the DESeq2 analysis are shown in columns C, D, and E, respectively. Read counts from the DESeq2 analysis for the three biological replicates of wildtype are listed in columns F, G and H. Read counts from the DESeq2 analysis for the three biological replicates of adr-1(-);adr-2(-) animals are listed in columns I, J and K . "	"Supplemental Table 4: Differentially expressed genes in adr-1(-);adr-2(-) animals grown on OP40 for 7 hours compared to wildtype animals grown on OP50 for 7 hours (Related to Figure 4). DESeq2 was used to identify differentially expressed genes in adr-1(-);adr-2(-) double mutant animals compared to wildtype animals. p-value<0.05 were taken as targets misregulated in absence of ad-1 and adr-2. Differentially expressed genes are listed as Wormbase annotations in Column A, Gene name in Column B. The log2fold change values, raw p values and the adjusted p values as obtained from the DESeq2 analysis are shown in columns C, D, and E, respectively. Read counts from the DESeq2 analysis for the three biological replicates of wildtype are listed in columns F, G and H. Read counts from the DESeq2 analysis for the three biological replicates of adr-1(-);adr-2(-) animals are listed in columns I, J and K . "	"Supplemental Table 4: Differentially expressed genes in adr-1(-);adr-2(-) animals grown on OP40 for 7 hours compared to wildtype animals grown on OP50 for 7 hours (Related to Figure 4). DESeq2 was used to identify differentially expressed genes in adr-1(-);adr-2(-) double mutant animals compared to wildtype animals. p-value<0.05 were taken as targets misregulated in absence of ad-1 and adr-2. Differentially expressed genes are listed as Wormbase annotations in Column A, Gene name in Column B. The log2fold change values, raw p values and the adjusted p values as obtained from the DESeq2 analysis are shown in columns C, D, and E, respectively. Read counts from the DESeq2 analysis for the three biological replicates of wildtype are listed in columns F, G and H. Read counts from the DESeq2 analysis for the three biological replicates of adr-1(-);adr-2(-) animals are listed in columns I, J and K . "	"Supplemental Table 4: Differentially expressed genes in adr-1(-);adr-2(-) animals grown on OP40 for 7 hours compared to wildtype animals grown on OP50 for 7 hours (Related to Figure 4). DESeq2 was used to identify differentially expressed genes in adr-1(-);adr-2(-) double mutant animals compared to wildtype animals. p-value<0.05 were taken as targets misregulated in absence of ad-1 and adr-2. Differentially expressed genes are listed as Wormbase annotations in Column A, Gene name in Column B. The log2fold change values, raw p values and the adjusted p values as obtained from the DESeq2 analysis are shown in columns C, D, and E, respectively. Read counts from the DESeq2 analysis for the three biological replicates of wildtype are listed in columns F, G and H. Read counts from the DESeq2 analysis for the three biological replicates of adr-1(-);adr-2(-) animals are listed in columns I, J and K . "	"Supplemental Table 4: Differentially expressed genes in adr-1(-);adr-2(-) animals grown on OP40 for 7 hours compared to wildtype animals grown on OP50 for 7 hours (Related to Figure 4). DESeq2 was used to identify differentially expressed genes in adr-1(-);adr-2(-) double mutant animals compared to wildtype animals. p-value<0.05 were taken as targets misregulated in absence of ad-1 and adr-2. Differentially expressed genes are listed as Wormbase annotations in Column A, Gene name in Column B. The log2fold change values, raw p values and the adjusted p values as obtained from the DESeq2 analysis are shown in columns C, D, and E, respectively. Read counts from the DESeq2 analysis for the three biological replicates of wildtype are listed in columns F, G and H. Read counts from the DESeq2 analysis for the three biological replicates of adr-1(-);adr-2(-) animals are listed in columns I, J and K . "	"Supplemental Table 4: Differentially expressed genes in adr-1(-);adr-2(-) animals grown on OP40 for 7 hours compared to wildtype animals grown on OP50 for 7 hours (Related to Figure 4). DESeq2 was used to identify differentially expressed genes in adr-1(-);adr-2(-) double mutant animals compared to wildtype animals. p-value<0.05 were taken as targets misregulated in absence of ad-1 and adr-2. Differentially expressed genes are listed as Wormbase annotations in Column A, Gene name in Column B. The log2fold change values, raw p values and the adjusted p values as obtained from the DESeq2 analysis are shown in columns C, D, and E, respectively. Read counts from the DESeq2 analysis for the three biological replicates of wildtype are listed in columns F, G and H. Read counts from the DESeq2 analysis for the three biological replicates of adr-1(-);adr-2(-) animals are listed in columns I, J and K . "	"Supplemental Table 4: Differentially expressed genes in adr-1(-);adr-2(-) animals grown on OP40 for 7 hours compared to wildtype animals grown on OP50 for 7 hours (Related to Figure 4). DESeq2 was used to identify differentially expressed genes in adr-1(-);adr-2(-) double mutant animals compared to wildtype animals. p-value<0.05 were taken as targets misregulated in absence of ad-1 and adr-2. Differentially expressed genes are listed as Wormbase annotations in Column A, Gene name in Column B. The log2fold change values, raw p values and the adjusted p values as obtained from the DESeq2 analysis are shown in columns C, D, and E, respectively. Read counts from the DESeq2 analysis for the three biological replicates of wildtype are listed in columns F, G and H. Read counts from the DESeq2 analysis for the three biological replicates of adr-1(-);adr-2(-) animals are listed in columns I, J and K . "	"Supplemental Table 4: Differentially expressed genes in adr-1(-);adr-2(-) animals grown on OP40 for 7 hours compared to wildtype animals grown on OP50 for 7 hours (Related to Figure 4). DESeq2 was used to identify differentially expressed genes in adr-1(-);adr-2(-) double mutant animals compared to wildtype animals. p-value<0.05 were taken as targets misregulated in absence of ad-1 and adr-2. Differentially expressed genes are listed as Wormbase annotations in Column A, Gene name in Column B. The log2fold change values, raw p values and the adjusted p values as obtained from the DESeq2 analysis are shown in columns C, D, and E, respectively. Read counts from the DESeq2 analysis for the three biological replicates of wildtype are listed in columns F, G and H. Read counts from the DESeq2 analysis for the three biological replicates of adr-1(-);adr-2(-) animals are listed in columns I, J and K . "	"Supplemental Table 4: Differentially expressed genes in adr-1(-);adr-2(-) animals grown on OP40 for 7 hours compared to wildtype animals grown on OP50 for 7 hours (Related to Figure 4). DESeq2 was used to identify differentially expressed genes in adr-1(-);adr-2(-) double mutant animals compared to wildtype animals. p-value<0.05 were taken as targets misregulated in absence of ad-1 and adr-2. Differentially expressed genes are listed as Wormbase annotations in Column A, Gene name in Column B. The log2fold change values, raw p values and the adjusted p values as obtained from the DESeq2 analysis are shown in columns C, D, and E, respectively. Read counts from the DESeq2 analysis for the three biological replicates of wildtype are listed in columns F, G and H. Read counts from the DESeq2 analysis for the three biological replicates of adr-1(-);adr-2(-) animals are listed in columns I, J and K . "	"Supplemental Table 4: Differentially expressed genes in adr-1(-);adr-2(-) animals grown on OP40 for 7 hours compared to wildtype animals grown on OP50 for 7 hours (Related to Figure 4). DESeq2 was used to identify differentially expressed genes in adr-1(-);adr-2(-) double mutant animals compared to wildtype animals. p-value<0.05 were taken as targets misregulated in absence of ad-1 and adr-2. Differentially expressed genes are listed as Wormbase annotations in Column A, Gene name in Column B. The log2fold change values, raw p values and the adjusted p values as obtained from the DESeq2 analysis are shown in columns C, D, and E, respectively. Read counts from the DESeq2 analysis for the three biological replicates of wildtype are listed in columns F, G and H. Read counts from the DESeq2 analysis for the three biological replicates of adr-1(-);adr-2(-) animals are listed in columns I, J and K . "	"Supplemental Table 4: Differentially expressed genes in adr-1(-);adr-2(-) animals grown on OP40 for 7 hours compared to wildtype animals grown on OP50 for 7 hours (Related to Figure 4). DESeq2 was used to identify differentially expressed genes in adr-1(-);adr-2(-) double mutant animals compared to wildtype animals. p-value<0.05 were taken as targets misregulated in absence of ad-1 and adr-2. Differentially expressed genes are listed as Wormbase annotations in Column A, Gene name in Column B. The log2fold change values, raw p values and the adjusted p values as obtained from the DESeq2 analysis are shown in columns C, D, and E, respectively. Read counts from the DESeq2 analysis for the three biological replicates of wildtype are listed in columns F, G and H. Read counts from the DESeq2 analysis for the three biological replicates of adr-1(-);adr-2(-) animals are listed in columns I, J and K . "	"Supplemental Table 4: Differentially expressed genes in adr-1(-);adr-2(-) animals grown on OP40 for 7 hours compared to wildtype animals grown on OP50 for 7 hours (Related to Figure 4). DESeq2 was used to identify differentially expressed genes in adr-1(-);adr-2(-) double mutant animals compared to wildtype animals. p-value<0.05 were taken as targets misregulated in absence of ad-1 and adr-2. Differentially expressed genes are listed as Wormbase annotations in Column A, Gene name in Column B. The log2fold change values, raw p values and the adjusted p values as obtained from the DESeq2 analysis are shown in columns C, D, and E, respectively. Read counts from the DESeq2 analysis for the three biological replicates of wildtype are listed in columns F, G and H. Read counts from the DESeq2 analysis for the three biological replicates of adr-1(-);adr-2(-) animals are listed in columns I, J and K . "	"Supplemental Table 4: Differentially expressed genes in adr-1(-);adr-2(-) animals grown on OP40 for 7 hours compared to wildtype animals grown on OP50 for 7 hours (Related to Figure 4). DESeq2 was used to identify differentially expressed genes in adr-1(-);adr-2(-) double mutant animals compared to wildtype animals. p-value<0.05 were taken as targets misregulated in absence of ad-1 and adr-2. Differentially expressed genes are listed as Wormbase annotations in Column A, Gene name in Column B. The log2fold change values, raw p values and the adjusted p values as obtained from the DESeq2 analysis are shown in columns C, D, and E, respectively. Read counts from the DESeq2 analysis for the three biological replicates of wildtype are listed in columns F, G and H. Read counts from the DESeq2 analysis for the three biological replicates of adr-1(-);adr-2(-) animals are listed in columns I, J and K . "	"Supplemental Table 4: Differentially expressed genes in adr-1(-);adr-2(-) animals grown on OP40 for 7 hours compared to wildtype animals grown on OP50 for 7 hours (Related to Figure 4). DESeq2 was used to identify differentially expressed genes in adr-1(-);adr-2(-) double mutant animals compared to wildtype animals. p-value<0.05 were taken as targets misregulated in absence of ad-1 and adr-2. Differentially expressed genes are listed as Wormbase annotations in Column A, Gene name in Column B. The log2fold change values, raw p values and the adjusted p values as obtained from the DESeq2 analysis are shown in columns C, D, and E, respectively. Read counts from the DESeq2 analysis for the three biological replicates of wildtype are listed in columns F, G and H. Read counts from the DESeq2 analysis for the three biological replicates of adr-1(-);adr-2(-) animals are listed in columns I, J and K . "	"Supplemental Table 4: Differentially expressed genes in adr-1(-);adr-2(-) animals grown on OP40 for 7 hours compared to wildtype animals grown on OP50 for 7 hours (Related to Figure 4). DESeq2 was used to identify differentially expressed genes in adr-1(-);adr-2(-) double mutant animals compared to wildtype animals. p-value<0.05 were taken as targets misregulated in absence of ad-1 and adr-2. Differentially expressed genes are listed as Wormbase annotations in Column A, Gene name in Column B. The log2fold change values, raw p values and the adjusted p values as obtained from the DESeq2 analysis are shown in columns C, D, and E, respectively. Read counts from the DESeq2 analysis for the three biological replicates of wildtype are listed in columns F, G and H. Read counts from the DESeq2 analysis for the three biological replicates of adr-1(-);adr-2(-) animals are listed in columns I, J and K . "	"Supplemental Table 4: Differentially expressed genes in adr-1(-);adr-2(-) animals grown on OP40 for 7 hours compared to wildtype animals grown on OP50 for 7 hours (Related to Figure 4). DESeq2 was used to identify differentially expressed genes in adr-1(-);adr-2(-) double mutant animals compared to wildtype animals. p-value<0.05 were taken as targets misregulated in absence of ad-1 and adr-2. Differentially expressed genes are listed as Wormbase annotations in Column A, Gene name in Column B. The log2fold change values, raw p values and the adjusted p values as obtained from the DESeq2 analysis are shown in columns C, D, and E, respectively. Read counts from the DESeq2 analysis for the three biological replicates of wildtype are listed in columns F, G and H. Read counts from the DESeq2 analysis for the three biological replicates of adr-1(-);adr-2(-) animals are listed in columns I, J and K . "	"Supplemental Table 4: Differentially expressed genes in adr-1(-);adr-2(-) animals grown on OP40 for 7 hours compared to wildtype animals grown on OP50 for 7 hours (Related to Figure 4). DESeq2 was used to identify differentially expressed genes in adr-1(-);adr-2(-) double mutant animals compared to wildtype animals. p-value<0.05 were taken as targets misregulated in absence of ad-1 and adr-2. Differentially expressed genes are listed as Wormbase annotations in Column A, Gene name in Column B. The log2fold change values, raw p values and the adjusted p values as obtained from the DESeq2 analysis are shown in columns C, D, and E, respectively. Read counts from the DESeq2 analysis for the three biological replicates of wildtype are listed in columns F, G and H. Read counts from the DESeq2 analysis for the three biological replicates of adr-1(-);adr-2(-) animals are listed in columns I, J and K . "	"Supplemental Table 4: Differentially expressed genes in adr-1(-);adr-2(-) animals grown on OP40 for 7 hours compared to wildtype animals grown on OP50 for 7 hours (Related to Figure 4). DESeq2 was used to identify differentially expressed genes in adr-1(-);adr-2(-) double mutant animals compared to wildtype animals. p-value<0.05 were taken as targets misregulated in absence of ad-1 and adr-2. Differentially expressed genes are listed as Wormbase annotations in Column A, Gene name in Column B. The log2fold change values, raw p values and the adjusted p values as obtained from the DESeq2 analysis are shown in columns C, D, and E, respectively. Read counts from the DESeq2 analysis for the three biological replicates of wildtype are listed in columns F, G and H. Read counts from the DESeq2 analysis for the three biological replicates of adr-1(-);adr-2(-) animals are listed in columns I, J and K . "	"Supplemental Table 4: Differentially expressed genes in adr-1(-);adr-2(-) animals grown on OP40 for 7 hours compared to wildtype animals grown on OP50 for 7 hours (Related to Figure 4). DESeq2 was used to identify differentially expressed genes in adr-1(-);adr-2(-) double mutant animals compared to wildtype animals. p-value<0.05 were taken as targets misregulated in absence of ad-1 and adr-2. Differentially expressed genes are listed as Wormbase annotations in Column A, Gene name in Column B. The log2fold change values, raw p values and the adjusted p values as obtained from the DESeq2 analysis are shown in columns C, D, and E, respectively. Read counts from the DESeq2 analysis for the three biological replicates of wildtype are listed in columns F, G and H. Read counts from the DESeq2 analysis for the three biological replicates of adr-1(-);adr-2(-) animals are listed in columns I, J and K . "	"Supplemental Table 4: Differentially expressed genes in adr-1(-);adr-2(-) animals grown on OP40 for 7 hours compared to wildtype animals grown on OP50 for 7 hours (Related to Figure 4). DESeq2 was used to identify differentially expressed genes in adr-1(-);adr-2(-) double mutant animals compared to wildtype animals. p-value<0.05 were taken as targets misregulated in absence of ad-1 and adr-2. Differentially expressed genes are listed as Wormbase annotations in Column A, Gene name in Column B. The log2fold change values, raw p values and the adjusted p values as obtained from the DESeq2 analysis are shown in columns C, D, and E, respectively. Read counts from the DESeq2 analysis for the three biological replicates of wildtype are listed in columns F, G and H. Read counts from the DESeq2 analysis for the three biological replicates of adr-1(-);adr-2(-) animals are listed in columns I, J and K . "	"Supplemental Table 4: Differentially expressed genes in adr-1(-);adr-2(-) animals grown on OP40 for 7 hours compared to wildtype animals grown on OP50 for 7 hours (Related to Figure 4). DESeq2 was used to identify differentially expressed genes in adr-1(-);adr-2(-) double mutant animals compared to wildtype animals. p-value<0.05 were taken as targets misregulated in absence of ad-1 and adr-2. Differentially expressed genes are listed as Wormbase annotations in Column A, Gene name in Column B. The log2fold change values, raw p values and the adjusted p values as obtained from the DESeq2 analysis are shown in columns C, D, and E, respectively. Read counts from the DESeq2 analysis for the three biological replicates of wildtype are listed in columns F, G and H. Read counts from the DESeq2 analysis for the three biological replicates of adr-1(-);adr-2(-) animals are listed in columns I, J and K . "	"Supplemental Table 4: Differentially expressed genes in adr-1(-);adr-2(-) animals grown on OP40 for 7 hours compared to wildtype animals grown on OP50 for 7 hours (Related to Figure 4). DESeq2 was used to identify differentially expressed genes in adr-1(-);adr-2(-) double mutant animals compared to wildtype animals. p-value<0.05 were taken as targets misregulated in absence of ad-1 and adr-2. Differentially expressed genes are listed as Wormbase annotations in Column A, Gene name in Column B. The log2fold change values, raw p values and the adjusted p values as obtained from the DESeq2 analysis are shown in columns C, D, and E, respectively. Read counts from the DESeq2 analysis for the three biological replicates of wildtype are listed in columns F, G and H. Read counts from the DESeq2 analysis for the three biological replicates of adr-1(-);adr-2(-) animals are listed in columns I, J and K . "	"Supplemental Table 4: Differentially expressed genes in adr-1(-);adr-2(-) animals grown on OP40 for 7 hours compared to wildtype animals grown on OP50 for 7 hours (Related to Figure 4). DESeq2 was used to identify differentially expressed genes in adr-1(-);adr-2(-) double mutant animals compared to wildtype animals. p-value<0.05 were taken as targets misregulated in absence of ad-1 and adr-2. Differentially expressed genes are listed as Wormbase annotations in Column A, Gene name in Column B. The log2fold change values, raw p values and the adjusted p values as obtained from the DESeq2 analysis are shown in columns C, D, and E, respectively. Read counts from the DESeq2 analysis for the three biological replicates of wildtype are listed in columns F, G and H. Read counts from the DESeq2 analysis for the three biological replicates of adr-1(-);adr-2(-) animals are listed in columns I, J and K . "	"Supplemental Table 4: Differentially expressed genes in adr-1(-);adr-2(-) animals grown on OP40 for 7 hours compared to wildtype animals grown on OP50 for 7 hours (Related to Figure 4). DESeq2 was used to identify differentially expressed genes in adr-1(-);adr-2(-) double mutant animals compared to wildtype animals. p-value<0.05 were taken as targets misregulated in absence of ad-1 and adr-2. Differentially expressed genes are listed as Wormbase annotations in Column A, Gene name in Column B. The log2fold change values, raw p values and the adjusted p values as obtained from the DESeq2 analysis are shown in columns C, D, and E, respectively. Read counts from the DESeq2 analysis for the three biological replicates of wildtype are listed in columns F, G and H. Read counts from the DESeq2 analysis for the three biological replicates of adr-1(-);adr-2(-) animals are listed in columns I, J and K . "	"Supplemental Table 4: Differentially expressed genes in adr-1(-);adr-2(-) animals grown on OP40 for 7 hours compared to wildtype animals grown on OP50 for 7 hours (Related to Figure 4). DESeq2 was used to identify differentially expressed genes in adr-1(-);adr-2(-) double mutant animals compared to wildtype animals. p-value<0.05 were taken as targets misregulated in absence of ad-1 and adr-2. Differentially expressed genes are listed as Wormbase annotations in Column A, Gene name in Column B. The log2fold change values, raw p values and the adjusted p values as obtained from the DESeq2 analysis are shown in columns C, D, and E, respectively. Read counts from the DESeq2 analysis for the three biological replicates of wildtype are listed in columns F, G and H. Read counts from the DESeq2 analysis for the three biological replicates of adr-1(-);adr-2(-) animals are listed in columns I, J and K . "	"Supplemental Table 4: Differentially expressed genes in adr-1(-);adr-2(-) animals grown on OP40 for 7 hours compared to wildtype animals grown on OP50 for 7 hours (Related to Figure 4). DESeq2 was used to identify differentially expressed genes in adr-1(-);adr-2(-) double mutant animals compared to wildtype animals. p-value<0.05 were taken as targets misregulated in absence of ad-1 and adr-2. Differentially expressed genes are listed as Wormbase annotations in Column A, Gene name in Column B. The log2fold change values, raw p values and the adjusted p values as obtained from the DESeq2 analysis are shown in columns C, D, and E, respectively. Read counts from the DESeq2 analysis for the three biological replicates of wildtype are listed in columns F, G and H. Read counts from the DESeq2 analysis for the three biological replicates of adr-1(-);adr-2(-) animals are listed in columns I, J and K . "	"Supplemental Table 4: Differentially expressed genes in adr-1(-);adr-2(-) animals grown on OP40 for 7 hours compared to wildtype animals grown on OP50 for 7 hours (Related to Figure 4). DESeq2 was used to identify differentially expressed genes in adr-1(-);adr-2(-) double mutant animals compared to wildtype animals. p-value<0.05 were taken as targets misregulated in absence of ad-1 and adr-2. Differentially expressed genes are listed as Wormbase annotations in Column A, Gene name in Column B. The log2fold change values, raw p values and the adjusted p values as obtained from the DESeq2 analysis are shown in columns C, D, and E, respectively. Read counts from the DESeq2 analysis for the three biological replicates of wildtype are listed in columns F, G and H. Read counts from the DESeq2 analysis for the three biological replicates of adr-1(-);adr-2(-) animals are listed in columns I, J and K . "	"Supplemental Table 4: Differentially expressed genes in adr-1(-);adr-2(-) animals grown on OP40 for 7 hours compared to wildtype animals grown on OP50 for 7 hours (Related to Figure 4). DESeq2 was used to identify differentially expressed genes in adr-1(-);adr-2(-) double mutant animals compared to wildtype animals. p-value<0.05 were taken as targets misregulated in absence of ad-1 and adr-2. Differentially expressed genes are listed as Wormbase annotations in Column A, Gene name in Column B. The log2fold change values, raw p values and the adjusted p values as obtained from the DESeq2 analysis are shown in columns C, D, and E, respectively. Read counts from the DESeq2 analysis for the three biological replicates of wildtype are listed in columns F, G and H. Read counts from the DESeq2 analysis for the three biological replicates of adr-1(-);adr-2(-) animals are listed in columns I, J and K . "	"Supplemental Table 4: Differentially expressed genes in adr-1(-);adr-2(-) animals grown on OP40 for 7 hours compared to wildtype animals grown on OP50 for 7 hours (Related to Figure 4). DESeq2 was used to identify differentially expressed genes in adr-1(-);adr-2(-) double mutant animals compared to wildtype animals. p-value<0.05 were taken as targets misregulated in absence of ad-1 and adr-2. Differentially expressed genes are listed as Wormbase annotations in Column A, Gene name in Column B. The log2fold change values, raw p values and the adjusted p values as obtained from the DESeq2 analysis are shown in columns C, D, and E, respectively. Read counts from the DESeq2 analysis for the three biological replicates of wildtype are listed in columns F, G and H. Read counts from the DESeq2 analysis for the three biological replicates of adr-1(-);adr-2(-) animals are listed in columns I, J and K . "	"Supplemental Table 4: Differentially expressed genes in adr-1(-);adr-2(-) animals grown on OP40 for 7 hours compared to wildtype animals grown on OP50 for 7 hours (Related to Figure 4). DESeq2 was used to identify differentially expressed genes in adr-1(-);adr-2(-) double mutant animals compared to wildtype animals. p-value<0.05 were taken as targets misregulated in absence of ad-1 and adr-2. Differentially expressed genes are listed as Wormbase annotations in Column A, Gene name in Column B. The log2fold change values, raw p values and the adjusted p values as obtained from the DESeq2 analysis are shown in columns C, D, and E, respectively. Read counts from the DESeq2 analysis for the three biological replicates of wildtype are listed in columns F, G and H. Read counts from the DESeq2 analysis for the three biological replicates of adr-1(-);adr-2(-) animals are listed in columns I, J and K . "	"Supplemental Table 4: Differentially expressed genes in adr-1(-);adr-2(-) animals grown on OP40 for 7 hours compared to wildtype animals grown on OP50 for 7 hours (Related to Figure 4). DESeq2 was used to identify differentially expressed genes in adr-1(-);adr-2(-) double mutant animals compared to wildtype animals. p-value<0.05 were taken as targets misregulated in absence of ad-1 and adr-2. Differentially expressed genes are listed as Wormbase annotations in Column A, Gene name in Column B. The log2fold change values, raw p values and the adjusted p values as obtained from the DESeq2 analysis are shown in columns C, D, and E, respectively. Read counts from the DESeq2 analysis for the three biological replicates of wildtype are listed in columns F, G and H. Read counts from the DESeq2 analysis for the three biological replicates of adr-1(-);adr-2(-) animals are listed in columns I, J and K . "	"Supplemental Table 4: Differentially expressed genes in adr-1(-);adr-2(-) animals grown on OP40 for 7 hours compared to wildtype animals grown on OP50 for 7 hours (Related to Figure 4). DESeq2 was used to identify differentially expressed genes in adr-1(-);adr-2(-) double mutant animals compared to wildtype animals. p-value<0.05 were taken as targets misregulated in absence of ad-1 and adr-2. Differentially expressed genes are listed as Wormbase annotations in Column A, Gene name in Column B. The log2fold change values, raw p values and the adjusted p values as obtained from the DESeq2 analysis are shown in columns C, D, and E, respectively. Read counts from the DESeq2 analysis for the three biological replicates of wildtype are listed in columns F, G and H. Read counts from the DESeq2 analysis for the three biological replicates of adr-1(-);adr-2(-) animals are listed in columns I, J and K . "	"Supplemental Table 4: Differentially expressed genes in adr-1(-);adr-2(-) animals grown on OP40 for 7 hours compared to wildtype animals grown on OP50 for 7 hours (Related to Figure 4). DESeq2 was used to identify differentially expressed genes in adr-1(-);adr-2(-) double mutant animals compared to wildtype animals. p-value<0.05 were taken as targets misregulated in absence of ad-1 and adr-2. Differentially expressed genes are listed as Wormbase annotations in Column A, Gene name in Column B. The log2fold change values, raw p values and the adjusted p values as obtained from the DESeq2 analysis are shown in columns C, D, and E, respectively. Read counts from the DESeq2 analysis for the three biological replicates of wildtype are listed in columns F, G and H. Read counts from the DESeq2 analysis for the three biological replicates of adr-1(-);adr-2(-) animals are listed in columns I, J and K . "	"Supplemental Table 4: Differentially expressed genes in adr-1(-);adr-2(-) animals grown on OP40 for 7 hours compared to wildtype animals grown on OP50 for 7 hours (Related to Figure 4). DESeq2 was used to identify differentially expressed genes in adr-1(-);adr-2(-) double mutant animals compared to wildtype animals. p-value<0.05 were taken as targets misregulated in absence of ad-1 and adr-2. Differentially expressed genes are listed as Wormbase annotations in Column A, Gene name in Column B. The log2fold change values, raw p values and the adjusted p values as obtained from the DESeq2 analysis are shown in columns C, D, and E, respectively. Read counts from the DESeq2 analysis for the three biological replicates of wildtype are listed in columns F, G and H. Read counts from the DESeq2 analysis for the three biological replicates of adr-1(-);adr-2(-) animals are listed in columns I, J and K . "	"Supplemental Table 4: Differentially expressed genes in adr-1(-);adr-2(-) animals grown on OP40 for 7 hours compared to wildtype animals grown on OP50 for 7 hours (Related to Figure 4). DESeq2 was used to identify differentially expressed genes in adr-1(-);adr-2(-) double mutant animals compared to wildtype animals. p-value<0.05 were taken as targets misregulated in absence of ad-1 and adr-2. Differentially expressed genes are listed as Wormbase annotations in Column A, Gene name in Column B. The log2fold change values, raw p values and the adjusted p values as obtained from the DESeq2 analysis are shown in columns C, D, and E, respectively. Read counts from the DESeq2 analysis for the three biological replicates of wildtype are listed in columns F, G and H. Read counts from the DESeq2 analysis for the three biological replicates of adr-1(-);adr-2(-) animals are listed in columns I, J and K . "	"Supplemental Table 4: Differentially expressed genes in adr-1(-);adr-2(-) animals grown on OP40 for 7 hours compared to wildtype animals grown on OP50 for 7 hours (Related to Figure 4). DESeq2 was used to identify differentially expressed genes in adr-1(-);adr-2(-) double mutant animals compared to wildtype animals. p-value<0.05 were taken as targets misregulated in absence of ad-1 and adr-2. Differentially expressed genes are listed as Wormbase annotations in Column A, Gene name in Column B. The log2fold change values, raw p values and the adjusted p values as obtained from the DESeq2 analysis are shown in columns C, D, and E, respectively. Read counts from the DESeq2 analysis for the three biological replicates of wildtype are listed in columns F, G and H. Read counts from the DESeq2 analysis for the three biological replicates of adr-1(-);adr-2(-) animals are listed in columns I, J and K . "	"Supplemental Table 4: Differentially expressed genes in adr-1(-);adr-2(-) animals grown on OP40 for 7 hours compared to wildtype animals grown on OP50 for 7 hours (Related to Figure 4). DESeq2 was used to identify differentially expressed genes in adr-1(-);adr-2(-) double mutant animals compared to wildtype animals. p-value<0.05 were taken as targets misregulated in absence of ad-1 and adr-2. Differentially expressed genes are listed as Wormbase annotations in Column A, Gene name in Column B. The log2fold change values, raw p values and the adjusted p values as obtained from the DESeq2 analysis are shown in columns C, D, and E, respectively. Read counts from the DESeq2 analysis for the three biological replicates of wildtype are listed in columns F, G and H. Read counts from the DESeq2 analysis for the three biological replicates of adr-1(-);adr-2(-) animals are listed in columns I, J and K . "	"Supplemental Table 4: Differentially expressed genes in adr-1(-);adr-2(-) animals grown on OP40 for 7 hours compared to wildtype animals grown on OP50 for 7 hours (Related to Figure 4). DESeq2 was used to identify differentially expressed genes in adr-1(-);adr-2(-) double mutant animals compared to wildtype animals. p-value<0.05 were taken as targets misregulated in absence of ad-1 and adr-2. Differentially expressed genes are listed as Wormbase annotations in Column A, Gene name in Column B. The log2fold change values, raw p values and the adjusted p values as obtained from the DESeq2 analysis are shown in columns C, D, and E, respectively. Read counts from the DESeq2 analysis for the three biological replicates of wildtype are listed in columns F, G and H. Read counts from the DESeq2 analysis for the three biological replicates of adr-1(-);adr-2(-) animals are listed in columns I, J and K . "	"Supplemental Table 4: Differentially expressed genes in adr-1(-);adr-2(-) animals grown on OP40 for 7 hours compared to wildtype animals grown on OP50 for 7 hours (Related to Figure 4). DESeq2 was used to identify differentially expressed genes in adr-1(-);adr-2(-) double mutant animals compared to wildtype animals. p-value<0.05 were taken as targets misregulated in absence of ad-1 and adr-2. Differentially expressed genes are listed as Wormbase annotations in Column A, Gene name in Column B. The log2fold change values, raw p values and the adjusted p values as obtained from the DESeq2 analysis are shown in columns C, D, and E, respectively. Read counts from the DESeq2 analysis for the three biological replicates of wildtype are listed in columns F, G and H. Read counts from the DESeq2 analysis for the three biological replicates of adr-1(-);adr-2(-) animals are listed in columns I, J and K . "	"Supplemental Table 4: Differentially expressed genes in adr-1(-);adr-2(-) animals grown on OP40 for 7 hours compared to wildtype animals grown on OP50 for 7 hours (Related to Figure 4). DESeq2 was used to identify differentially expressed genes in adr-1(-);adr-2(-) double mutant animals compared to wildtype animals. p-value<0.05 were taken as targets misregulated in absence of ad-1 and adr-2. Differentially expressed genes are listed as Wormbase annotations in Column A, Gene name in Column B. The log2fold change values, raw p values and the adjusted p values as obtained from the DESeq2 analysis are shown in columns C, D, and E, respectively. Read counts from the DESeq2 analysis for the three biological replicates of wildtype are listed in columns F, G and H. Read counts from the DESeq2 analysis for the three biological replicates of adr-1(-);adr-2(-) animals are listed in columns I, J and K . "	"Supplemental Table 4: Differentially expressed genes in adr-1(-);adr-2(-) animals grown on OP40 for 7 hours compared to wildtype animals grown on OP50 for 7 hours (Related to Figure 4). DESeq2 was used to identify differentially expressed genes in adr-1(-);adr-2(-) double mutant animals compared to wildtype animals. p-value<0.05 were taken as targets misregulated in absence of ad-1 and adr-2. Differentially expressed genes are listed as Wormbase annotations in Column A, Gene name in Column B. The log2fold change values, raw p values and the adjusted p values as obtained from the DESeq2 analysis are shown in columns C, D, and E, respectively. Read counts from the DESeq2 analysis for the three biological replicates of wildtype are listed in columns F, G and H. Read counts from the DESeq2 analysis for the three biological replicates of adr-1(-);adr-2(-) animals are listed in columns I, J and K . "	"Supplemental Table 4: Differentially expressed genes in adr-1(-);adr-2(-) animals grown on OP40 for 7 hours compared to wildtype animals grown on OP50 for 7 hours (Related to Figure 4). DESeq2 was used to identify differentially expressed genes in adr-1(-);adr-2(-) double mutant animals compared to wildtype animals. p-value<0.05 were taken as targets misregulated in absence of ad-1 and adr-2. Differentially expressed genes are listed as Wormbase annotations in Column A, Gene name in Column B. The log2fold change values, raw p values and the adjusted p values as obtained from the DESeq2 analysis are shown in columns C, D, and E, respectively. Read counts from the DESeq2 analysis for the three biological replicates of wildtype are listed in columns F, G and H. Read counts from the DESeq2 analysis for the three biological replicates of adr-1(-);adr-2(-) animals are listed in columns I, J and K . "	"Supplemental Table 4: Differentially expressed genes in adr-1(-);adr-2(-) animals grown on OP40 for 7 hours compared to wildtype animals grown on OP50 for 7 hours (Related to Figure 4). DESeq2 was used to identify differentially expressed genes in adr-1(-);adr-2(-) double mutant animals compared to wildtype animals. p-value<0.05 were taken as targets misregulated in absence of ad-1 and adr-2. Differentially expressed genes are listed as Wormbase annotations in Column A, Gene name in Column B. The log2fold change values, raw p values and the adjusted p values as obtained from the DESeq2 analysis are shown in columns C, D, and E, respectively. Read counts from the DESeq2 analysis for the three biological replicates of wildtype are listed in columns F, G and H. Read counts from the DESeq2 analysis for the three biological replicates of adr-1(-);adr-2(-) animals are listed in columns I, J and K . "	"Supplemental Table 4: Differentially expressed genes in adr-1(-);adr-2(-) animals grown on OP40 for 7 hours compared to wildtype animals grown on OP50 for 7 hours (Related to Figure 4). DESeq2 was used to identify differentially expressed genes in adr-1(-);adr-2(-) double mutant animals compared to wildtype animals. p-value<0.05 were taken as targets misregulated in absence of ad-1 and adr-2. Differentially expressed genes are listed as Wormbase annotations in Column A, Gene name in Column B. The log2fold change values, raw p values and the adjusted p values as obtained from the DESeq2 analysis are shown in columns C, D, and E, respectively. Read counts from the DESeq2 analysis for the three biological replicates of wildtype are listed in columns F, G and H. Read counts from the DESeq2 analysis for the three biological replicates of adr-1(-);adr-2(-) animals are listed in columns I, J and K . "	"Supplemental Table 4: Differentially expressed genes in adr-1(-);adr-2(-) animals grown on OP40 for 7 hours compared to wildtype animals grown on OP50 for 7 hours (Related to Figure 4). DESeq2 was used to identify differentially expressed genes in adr-1(-);adr-2(-) double mutant animals compared to wildtype animals. p-value<0.05 were taken as targets misregulated in absence of ad-1 and adr-2. Differentially expressed genes are listed as Wormbase annotations in Column A, Gene name in Column B. The log2fold change values, raw p values and the adjusted p values as obtained from the DESeq2 analysis are shown in columns C, D, and E, respectively. Read counts from the DESeq2 analysis for the three biological replicates of wildtype are listed in columns F, G and H. Read counts from the DESeq2 analysis for the three biological replicates of adr-1(-);adr-2(-) animals are listed in columns I, J and K . "	"Supplemental Table 4: Differentially expressed genes in adr-1(-);adr-2(-) animals grown on OP40 for 7 hours compared to wildtype animals grown on OP50 for 7 hours (Related to Figure 4). DESeq2 was used to identify differentially expressed genes in adr-1(-);adr-2(-) double mutant animals compared to wildtype animals. p-value<0.05 were taken as targets misregulated in absence of ad-1 and adr-2. Differentially expressed genes are listed as Wormbase annotations in Column A, Gene name in Column B. The log2fold change values, raw p values and the adjusted p values as obtained from the DESeq2 analysis are shown in columns C, D, and E, respectively. Read counts from the DESeq2 analysis for the three biological replicates of wildtype are listed in columns F, G and H. Read counts from the DESeq2 analysis for the three biological replicates of adr-1(-);adr-2(-) animals are listed in columns I, J and K . "	"Supplemental Table 4: Differentially expressed genes in adr-1(-);adr-2(-) animals grown on OP40 for 7 hours compared to wildtype animals grown on OP50 for 7 hours (Related to Figure 4). DESeq2 was used to identify differentially expressed genes in adr-1(-);adr-2(-) double mutant animals compared to wildtype animals. p-value<0.05 were taken as targets misregulated in absence of ad-1 and adr-2. Differentially expressed genes are listed as Wormbase annotations in Column A, Gene name in Column B. The log2fold change values, raw p values and the adjusted p values as obtained from the DESeq2 analysis are shown in columns C, D, and E, respectively. Read counts from the DESeq2 analysis for the three biological replicates of wildtype are listed in columns F, G and H. Read counts from the DESeq2 analysis for the three biological replicates of adr-1(-);adr-2(-) animals are listed in columns I, J and K . "	"Supplemental Table 4: Differentially expressed genes in adr-1(-);adr-2(-) animals grown on OP40 for 7 hours compared to wildtype animals grown on OP50 for 7 hours (Related to Figure 4). DESeq2 was used to identify differentially expressed genes in adr-1(-);adr-2(-) double mutant animals compared to wildtype animals. p-value<0.05 were taken as targets misregulated in absence of ad-1 and adr-2. Differentially expressed genes are listed as Wormbase annotations in Column A, Gene name in Column B. The log2fold change values, raw p values and the adjusted p values as obtained from the DESeq2 analysis are shown in columns C, D, and E, respectively. Read counts from the DESeq2 analysis for the three biological replicates of wildtype are listed in columns F, G and H. Read counts from the DESeq2 analysis for the three biological replicates of adr-1(-);adr-2(-) animals are listed in columns I, J and K . "	"Supplemental Table 4: Differentially expressed genes in adr-1(-);adr-2(-) animals grown on OP40 for 7 hours compared to wildtype animals grown on OP50 for 7 hours (Related to Figure 4). DESeq2 was used to identify differentially expressed genes in adr-1(-);adr-2(-) double mutant animals compared to wildtype animals. p-value<0.05 were taken as targets misregulated in absence of ad-1 and adr-2. Differentially expressed genes are listed as Wormbase annotations in Column A, Gene name in Column B. The log2fold change values, raw p values and the adjusted p values as obtained from the DESeq2 analysis are shown in columns C, D, and E, respectively. Read counts from the DESeq2 analysis for the three biological replicates of wildtype are listed in columns F, G and H. Read counts from the DESeq2 analysis for the three biological replicates of adr-1(-);adr-2(-) animals are listed in columns I, J and K . "	"Supplemental Table 4: Differentially expressed genes in adr-1(-);adr-2(-) animals grown on OP40 for 7 hours compared to wildtype animals grown on OP50 for 7 hours (Related to Figure 4). DESeq2 was used to identify differentially expressed genes in adr-1(-);adr-2(-) double mutant animals compared to wildtype animals. p-value<0.05 were taken as targets misregulated in absence of ad-1 and adr-2. Differentially expressed genes are listed as Wormbase annotations in Column A, Gene name in Column B. The log2fold change values, raw p values and the adjusted p values as obtained from the DESeq2 analysis are shown in columns C, D, and E, respectively. Read counts from the DESeq2 analysis for the three biological replicates of wildtype are listed in columns F, G and H. Read counts from the DESeq2 analysis for the three biological replicates of adr-1(-);adr-2(-) animals are listed in columns I, J and K . "	"Supplemental Table 4: Differentially expressed genes in adr-1(-);adr-2(-) animals grown on OP40 for 7 hours compared to wildtype animals grown on OP50 for 7 hours (Related to Figure 4). DESeq2 was used to identify differentially expressed genes in adr-1(-);adr-2(-) double mutant animals compared to wildtype animals. p-value<0.05 were taken as targets misregulated in absence of ad-1 and adr-2. Differentially expressed genes are listed as Wormbase annotations in Column A, Gene name in Column B. The log2fold change values, raw p values and the adjusted p values as obtained from the DESeq2 analysis are shown in columns C, D, and E, respectively. Read counts from the DESeq2 analysis for the three biological replicates of wildtype are listed in columns F, G and H. Read counts from the DESeq2 analysis for the three biological replicates of adr-1(-);adr-2(-) animals are listed in columns I, J and K . "	"Supplemental Table 4: Differentially expressed genes in adr-1(-);adr-2(-) animals grown on OP40 for 7 hours compared to wildtype animals grown on OP50 for 7 hours (Related to Figure 4). DESeq2 was used to identify differentially expressed genes in adr-1(-);adr-2(-) double mutant animals compared to wildtype animals. p-value<0.05 were taken as targets misregulated in absence of ad-1 and adr-2. Differentially expressed genes are listed as Wormbase annotations in Column A, Gene name in Column B. The log2fold change values, raw p values and the adjusted p values as obtained from the DESeq2 analysis are shown in columns C, D, and E, respectively. Read counts from the DESeq2 analysis for the three biological replicates of wildtype are listed in columns F, G and H. Read counts from the DESeq2 analysis for the three biological replicates of adr-1(-);adr-2(-) animals are listed in columns I, J and K . "	"Supplemental Table 4: Differentially expressed genes in adr-1(-);adr-2(-) animals grown on OP40 for 7 hours compared to wildtype animals grown on OP50 for 7 hours (Related to Figure 4). DESeq2 was used to identify differentially expressed genes in adr-1(-);adr-2(-) double mutant animals compared to wildtype animals. p-value<0.05 were taken as targets misregulated in absence of ad-1 and adr-2. Differentially expressed genes are listed as Wormbase annotations in Column A, Gene name in Column B. The log2fold change values, raw p values and the adjusted p values as obtained from the DESeq2 analysis are shown in columns C, D, and E, respectively. Read counts from the DESeq2 analysis for the three biological replicates of wildtype are listed in columns F, G and H. Read counts from the DESeq2 analysis for the three biological replicates of adr-1(-);adr-2(-) animals are listed in columns I, J and K . "	"Supplemental Table 4: Differentially expressed genes in adr-1(-);adr-2(-) animals grown on OP40 for 7 hours compared to wildtype animals grown on OP50 for 7 hours (Related to Figure 4). DESeq2 was used to identify differentially expressed genes in adr-1(-);adr-2(-) double mutant animals compared to wildtype animals. p-value<0.05 were taken as targets misregulated in absence of ad-1 and adr-2. Differentially expressed genes are listed as Wormbase annotations in Column A, Gene name in Column B. The log2fold change values, raw p values and the adjusted p values as obtained from the DESeq2 analysis are shown in columns C, D, and E, respectively. Read counts from the DESeq2 analysis for the three biological replicates of wildtype are listed in columns F, G and H. Read counts from the DESeq2 analysis for the three biological replicates of adr-1(-);adr-2(-) animals are listed in columns I, J and K . "	"Supplemental Table 4: Differentially expressed genes in adr-1(-);adr-2(-) animals grown on OP40 for 7 hours compared to wildtype animals grown on OP50 for 7 hours (Related to Figure 4). DESeq2 was used to identify differentially expressed genes in adr-1(-);adr-2(-) double mutant animals compared to wildtype animals. p-value<0.05 were taken as targets misregulated in absence of ad-1 and adr-2. Differentially expressed genes are listed as Wormbase annotations in Column A, Gene name in Column B. The log2fold change values, raw p values and the adjusted p values as obtained from the DESeq2 analysis are shown in columns C, D, and E, respectively. Read counts from the DESeq2 analysis for the three biological replicates of wildtype are listed in columns F, G and H. Read counts from the DESeq2 analysis for the three biological replicates of adr-1(-);adr-2(-) animals are listed in columns I, J and K . "	"Supplemental Table 4: Differentially expressed genes in adr-1(-);adr-2(-) animals grown on OP40 for 7 hours compared to wildtype animals grown on OP50 for 7 hours (Related to Figure 4). DESeq2 was used to identify differentially expressed genes in adr-1(-);adr-2(-) double mutant animals compared to wildtype animals. p-value<0.05 were taken as targets misregulated in absence of ad-1 and adr-2. Differentially expressed genes are listed as Wormbase annotations in Column A, Gene name in Column B. The log2fold change values, raw p values and the adjusted p values as obtained from the DESeq2 analysis are shown in columns C, D, and E, respectively. Read counts from the DESeq2 analysis for the three biological replicates of wildtype are listed in columns F, G and H. Read counts from the DESeq2 analysis for the three biological replicates of adr-1(-);adr-2(-) animals are listed in columns I, J and K . "	"Supplemental Table 4: Differentially expressed genes in adr-1(-);adr-2(-) animals grown on OP40 for 7 hours compared to wildtype animals grown on OP50 for 7 hours (Related to Figure 4). DESeq2 was used to identify differentially expressed genes in adr-1(-);adr-2(-) double mutant animals compared to wildtype animals. p-value<0.05 were taken as targets misregulated in absence of ad-1 and adr-2. Differentially expressed genes are listed as Wormbase annotations in Column A, Gene name in Column B. The log2fold change values, raw p values and the adjusted p values as obtained from the DESeq2 analysis are shown in columns C, D, and E, respectively. Read counts from the DESeq2 analysis for the three biological replicates of wildtype are listed in columns F, G and H. Read counts from the DESeq2 analysis for the three biological replicates of adr-1(-);adr-2(-) animals are listed in columns I, J and K . "	"Supplemental Table 4: Differentially expressed genes in adr-1(-);adr-2(-) animals grown on OP40 for 7 hours compared to wildtype animals grown on OP50 for 7 hours (Related to Figure 4). DESeq2 was used to identify differentially expressed genes in adr-1(-);adr-2(-) double mutant animals compared to wildtype animals. p-value<0.05 were taken as targets misregulated in absence of ad-1 and adr-2. Differentially expressed genes are listed as Wormbase annotations in Column A, Gene name in Column B. The log2fold change values, raw p values and the adjusted p values as obtained from the DESeq2 analysis are shown in columns C, D, and E, respectively. Read counts from the DESeq2 analysis for the three biological replicates of wildtype are listed in columns F, G and H. Read counts from the DESeq2 analysis for the three biological replicates of adr-1(-);adr-2(-) animals are listed in columns I, J and K . "	"Supplemental Table 4: Differentially expressed genes in adr-1(-);adr-2(-) animals grown on OP40 for 7 hours compared to wildtype animals grown on OP50 for 7 hours (Related to Figure 4). DESeq2 was used to identify differentially expressed genes in adr-1(-);adr-2(-) double mutant animals compared to wildtype animals. p-value<0.05 were taken as targets misregulated in absence of ad-1 and adr-2. Differentially expressed genes are listed as Wormbase annotations in Column A, Gene name in Column B. The log2fold change values, raw p values and the adjusted p values as obtained from the DESeq2 analysis are shown in columns C, D, and E, respectively. Read counts from the DESeq2 analysis for the three biological replicates of wildtype are listed in columns F, G and H. Read counts from the DESeq2 analysis for the three biological replicates of adr-1(-);adr-2(-) animals are listed in columns I, J and K . "	"Supplemental Table 4: Differentially expressed genes in adr-1(-);adr-2(-) animals grown on OP40 for 7 hours compared to wildtype animals grown on OP50 for 7 hours (Related to Figure 4). DESeq2 was used to identify differentially expressed genes in adr-1(-);adr-2(-) double mutant animals compared to wildtype animals. p-value<0.05 were taken as targets misregulated in absence of ad-1 and adr-2. Differentially expressed genes are listed as Wormbase annotations in Column A, Gene name in Column B. The log2fold change values, raw p values and the adjusted p values as obtained from the DESeq2 analysis are shown in columns C, D, and E, respectively. Read counts from the DESeq2 analysis for the three biological replicates of wildtype are listed in columns F, G and H. Read counts from the DESeq2 analysis for the three biological replicates of adr-1(-);adr-2(-) animals are listed in columns I, J and K . "	"Supplemental Table 4: Differentially expressed genes in adr-1(-);adr-2(-) animals grown on OP40 for 7 hours compared to wildtype animals grown on OP50 for 7 hours (Related to Figure 4). DESeq2 was used to identify differentially expressed genes in adr-1(-);adr-2(-) double mutant animals compared to wildtype animals. p-value<0.05 were taken as targets misregulated in absence of ad-1 and adr-2. Differentially expressed genes are listed as Wormbase annotations in Column A, Gene name in Column B. The log2fold change values, raw p values and the adjusted p values as obtained from the DESeq2 analysis are shown in columns C, D, and E, respectively. Read counts from the DESeq2 analysis for the three biological replicates of wildtype are listed in columns F, G and H. Read counts from the DESeq2 analysis for the three biological replicates of adr-1(-);adr-2(-) animals are listed in columns I, J and K . "	"Supplemental Table 4: Differentially expressed genes in adr-1(-);adr-2(-) animals grown on OP40 for 7 hours compared to wildtype animals grown on OP50 for 7 hours (Related to Figure 4). DESeq2 was used to identify differentially expressed genes in adr-1(-);adr-2(-) double mutant animals compared to wildtype animals. p-value<0.05 were taken as targets misregulated in absence of ad-1 and adr-2. Differentially expressed genes are listed as Wormbase annotations in Column A, Gene name in Column B. The log2fold change values, raw p values and the adjusted p values as obtained from the DESeq2 analysis are shown in columns C, D, and E, respectively. Read counts from the DESeq2 analysis for the three biological replicates of wildtype are listed in columns F, G and H. Read counts from the DESeq2 analysis for the three biological replicates of adr-1(-);adr-2(-) animals are listed in columns I, J and K . "	"Supplemental Table 4: Differentially expressed genes in adr-1(-);adr-2(-) animals grown on OP40 for 7 hours compared to wildtype animals grown on OP50 for 7 hours (Related to Figure 4). DESeq2 was used to identify differentially expressed genes in adr-1(-);adr-2(-) double mutant animals compared to wildtype animals. p-value<0.05 were taken as targets misregulated in absence of ad-1 and adr-2. Differentially expressed genes are listed as Wormbase annotations in Column A, Gene name in Column B. The log2fold change values, raw p values and the adjusted p values as obtained from the DESeq2 analysis are shown in columns C, D, and E, respectively. Read counts from the DESeq2 analysis for the three biological replicates of wildtype are listed in columns F, G and H. Read counts from the DESeq2 analysis for the three biological replicates of adr-1(-);adr-2(-) animals are listed in columns I, J and K . "	"Supplemental Table 4: Differentially expressed genes in adr-1(-);adr-2(-) animals grown on OP40 for 7 hours compared to wildtype animals grown on OP50 for 7 hours (Related to Figure 4). DESeq2 was used to identify differentially expressed genes in adr-1(-);adr-2(-) double mutant animals compared to wildtype animals. p-value<0.05 were taken as targets misregulated in absence of ad-1 and adr-2. Differentially expressed genes are listed as Wormbase annotations in Column A, Gene name in Column B. The log2fold change values, raw p values and the adjusted p values as obtained from the DESeq2 analysis are shown in columns C, D, and E, respectively. Read counts from the DESeq2 analysis for the three biological replicates of wildtype are listed in columns F, G and H. Read counts from the DESeq2 analysis for the three biological replicates of adr-1(-);adr-2(-) animals are listed in columns I, J and K . "	"Supplemental Table 4: Differentially expressed genes in adr-1(-);adr-2(-) animals grown on OP40 for 7 hours compared to wildtype animals grown on OP50 for 7 hours (Related to Figure 4). DESeq2 was used to identify differentially expressed genes in adr-1(-);adr-2(-) double mutant animals compared to wildtype animals. p-value<0.05 were taken as targets misregulated in absence of ad-1 and adr-2. Differentially expressed genes are listed as Wormbase annotations in Column A, Gene name in Column B. The log2fold change values, raw p values and the adjusted p values as obtained from the DESeq2 analysis are shown in columns C, D, and E, respectively. Read counts from the DESeq2 analysis for the three biological replicates of wildtype are listed in columns F, G and H. Read counts from the DESeq2 analysis for the three biological replicates of adr-1(-);adr-2(-) animals are listed in columns I, J and K . "	"Supplemental Table 4: Differentially expressed genes in adr-1(-);adr-2(-) animals grown on OP40 for 7 hours compared to wildtype animals grown on OP50 for 7 hours (Related to Figure 4). DESeq2 was used to identify differentially expressed genes in adr-1(-);adr-2(-) double mutant animals compared to wildtype animals. p-value<0.05 were taken as targets misregulated in absence of ad-1 and adr-2. Differentially expressed genes are listed as Wormbase annotations in Column A, Gene name in Column B. The log2fold change values, raw p values and the adjusted p values as obtained from the DESeq2 analysis are shown in columns C, D, and E, respectively. Read counts from the DESeq2 analysis for the three biological replicates of wildtype are listed in columns F, G and H. Read counts from the DESeq2 analysis for the three biological replicates of adr-1(-);adr-2(-) animals are listed in columns I, J and K . "	"Supplemental Table 4: Differentially expressed genes in adr-1(-);adr-2(-) animals grown on OP40 for 7 hours compared to wildtype animals grown on OP50 for 7 hours (Related to Figure 4). DESeq2 was used to identify differentially expressed genes in adr-1(-);adr-2(-) double mutant animals compared to wildtype animals. p-value<0.05 were taken as targets misregulated in absence of ad-1 and adr-2. Differentially expressed genes are listed as Wormbase annotations in Column A, Gene name in Column B. The log2fold change values, raw p values and the adjusted p values as obtained from the DESeq2 analysis are shown in columns C, D, and E, respectively. Read counts from the DESeq2 analysis for the three biological replicates of wildtype are listed in columns F, G and H. Read counts from the DESeq2 analysis for the three biological replicates of adr-1(-);adr-2(-) animals are listed in columns I, J and K . "	"Supplemental Table 4: Differentially expressed genes in adr-1(-);adr-2(-) animals grown on OP40 for 7 hours compared to wildtype animals grown on OP50 for 7 hours (Related to Figure 4). DESeq2 was used to identify differentially expressed genes in adr-1(-);adr-2(-) double mutant animals compared to wildtype animals. p-value<0.05 were taken as targets misregulated in absence of ad-1 and adr-2. Differentially expressed genes are listed as Wormbase annotations in Column A, Gene name in Column B. The log2fold change values, raw p values and the adjusted p values as obtained from the DESeq2 analysis are shown in columns C, D, and E, respectively. Read counts from the DESeq2 analysis for the three biological replicates of wildtype are listed in columns F, G and H. Read counts from the DESeq2 analysis for the three biological replicates of adr-1(-);adr-2(-) animals are listed in columns I, J and K . "	"Supplemental Table 4: Differentially expressed genes in adr-1(-);adr-2(-) animals grown on OP40 for 7 hours compared to wildtype animals grown on OP50 for 7 hours (Related to Figure 4). DESeq2 was used to identify differentially expressed genes in adr-1(-);adr-2(-) double mutant animals compared to wildtype animals. p-value<0.05 were taken as targets misregulated in absence of ad-1 and adr-2. Differentially expressed genes are listed as Wormbase annotations in Column A, Gene name in Column B. The log2fold change values, raw p values and the adjusted p values as obtained from the DESeq2 analysis are shown in columns C, D, and E, respectively. Read counts from the DESeq2 analysis for the three biological replicates of wildtype are listed in columns F, G and H. Read counts from the DESeq2 analysis for the three biological replicates of adr-1(-);adr-2(-) animals are listed in columns I, J and K . "	"Supplemental Table 4: Differentially expressed genes in adr-1(-);adr-2(-) animals grown on OP40 for 7 hours compared to wildtype animals grown on OP50 for 7 hours (Related to Figure 4). DESeq2 was used to identify differentially expressed genes in adr-1(-);adr-2(-) double mutant animals compared to wildtype animals. p-value<0.05 were taken as targets misregulated in absence of ad-1 and adr-2. Differentially expressed genes are listed as Wormbase annotations in Column A, Gene name in Column B. The log2fold change values, raw p values and the adjusted p values as obtained from the DESeq2 analysis are shown in columns C, D, and E, respectively. Read counts from the DESeq2 analysis for the three biological replicates of wildtype are listed in columns F, G and H. Read counts from the DESeq2 analysis for the three biological replicates of adr-1(-);adr-2(-) animals are listed in columns I, J and K . "	"Supplemental Table 4: Differentially expressed genes in adr-1(-);adr-2(-) animals grown on OP40 for 7 hours compared to wildtype animals grown on OP50 for 7 hours (Related to Figure 4). DESeq2 was used to identify differentially expressed genes in adr-1(-);adr-2(-) double mutant animals compared to wildtype animals. p-value<0.05 were taken as targets misregulated in absence of ad-1 and adr-2. Differentially expressed genes are listed as Wormbase annotations in Column A, Gene name in Column B. The log2fold change values, raw p values and the adjusted p values as obtained from the DESeq2 analysis are shown in columns C, D, and E, respectively. Read counts from the DESeq2 analysis for the three biological replicates of wildtype are listed in columns F, G and H. Read counts from the DESeq2 analysis for the three biological replicates of adr-1(-);adr-2(-) animals are listed in columns I, J and K . "	"Supplemental Table 4: Differentially expressed genes in adr-1(-);adr-2(-) animals grown on OP40 for 7 hours compared to wildtype animals grown on OP50 for 7 hours (Related to Figure 4). DESeq2 was used to identify differentially expressed genes in adr-1(-);adr-2(-) double mutant animals compared to wildtype animals. p-value<0.05 were taken as targets misregulated in absence of ad-1 and adr-2. Differentially expressed genes are listed as Wormbase annotations in Column A, Gene name in Column B. The log2fold change values, raw p values and the adjusted p values as obtained from the DESeq2 analysis are shown in columns C, D, and E, respectively. Read counts from the DESeq2 analysis for the three biological replicates of wildtype are listed in columns F, G and H. Read counts from the DESeq2 analysis for the three biological replicates of adr-1(-);adr-2(-) animals are listed in columns I, J and K . "	"Supplemental Table 4: Differentially expressed genes in adr-1(-);adr-2(-) animals grown on OP40 for 7 hours compared to wildtype animals grown on OP50 for 7 hours (Related to Figure 4). DESeq2 was used to identify differentially expressed genes in adr-1(-);adr-2(-) double mutant animals compared to wildtype animals. p-value<0.05 were taken as targets misregulated in absence of ad-1 and adr-2. Differentially expressed genes are listed as Wormbase annotations in Column A, Gene name in Column B. The log2fold change values, raw p values and the adjusted p values as obtained from the DESeq2 analysis are shown in columns C, D, and E, respectively. Read counts from the DESeq2 analysis for the three biological replicates of wildtype are listed in columns F, G and H. Read counts from the DESeq2 analysis for the three biological replicates of adr-1(-);adr-2(-) animals are listed in columns I, J and K . "	"Supplemental Table 4: Differentially expressed genes in adr-1(-);adr-2(-) animals grown on OP40 for 7 hours compared to wildtype animals grown on OP50 for 7 hours (Related to Figure 4). DESeq2 was used to identify differentially expressed genes in adr-1(-);adr-2(-) double mutant animals compared to wildtype animals. p-value<0.05 were taken as targets misregulated in absence of ad-1 and adr-2. Differentially expressed genes are listed as Wormbase annotations in Column A, Gene name in Column B. The log2fold change values, raw p values and the adjusted p values as obtained from the DESeq2 analysis are shown in columns C, D, and E, respectively. Read counts from the DESeq2 analysis for the three biological replicates of wildtype are listed in columns F, G and H. Read counts from the DESeq2 analysis for the three biological replicates of adr-1(-);adr-2(-) animals are listed in columns I, J and K . "	"Supplemental Table 4: Differentially expressed genes in adr-1(-);adr-2(-) animals grown on OP40 for 7 hours compared to wildtype animals grown on OP50 for 7 hours (Related to Figure 4). DESeq2 was used to identify differentially expressed genes in adr-1(-);adr-2(-) double mutant animals compared to wildtype animals. p-value<0.05 were taken as targets misregulated in absence of ad-1 and adr-2. Differentially expressed genes are listed as Wormbase annotations in Column A, Gene name in Column B. The log2fold change values, raw p values and the adjusted p values as obtained from the DESeq2 analysis are shown in columns C, D, and E, respectively. Read counts from the DESeq2 analysis for the three biological replicates of wildtype are listed in columns F, G and H. Read counts from the DESeq2 analysis for the three biological replicates of adr-1(-);adr-2(-) animals are listed in columns I, J and K . "	"Supplemental Table 4: Differentially expressed genes in adr-1(-);adr-2(-) animals grown on OP40 for 7 hours compared to wildtype animals grown on OP50 for 7 hours (Related to Figure 4). DESeq2 was used to identify differentially expressed genes in adr-1(-);adr-2(-) double mutant animals compared to wildtype animals. p-value<0.05 were taken as targets misregulated in absence of ad-1 and adr-2. Differentially expressed genes are listed as Wormbase annotations in Column A, Gene name in Column B. The log2fold change values, raw p values and the adjusted p values as obtained from the DESeq2 analysis are shown in columns C, D, and E, respectively. Read counts from the DESeq2 analysis for the three biological replicates of wildtype are listed in columns F, G and H. Read counts from the DESeq2 analysis for the three biological replicates of adr-1(-);adr-2(-) animals are listed in columns I, J and K . "	"Supplemental Table 4: Differentially expressed genes in adr-1(-);adr-2(-) animals grown on OP40 for 7 hours compared to wildtype animals grown on OP50 for 7 hours (Related to Figure 4). DESeq2 was used to identify differentially expressed genes in adr-1(-);adr-2(-) double mutant animals compared to wildtype animals. p-value<0.05 were taken as targets misregulated in absence of ad-1 and adr-2. Differentially expressed genes are listed as Wormbase annotations in Column A, Gene name in Column B. The log2fold change values, raw p values and the adjusted p values as obtained from the DESeq2 analysis are shown in columns C, D, and E, respectively. Read counts from the DESeq2 analysis for the three biological replicates of wildtype are listed in columns F, G and H. Read counts from the DESeq2 analysis for the three biological replicates of adr-1(-);adr-2(-) animals are listed in columns I, J and K . "	"Supplemental Table 4: Differentially expressed genes in adr-1(-);adr-2(-) animals grown on OP40 for 7 hours compared to wildtype animals grown on OP50 for 7 hours (Related to Figure 4). DESeq2 was used to identify differentially expressed genes in adr-1(-);adr-2(-) double mutant animals compared to wildtype animals. p-value<0.05 were taken as targets misregulated in absence of ad-1 and adr-2. Differentially expressed genes are listed as Wormbase annotations in Column A, Gene name in Column B. The log2fold change values, raw p values and the adjusted p values as obtained from the DESeq2 analysis are shown in columns C, D, and E, respectively. Read counts from the DESeq2 analysis for the three biological replicates of wildtype are listed in columns F, G and H. Read counts from the DESeq2 analysis for the three biological replicates of adr-1(-);adr-2(-) animals are listed in columns I, J and K . "	"Supplemental Table 4: Differentially expressed genes in adr-1(-);adr-2(-) animals grown on OP40 for 7 hours compared to wildtype animals grown on OP50 for 7 hours (Related to Figure 4). DESeq2 was used to identify differentially expressed genes in adr-1(-);adr-2(-) double mutant animals compared to wildtype animals. p-value<0.05 were taken as targets misregulated in absence of ad-1 and adr-2. Differentially expressed genes are listed as Wormbase annotations in Column A, Gene name in Column B. The log2fold change values, raw p values and the adjusted p values as obtained from the DESeq2 analysis are shown in columns C, D, and E, respectively. Read counts from the DESeq2 analysis for the three biological replicates of wildtype are listed in columns F, G and H. Read counts from the DESeq2 analysis for the three biological replicates of adr-1(-);adr-2(-) animals are listed in columns I, J and K . "	"Supplemental Table 4: Differentially expressed genes in adr-1(-);adr-2(-) animals grown on OP40 for 7 hours compared to wildtype animals grown on OP50 for 7 hours (Related to Figure 4). DESeq2 was used to identify differentially expressed genes in adr-1(-);adr-2(-) double mutant animals compared to wildtype animals. p-value<0.05 were taken as targets misregulated in absence of ad-1 and adr-2. Differentially expressed genes are listed as Wormbase annotations in Column A, Gene name in Column B. The log2fold change values, raw p values and the adjusted p values as obtained from the DESeq2 analysis are shown in columns C, D, and E, respectively. Read counts from the DESeq2 analysis for the three biological replicates of wildtype are listed in columns F, G and H. Read counts from the DESeq2 analysis for the three biological replicates of adr-1(-);adr-2(-) animals are listed in columns I, J and K . "	"Supplemental Table 4: Differentially expressed genes in adr-1(-);adr-2(-) animals grown on OP40 for 7 hours compared to wildtype animals grown on OP50 for 7 hours (Related to Figure 4). DESeq2 was used to identify differentially expressed genes in adr-1(-);adr-2(-) double mutant animals compared to wildtype animals. p-value<0.05 were taken as targets misregulated in absence of ad-1 and adr-2. Differentially expressed genes are listed as Wormbase annotations in Column A, Gene name in Column B. The log2fold change values, raw p values and the adjusted p values as obtained from the DESeq2 analysis are shown in columns C, D, and E, respectively. Read counts from the DESeq2 analysis for the three biological replicates of wildtype are listed in columns F, G and H. Read counts from the DESeq2 analysis for the three biological replicates of adr-1(-);adr-2(-) animals are listed in columns I, J and K . "	"Supplemental Table 4: Differentially expressed genes in adr-1(-);adr-2(-) animals grown on OP40 for 7 hours compared to wildtype animals grown on OP50 for 7 hours (Related to Figure 4). DESeq2 was used to identify differentially expressed genes in adr-1(-);adr-2(-) double mutant animals compared to wildtype animals. p-value<0.05 were taken as targets misregulated in absence of ad-1 and adr-2. Differentially expressed genes are listed as Wormbase annotations in Column A, Gene name in Column B. The log2fold change values, raw p values and the adjusted p values as obtained from the DESeq2 analysis are shown in columns C, D, and E, respectively. Read counts from the DESeq2 analysis for the three biological replicates of wildtype are listed in columns F, G and H. Read counts from the DESeq2 analysis for the three biological replicates of adr-1(-);adr-2(-) animals are listed in columns I, J and K . "	"Supplemental Table 4: Differentially expressed genes in adr-1(-);adr-2(-) animals grown on OP40 for 7 hours compared to wildtype animals grown on OP50 for 7 hours (Related to Figure 4). DESeq2 was used to identify differentially expressed genes in adr-1(-);adr-2(-) double mutant animals compared to wildtype animals. p-value<0.05 were taken as targets misregulated in absence of ad-1 and adr-2. Differentially expressed genes are listed as Wormbase annotations in Column A, Gene name in Column B. The log2fold change values, raw p values and the adjusted p values as obtained from the DESeq2 analysis are shown in columns C, D, and E, respectively. Read counts from the DESeq2 analysis for the three biological replicates of wildtype are listed in columns F, G and H. Read counts from the DESeq2 analysis for the three biological replicates of adr-1(-);adr-2(-) animals are listed in columns I, J and K . "	"Supplemental Table 4: Differentially expressed genes in adr-1(-);adr-2(-) animals grown on OP40 for 7 hours compared to wildtype animals grown on OP50 for 7 hours (Related to Figure 4). DESeq2 was used to identify differentially expressed genes in adr-1(-);adr-2(-) double mutant animals compared to wildtype animals. p-value<0.05 were taken as targets misregulated in absence of ad-1 and adr-2. Differentially expressed genes are listed as Wormbase annotations in Column A, Gene name in Column B. The log2fold change values, raw p values and the adjusted p values as obtained from the DESeq2 analysis are shown in columns C, D, and E, respectively. Read counts from the DESeq2 analysis for the three biological replicates of wildtype are listed in columns F, G and H. Read counts from the DESeq2 analysis for the three biological replicates of adr-1(-);adr-2(-) animals are listed in columns I, J and K . "	"Supplemental Table 4: Differentially expressed genes in adr-1(-);adr-2(-) animals grown on OP40 for 7 hours compared to wildtype animals grown on OP50 for 7 hours (Related to Figure 4). DESeq2 was used to identify differentially expressed genes in adr-1(-);adr-2(-) double mutant animals compared to wildtype animals. p-value<0.05 were taken as targets misregulated in absence of ad-1 and adr-2. Differentially expressed genes are listed as Wormbase annotations in Column A, Gene name in Column B. The log2fold change values, raw p values and the adjusted p values as obtained from the DESeq2 analysis are shown in columns C, D, and E, respectively. Read counts from the DESeq2 analysis for the three biological replicates of wildtype are listed in columns F, G and H. Read counts from the DESeq2 analysis for the three biological replicates of adr-1(-);adr-2(-) animals are listed in columns I, J and K . "	"Supplemental Table 4: Differentially expressed genes in adr-1(-);adr-2(-) animals grown on OP40 for 7 hours compared to wildtype animals grown on OP50 for 7 hours (Related to Figure 4). DESeq2 was used to identify differentially expressed genes in adr-1(-);adr-2(-) double mutant animals compared to wildtype animals. p-value<0.05 were taken as targets misregulated in absence of ad-1 and adr-2. Differentially expressed genes are listed as Wormbase annotations in Column A, Gene name in Column B. The log2fold change values, raw p values and the adjusted p values as obtained from the DESeq2 analysis are shown in columns C, D, and E, respectively. Read counts from the DESeq2 analysis for the three biological replicates of wildtype are listed in columns F, G and H. Read counts from the DESeq2 analysis for the three biological replicates of adr-1(-);adr-2(-) animals are listed in columns I, J and K . "	"Supplemental Table 4: Differentially expressed genes in adr-1(-);adr-2(-) animals grown on OP40 for 7 hours compared to wildtype animals grown on OP50 for 7 hours (Related to Figure 4). DESeq2 was used to identify differentially expressed genes in adr-1(-);adr-2(-) double mutant animals compared to wildtype animals. p-value<0.05 were taken as targets misregulated in absence of ad-1 and adr-2. Differentially expressed genes are listed as Wormbase annotations in Column A, Gene name in Column B. The log2fold change values, raw p values and the adjusted p values as obtained from the DESeq2 analysis are shown in columns C, D, and E, respectively. Read counts from the DESeq2 analysis for the three biological replicates of wildtype are listed in columns F, G and H. Read counts from the DESeq2 analysis for the three biological replicates of adr-1(-);adr-2(-) animals are listed in columns I, J and K . "	"Supplemental Table 4: Differentially expressed genes in adr-1(-);adr-2(-) animals grown on OP40 for 7 hours compared to wildtype animals grown on OP50 for 7 hours (Related to Figure 4). DESeq2 was used to identify differentially expressed genes in adr-1(-);adr-2(-) double mutant animals compared to wildtype animals. p-value<0.05 were taken as targets misregulated in absence of ad-1 and adr-2. Differentially expressed genes are listed as Wormbase annotations in Column A, Gene name in Column B. The log2fold change values, raw p values and the adjusted p values as obtained from the DESeq2 analysis are shown in columns C, D, and E, respectively. Read counts from the DESeq2 analysis for the three biological replicates of wildtype are listed in columns F, G and H. Read counts from the DESeq2 analysis for the three biological replicates of adr-1(-);adr-2(-) animals are listed in columns I, J and K . "	"Supplemental Table 4: Differentially expressed genes in adr-1(-);adr-2(-) animals grown on OP40 for 7 hours compared to wildtype animals grown on OP50 for 7 hours (Related to Figure 4). DESeq2 was used to identify differentially expressed genes in adr-1(-);adr-2(-) double mutant animals compared to wildtype animals. p-value<0.05 were taken as targets misregulated in absence of ad-1 and adr-2. Differentially expressed genes are listed as Wormbase annotations in Column A, Gene name in Column B. The log2fold change values, raw p values and the adjusted p values as obtained from the DESeq2 analysis are shown in columns C, D, and E, respectively. Read counts from the DESeq2 analysis for the three biological replicates of wildtype are listed in columns F, G and H. Read counts from the DESeq2 analysis for the three biological replicates of adr-1(-);adr-2(-) animals are listed in columns I, J and K . "	"Supplemental Table 4: Differentially expressed genes in adr-1(-);adr-2(-) animals grown on OP40 for 7 hours compared to wildtype animals grown on OP50 for 7 hours (Related to Figure 4). DESeq2 was used to identify differentially expressed genes in adr-1(-);adr-2(-) double mutant animals compared to wildtype animals. p-value<0.05 were taken as targets misregulated in absence of ad-1 and adr-2. Differentially expressed genes are listed as Wormbase annotations in Column A, Gene name in Column B. The log2fold change values, raw p values and the adjusted p values as obtained from the DESeq2 analysis are shown in columns C, D, and E, respectively. Read counts from the DESeq2 analysis for the three biological replicates of wildtype are listed in columns F, G and H. Read counts from the DESeq2 analysis for the three biological replicates of adr-1(-);adr-2(-) animals are listed in columns I, J and K . "	"Supplemental Table 4: Differentially expressed genes in adr-1(-);adr-2(-) animals grown on OP40 for 7 hours compared to wildtype animals grown on OP50 for 7 hours (Related to Figure 4). DESeq2 was used to identify differentially expressed genes in adr-1(-);adr-2(-) double mutant animals compared to wildtype animals. p-value<0.05 were taken as targets misregulated in absence of ad-1 and adr-2. Differentially expressed genes are listed as Wormbase annotations in Column A, Gene name in Column B. The log2fold change values, raw p values and the adjusted p values as obtained from the DESeq2 analysis are shown in columns C, D, and E, respectively. Read counts from the DESeq2 analysis for the three biological replicates of wildtype are listed in columns F, G and H. Read counts from the DESeq2 analysis for the three biological replicates of adr-1(-);adr-2(-) animals are listed in columns I, J and K . "	"Supplemental Table 4: Differentially expressed genes in adr-1(-);adr-2(-) animals grown on OP40 for 7 hours compared to wildtype animals grown on OP50 for 7 hours (Related to Figure 4). DESeq2 was used to identify differentially expressed genes in adr-1(-);adr-2(-) double mutant animals compared to wildtype animals. p-value<0.05 were taken as targets misregulated in absence of ad-1 and adr-2. Differentially expressed genes are listed as Wormbase annotations in Column A, Gene name in Column B. The log2fold change values, raw p values and the adjusted p values as obtained from the DESeq2 analysis are shown in columns C, D, and E, respectively. Read counts from the DESeq2 analysis for the three biological replicates of wildtype are listed in columns F, G and H. Read counts from the DESeq2 analysis for the three biological replicates of adr-1(-);adr-2(-) animals are listed in columns I, J and K . "	"Supplemental Table 4: Differentially expressed genes in adr-1(-);adr-2(-) animals grown on OP40 for 7 hours compared to wildtype animals grown on OP50 for 7 hours (Related to Figure 4). DESeq2 was used to identify differentially expressed genes in adr-1(-);adr-2(-) double mutant animals compared to wildtype animals. p-value<0.05 were taken as targets misregulated in absence of ad-1 and adr-2. Differentially expressed genes are listed as Wormbase annotations in Column A, Gene name in Column B. The log2fold change values, raw p values and the adjusted p values as obtained from the DESeq2 analysis are shown in columns C, D, and E, respectively. Read counts from the DESeq2 analysis for the three biological replicates of wildtype are listed in columns F, G and H. Read counts from the DESeq2 analysis for the three biological replicates of adr-1(-);adr-2(-) animals are listed in columns I, J and K . "	"Supplemental Table 4: Differentially expressed genes in adr-1(-);adr-2(-) animals grown on OP40 for 7 hours compared to wildtype animals grown on OP50 for 7 hours (Related to Figure 4). DESeq2 was used to identify differentially expressed genes in adr-1(-);adr-2(-) double mutant animals compared to wildtype animals. p-value<0.05 were taken as targets misregulated in absence of ad-1 and adr-2. Differentially expressed genes are listed as Wormbase annotations in Column A, Gene name in Column B. The log2fold change values, raw p values and the adjusted p values as obtained from the DESeq2 analysis are shown in columns C, D, and E, respectively. Read counts from the DESeq2 analysis for the three biological replicates of wildtype are listed in columns F, G and H. Read counts from the DESeq2 analysis for the three biological replicates of adr-1(-);adr-2(-) animals are listed in columns I, J and K . "	"Supplemental Table 4: Differentially expressed genes in adr-1(-);adr-2(-) animals grown on OP40 for 7 hours compared to wildtype animals grown on OP50 for 7 hours (Related to Figure 4). DESeq2 was used to identify differentially expressed genes in adr-1(-);adr-2(-) double mutant animals compared to wildtype animals. p-value<0.05 were taken as targets misregulated in absence of ad-1 and adr-2. Differentially expressed genes are listed as Wormbase annotations in Column A, Gene name in Column B. The log2fold change values, raw p values and the adjusted p values as obtained from the DESeq2 analysis are shown in columns C, D, and E, respectively. Read counts from the DESeq2 analysis for the three biological replicates of wildtype are listed in columns F, G and H. Read counts from the DESeq2 analysis for the three biological replicates of adr-1(-);adr-2(-) animals are listed in columns I, J and K . "	"Supplemental Table 4: Differentially expressed genes in adr-1(-);adr-2(-) animals grown on OP40 for 7 hours compared to wildtype animals grown on OP50 for 7 hours (Related to Figure 4). DESeq2 was used to identify differentially expressed genes in adr-1(-);adr-2(-) double mutant animals compared to wildtype animals. p-value<0.05 were taken as targets misregulated in absence of ad-1 and adr-2. Differentially expressed genes are listed as Wormbase annotations in Column A, Gene name in Column B. The log2fold change values, raw p values and the adjusted p values as obtained from the DESeq2 analysis are shown in columns C, D, and E, respectively. Read counts from the DESeq2 analysis for the three biological replicates of wildtype are listed in columns F, G and H. Read counts from the DESeq2 analysis for the three biological replicates of adr-1(-);adr-2(-) animals are listed in columns I, J and K . "	"Supplemental Table 4: Differentially expressed genes in adr-1(-);adr-2(-) animals grown on OP40 for 7 hours compared to wildtype animals grown on OP50 for 7 hours (Related to Figure 4). DESeq2 was used to identify differentially expressed genes in adr-1(-);adr-2(-) double mutant animals compared to wildtype animals. p-value<0.05 were taken as targets misregulated in absence of ad-1 and adr-2. Differentially expressed genes are listed as Wormbase annotations in Column A, Gene name in Column B. The log2fold change values, raw p values and the adjusted p values as obtained from the DESeq2 analysis are shown in columns C, D, and E, respectively. Read counts from the DESeq2 analysis for the three biological replicates of wildtype are listed in columns F, G and H. Read counts from the DESeq2 analysis for the three biological replicates of adr-1(-);adr-2(-) animals are listed in columns I, J and K . "	"Supplemental Table 4: Differentially expressed genes in adr-1(-);adr-2(-) animals grown on OP40 for 7 hours compared to wildtype animals grown on OP50 for 7 hours (Related to Figure 4). DESeq2 was used to identify differentially expressed genes in adr-1(-);adr-2(-) double mutant animals compared to wildtype animals. p-value<0.05 were taken as targets misregulated in absence of ad-1 and adr-2. Differentially expressed genes are listed as Wormbase annotations in Column A, Gene name in Column B. The log2fold change values, raw p values and the adjusted p values as obtained from the DESeq2 analysis are shown in columns C, D, and E, respectively. Read counts from the DESeq2 analysis for the three biological replicates of wildtype are listed in columns F, G and H. Read counts from the DESeq2 analysis for the three biological replicates of adr-1(-);adr-2(-) animals are listed in columns I, J and K . "	"Supplemental Table 4: Differentially expressed genes in adr-1(-);adr-2(-) animals grown on OP40 for 7 hours compared to wildtype animals grown on OP50 for 7 hours (Related to Figure 4). DESeq2 was used to identify differentially expressed genes in adr-1(-);adr-2(-) double mutant animals compared to wildtype animals. p-value<0.05 were taken as targets misregulated in absence of ad-1 and adr-2. Differentially expressed genes are listed as Wormbase annotations in Column A, Gene name in Column B. The log2fold change values, raw p values and the adjusted p values as obtained from the DESeq2 analysis are shown in columns C, D, and E, respectively. Read counts from the DESeq2 analysis for the three biological replicates of wildtype are listed in columns F, G and H. Read counts from the DESeq2 analysis for the three biological replicates of adr-1(-);adr-2(-) animals are listed in columns I, J and K . "	"Supplemental Table 4: Differentially expressed genes in adr-1(-);adr-2(-) animals grown on OP40 for 7 hours compared to wildtype animals grown on OP50 for 7 hours (Related to Figure 4). DESeq2 was used to identify differentially expressed genes in adr-1(-);adr-2(-) double mutant animals compared to wildtype animals. p-value<0.05 were taken as targets misregulated in absence of ad-1 and adr-2. Differentially expressed genes are listed as Wormbase annotations in Column A, Gene name in Column B. The log2fold change values, raw p values and the adjusted p values as obtained from the DESeq2 analysis are shown in columns C, D, and E, respectively. Read counts from the DESeq2 analysis for the three biological replicates of wildtype are listed in columns F, G and H. Read counts from the DESeq2 analysis for the three biological replicates of adr-1(-);adr-2(-) animals are listed in columns I, J and K . "	"Supplemental Table 4: Differentially expressed genes in adr-1(-);adr-2(-) animals grown on OP40 for 7 hours compared to wildtype animals grown on OP50 for 7 hours (Related to Figure 4). DESeq2 was used to identify differentially expressed genes in adr-1(-);adr-2(-) double mutant animals compared to wildtype animals. p-value<0.05 were taken as targets misregulated in absence of ad-1 and adr-2. Differentially expressed genes are listed as Wormbase annotations in Column A, Gene name in Column B. The log2fold change values, raw p values and the adjusted p values as obtained from the DESeq2 analysis are shown in columns C, D, and E, respectively. Read counts from the DESeq2 analysis for the three biological replicates of wildtype are listed in columns F, G and H. Read counts from the DESeq2 analysis for the three biological replicates of adr-1(-);adr-2(-) animals are listed in columns I, J and K . "	"Supplemental Table 4: Differentially expressed genes in adr-1(-);adr-2(-) animals grown on OP40 for 7 hours compared to wildtype animals grown on OP50 for 7 hours (Related to Figure 4). DESeq2 was used to identify differentially expressed genes in adr-1(-);adr-2(-) double mutant animals compared to wildtype animals. p-value<0.05 were taken as targets misregulated in absence of ad-1 and adr-2. Differentially expressed genes are listed as Wormbase annotations in Column A, Gene name in Column B. The log2fold change values, raw p values and the adjusted p values as obtained from the DESeq2 analysis are shown in columns C, D, and E, respectively. Read counts from the DESeq2 analysis for the three biological replicates of wildtype are listed in columns F, G and H. Read counts from the DESeq2 analysis for the three biological replicates of adr-1(-);adr-2(-) animals are listed in columns I, J and K . "	"Supplemental Table 4: Differentially expressed genes in adr-1(-);adr-2(-) animals grown on OP40 for 7 hours compared to wildtype animals grown on OP50 for 7 hours (Related to Figure 4). DESeq2 was used to identify differentially expressed genes in adr-1(-);adr-2(-) double mutant animals compared to wildtype animals. p-value<0.05 were taken as targets misregulated in absence of ad-1 and adr-2. Differentially expressed genes are listed as Wormbase annotations in Column A, Gene name in Column B. The log2fold change values, raw p values and the adjusted p values as obtained from the DESeq2 analysis are shown in columns C, D, and E, respectively. Read counts from the DESeq2 analysis for the three biological replicates of wildtype are listed in columns F, G and H. Read counts from the DESeq2 analysis for the three biological replicates of adr-1(-);adr-2(-) animals are listed in columns I, J and K . "	"Supplemental Table 4: Differentially expressed genes in adr-1(-);adr-2(-) animals grown on OP40 for 7 hours compared to wildtype animals grown on OP50 for 7 hours (Related to Figure 4). DESeq2 was used to identify differentially expressed genes in adr-1(-);adr-2(-) double mutant animals compared to wildtype animals. p-value<0.05 were taken as targets misregulated in absence of ad-1 and adr-2. Differentially expressed genes are listed as Wormbase annotations in Column A, Gene name in Column B. The log2fold change values, raw p values and the adjusted p values as obtained from the DESeq2 analysis are shown in columns C, D, and E, respectively. Read counts from the DESeq2 analysis for the three biological replicates of wildtype are listed in columns F, G and H. Read counts from the DESeq2 analysis for the three biological replicates of adr-1(-);adr-2(-) animals are listed in columns I, J and K . "	"Supplemental Table 4: Differentially expressed genes in adr-1(-);adr-2(-) animals grown on OP40 for 7 hours compared to wildtype animals grown on OP50 for 7 hours (Related to Figure 4). DESeq2 was used to identify differentially expressed genes in adr-1(-);adr-2(-) double mutant animals compared to wildtype animals. p-value<0.05 were taken as targets misregulated in absence of ad-1 and adr-2. Differentially expressed genes are listed as Wormbase annotations in Column A, Gene name in Column B. The log2fold change values, raw p values and the adjusted p values as obtained from the DESeq2 analysis are shown in columns C, D, and E, respectively. Read counts from the DESeq2 analysis for the three biological replicates of wildtype are listed in columns F, G and H. Read counts from the DESeq2 analysis for the three biological replicates of adr-1(-);adr-2(-) animals are listed in columns I, J and K . "	"Supplemental Table 4: Differentially expressed genes in adr-1(-);adr-2(-) animals grown on OP40 for 7 hours compared to wildtype animals grown on OP50 for 7 hours (Related to Figure 4). DESeq2 was used to identify differentially expressed genes in adr-1(-);adr-2(-) double mutant animals compared to wildtype animals. p-value<0.05 were taken as targets misregulated in absence of ad-1 and adr-2. Differentially expressed genes are listed as Wormbase annotations in Column A, Gene name in Column B. The log2fold change values, raw p values and the adjusted p values as obtained from the DESeq2 analysis are shown in columns C, D, and E, respectively. Read counts from the DESeq2 analysis for the three biological replicates of wildtype are listed in columns F, G and H. Read counts from the DESeq2 analysis for the three biological replicates of adr-1(-);adr-2(-) animals are listed in columns I, J and K . "	"Supplemental Table 4: Differentially expressed genes in adr-1(-);adr-2(-) animals grown on OP40 for 7 hours compared to wildtype animals grown on OP50 for 7 hours (Related to Figure 4). DESeq2 was used to identify differentially expressed genes in adr-1(-);adr-2(-) double mutant animals compared to wildtype animals. p-value<0.05 were taken as targets misregulated in absence of ad-1 and adr-2. Differentially expressed genes are listed as Wormbase annotations in Column A, Gene name in Column B. The log2fold change values, raw p values and the adjusted p values as obtained from the DESeq2 analysis are shown in columns C, D, and E, respectively. Read counts from the DESeq2 analysis for the three biological replicates of wildtype are listed in columns F, G and H. Read counts from the DESeq2 analysis for the three biological replicates of adr-1(-);adr-2(-) animals are listed in columns I, J and K . "	"Supplemental Table 4: Differentially expressed genes in adr-1(-);adr-2(-) animals grown on OP40 for 7 hours compared to wildtype animals grown on OP50 for 7 hours (Related to Figure 4). DESeq2 was used to identify differentially expressed genes in adr-1(-);adr-2(-) double mutant animals compared to wildtype animals. p-value<0.05 were taken as targets misregulated in absence of ad-1 and adr-2. Differentially expressed genes are listed as Wormbase annotations in Column A, Gene name in Column B. The log2fold change values, raw p values and the adjusted p values as obtained from the DESeq2 analysis are shown in columns C, D, and E, respectively. Read counts from the DESeq2 analysis for the three biological replicates of wildtype are listed in columns F, G and H. Read counts from the DESeq2 analysis for the three biological replicates of adr-1(-);adr-2(-) animals are listed in columns I, J and K . "	"Supplemental Table 4: Differentially expressed genes in adr-1(-);adr-2(-) animals grown on OP40 for 7 hours compared to wildtype animals grown on OP50 for 7 hours (Related to Figure 4). DESeq2 was used to identify differentially expressed genes in adr-1(-);adr-2(-) double mutant animals compared to wildtype animals. p-value<0.05 were taken as targets misregulated in absence of ad-1 and adr-2. Differentially expressed genes are listed as Wormbase annotations in Column A, Gene name in Column B. The log2fold change values, raw p values and the adjusted p values as obtained from the DESeq2 analysis are shown in columns C, D, and E, respectively. Read counts from the DESeq2 analysis for the three biological replicates of wildtype are listed in columns F, G and H. Read counts from the DESeq2 analysis for the three biological replicates of adr-1(-);adr-2(-) animals are listed in columns I, J and K . "	"Supplemental Table 4: Differentially expressed genes in adr-1(-);adr-2(-) animals grown on OP40 for 7 hours compared to wildtype animals grown on OP50 for 7 hours (Related to Figure 4). DESeq2 was used to identify differentially expressed genes in adr-1(-);adr-2(-) double mutant animals compared to wildtype animals. p-value<0.05 were taken as targets misregulated in absence of ad-1 and adr-2. Differentially expressed genes are listed as Wormbase annotations in Column A, Gene name in Column B. The log2fold change values, raw p values and the adjusted p values as obtained from the DESeq2 analysis are shown in columns C, D, and E, respectively. Read counts from the DESeq2 analysis for the three biological replicates of wildtype are listed in columns F, G and H. Read counts from the DESeq2 analysis for the three biological replicates of adr-1(-);adr-2(-) animals are listed in columns I, J and K . "	"Supplemental Table 4: Differentially expressed genes in adr-1(-);adr-2(-) animals grown on OP40 for 7 hours compared to wildtype animals grown on OP50 for 7 hours (Related to Figure 4). DESeq2 was used to identify differentially expressed genes in adr-1(-);adr-2(-) double mutant animals compared to wildtype animals. p-value<0.05 were taken as targets misregulated in absence of ad-1 and adr-2. Differentially expressed genes are listed as Wormbase annotations in Column A, Gene name in Column B. The log2fold change values, raw p values and the adjusted p values as obtained from the DESeq2 analysis are shown in columns C, D, and E, respectively. Read counts from the DESeq2 analysis for the three biological replicates of wildtype are listed in columns F, G and H. Read counts from the DESeq2 analysis for the three biological replicates of adr-1(-);adr-2(-) animals are listed in columns I, J and K . "	"Supplemental Table 4: Differentially expressed genes in adr-1(-);adr-2(-) animals grown on OP40 for 7 hours compared to wildtype animals grown on OP50 for 7 hours (Related to Figure 4). DESeq2 was used to identify differentially expressed genes in adr-1(-);adr-2(-) double mutant animals compared to wildtype animals. p-value<0.05 were taken as targets misregulated in absence of ad-1 and adr-2. Differentially expressed genes are listed as Wormbase annotations in Column A, Gene name in Column B. The log2fold change values, raw p values and the adjusted p values as obtained from the DESeq2 analysis are shown in columns C, D, and E, respectively. Read counts from the DESeq2 analysis for the three biological replicates of wildtype are listed in columns F, G and H. Read counts from the DESeq2 analysis for the three biological replicates of adr-1(-);adr-2(-) animals are listed in columns I, J and K . "	"Supplemental Table 4: Differentially expressed genes in adr-1(-);adr-2(-) animals grown on OP40 for 7 hours compared to wildtype animals grown on OP50 for 7 hours (Related to Figure 4). DESeq2 was used to identify differentially expressed genes in adr-1(-);adr-2(-) double mutant animals compared to wildtype animals. p-value<0.05 were taken as targets misregulated in absence of ad-1 and adr-2. Differentially expressed genes are listed as Wormbase annotations in Column A, Gene name in Column B. The log2fold change values, raw p values and the adjusted p values as obtained from the DESeq2 analysis are shown in columns C, D, and E, respectively. Read counts from the DESeq2 analysis for the three biological replicates of wildtype are listed in columns F, G and H. Read counts from the DESeq2 analysis for the three biological replicates of adr-1(-);adr-2(-) animals are listed in columns I, J and K . "	"Supplemental Table 4: Differentially expressed genes in adr-1(-);adr-2(-) animals grown on OP40 for 7 hours compared to wildtype animals grown on OP50 for 7 hours (Related to Figure 4). DESeq2 was used to identify differentially expressed genes in adr-1(-);adr-2(-) double mutant animals compared to wildtype animals. p-value<0.05 were taken as targets misregulated in absence of ad-1 and adr-2. Differentially expressed genes are listed as Wormbase annotations in Column A, Gene name in Column B. The log2fold change values, raw p values and the adjusted p values as obtained from the DESeq2 analysis are shown in columns C, D, and E, respectively. Read counts from the DESeq2 analysis for the three biological replicates of wildtype are listed in columns F, G and H. Read counts from the DESeq2 analysis for the three biological replicates of adr-1(-);adr-2(-) animals are listed in columns I, J and K . "	"Supplemental Table 4: Differentially expressed genes in adr-1(-);adr-2(-) animals grown on OP40 for 7 hours compared to wildtype animals grown on OP50 for 7 hours (Related to Figure 4). DESeq2 was used to identify differentially expressed genes in adr-1(-);adr-2(-) double mutant animals compared to wildtype animals. p-value<0.05 were taken as targets misregulated in absence of ad-1 and adr-2. Differentially expressed genes are listed as Wormbase annotations in Column A, Gene name in Column B. The log2fold change values, raw p values and the adjusted p values as obtained from the DESeq2 analysis are shown in columns C, D, and E, respectively. Read counts from the DESeq2 analysis for the three biological replicates of wildtype are listed in columns F, G and H. Read counts from the DESeq2 analysis for the three biological replicates of adr-1(-);adr-2(-) animals are listed in columns I, J and K . "	"Supplemental Table 4: Differentially expressed genes in adr-1(-);adr-2(-) animals grown on OP40 for 7 hours compared to wildtype animals grown on OP50 for 7 hours (Related to Figure 4). DESeq2 was used to identify differentially expressed genes in adr-1(-);adr-2(-) double mutant animals compared to wildtype animals. p-value<0.05 were taken as targets misregulated in absence of ad-1 and adr-2. Differentially expressed genes are listed as Wormbase annotations in Column A, Gene name in Column B. The log2fold change values, raw p values and the adjusted p values as obtained from the DESeq2 analysis are shown in columns C, D, and E, respectively. Read counts from the DESeq2 analysis for the three biological replicates of wildtype are listed in columns F, G and H. Read counts from the DESeq2 analysis for the three biological replicates of adr-1(-);adr-2(-) animals are listed in columns I, J and K . "	"Supplemental Table 4: Differentially expressed genes in adr-1(-);adr-2(-) animals grown on OP40 for 7 hours compared to wildtype animals grown on OP50 for 7 hours (Related to Figure 4). DESeq2 was used to identify differentially expressed genes in adr-1(-);adr-2(-) double mutant animals compared to wildtype animals. p-value<0.05 were taken as targets misregulated in absence of ad-1 and adr-2. Differentially expressed genes are listed as Wormbase annotations in Column A, Gene name in Column B. The log2fold change values, raw p values and the adjusted p values as obtained from the DESeq2 analysis are shown in columns C, D, and E, respectively. Read counts from the DESeq2 analysis for the three biological replicates of wildtype are listed in columns F, G and H. Read counts from the DESeq2 analysis for the three biological replicates of adr-1(-);adr-2(-) animals are listed in columns I, J and K . "	"Supplemental Table 4: Differentially expressed genes in adr-1(-);adr-2(-) animals grown on OP40 for 7 hours compared to wildtype animals grown on OP50 for 7 hours (Related to Figure 4). DESeq2 was used to identify differentially expressed genes in adr-1(-);adr-2(-) double mutant animals compared to wildtype animals. p-value<0.05 were taken as targets misregulated in absence of ad-1 and adr-2. Differentially expressed genes are listed as Wormbase annotations in Column A, Gene name in Column B. The log2fold change values, raw p values and the adjusted p values as obtained from the DESeq2 analysis are shown in columns C, D, and E, respectively. Read counts from the DESeq2 analysis for the three biological replicates of wildtype are listed in columns F, G and H. Read counts from the DESeq2 analysis for the three biological replicates of adr-1(-);adr-2(-) animals are listed in columns I, J and K . "	"Supplemental Table 4: Differentially expressed genes in adr-1(-);adr-2(-) animals grown on OP40 for 7 hours compared to wildtype animals grown on OP50 for 7 hours (Related to Figure 4). DESeq2 was used to identify differentially expressed genes in adr-1(-);adr-2(-) double mutant animals compared to wildtype animals. p-value<0.05 were taken as targets misregulated in absence of ad-1 and adr-2. Differentially expressed genes are listed as Wormbase annotations in Column A, Gene name in Column B. The log2fold change values, raw p values and the adjusted p values as obtained from the DESeq2 analysis are shown in columns C, D, and E, respectively. Read counts from the DESeq2 analysis for the three biological replicates of wildtype are listed in columns F, G and H. Read counts from the DESeq2 analysis for the three biological replicates of adr-1(-);adr-2(-) animals are listed in columns I, J and K . "	"Supplemental Table 4: Differentially expressed genes in adr-1(-);adr-2(-) animals grown on OP40 for 7 hours compared to wildtype animals grown on OP50 for 7 hours (Related to Figure 4). DESeq2 was used to identify differentially expressed genes in adr-1(-);adr-2(-) double mutant animals compared to wildtype animals. p-value<0.05 were taken as targets misregulated in absence of ad-1 and adr-2. Differentially expressed genes are listed as Wormbase annotations in Column A, Gene name in Column B. The log2fold change values, raw p values and the adjusted p values as obtained from the DESeq2 analysis are shown in columns C, D, and E, respectively. Read counts from the DESeq2 analysis for the three biological replicates of wildtype are listed in columns F, G and H. Read counts from the DESeq2 analysis for the three biological replicates of adr-1(-);adr-2(-) animals are listed in columns I, J and K . "	"Supplemental Table 4: Differentially expressed genes in adr-1(-);adr-2(-) animals grown on OP40 for 7 hours compared to wildtype animals grown on OP50 for 7 hours (Related to Figure 4). DESeq2 was used to identify differentially expressed genes in adr-1(-);adr-2(-) double mutant animals compared to wildtype animals. p-value<0.05 were taken as targets misregulated in absence of ad-1 and adr-2. Differentially expressed genes are listed as Wormbase annotations in Column A, Gene name in Column B. The log2fold change values, raw p values and the adjusted p values as obtained from the DESeq2 analysis are shown in columns C, D, and E, respectively. Read counts from the DESeq2 analysis for the three biological replicates of wildtype are listed in columns F, G and H. Read counts from the DESeq2 analysis for the three biological replicates of adr-1(-);adr-2(-) animals are listed in columns I, J and K . "	"Supplemental Table 4: Differentially expressed genes in adr-1(-);adr-2(-) animals grown on OP40 for 7 hours compared to wildtype animals grown on OP50 for 7 hours (Related to Figure 4). DESeq2 was used to identify differentially expressed genes in adr-1(-);adr-2(-) double mutant animals compared to wildtype animals. p-value<0.05 were taken as targets misregulated in absence of ad-1 and adr-2. Differentially expressed genes are listed as Wormbase annotations in Column A, Gene name in Column B. The log2fold change values, raw p values and the adjusted p values as obtained from the DESeq2 analysis are shown in columns C, D, and E, respectively. Read counts from the DESeq2 analysis for the three biological replicates of wildtype are listed in columns F, G and H. Read counts from the DESeq2 analysis for the three biological replicates of adr-1(-);adr-2(-) animals are listed in columns I, J and K . "	"Supplemental Table 4: Differentially expressed genes in adr-1(-);adr-2(-) animals grown on OP40 for 7 hours compared to wildtype animals grown on OP50 for 7 hours (Related to Figure 4). DESeq2 was used to identify differentially expressed genes in adr-1(-);adr-2(-) double mutant animals compared to wildtype animals. p-value<0.05 were taken as targets misregulated in absence of ad-1 and adr-2. Differentially expressed genes are listed as Wormbase annotations in Column A, Gene name in Column B. The log2fold change values, raw p values and the adjusted p values as obtained from the DESeq2 analysis are shown in columns C, D, and E, respectively. Read counts from the DESeq2 analysis for the three biological replicates of wildtype are listed in columns F, G and H. Read counts from the DESeq2 analysis for the three biological replicates of adr-1(-);adr-2(-) animals are listed in columns I, J and K . "	"Supplemental Table 4: Differentially expressed genes in adr-1(-);adr-2(-) animals grown on OP40 for 7 hours compared to wildtype animals grown on OP50 for 7 hours (Related to Figure 4). DESeq2 was used to identify differentially expressed genes in adr-1(-);adr-2(-) double mutant animals compared to wildtype animals. p-value<0.05 were taken as targets misregulated in absence of ad-1 and adr-2. Differentially expressed genes are listed as Wormbase annotations in Column A, Gene name in Column B. The log2fold change values, raw p values and the adjusted p values as obtained from the DESeq2 analysis are shown in columns C, D, and E, respectively. Read counts from the DESeq2 analysis for the three biological replicates of wildtype are listed in columns F, G and H. Read counts from the DESeq2 analysis for the three biological replicates of adr-1(-);adr-2(-) animals are listed in columns I, J and K . "	"Supplemental Table 4: Differentially expressed genes in adr-1(-);adr-2(-) animals grown on OP40 for 7 hours compared to wildtype animals grown on OP50 for 7 hours (Related to Figure 4). DESeq2 was used to identify differentially expressed genes in adr-1(-);adr-2(-) double mutant animals compared to wildtype animals. p-value<0.05 were taken as targets misregulated in absence of ad-1 and adr-2. Differentially expressed genes are listed as Wormbase annotations in Column A, Gene name in Column B. The log2fold change values, raw p values and the adjusted p values as obtained from the DESeq2 analysis are shown in columns C, D, and E, respectively. Read counts from the DESeq2 analysis for the three biological replicates of wildtype are listed in columns F, G and H. Read counts from the DESeq2 analysis for the three biological replicates of adr-1(-);adr-2(-) animals are listed in columns I, J and K . "	"Supplemental Table 4: Differentially expressed genes in adr-1(-);adr-2(-) animals grown on OP40 for 7 hours compared to wildtype animals grown on OP50 for 7 hours (Related to Figure 4). DESeq2 was used to identify differentially expressed genes in adr-1(-);adr-2(-) double mutant animals compared to wildtype animals. p-value<0.05 were taken as targets misregulated in absence of ad-1 and adr-2. Differentially expressed genes are listed as Wormbase annotations in Column A, Gene name in Column B. The log2fold change values, raw p values and the adjusted p values as obtained from the DESeq2 analysis are shown in columns C, D, and E, respectively. Read counts from the DESeq2 analysis for the three biological replicates of wildtype are listed in columns F, G and H. Read counts from the DESeq2 analysis for the three biological replicates of adr-1(-);adr-2(-) animals are listed in columns I, J and K . "	"Supplemental Table 4: Differentially expressed genes in adr-1(-);adr-2(-) animals grown on OP40 for 7 hours compared to wildtype animals grown on OP50 for 7 hours (Related to Figure 4). DESeq2 was used to identify differentially expressed genes in adr-1(-);adr-2(-) double mutant animals compared to wildtype animals. p-value<0.05 were taken as targets misregulated in absence of ad-1 and adr-2. Differentially expressed genes are listed as Wormbase annotations in Column A, Gene name in Column B. The log2fold change values, raw p values and the adjusted p values as obtained from the DESeq2 analysis are shown in columns C, D, and E, respectively. Read counts from the DESeq2 analysis for the three biological replicates of wildtype are listed in columns F, G and H. Read counts from the DESeq2 analysis for the three biological replicates of adr-1(-);adr-2(-) animals are listed in columns I, J and K . "	"Supplemental Table 4: Differentially expressed genes in adr-1(-);adr-2(-) animals grown on OP40 for 7 hours compared to wildtype animals grown on OP50 for 7 hours (Related to Figure 4). DESeq2 was used to identify differentially expressed genes in adr-1(-);adr-2(-) double mutant animals compared to wildtype animals. p-value<0.05 were taken as targets misregulated in absence of ad-1 and adr-2. Differentially expressed genes are listed as Wormbase annotations in Column A, Gene name in Column B. The log2fold change values, raw p values and the adjusted p values as obtained from the DESeq2 analysis are shown in columns C, D, and E, respectively. Read counts from the DESeq2 analysis for the three biological replicates of wildtype are listed in columns F, G and H. Read counts from the DESeq2 analysis for the three biological replicates of adr-1(-);adr-2(-) animals are listed in columns I, J and K . "	"Supplemental Table 4: Differentially expressed genes in adr-1(-);adr-2(-) animals grown on OP40 for 7 hours compared to wildtype animals grown on OP50 for 7 hours (Related to Figure 4). DESeq2 was used to identify differentially expressed genes in adr-1(-);adr-2(-) double mutant animals compared to wildtype animals. p-value<0.05 were taken as targets misregulated in absence of ad-1 and adr-2. Differentially expressed genes are listed as Wormbase annotations in Column A, Gene name in Column B. The log2fold change values, raw p values and the adjusted p values as obtained from the DESeq2 analysis are shown in columns C, D, and E, respectively. Read counts from the DESeq2 analysis for the three biological replicates of wildtype are listed in columns F, G and H. Read counts from the DESeq2 analysis for the three biological replicates of adr-1(-);adr-2(-) animals are listed in columns I, J and K . "	"Supplemental Table 4: Differentially expressed genes in adr-1(-);adr-2(-) animals grown on OP40 for 7 hours compared to wildtype animals grown on OP50 for 7 hours (Related to Figure 4). DESeq2 was used to identify differentially expressed genes in adr-1(-);adr-2(-) double mutant animals compared to wildtype animals. p-value<0.05 were taken as targets misregulated in absence of ad-1 and adr-2. Differentially expressed genes are listed as Wormbase annotations in Column A, Gene name in Column B. The log2fold change values, raw p values and the adjusted p values as obtained from the DESeq2 analysis are shown in columns C, D, and E, respectively. Read counts from the DESeq2 analysis for the three biological replicates of wildtype are listed in columns F, G and H. Read counts from the DESeq2 analysis for the three biological replicates of adr-1(-);adr-2(-) animals are listed in columns I, J and K . "	"Supplemental Table 4: Differentially expressed genes in adr-1(-);adr-2(-) animals grown on OP40 for 7 hours compared to wildtype animals grown on OP50 for 7 hours (Related to Figure 4). DESeq2 was used to identify differentially expressed genes in adr-1(-);adr-2(-) double mutant animals compared to wildtype animals. p-value<0.05 were taken as targets misregulated in absence of ad-1 and adr-2. Differentially expressed genes are listed as Wormbase annotations in Column A, Gene name in Column B. The log2fold change values, raw p values and the adjusted p values as obtained from the DESeq2 analysis are shown in columns C, D, and E, respectively. Read counts from the DESeq2 analysis for the three biological replicates of wildtype are listed in columns F, G and H. Read counts from the DESeq2 analysis for the three biological replicates of adr-1(-);adr-2(-) animals are listed in columns I, J and K . "	"Supplemental Table 4: Differentially expressed genes in adr-1(-);adr-2(-) animals grown on OP40 for 7 hours compared to wildtype animals grown on OP50 for 7 hours (Related to Figure 4). DESeq2 was used to identify differentially expressed genes in adr-1(-);adr-2(-) double mutant animals compared to wildtype animals. p-value<0.05 were taken as targets misregulated in absence of ad-1 and adr-2. Differentially expressed genes are listed as Wormbase annotations in Column A, Gene name in Column B. The log2fold change values, raw p values and the adjusted p values as obtained from the DESeq2 analysis are shown in columns C, D, and E, respectively. Read counts from the DESeq2 analysis for the three biological replicates of wildtype are listed in columns F, G and H. Read counts from the DESeq2 analysis for the three biological replicates of adr-1(-);adr-2(-) animals are listed in columns I, J and K . "	"Supplemental Table 4: Differentially expressed genes in adr-1(-);adr-2(-) animals grown on OP40 for 7 hours compared to wildtype animals grown on OP50 for 7 hours (Related to Figure 4). DESeq2 was used to identify differentially expressed genes in adr-1(-);adr-2(-) double mutant animals compared to wildtype animals. p-value<0.05 were taken as targets misregulated in absence of ad-1 and adr-2. Differentially expressed genes are listed as Wormbase annotations in Column A, Gene name in Column B. The log2fold change values, raw p values and the adjusted p values as obtained from the DESeq2 analysis are shown in columns C, D, and E, respectively. Read counts from the DESeq2 analysis for the three biological replicates of wildtype are listed in columns F, G and H. Read counts from the DESeq2 analysis for the three biological replicates of adr-1(-);adr-2(-) animals are listed in columns I, J and K . "	"Supplemental Table 4: Differentially expressed genes in adr-1(-);adr-2(-) animals grown on OP40 for 7 hours compared to wildtype animals grown on OP50 for 7 hours (Related to Figure 4). DESeq2 was used to identify differentially expressed genes in adr-1(-);adr-2(-) double mutant animals compared to wildtype animals. p-value<0.05 were taken as targets misregulated in absence of ad-1 and adr-2. Differentially expressed genes are listed as Wormbase annotations in Column A, Gene name in Column B. The log2fold change values, raw p values and the adjusted p values as obtained from the DESeq2 analysis are shown in columns C, D, and E, respectively. Read counts from the DESeq2 analysis for the three biological replicates of wildtype are listed in columns F, G and H. Read counts from the DESeq2 analysis for the three biological replicates of adr-1(-);adr-2(-) animals are listed in columns I, J and K . "	"Supplemental Table 4: Differentially expressed genes in adr-1(-);adr-2(-) animals grown on OP40 for 7 hours compared to wildtype animals grown on OP50 for 7 hours (Related to Figure 4). DESeq2 was used to identify differentially expressed genes in adr-1(-);adr-2(-) double mutant animals compared to wildtype animals. p-value<0.05 were taken as targets misregulated in absence of ad-1 and adr-2. Differentially expressed genes are listed as Wormbase annotations in Column A, Gene name in Column B. The log2fold change values, raw p values and the adjusted p values as obtained from the DESeq2 analysis are shown in columns C, D, and E, respectively. Read counts from the DESeq2 analysis for the three biological replicates of wildtype are listed in columns F, G and H. Read counts from the DESeq2 analysis for the three biological replicates of adr-1(-);adr-2(-) animals are listed in columns I, J and K . "	"Supplemental Table 4: Differentially expressed genes in adr-1(-);adr-2(-) animals grown on OP40 for 7 hours compared to wildtype animals grown on OP50 for 7 hours (Related to Figure 4). DESeq2 was used to identify differentially expressed genes in adr-1(-);adr-2(-) double mutant animals compared to wildtype animals. p-value<0.05 were taken as targets misregulated in absence of ad-1 and adr-2. Differentially expressed genes are listed as Wormbase annotations in Column A, Gene name in Column B. The log2fold change values, raw p values and the adjusted p values as obtained from the DESeq2 analysis are shown in columns C, D, and E, respectively. Read counts from the DESeq2 analysis for the three biological replicates of wildtype are listed in columns F, G and H. Read counts from the DESeq2 analysis for the three biological replicates of adr-1(-);adr-2(-) animals are listed in columns I, J and K . "	"Supplemental Table 4: Differentially expressed genes in adr-1(-);adr-2(-) animals grown on OP40 for 7 hours compared to wildtype animals grown on OP50 for 7 hours (Related to Figure 4). DESeq2 was used to identify differentially expressed genes in adr-1(-);adr-2(-) double mutant animals compared to wildtype animals. p-value<0.05 were taken as targets misregulated in absence of ad-1 and adr-2. Differentially expressed genes are listed as Wormbase annotations in Column A, Gene name in Column B. The log2fold change values, raw p values and the adjusted p values as obtained from the DESeq2 analysis are shown in columns C, D, and E, respectively. Read counts from the DESeq2 analysis for the three biological replicates of wildtype are listed in columns F, G and H. Read counts from the DESeq2 analysis for the three biological replicates of adr-1(-);adr-2(-) animals are listed in columns I, J and K . "	"Supplemental Table 4: Differentially expressed genes in adr-1(-);adr-2(-) animals grown on OP40 for 7 hours compared to wildtype animals grown on OP50 for 7 hours (Related to Figure 4). DESeq2 was used to identify differentially expressed genes in adr-1(-);adr-2(-) double mutant animals compared to wildtype animals. p-value<0.05 were taken as targets misregulated in absence of ad-1 and adr-2. Differentially expressed genes are listed as Wormbase annotations in Column A, Gene name in Column B. The log2fold change values, raw p values and the adjusted p values as obtained from the DESeq2 analysis are shown in columns C, D, and E, respectively. Read counts from the DESeq2 analysis for the three biological replicates of wildtype are listed in columns F, G and H. Read counts from the DESeq2 analysis for the three biological replicates of adr-1(-);adr-2(-) animals are listed in columns I, J and K . "	"Supplemental Table 4: Differentially expressed genes in adr-1(-);adr-2(-) animals grown on OP40 for 7 hours compared to wildtype animals grown on OP50 for 7 hours (Related to Figure 4). DESeq2 was used to identify differentially expressed genes in adr-1(-);adr-2(-) double mutant animals compared to wildtype animals. p-value<0.05 were taken as targets misregulated in absence of ad-1 and adr-2. Differentially expressed genes are listed as Wormbase annotations in Column A, Gene name in Column B. The log2fold change values, raw p values and the adjusted p values as obtained from the DESeq2 analysis are shown in columns C, D, and E, respectively. Read counts from the DESeq2 analysis for the three biological replicates of wildtype are listed in columns F, G and H. Read counts from the DESeq2 analysis for the three biological replicates of adr-1(-);adr-2(-) animals are listed in columns I, J and K . "	"Supplemental Table 4: Differentially expressed genes in adr-1(-);adr-2(-) animals grown on OP40 for 7 hours compared to wildtype animals grown on OP50 for 7 hours (Related to Figure 4). DESeq2 was used to identify differentially expressed genes in adr-1(-);adr-2(-) double mutant animals compared to wildtype animals. p-value<0.05 were taken as targets misregulated in absence of ad-1 and adr-2. Differentially expressed genes are listed as Wormbase annotations in Column A, Gene name in Column B. The log2fold change values, raw p values and the adjusted p values as obtained from the DESeq2 analysis are shown in columns C, D, and E, respectively. Read counts from the DESeq2 analysis for the three biological replicates of wildtype are listed in columns F, G and H. Read counts from the DESeq2 analysis for the three biological replicates of adr-1(-);adr-2(-) animals are listed in columns I, J and K . "	"Supplemental Table 4: Differentially expressed genes in adr-1(-);adr-2(-) animals grown on OP40 for 7 hours compared to wildtype animals grown on OP50 for 7 hours (Related to Figure 4). DESeq2 was used to identify differentially expressed genes in adr-1(-);adr-2(-) double mutant animals compared to wildtype animals. p-value<0.05 were taken as targets misregulated in absence of ad-1 and adr-2. Differentially expressed genes are listed as Wormbase annotations in Column A, Gene name in Column B. The log2fold change values, raw p values and the adjusted p values as obtained from the DESeq2 analysis are shown in columns C, D, and E, respectively. Read counts from the DESeq2 analysis for the three biological replicates of wildtype are listed in columns F, G and H. Read counts from the DESeq2 analysis for the three biological replicates of adr-1(-);adr-2(-) animals are listed in columns I, J and K . "	"Supplemental Table 4: Differentially expressed genes in adr-1(-);adr-2(-) animals grown on OP40 for 7 hours compared to wildtype animals grown on OP50 for 7 hours (Related to Figure 4). DESeq2 was used to identify differentially expressed genes in adr-1(-);adr-2(-) double mutant animals compared to wildtype animals. p-value<0.05 were taken as targets misregulated in absence of ad-1 and adr-2. Differentially expressed genes are listed as Wormbase annotations in Column A, Gene name in Column B. The log2fold change values, raw p values and the adjusted p values as obtained from the DESeq2 analysis are shown in columns C, D, and E, respectively. Read counts from the DESeq2 analysis for the three biological replicates of wildtype are listed in columns F, G and H. Read counts from the DESeq2 analysis for the three biological replicates of adr-1(-);adr-2(-) animals are listed in columns I, J and K . "	"Supplemental Table 4: Differentially expressed genes in adr-1(-);adr-2(-) animals grown on OP40 for 7 hours compared to wildtype animals grown on OP50 for 7 hours (Related to Figure 4). DESeq2 was used to identify differentially expressed genes in adr-1(-);adr-2(-) double mutant animals compared to wildtype animals. p-value<0.05 were taken as targets misregulated in absence of ad-1 and adr-2. Differentially expressed genes are listed as Wormbase annotations in Column A, Gene name in Column B. The log2fold change values, raw p values and the adjusted p values as obtained from the DESeq2 analysis are shown in columns C, D, and E, respectively. Read counts from the DESeq2 analysis for the three biological replicates of wildtype are listed in columns F, G and H. Read counts from the DESeq2 analysis for the three biological replicates of adr-1(-);adr-2(-) animals are listed in columns I, J and K . "	"Supplemental Table 4: Differentially expressed genes in adr-1(-);adr-2(-) animals grown on OP40 for 7 hours compared to wildtype animals grown on OP50 for 7 hours (Related to Figure 4). DESeq2 was used to identify differentially expressed genes in adr-1(-);adr-2(-) double mutant animals compared to wildtype animals. p-value<0.05 were taken as targets misregulated in absence of ad-1 and adr-2. Differentially expressed genes are listed as Wormbase annotations in Column A, Gene name in Column B. The log2fold change values, raw p values and the adjusted p values as obtained from the DESeq2 analysis are shown in columns C, D, and E, respectively. Read counts from the DESeq2 analysis for the three biological replicates of wildtype are listed in columns F, G and H. Read counts from the DESeq2 analysis for the three biological replicates of adr-1(-);adr-2(-) animals are listed in columns I, J and K . "	"Supplemental Table 4: Differentially expressed genes in adr-1(-);adr-2(-) animals grown on OP40 for 7 hours compared to wildtype animals grown on OP50 for 7 hours (Related to Figure 4). DESeq2 was used to identify differentially expressed genes in adr-1(-);adr-2(-) double mutant animals compared to wildtype animals. p-value<0.05 were taken as targets misregulated in absence of ad-1 and adr-2. Differentially expressed genes are listed as Wormbase annotations in Column A, Gene name in Column B. The log2fold change values, raw p values and the adjusted p values as obtained from the DESeq2 analysis are shown in columns C, D, and E, respectively. Read counts from the DESeq2 analysis for the three biological replicates of wildtype are listed in columns F, G and H. Read counts from the DESeq2 analysis for the three biological replicates of adr-1(-);adr-2(-) animals are listed in columns I, J and K . "	"Supplemental Table 4: Differentially expressed genes in adr-1(-);adr-2(-) animals grown on OP40 for 7 hours compared to wildtype animals grown on OP50 for 7 hours (Related to Figure 4). DESeq2 was used to identify differentially expressed genes in adr-1(-);adr-2(-) double mutant animals compared to wildtype animals. p-value<0.05 were taken as targets misregulated in absence of ad-1 and adr-2. Differentially expressed genes are listed as Wormbase annotations in Column A, Gene name in Column B. The log2fold change values, raw p values and the adjusted p values as obtained from the DESeq2 analysis are shown in columns C, D, and E, respectively. Read counts from the DESeq2 analysis for the three biological replicates of wildtype are listed in columns F, G and H. Read counts from the DESeq2 analysis for the three biological replicates of adr-1(-);adr-2(-) animals are listed in columns I, J and K . "	"Supplemental Table 4: Differentially expressed genes in adr-1(-);adr-2(-) animals grown on OP40 for 7 hours compared to wildtype animals grown on OP50 for 7 hours (Related to Figure 4). DESeq2 was used to identify differentially expressed genes in adr-1(-);adr-2(-) double mutant animals compared to wildtype animals. p-value<0.05 were taken as targets misregulated in absence of ad-1 and adr-2. Differentially expressed genes are listed as Wormbase annotations in Column A, Gene name in Column B. The log2fold change values, raw p values and the adjusted p values as obtained from the DESeq2 analysis are shown in columns C, D, and E, respectively. Read counts from the DESeq2 analysis for the three biological replicates of wildtype are listed in columns F, G and H. Read counts from the DESeq2 analysis for the three biological replicates of adr-1(-);adr-2(-) animals are listed in columns I, J and K . "	"Supplemental Table 4: Differentially expressed genes in adr-1(-);adr-2(-) animals grown on OP40 for 7 hours compared to wildtype animals grown on OP50 for 7 hours (Related to Figure 4). DESeq2 was used to identify differentially expressed genes in adr-1(-);adr-2(-) double mutant animals compared to wildtype animals. p-value<0.05 were taken as targets misregulated in absence of ad-1 and adr-2. Differentially expressed genes are listed as Wormbase annotations in Column A, Gene name in Column B. The log2fold change values, raw p values and the adjusted p values as obtained from the DESeq2 analysis are shown in columns C, D, and E, respectively. Read counts from the DESeq2 analysis for the three biological replicates of wildtype are listed in columns F, G and H. Read counts from the DESeq2 analysis for the three biological replicates of adr-1(-);adr-2(-) animals are listed in columns I, J and K . "	"Supplemental Table 4: Differentially expressed genes in adr-1(-);adr-2(-) animals grown on OP40 for 7 hours compared to wildtype animals grown on OP50 for 7 hours (Related to Figure 4). DESeq2 was used to identify differentially expressed genes in adr-1(-);adr-2(-) double mutant animals compared to wildtype animals. p-value<0.05 were taken as targets misregulated in absence of ad-1 and adr-2. Differentially expressed genes are listed as Wormbase annotations in Column A, Gene name in Column B. The log2fold change values, raw p values and the adjusted p values as obtained from the DESeq2 analysis are shown in columns C, D, and E, respectively. Read counts from the DESeq2 analysis for the three biological replicates of wildtype are listed in columns F, G and H. Read counts from the DESeq2 analysis for the three biological replicates of adr-1(-);adr-2(-) animals are listed in columns I, J and K . "	"Supplemental Table 4: Differentially expressed genes in adr-1(-);adr-2(-) animals grown on OP40 for 7 hours compared to wildtype animals grown on OP50 for 7 hours (Related to Figure 4). DESeq2 was used to identify differentially expressed genes in adr-1(-);adr-2(-) double mutant animals compared to wildtype animals. p-value<0.05 were taken as targets misregulated in absence of ad-1 and adr-2. Differentially expressed genes are listed as Wormbase annotations in Column A, Gene name in Column B. The log2fold change values, raw p values and the adjusted p values as obtained from the DESeq2 analysis are shown in columns C, D, and E, respectively. Read counts from the DESeq2 analysis for the three biological replicates of wildtype are listed in columns F, G and H. Read counts from the DESeq2 analysis for the three biological replicates of adr-1(-);adr-2(-) animals are listed in columns I, J and K . "	"Supplemental Table 4: Differentially expressed genes in adr-1(-);adr-2(-) animals grown on OP40 for 7 hours compared to wildtype animals grown on OP50 for 7 hours (Related to Figure 4). DESeq2 was used to identify differentially expressed genes in adr-1(-);adr-2(-) double mutant animals compared to wildtype animals. p-value<0.05 were taken as targets misregulated in absence of ad-1 and adr-2. Differentially expressed genes are listed as Wormbase annotations in Column A, Gene name in Column B. The log2fold change values, raw p values and the adjusted p values as obtained from the DESeq2 analysis are shown in columns C, D, and E, respectively. Read counts from the DESeq2 analysis for the three biological replicates of wildtype are listed in columns F, G and H. Read counts from the DESeq2 analysis for the three biological replicates of adr-1(-);adr-2(-) animals are listed in columns I, J and K . "	"Supplemental Table 4: Differentially expressed genes in adr-1(-);adr-2(-) animals grown on OP40 for 7 hours compared to wildtype animals grown on OP50 for 7 hours (Related to Figure 4). DESeq2 was used to identify differentially expressed genes in adr-1(-);adr-2(-) double mutant animals compared to wildtype animals. p-value<0.05 were taken as targets misregulated in absence of ad-1 and adr-2. Differentially expressed genes are listed as Wormbase annotations in Column A, Gene name in Column B. The log2fold change values, raw p values and the adjusted p values as obtained from the DESeq2 analysis are shown in columns C, D, and E, respectively. Read counts from the DESeq2 analysis for the three biological replicates of wildtype are listed in columns F, G and H. Read counts from the DESeq2 analysis for the three biological replicates of adr-1(-);adr-2(-) animals are listed in columns I, J and K . "	"Supplemental Table 4: Differentially expressed genes in adr-1(-);adr-2(-) animals grown on OP40 for 7 hours compared to wildtype animals grown on OP50 for 7 hours (Related to Figure 4). DESeq2 was used to identify differentially expressed genes in adr-1(-);adr-2(-) double mutant animals compared to wildtype animals. p-value<0.05 were taken as targets misregulated in absence of ad-1 and adr-2. Differentially expressed genes are listed as Wormbase annotations in Column A, Gene name in Column B. The log2fold change values, raw p values and the adjusted p values as obtained from the DESeq2 analysis are shown in columns C, D, and E, respectively. Read counts from the DESeq2 analysis for the three biological replicates of wildtype are listed in columns F, G and H. Read counts from the DESeq2 analysis for the three biological replicates of adr-1(-);adr-2(-) animals are listed in columns I, J and K . "	"Supplemental Table 4: Differentially expressed genes in adr-1(-);adr-2(-) animals grown on OP40 for 7 hours compared to wildtype animals grown on OP50 for 7 hours (Related to Figure 4). DESeq2 was used to identify differentially expressed genes in adr-1(-);adr-2(-) double mutant animals compared to wildtype animals. p-value<0.05 were taken as targets misregulated in absence of ad-1 and adr-2. Differentially expressed genes are listed as Wormbase annotations in Column A, Gene name in Column B. The log2fold change values, raw p values and the adjusted p values as obtained from the DESeq2 analysis are shown in columns C, D, and E, respectively. Read counts from the DESeq2 analysis for the three biological replicates of wildtype are listed in columns F, G and H. Read counts from the DESeq2 analysis for the three biological replicates of adr-1(-);adr-2(-) animals are listed in columns I, J and K . "	"Supplemental Table 4: Differentially expressed genes in adr-1(-);adr-2(-) animals grown on OP40 for 7 hours compared to wildtype animals grown on OP50 for 7 hours (Related to Figure 4). DESeq2 was used to identify differentially expressed genes in adr-1(-);adr-2(-) double mutant animals compared to wildtype animals. p-value<0.05 were taken as targets misregulated in absence of ad-1 and adr-2. Differentially expressed genes are listed as Wormbase annotations in Column A, Gene name in Column B. The log2fold change values, raw p values and the adjusted p values as obtained from the DESeq2 analysis are shown in columns C, D, and E, respectively. Read counts from the DESeq2 analysis for the three biological replicates of wildtype are listed in columns F, G and H. Read counts from the DESeq2 analysis for the three biological replicates of adr-1(-);adr-2(-) animals are listed in columns I, J and K . "	"Supplemental Table 4: Differentially expressed genes in adr-1(-);adr-2(-) animals grown on OP40 for 7 hours compared to wildtype animals grown on OP50 for 7 hours (Related to Figure 4). DESeq2 was used to identify differentially expressed genes in adr-1(-);adr-2(-) double mutant animals compared to wildtype animals. p-value<0.05 were taken as targets misregulated in absence of ad-1 and adr-2. Differentially expressed genes are listed as Wormbase annotations in Column A, Gene name in Column B. The log2fold change values, raw p values and the adjusted p values as obtained from the DESeq2 analysis are shown in columns C, D, and E, respectively. Read counts from the DESeq2 analysis for the three biological replicates of wildtype are listed in columns F, G and H. Read counts from the DESeq2 analysis for the three biological replicates of adr-1(-);adr-2(-) animals are listed in columns I, J and K . "	"Supplemental Table 4: Differentially expressed genes in adr-1(-);adr-2(-) animals grown on OP40 for 7 hours compared to wildtype animals grown on OP50 for 7 hours (Related to Figure 4). DESeq2 was used to identify differentially expressed genes in adr-1(-);adr-2(-) double mutant animals compared to wildtype animals. p-value<0.05 were taken as targets misregulated in absence of ad-1 and adr-2. Differentially expressed genes are listed as Wormbase annotations in Column A, Gene name in Column B. The log2fold change values, raw p values and the adjusted p values as obtained from the DESeq2 analysis are shown in columns C, D, and E, respectively. Read counts from the DESeq2 analysis for the three biological replicates of wildtype are listed in columns F, G and H. Read counts from the DESeq2 analysis for the three biological replicates of adr-1(-);adr-2(-) animals are listed in columns I, J and K . "	"Supplemental Table 4: Differentially expressed genes in adr-1(-);adr-2(-) animals grown on OP40 for 7 hours compared to wildtype animals grown on OP50 for 7 hours (Related to Figure 4). DESeq2 was used to identify differentially expressed genes in adr-1(-);adr-2(-) double mutant animals compared to wildtype animals. p-value<0.05 were taken as targets misregulated in absence of ad-1 and adr-2. Differentially expressed genes are listed as Wormbase annotations in Column A, Gene name in Column B. The log2fold change values, raw p values and the adjusted p values as obtained from the DESeq2 analysis are shown in columns C, D, and E, respectively. Read counts from the DESeq2 analysis for the three biological replicates of wildtype are listed in columns F, G and H. Read counts from the DESeq2 analysis for the three biological replicates of adr-1(-);adr-2(-) animals are listed in columns I, J and K . "	"Supplemental Table 4: Differentially expressed genes in adr-1(-);adr-2(-) animals grown on OP40 for 7 hours compared to wildtype animals grown on OP50 for 7 hours (Related to Figure 4). DESeq2 was used to identify differentially expressed genes in adr-1(-);adr-2(-) double mutant animals compared to wildtype animals. p-value<0.05 were taken as targets misregulated in absence of ad-1 and adr-2. Differentially expressed genes are listed as Wormbase annotations in Column A, Gene name in Column B. The log2fold change values, raw p values and the adjusted p values as obtained from the DESeq2 analysis are shown in columns C, D, and E, respectively. Read counts from the DESeq2 analysis for the three biological replicates of wildtype are listed in columns F, G and H. Read counts from the DESeq2 analysis for the three biological replicates of adr-1(-);adr-2(-) animals are listed in columns I, J and K . "	"Supplemental Table 4: Differentially expressed genes in adr-1(-);adr-2(-) animals grown on OP40 for 7 hours compared to wildtype animals grown on OP50 for 7 hours (Related to Figure 4). DESeq2 was used to identify differentially expressed genes in adr-1(-);adr-2(-) double mutant animals compared to wildtype animals. p-value<0.05 were taken as targets misregulated in absence of ad-1 and adr-2. Differentially expressed genes are listed as Wormbase annotations in Column A, Gene name in Column B. The log2fold change values, raw p values and the adjusted p values as obtained from the DESeq2 analysis are shown in columns C, D, and E, respectively. Read counts from the DESeq2 analysis for the three biological replicates of wildtype are listed in columns F, G and H. Read counts from the DESeq2 analysis for the three biological replicates of adr-1(-);adr-2(-) animals are listed in columns I, J and K . "	"Supplemental Table 4: Differentially expressed genes in adr-1(-);adr-2(-) animals grown on OP40 for 7 hours compared to wildtype animals grown on OP50 for 7 hours (Related to Figure 4). DESeq2 was used to identify differentially expressed genes in adr-1(-);adr-2(-) double mutant animals compared to wildtype animals. p-value<0.05 were taken as targets misregulated in absence of ad-1 and adr-2. Differentially expressed genes are listed as Wormbase annotations in Column A, Gene name in Column B. The log2fold change values, raw p values and the adjusted p values as obtained from the DESeq2 analysis are shown in columns C, D, and E, respectively. Read counts from the DESeq2 analysis for the three biological replicates of wildtype are listed in columns F, G and H. Read counts from the DESeq2 analysis for the three biological replicates of adr-1(-);adr-2(-) animals are listed in columns I, J and K . "	"Supplemental Table 4: Differentially expressed genes in adr-1(-);adr-2(-) animals grown on OP40 for 7 hours compared to wildtype animals grown on OP50 for 7 hours (Related to Figure 4). DESeq2 was used to identify differentially expressed genes in adr-1(-);adr-2(-) double mutant animals compared to wildtype animals. p-value<0.05 were taken as targets misregulated in absence of ad-1 and adr-2. Differentially expressed genes are listed as Wormbase annotations in Column A, Gene name in Column B. The log2fold change values, raw p values and the adjusted p values as obtained from the DESeq2 analysis are shown in columns C, D, and E, respectively. Read counts from the DESeq2 analysis for the three biological replicates of wildtype are listed in columns F, G and H. Read counts from the DESeq2 analysis for the three biological replicates of adr-1(-);adr-2(-) animals are listed in columns I, J and K . "	"Supplemental Table 4: Differentially expressed genes in adr-1(-);adr-2(-) animals grown on OP40 for 7 hours compared to wildtype animals grown on OP50 for 7 hours (Related to Figure 4). DESeq2 was used to identify differentially expressed genes in adr-1(-);adr-2(-) double mutant animals compared to wildtype animals. p-value<0.05 were taken as targets misregulated in absence of ad-1 and adr-2. Differentially expressed genes are listed as Wormbase annotations in Column A, Gene name in Column B. The log2fold change values, raw p values and the adjusted p values as obtained from the DESeq2 analysis are shown in columns C, D, and E, respectively. Read counts from the DESeq2 analysis for the three biological replicates of wildtype are listed in columns F, G and H. Read counts from the DESeq2 analysis for the three biological replicates of adr-1(-);adr-2(-) animals are listed in columns I, J and K . "	"Supplemental Table 4: Differentially expressed genes in adr-1(-);adr-2(-) animals grown on OP40 for 7 hours compared to wildtype animals grown on OP50 for 7 hours (Related to Figure 4). DESeq2 was used to identify differentially expressed genes in adr-1(-);adr-2(-) double mutant animals compared to wildtype animals. p-value<0.05 were taken as targets misregulated in absence of ad-1 and adr-2. Differentially expressed genes are listed as Wormbase annotations in Column A, Gene name in Column B. The log2fold change values, raw p values and the adjusted p values as obtained from the DESeq2 analysis are shown in columns C, D, and E, respectively. Read counts from the DESeq2 analysis for the three biological replicates of wildtype are listed in columns F, G and H. Read counts from the DESeq2 analysis for the three biological replicates of adr-1(-);adr-2(-) animals are listed in columns I, J and K . "	"Supplemental Table 4: Differentially expressed genes in adr-1(-);adr-2(-) animals grown on OP40 for 7 hours compared to wildtype animals grown on OP50 for 7 hours (Related to Figure 4). DESeq2 was used to identify differentially expressed genes in adr-1(-);adr-2(-) double mutant animals compared to wildtype animals. p-value<0.05 were taken as targets misregulated in absence of ad-1 and adr-2. Differentially expressed genes are listed as Wormbase annotations in Column A, Gene name in Column B. The log2fold change values, raw p values and the adjusted p values as obtained from the DESeq2 analysis are shown in columns C, D, and E, respectively. Read counts from the DESeq2 analysis for the three biological replicates of wildtype are listed in columns F, G and H. Read counts from the DESeq2 analysis for the three biological replicates of adr-1(-);adr-2(-) animals are listed in columns I, J and K . "	"Supplemental Table 4: Differentially expressed genes in adr-1(-);adr-2(-) animals grown on OP40 for 7 hours compared to wildtype animals grown on OP50 for 7 hours (Related to Figure 4). DESeq2 was used to identify differentially expressed genes in adr-1(-);adr-2(-) double mutant animals compared to wildtype animals. p-value<0.05 were taken as targets misregulated in absence of ad-1 and adr-2. Differentially expressed genes are listed as Wormbase annotations in Column A, Gene name in Column B. The log2fold change values, raw p values and the adjusted p values as obtained from the DESeq2 analysis are shown in columns C, D, and E, respectively. Read counts from the DESeq2 analysis for the three biological replicates of wildtype are listed in columns F, G and H. Read counts from the DESeq2 analysis for the three biological replicates of adr-1(-);adr-2(-) animals are listed in columns I, J and K . "	"Supplemental Table 4: Differentially expressed genes in adr-1(-);adr-2(-) animals grown on OP40 for 7 hours compared to wildtype animals grown on OP50 for 7 hours (Related to Figure 4). DESeq2 was used to identify differentially expressed genes in adr-1(-);adr-2(-) double mutant animals compared to wildtype animals. p-value<0.05 were taken as targets misregulated in absence of ad-1 and adr-2. Differentially expressed genes are listed as Wormbase annotations in Column A, Gene name in Column B. The log2fold change values, raw p values and the adjusted p values as obtained from the DESeq2 analysis are shown in columns C, D, and E, respectively. Read counts from the DESeq2 analysis for the three biological replicates of wildtype are listed in columns F, G and H. Read counts from the DESeq2 analysis for the three biological replicates of adr-1(-);adr-2(-) animals are listed in columns I, J and K . "	"Supplemental Table 4: Differentially expressed genes in adr-1(-);adr-2(-) animals grown on OP40 for 7 hours compared to wildtype animals grown on OP50 for 7 hours (Related to Figure 4). DESeq2 was used to identify differentially expressed genes in adr-1(-);adr-2(-) double mutant animals compared to wildtype animals. p-value<0.05 were taken as targets misregulated in absence of ad-1 and adr-2. Differentially expressed genes are listed as Wormbase annotations in Column A, Gene name in Column B. The log2fold change values, raw p values and the adjusted p values as obtained from the DESeq2 analysis are shown in columns C, D, and E, respectively. Read counts from the DESeq2 analysis for the three biological replicates of wildtype are listed in columns F, G and H. Read counts from the DESeq2 analysis for the three biological replicates of adr-1(-);adr-2(-) animals are listed in columns I, J and K . "	"Supplemental Table 4: Differentially expressed genes in adr-1(-);adr-2(-) animals grown on OP40 for 7 hours compared to wildtype animals grown on OP50 for 7 hours (Related to Figure 4). DESeq2 was used to identify differentially expressed genes in adr-1(-);adr-2(-) double mutant animals compared to wildtype animals. p-value<0.05 were taken as targets misregulated in absence of ad-1 and adr-2. Differentially expressed genes are listed as Wormbase annotations in Column A, Gene name in Column B. The log2fold change values, raw p values and the adjusted p values as obtained from the DESeq2 analysis are shown in columns C, D, and E, respectively. Read counts from the DESeq2 analysis for the three biological replicates of wildtype are listed in columns F, G and H. Read counts from the DESeq2 analysis for the three biological replicates of adr-1(-);adr-2(-) animals are listed in columns I, J and K . "	"Supplemental Table 4: Differentially expressed genes in adr-1(-);adr-2(-) animals grown on OP40 for 7 hours compared to wildtype animals grown on OP50 for 7 hours (Related to Figure 4). DESeq2 was used to identify differentially expressed genes in adr-1(-);adr-2(-) double mutant animals compared to wildtype animals. p-value<0.05 were taken as targets misregulated in absence of ad-1 and adr-2. Differentially expressed genes are listed as Wormbase annotations in Column A, Gene name in Column B. The log2fold change values, raw p values and the adjusted p values as obtained from the DESeq2 analysis are shown in columns C, D, and E, respectively. Read counts from the DESeq2 analysis for the three biological replicates of wildtype are listed in columns F, G and H. Read counts from the DESeq2 analysis for the three biological replicates of adr-1(-);adr-2(-) animals are listed in columns I, J and K . "	"Supplemental Table 4: Differentially expressed genes in adr-1(-);adr-2(-) animals grown on OP40 for 7 hours compared to wildtype animals grown on OP50 for 7 hours (Related to Figure 4). DESeq2 was used to identify differentially expressed genes in adr-1(-);adr-2(-) double mutant animals compared to wildtype animals. p-value<0.05 were taken as targets misregulated in absence of ad-1 and adr-2. Differentially expressed genes are listed as Wormbase annotations in Column A, Gene name in Column B. The log2fold change values, raw p values and the adjusted p values as obtained from the DESeq2 analysis are shown in columns C, D, and E, respectively. Read counts from the DESeq2 analysis for the three biological replicates of wildtype are listed in columns F, G and H. Read counts from the DESeq2 analysis for the three biological replicates of adr-1(-);adr-2(-) animals are listed in columns I, J and K . "	"Supplemental Table 4: Differentially expressed genes in adr-1(-);adr-2(-) animals grown on OP40 for 7 hours compared to wildtype animals grown on OP50 for 7 hours (Related to Figure 4). DESeq2 was used to identify differentially expressed genes in adr-1(-);adr-2(-) double mutant animals compared to wildtype animals. p-value<0.05 were taken as targets misregulated in absence of ad-1 and adr-2. Differentially expressed genes are listed as Wormbase annotations in Column A, Gene name in Column B. The log2fold change values, raw p values and the adjusted p values as obtained from the DESeq2 analysis are shown in columns C, D, and E, respectively. Read counts from the DESeq2 analysis for the three biological replicates of wildtype are listed in columns F, G and H. Read counts from the DESeq2 analysis for the three biological replicates of adr-1(-);adr-2(-) animals are listed in columns I, J and K . "	"Supplemental Table 4: Differentially expressed genes in adr-1(-);adr-2(-) animals grown on OP40 for 7 hours compared to wildtype animals grown on OP50 for 7 hours (Related to Figure 4). DESeq2 was used to identify differentially expressed genes in adr-1(-);adr-2(-) double mutant animals compared to wildtype animals. p-value<0.05 were taken as targets misregulated in absence of ad-1 and adr-2. Differentially expressed genes are listed as Wormbase annotations in Column A, Gene name in Column B. The log2fold change values, raw p values and the adjusted p values as obtained from the DESeq2 analysis are shown in columns C, D, and E, respectively. Read counts from the DESeq2 analysis for the three biological replicates of wildtype are listed in columns F, G and H. Read counts from the DESeq2 analysis for the three biological replicates of adr-1(-);adr-2(-) animals are listed in columns I, J and K . "	"Supplemental Table 4: Differentially expressed genes in adr-1(-);adr-2(-) animals grown on OP40 for 7 hours compared to wildtype animals grown on OP50 for 7 hours (Related to Figure 4). DESeq2 was used to identify differentially expressed genes in adr-1(-);adr-2(-) double mutant animals compared to wildtype animals. p-value<0.05 were taken as targets misregulated in absence of ad-1 and adr-2. Differentially expressed genes are listed as Wormbase annotations in Column A, Gene name in Column B. The log2fold change values, raw p values and the adjusted p values as obtained from the DESeq2 analysis are shown in columns C, D, and E, respectively. Read counts from the DESeq2 analysis for the three biological replicates of wildtype are listed in columns F, G and H. Read counts from the DESeq2 analysis for the three biological replicates of adr-1(-);adr-2(-) animals are listed in columns I, J and K . "	"Supplemental Table 4: Differentially expressed genes in adr-1(-);adr-2(-) animals grown on OP40 for 7 hours compared to wildtype animals grown on OP50 for 7 hours (Related to Figure 4). DESeq2 was used to identify differentially expressed genes in adr-1(-);adr-2(-) double mutant animals compared to wildtype animals. p-value<0.05 were taken as targets misregulated in absence of ad-1 and adr-2. Differentially expressed genes are listed as Wormbase annotations in Column A, Gene name in Column B. The log2fold change values, raw p values and the adjusted p values as obtained from the DESeq2 analysis are shown in columns C, D, and E, respectively. Read counts from the DESeq2 analysis for the three biological replicates of wildtype are listed in columns F, G and H. Read counts from the DESeq2 analysis for the three biological replicates of adr-1(-);adr-2(-) animals are listed in columns I, J and K . "	"Supplemental Table 4: Differentially expressed genes in adr-1(-);adr-2(-) animals grown on OP40 for 7 hours compared to wildtype animals grown on OP50 for 7 hours (Related to Figure 4). DESeq2 was used to identify differentially expressed genes in adr-1(-);adr-2(-) double mutant animals compared to wildtype animals. p-value<0.05 were taken as targets misregulated in absence of ad-1 and adr-2. Differentially expressed genes are listed as Wormbase annotations in Column A, Gene name in Column B. The log2fold change values, raw p values and the adjusted p values as obtained from the DESeq2 analysis are shown in columns C, D, and E, respectively. Read counts from the DESeq2 analysis for the three biological replicates of wildtype are listed in columns F, G and H. Read counts from the DESeq2 analysis for the three biological replicates of adr-1(-);adr-2(-) animals are listed in columns I, J and K . "	"Supplemental Table 4: Differentially expressed genes in adr-1(-);adr-2(-) animals grown on OP40 for 7 hours compared to wildtype animals grown on OP50 for 7 hours (Related to Figure 4). DESeq2 was used to identify differentially expressed genes in adr-1(-);adr-2(-) double mutant animals compared to wildtype animals. p-value<0.05 were taken as targets misregulated in absence of ad-1 and adr-2. Differentially expressed genes are listed as Wormbase annotations in Column A, Gene name in Column B. The log2fold change values, raw p values and the adjusted p values as obtained from the DESeq2 analysis are shown in columns C, D, and E, respectively. Read counts from the DESeq2 analysis for the three biological replicates of wildtype are listed in columns F, G and H. Read counts from the DESeq2 analysis for the three biological replicates of adr-1(-);adr-2(-) animals are listed in columns I, J and K . "	"Supplemental Table 4: Differentially expressed genes in adr-1(-);adr-2(-) animals grown on OP40 for 7 hours compared to wildtype animals grown on OP50 for 7 hours (Related to Figure 4). DESeq2 was used to identify differentially expressed genes in adr-1(-);adr-2(-) double mutant animals compared to wildtype animals. p-value<0.05 were taken as targets misregulated in absence of ad-1 and adr-2. Differentially expressed genes are listed as Wormbase annotations in Column A, Gene name in Column B. The log2fold change values, raw p values and the adjusted p values as obtained from the DESeq2 analysis are shown in columns C, D, and E, respectively. Read counts from the DESeq2 analysis for the three biological replicates of wildtype are listed in columns F, G and H. Read counts from the DESeq2 analysis for the three biological replicates of adr-1(-);adr-2(-) animals are listed in columns I, J and K . "	"Supplemental Table 4: Differentially expressed genes in adr-1(-);adr-2(-) animals grown on OP40 for 7 hours compared to wildtype animals grown on OP50 for 7 hours (Related to Figure 4). DESeq2 was used to identify differentially expressed genes in adr-1(-);adr-2(-) double mutant animals compared to wildtype animals. p-value<0.05 were taken as targets misregulated in absence of ad-1 and adr-2. Differentially expressed genes are listed as Wormbase annotations in Column A, Gene name in Column B. The log2fold change values, raw p values and the adjusted p values as obtained from the DESeq2 analysis are shown in columns C, D, and E, respectively. Read counts from the DESeq2 analysis for the three biological replicates of wildtype are listed in columns F, G and H. Read counts from the DESeq2 analysis for the three biological replicates of adr-1(-);adr-2(-) animals are listed in columns I, J and K . "	"Supplemental Table 4: Differentially expressed genes in adr-1(-);adr-2(-) animals grown on OP40 for 7 hours compared to wildtype animals grown on OP50 for 7 hours (Related to Figure 4). DESeq2 was used to identify differentially expressed genes in adr-1(-);adr-2(-) double mutant animals compared to wildtype animals. p-value<0.05 were taken as targets misregulated in absence of ad-1 and adr-2. Differentially expressed genes are listed as Wormbase annotations in Column A, Gene name in Column B. The log2fold change values, raw p values and the adjusted p values as obtained from the DESeq2 analysis are shown in columns C, D, and E, respectively. Read counts from the DESeq2 analysis for the three biological replicates of wildtype are listed in columns F, G and H. Read counts from the DESeq2 analysis for the three biological replicates of adr-1(-);adr-2(-) animals are listed in columns I, J and K . "	"Supplemental Table 4: Differentially expressed genes in adr-1(-);adr-2(-) animals grown on OP40 for 7 hours compared to wildtype animals grown on OP50 for 7 hours (Related to Figure 4). DESeq2 was used to identify differentially expressed genes in adr-1(-);adr-2(-) double mutant animals compared to wildtype animals. p-value<0.05 were taken as targets misregulated in absence of ad-1 and adr-2. Differentially expressed genes are listed as Wormbase annotations in Column A, Gene name in Column B. The log2fold change values, raw p values and the adjusted p values as obtained from the DESeq2 analysis are shown in columns C, D, and E, respectively. Read counts from the DESeq2 analysis for the three biological replicates of wildtype are listed in columns F, G and H. Read counts from the DESeq2 analysis for the three biological replicates of adr-1(-);adr-2(-) animals are listed in columns I, J and K . "	"Supplemental Table 4: Differentially expressed genes in adr-1(-);adr-2(-) animals grown on OP40 for 7 hours compared to wildtype animals grown on OP50 for 7 hours (Related to Figure 4). DESeq2 was used to identify differentially expressed genes in adr-1(-);adr-2(-) double mutant animals compared to wildtype animals. p-value<0.05 were taken as targets misregulated in absence of ad-1 and adr-2. Differentially expressed genes are listed as Wormbase annotations in Column A, Gene name in Column B. The log2fold change values, raw p values and the adjusted p values as obtained from the DESeq2 analysis are shown in columns C, D, and E, respectively. Read counts from the DESeq2 analysis for the three biological replicates of wildtype are listed in columns F, G and H. Read counts from the DESeq2 analysis for the three biological replicates of adr-1(-);adr-2(-) animals are listed in columns I, J and K . "	"Supplemental Table 4: Differentially expressed genes in adr-1(-);adr-2(-) animals grown on OP40 for 7 hours compared to wildtype animals grown on OP50 for 7 hours (Related to Figure 4). DESeq2 was used to identify differentially expressed genes in adr-1(-);adr-2(-) double mutant animals compared to wildtype animals. p-value<0.05 were taken as targets misregulated in absence of ad-1 and adr-2. Differentially expressed genes are listed as Wormbase annotations in Column A, Gene name in Column B. The log2fold change values, raw p values and the adjusted p values as obtained from the DESeq2 analysis are shown in columns C, D, and E, respectively. Read counts from the DESeq2 analysis for the three biological replicates of wildtype are listed in columns F, G and H. Read counts from the DESeq2 analysis for the three biological replicates of adr-1(-);adr-2(-) animals are listed in columns I, J and K . "	"Supplemental Table 4: Differentially expressed genes in adr-1(-);adr-2(-) animals grown on OP40 for 7 hours compared to wildtype animals grown on OP50 for 7 hours (Related to Figure 4). DESeq2 was used to identify differentially expressed genes in adr-1(-);adr-2(-) double mutant animals compared to wildtype animals. p-value<0.05 were taken as targets misregulated in absence of ad-1 and adr-2. Differentially expressed genes are listed as Wormbase annotations in Column A, Gene name in Column B. The log2fold change values, raw p values and the adjusted p values as obtained from the DESeq2 analysis are shown in columns C, D, and E, respectively. Read counts from the DESeq2 analysis for the three biological replicates of wildtype are listed in columns F, G and H. Read counts from the DESeq2 analysis for the three biological replicates of adr-1(-);adr-2(-) animals are listed in columns I, J and K . "	"Supplemental Table 4: Differentially expressed genes in adr-1(-);adr-2(-) animals grown on OP40 for 7 hours compared to wildtype animals grown on OP50 for 7 hours (Related to Figure 4). DESeq2 was used to identify differentially expressed genes in adr-1(-);adr-2(-) double mutant animals compared to wildtype animals. p-value<0.05 were taken as targets misregulated in absence of ad-1 and adr-2. Differentially expressed genes are listed as Wormbase annotations in Column A, Gene name in Column B. The log2fold change values, raw p values and the adjusted p values as obtained from the DESeq2 analysis are shown in columns C, D, and E, respectively. Read counts from the DESeq2 analysis for the three biological replicates of wildtype are listed in columns F, G and H. Read counts from the DESeq2 analysis for the three biological replicates of adr-1(-);adr-2(-) animals are listed in columns I, J and K . "	"Supplemental Table 4: Differentially expressed genes in adr-1(-);adr-2(-) animals grown on OP40 for 7 hours compared to wildtype animals grown on OP50 for 7 hours (Related to Figure 4). DESeq2 was used to identify differentially expressed genes in adr-1(-);adr-2(-) double mutant animals compared to wildtype animals. p-value<0.05 were taken as targets misregulated in absence of ad-1 and adr-2. Differentially expressed genes are listed as Wormbase annotations in Column A, Gene name in Column B. The log2fold change values, raw p values and the adjusted p values as obtained from the DESeq2 analysis are shown in columns C, D, and E, respectively. Read counts from the DESeq2 analysis for the three biological replicates of wildtype are listed in columns F, G and H. Read counts from the DESeq2 analysis for the three biological replicates of adr-1(-);adr-2(-) animals are listed in columns I, J and K . "	"Supplemental Table 4: Differentially expressed genes in adr-1(-);adr-2(-) animals grown on OP40 for 7 hours compared to wildtype animals grown on OP50 for 7 hours (Related to Figure 4). DESeq2 was used to identify differentially expressed genes in adr-1(-);adr-2(-) double mutant animals compared to wildtype animals. p-value<0.05 were taken as targets misregulated in absence of ad-1 and adr-2. Differentially expressed genes are listed as Wormbase annotations in Column A, Gene name in Column B. The log2fold change values, raw p values and the adjusted p values as obtained from the DESeq2 analysis are shown in columns C, D, and E, respectively. Read counts from the DESeq2 analysis for the three biological replicates of wildtype are listed in columns F, G and H. Read counts from the DESeq2 analysis for the three biological replicates of adr-1(-);adr-2(-) animals are listed in columns I, J and K . "	"Supplemental Table 4: Differentially expressed genes in adr-1(-);adr-2(-) animals grown on OP40 for 7 hours compared to wildtype animals grown on OP50 for 7 hours (Related to Figure 4). DESeq2 was used to identify differentially expressed genes in adr-1(-);adr-2(-) double mutant animals compared to wildtype animals. p-value<0.05 were taken as targets misregulated in absence of ad-1 and adr-2. Differentially expressed genes are listed as Wormbase annotations in Column A, Gene name in Column B. The log2fold change values, raw p values and the adjusted p values as obtained from the DESeq2 analysis are shown in columns C, D, and E, respectively. Read counts from the DESeq2 analysis for the three biological replicates of wildtype are listed in columns F, G and H. Read counts from the DESeq2 analysis for the three biological replicates of adr-1(-);adr-2(-) animals are listed in columns I, J and K . "	"Supplemental Table 4: Differentially expressed genes in adr-1(-);adr-2(-) animals grown on OP40 for 7 hours compared to wildtype animals grown on OP50 for 7 hours (Related to Figure 4). DESeq2 was used to identify differentially expressed genes in adr-1(-);adr-2(-) double mutant animals compared to wildtype animals. p-value<0.05 were taken as targets misregulated in absence of ad-1 and adr-2. Differentially expressed genes are listed as Wormbase annotations in Column A, Gene name in Column B. The log2fold change values, raw p values and the adjusted p values as obtained from the DESeq2 analysis are shown in columns C, D, and E, respectively. Read counts from the DESeq2 analysis for the three biological replicates of wildtype are listed in columns F, G and H. Read counts from the DESeq2 analysis for the three biological replicates of adr-1(-);adr-2(-) animals are listed in columns I, J and K . "	"Supplemental Table 4: Differentially expressed genes in adr-1(-);adr-2(-) animals grown on OP40 for 7 hours compared to wildtype animals grown on OP50 for 7 hours (Related to Figure 4). DESeq2 was used to identify differentially expressed genes in adr-1(-);adr-2(-) double mutant animals compared to wildtype animals. p-value<0.05 were taken as targets misregulated in absence of ad-1 and adr-2. Differentially expressed genes are listed as Wormbase annotations in Column A, Gene name in Column B. The log2fold change values, raw p values and the adjusted p values as obtained from the DESeq2 analysis are shown in columns C, D, and E, respectively. Read counts from the DESeq2 analysis for the three biological replicates of wildtype are listed in columns F, G and H. Read counts from the DESeq2 analysis for the three biological replicates of adr-1(-);adr-2(-) animals are listed in columns I, J and K . "	"Supplemental Table 4: Differentially expressed genes in adr-1(-);adr-2(-) animals grown on OP40 for 7 hours compared to wildtype animals grown on OP50 for 7 hours (Related to Figure 4). DESeq2 was used to identify differentially expressed genes in adr-1(-);adr-2(-) double mutant animals compared to wildtype animals. p-value<0.05 were taken as targets misregulated in absence of ad-1 and adr-2. Differentially expressed genes are listed as Wormbase annotations in Column A, Gene name in Column B. The log2fold change values, raw p values and the adjusted p values as obtained from the DESeq2 analysis are shown in columns C, D, and E, respectively. Read counts from the DESeq2 analysis for the three biological replicates of wildtype are listed in columns F, G and H. Read counts from the DESeq2 analysis for the three biological replicates of adr-1(-);adr-2(-) animals are listed in columns I, J and K . "	"Supplemental Table 4: Differentially expressed genes in adr-1(-);adr-2(-) animals grown on OP40 for 7 hours compared to wildtype animals grown on OP50 for 7 hours (Related to Figure 4). DESeq2 was used to identify differentially expressed genes in adr-1(-);adr-2(-) double mutant animals compared to wildtype animals. p-value<0.05 were taken as targets misregulated in absence of ad-1 and adr-2. Differentially expressed genes are listed as Wormbase annotations in Column A, Gene name in Column B. The log2fold change values, raw p values and the adjusted p values as obtained from the DESeq2 analysis are shown in columns C, D, and E, respectively. Read counts from the DESeq2 analysis for the three biological replicates of wildtype are listed in columns F, G and H. Read counts from the DESeq2 analysis for the three biological replicates of adr-1(-);adr-2(-) animals are listed in columns I, J and K . "	"Supplemental Table 4: Differentially expressed genes in adr-1(-);adr-2(-) animals grown on OP40 for 7 hours compared to wildtype animals grown on OP50 for 7 hours (Related to Figure 4). DESeq2 was used to identify differentially expressed genes in adr-1(-);adr-2(-) double mutant animals compared to wildtype animals. p-value<0.05 were taken as targets misregulated in absence of ad-1 and adr-2. Differentially expressed genes are listed as Wormbase annotations in Column A, Gene name in Column B. The log2fold change values, raw p values and the adjusted p values as obtained from the DESeq2 analysis are shown in columns C, D, and E, respectively. Read counts from the DESeq2 analysis for the three biological replicates of wildtype are listed in columns F, G and H. Read counts from the DESeq2 analysis for the three biological replicates of adr-1(-);adr-2(-) animals are listed in columns I, J and K . "	"Supplemental Table 4: Differentially expressed genes in adr-1(-);adr-2(-) animals grown on OP40 for 7 hours compared to wildtype animals grown on OP50 for 7 hours (Related to Figure 4). DESeq2 was used to identify differentially expressed genes in adr-1(-);adr-2(-) double mutant animals compared to wildtype animals. p-value<0.05 were taken as targets misregulated in absence of ad-1 and adr-2. Differentially expressed genes are listed as Wormbase annotations in Column A, Gene name in Column B. The log2fold change values, raw p values and the adjusted p values as obtained from the DESeq2 analysis are shown in columns C, D, and E, respectively. Read counts from the DESeq2 analysis for the three biological replicates of wildtype are listed in columns F, G and H. Read counts from the DESeq2 analysis for the three biological replicates of adr-1(-);adr-2(-) animals are listed in columns I, J and K . "	"Supplemental Table 4: Differentially expressed genes in adr-1(-);adr-2(-) animals grown on OP40 for 7 hours compared to wildtype animals grown on OP50 for 7 hours (Related to Figure 4). DESeq2 was used to identify differentially expressed genes in adr-1(-);adr-2(-) double mutant animals compared to wildtype animals. p-value<0.05 were taken as targets misregulated in absence of ad-1 and adr-2. Differentially expressed genes are listed as Wormbase annotations in Column A, Gene name in Column B. The log2fold change values, raw p values and the adjusted p values as obtained from the DESeq2 analysis are shown in columns C, D, and E, respectively. Read counts from the DESeq2 analysis for the three biological replicates of wildtype are listed in columns F, G and H. Read counts from the DESeq2 analysis for the three biological replicates of adr-1(-);adr-2(-) animals are listed in columns I, J and K . "	"Supplemental Table 4: Differentially expressed genes in adr-1(-);adr-2(-) animals grown on OP40 for 7 hours compared to wildtype animals grown on OP50 for 7 hours (Related to Figure 4). DESeq2 was used to identify differentially expressed genes in adr-1(-);adr-2(-) double mutant animals compared to wildtype animals. p-value<0.05 were taken as targets misregulated in absence of ad-1 and adr-2. Differentially expressed genes are listed as Wormbase annotations in Column A, Gene name in Column B. The log2fold change values, raw p values and the adjusted p values as obtained from the DESeq2 analysis are shown in columns C, D, and E, respectively. Read counts from the DESeq2 analysis for the three biological replicates of wildtype are listed in columns F, G and H. Read counts from the DESeq2 analysis for the three biological replicates of adr-1(-);adr-2(-) animals are listed in columns I, J and K . "	"Supplemental Table 4: Differentially expressed genes in adr-1(-);adr-2(-) animals grown on OP40 for 7 hours compared to wildtype animals grown on OP50 for 7 hours (Related to Figure 4). DESeq2 was used to identify differentially expressed genes in adr-1(-);adr-2(-) double mutant animals compared to wildtype animals. p-value<0.05 were taken as targets misregulated in absence of ad-1 and adr-2. Differentially expressed genes are listed as Wormbase annotations in Column A, Gene name in Column B. The log2fold change values, raw p values and the adjusted p values as obtained from the DESeq2 analysis are shown in columns C, D, and E, respectively. Read counts from the DESeq2 analysis for the three biological replicates of wildtype are listed in columns F, G and H. Read counts from the DESeq2 analysis for the three biological replicates of adr-1(-);adr-2(-) animals are listed in columns I, J and K . "	"Supplemental Table 4: Differentially expressed genes in adr-1(-);adr-2(-) animals grown on OP40 for 7 hours compared to wildtype animals grown on OP50 for 7 hours (Related to Figure 4). DESeq2 was used to identify differentially expressed genes in adr-1(-);adr-2(-) double mutant animals compared to wildtype animals. p-value<0.05 were taken as targets misregulated in absence of ad-1 and adr-2. Differentially expressed genes are listed as Wormbase annotations in Column A, Gene name in Column B. The log2fold change values, raw p values and the adjusted p values as obtained from the DESeq2 analysis are shown in columns C, D, and E, respectively. Read counts from the DESeq2 analysis for the three biological replicates of wildtype are listed in columns F, G and H. Read counts from the DESeq2 analysis for the three biological replicates of adr-1(-);adr-2(-) animals are listed in columns I, J and K . "	"Supplemental Table 4: Differentially expressed genes in adr-1(-);adr-2(-) animals grown on OP40 for 7 hours compared to wildtype animals grown on OP50 for 7 hours (Related to Figure 4). DESeq2 was used to identify differentially expressed genes in adr-1(-);adr-2(-) double mutant animals compared to wildtype animals. p-value<0.05 were taken as targets misregulated in absence of ad-1 and adr-2. Differentially expressed genes are listed as Wormbase annotations in Column A, Gene name in Column B. The log2fold change values, raw p values and the adjusted p values as obtained from the DESeq2 analysis are shown in columns C, D, and E, respectively. Read counts from the DESeq2 analysis for the three biological replicates of wildtype are listed in columns F, G and H. Read counts from the DESeq2 analysis for the three biological replicates of adr-1(-);adr-2(-) animals are listed in columns I, J and K . "	"Supplemental Table 4: Differentially expressed genes in adr-1(-);adr-2(-) animals grown on OP40 for 7 hours compared to wildtype animals grown on OP50 for 7 hours (Related to Figure 4). DESeq2 was used to identify differentially expressed genes in adr-1(-);adr-2(-) double mutant animals compared to wildtype animals. p-value<0.05 were taken as targets misregulated in absence of ad-1 and adr-2. Differentially expressed genes are listed as Wormbase annotations in Column A, Gene name in Column B. The log2fold change values, raw p values and the adjusted p values as obtained from the DESeq2 analysis are shown in columns C, D, and E, respectively. Read counts from the DESeq2 analysis for the three biological replicates of wildtype are listed in columns F, G and H. Read counts from the DESeq2 analysis for the three biological replicates of adr-1(-);adr-2(-) animals are listed in columns I, J and K . "	"Supplemental Table 4: Differentially expressed genes in adr-1(-);adr-2(-) animals grown on OP40 for 7 hours compared to wildtype animals grown on OP50 for 7 hours (Related to Figure 4). DESeq2 was used to identify differentially expressed genes in adr-1(-);adr-2(-) double mutant animals compared to wildtype animals. p-value<0.05 were taken as targets misregulated in absence of ad-1 and adr-2. Differentially expressed genes are listed as Wormbase annotations in Column A, Gene name in Column B. The log2fold change values, raw p values and the adjusted p values as obtained from the DESeq2 analysis are shown in columns C, D, and E, respectively. Read counts from the DESeq2 analysis for the three biological replicates of wildtype are listed in columns F, G and H. Read counts from the DESeq2 analysis for the three biological replicates of adr-1(-);adr-2(-) animals are listed in columns I, J and K . "	"Supplemental Table 4: Differentially expressed genes in adr-1(-);adr-2(-) animals grown on OP40 for 7 hours compared to wildtype animals grown on OP50 for 7 hours (Related to Figure 4). DESeq2 was used to identify differentially expressed genes in adr-1(-);adr-2(-) double mutant animals compared to wildtype animals. p-value<0.05 were taken as targets misregulated in absence of ad-1 and adr-2. Differentially expressed genes are listed as Wormbase annotations in Column A, Gene name in Column B. The log2fold change values, raw p values and the adjusted p values as obtained from the DESeq2 analysis are shown in columns C, D, and E, respectively. Read counts from the DESeq2 analysis for the three biological replicates of wildtype are listed in columns F, G and H. Read counts from the DESeq2 analysis for the three biological replicates of adr-1(-);adr-2(-) animals are listed in columns I, J and K . "	"Supplemental Table 4: Differentially expressed genes in adr-1(-);adr-2(-) animals grown on OP40 for 7 hours compared to wildtype animals grown on OP50 for 7 hours (Related to Figure 4). DESeq2 was used to identify differentially expressed genes in adr-1(-);adr-2(-) double mutant animals compared to wildtype animals. p-value<0.05 were taken as targets misregulated in absence of ad-1 and adr-2. Differentially expressed genes are listed as Wormbase annotations in Column A, Gene name in Column B. The log2fold change values, raw p values and the adjusted p values as obtained from the DESeq2 analysis are shown in columns C, D, and E, respectively. Read counts from the DESeq2 analysis for the three biological replicates of wildtype are listed in columns F, G and H. Read counts from the DESeq2 analysis for the three biological replicates of adr-1(-);adr-2(-) animals are listed in columns I, J and K . "	"Supplemental Table 4: Differentially expressed genes in adr-1(-);adr-2(-) animals grown on OP40 for 7 hours compared to wildtype animals grown on OP50 for 7 hours (Related to Figure 4). DESeq2 was used to identify differentially expressed genes in adr-1(-);adr-2(-) double mutant animals compared to wildtype animals. p-value<0.05 were taken as targets misregulated in absence of ad-1 and adr-2. Differentially expressed genes are listed as Wormbase annotations in Column A, Gene name in Column B. The log2fold change values, raw p values and the adjusted p values as obtained from the DESeq2 analysis are shown in columns C, D, and E, respectively. Read counts from the DESeq2 analysis for the three biological replicates of wildtype are listed in columns F, G and H. Read counts from the DESeq2 analysis for the three biological replicates of adr-1(-);adr-2(-) animals are listed in columns I, J and K . "	"Supplemental Table 4: Differentially expressed genes in adr-1(-);adr-2(-) animals grown on OP40 for 7 hours compared to wildtype animals grown on OP50 for 7 hours (Related to Figure 4). DESeq2 was used to identify differentially expressed genes in adr-1(-);adr-2(-) double mutant animals compared to wildtype animals. p-value<0.05 were taken as targets misregulated in absence of ad-1 and adr-2. Differentially expressed genes are listed as Wormbase annotations in Column A, Gene name in Column B. The log2fold change values, raw p values and the adjusted p values as obtained from the DESeq2 analysis are shown in columns C, D, and E, respectively. Read counts from the DESeq2 analysis for the three biological replicates of wildtype are listed in columns F, G and H. Read counts from the DESeq2 analysis for the three biological replicates of adr-1(-);adr-2(-) animals are listed in columns I, J and K . "	"Supplemental Table 4: Differentially expressed genes in adr-1(-);adr-2(-) animals grown on OP40 for 7 hours compared to wildtype animals grown on OP50 for 7 hours (Related to Figure 4). DESeq2 was used to identify differentially expressed genes in adr-1(-);adr-2(-) double mutant animals compared to wildtype animals. p-value<0.05 were taken as targets misregulated in absence of ad-1 and adr-2. Differentially expressed genes are listed as Wormbase annotations in Column A, Gene name in Column B. The log2fold change values, raw p values and the adjusted p values as obtained from the DESeq2 analysis are shown in columns C, D, and E, respectively. Read counts from the DESeq2 analysis for the three biological replicates of wildtype are listed in columns F, G and H. Read counts from the DESeq2 analysis for the three biological replicates of adr-1(-);adr-2(-) animals are listed in columns I, J and K . "	"Supplemental Table 4: Differentially expressed genes in adr-1(-);adr-2(-) animals grown on OP40 for 7 hours compared to wildtype animals grown on OP50 for 7 hours (Related to Figure 4). DESeq2 was used to identify differentially expressed genes in adr-1(-);adr-2(-) double mutant animals compared to wildtype animals. p-value<0.05 were taken as targets misregulated in absence of ad-1 and adr-2. Differentially expressed genes are listed as Wormbase annotations in Column A, Gene name in Column B. The log2fold change values, raw p values and the adjusted p values as obtained from the DESeq2 analysis are shown in columns C, D, and E, respectively. Read counts from the DESeq2 analysis for the three biological replicates of wildtype are listed in columns F, G and H. Read counts from the DESeq2 analysis for the three biological replicates of adr-1(-);adr-2(-) animals are listed in columns I, J and K . "	"Supplemental Table 4: Differentially expressed genes in adr-1(-);adr-2(-) animals grown on OP40 for 7 hours compared to wildtype animals grown on OP50 for 7 hours (Related to Figure 4). DESeq2 was used to identify differentially expressed genes in adr-1(-);adr-2(-) double mutant animals compared to wildtype animals. p-value<0.05 were taken as targets misregulated in absence of ad-1 and adr-2. Differentially expressed genes are listed as Wormbase annotations in Column A, Gene name in Column B. The log2fold change values, raw p values and the adjusted p values as obtained from the DESeq2 analysis are shown in columns C, D, and E, respectively. Read counts from the DESeq2 analysis for the three biological replicates of wildtype are listed in columns F, G and H. Read counts from the DESeq2 analysis for the three biological replicates of adr-1(-);adr-2(-) animals are listed in columns I, J and K . "	"Supplemental Table 4: Differentially expressed genes in adr-1(-);adr-2(-) animals grown on OP40 for 7 hours compared to wildtype animals grown on OP50 for 7 hours (Related to Figure 4). DESeq2 was used to identify differentially expressed genes in adr-1(-);adr-2(-) double mutant animals compared to wildtype animals. p-value<0.05 were taken as targets misregulated in absence of ad-1 and adr-2. Differentially expressed genes are listed as Wormbase annotations in Column A, Gene name in Column B. The log2fold change values, raw p values and the adjusted p values as obtained from the DESeq2 analysis are shown in columns C, D, and E, respectively. Read counts from the DESeq2 analysis for the three biological replicates of wildtype are listed in columns F, G and H. Read counts from the DESeq2 analysis for the three biological replicates of adr-1(-);adr-2(-) animals are listed in columns I, J and K . "	"Supplemental Table 4: Differentially expressed genes in adr-1(-);adr-2(-) animals grown on OP40 for 7 hours compared to wildtype animals grown on OP50 for 7 hours (Related to Figure 4). DESeq2 was used to identify differentially expressed genes in adr-1(-);adr-2(-) double mutant animals compared to wildtype animals. p-value<0.05 were taken as targets misregulated in absence of ad-1 and adr-2. Differentially expressed genes are listed as Wormbase annotations in Column A, Gene name in Column B. The log2fold change values, raw p values and the adjusted p values as obtained from the DESeq2 analysis are shown in columns C, D, and E, respectively. Read counts from the DESeq2 analysis for the three biological replicates of wildtype are listed in columns F, G and H. Read counts from the DESeq2 analysis for the three biological replicates of adr-1(-);adr-2(-) animals are listed in columns I, J and K . "	"Supplemental Table 4: Differentially expressed genes in adr-1(-);adr-2(-) animals grown on OP40 for 7 hours compared to wildtype animals grown on OP50 for 7 hours (Related to Figure 4). DESeq2 was used to identify differentially expressed genes in adr-1(-);adr-2(-) double mutant animals compared to wildtype animals. p-value<0.05 were taken as targets misregulated in absence of ad-1 and adr-2. Differentially expressed genes are listed as Wormbase annotations in Column A, Gene name in Column B. The log2fold change values, raw p values and the adjusted p values as obtained from the DESeq2 analysis are shown in columns C, D, and E, respectively. Read counts from the DESeq2 analysis for the three biological replicates of wildtype are listed in columns F, G and H. Read counts from the DESeq2 analysis for the three biological replicates of adr-1(-);adr-2(-) animals are listed in columns I, J and K . "	"Supplemental Table 4: Differentially expressed genes in adr-1(-);adr-2(-) animals grown on OP40 for 7 hours compared to wildtype animals grown on OP50 for 7 hours (Related to Figure 4). DESeq2 was used to identify differentially expressed genes in adr-1(-);adr-2(-) double mutant animals compared to wildtype animals. p-value<0.05 were taken as targets misregulated in absence of ad-1 and adr-2. Differentially expressed genes are listed as Wormbase annotations in Column A, Gene name in Column B. The log2fold change values, raw p values and the adjusted p values as obtained from the DESeq2 analysis are shown in columns C, D, and E, respectively. Read counts from the DESeq2 analysis for the three biological replicates of wildtype are listed in columns F, G and H. Read counts from the DESeq2 analysis for the three biological replicates of adr-1(-);adr-2(-) animals are listed in columns I, J and K . "	"Supplemental Table 4: Differentially expressed genes in adr-1(-);adr-2(-) animals grown on OP40 for 7 hours compared to wildtype animals grown on OP50 for 7 hours (Related to Figure 4). DESeq2 was used to identify differentially expressed genes in adr-1(-);adr-2(-) double mutant animals compared to wildtype animals. p-value<0.05 were taken as targets misregulated in absence of ad-1 and adr-2. Differentially expressed genes are listed as Wormbase annotations in Column A, Gene name in Column B. The log2fold change values, raw p values and the adjusted p values as obtained from the DESeq2 analysis are shown in columns C, D, and E, respectively. Read counts from the DESeq2 analysis for the three biological replicates of wildtype are listed in columns F, G and H. Read counts from the DESeq2 analysis for the three biological replicates of adr-1(-);adr-2(-) animals are listed in columns I, J and K . "	"Supplemental Table 4: Differentially expressed genes in adr-1(-);adr-2(-) animals grown on OP40 for 7 hours compared to wildtype animals grown on OP50 for 7 hours (Related to Figure 4). DESeq2 was used to identify differentially expressed genes in adr-1(-);adr-2(-) double mutant animals compared to wildtype animals. p-value<0.05 were taken as targets misregulated in absence of ad-1 and adr-2. Differentially expressed genes are listed as Wormbase annotations in Column A, Gene name in Column B. The log2fold change values, raw p values and the adjusted p values as obtained from the DESeq2 analysis are shown in columns C, D, and E, respectively. Read counts from the DESeq2 analysis for the three biological replicates of wildtype are listed in columns F, G and H. Read counts from the DESeq2 analysis for the three biological replicates of adr-1(-);adr-2(-) animals are listed in columns I, J and K . "	"Supplemental Table 4: Differentially expressed genes in adr-1(-);adr-2(-) animals grown on OP40 for 7 hours compared to wildtype animals grown on OP50 for 7 hours (Related to Figure 4). DESeq2 was used to identify differentially expressed genes in adr-1(-);adr-2(-) double mutant animals compared to wildtype animals. p-value<0.05 were taken as targets misregulated in absence of ad-1 and adr-2. Differentially expressed genes are listed as Wormbase annotations in Column A, Gene name in Column B. The log2fold change values, raw p values and the adjusted p values as obtained from the DESeq2 analysis are shown in columns C, D, and E, respectively. Read counts from the DESeq2 analysis for the three biological replicates of wildtype are listed in columns F, G and H. Read counts from the DESeq2 analysis for the three biological replicates of adr-1(-);adr-2(-) animals are listed in columns I, J and K . "	"Supplemental Table 4: Differentially expressed genes in adr-1(-);adr-2(-) animals grown on OP40 for 7 hours compared to wildtype animals grown on OP50 for 7 hours (Related to Figure 4). DESeq2 was used to identify differentially expressed genes in adr-1(-);adr-2(-) double mutant animals compared to wildtype animals. p-value<0.05 were taken as targets misregulated in absence of ad-1 and adr-2. Differentially expressed genes are listed as Wormbase annotations in Column A, Gene name in Column B. The log2fold change values, raw p values and the adjusted p values as obtained from the DESeq2 analysis are shown in columns C, D, and E, respectively. Read counts from the DESeq2 analysis for the three biological replicates of wildtype are listed in columns F, G and H. Read counts from the DESeq2 analysis for the three biological replicates of adr-1(-);adr-2(-) animals are listed in columns I, J and K . "	"Supplemental Table 4: Differentially expressed genes in adr-1(-);adr-2(-) animals grown on OP40 for 7 hours compared to wildtype animals grown on OP50 for 7 hours (Related to Figure 4). DESeq2 was used to identify differentially expressed genes in adr-1(-);adr-2(-) double mutant animals compared to wildtype animals. p-value<0.05 were taken as targets misregulated in absence of ad-1 and adr-2. Differentially expressed genes are listed as Wormbase annotations in Column A, Gene name in Column B. The log2fold change values, raw p values and the adjusted p values as obtained from the DESeq2 analysis are shown in columns C, D, and E, respectively. Read counts from the DESeq2 analysis for the three biological replicates of wildtype are listed in columns F, G and H. Read counts from the DESeq2 analysis for the three biological replicates of adr-1(-);adr-2(-) animals are listed in columns I, J and K . "	"Supplemental Table 4: Differentially expressed genes in adr-1(-);adr-2(-) animals grown on OP40 for 7 hours compared to wildtype animals grown on OP50 for 7 hours (Related to Figure 4). DESeq2 was used to identify differentially expressed genes in adr-1(-);adr-2(-) double mutant animals compared to wildtype animals. p-value<0.05 were taken as targets misregulated in absence of ad-1 and adr-2. Differentially expressed genes are listed as Wormbase annotations in Column A, Gene name in Column B. The log2fold change values, raw p values and the adjusted p values as obtained from the DESeq2 analysis are shown in columns C, D, and E, respectively. Read counts from the DESeq2 analysis for the three biological replicates of wildtype are listed in columns F, G and H. Read counts from the DESeq2 analysis for the three biological replicates of adr-1(-);adr-2(-) animals are listed in columns I, J and K . "	"Supplemental Table 4: Differentially expressed genes in adr-1(-);adr-2(-) animals grown on OP40 for 7 hours compared to wildtype animals grown on OP50 for 7 hours (Related to Figure 4). DESeq2 was used to identify differentially expressed genes in adr-1(-);adr-2(-) double mutant animals compared to wildtype animals. p-value<0.05 were taken as targets misregulated in absence of ad-1 and adr-2. Differentially expressed genes are listed as Wormbase annotations in Column A, Gene name in Column B. The log2fold change values, raw p values and the adjusted p values as obtained from the DESeq2 analysis are shown in columns C, D, and E, respectively. Read counts from the DESeq2 analysis for the three biological replicates of wildtype are listed in columns F, G and H. Read counts from the DESeq2 analysis for the three biological replicates of adr-1(-);adr-2(-) animals are listed in columns I, J and K . "	"Supplemental Table 4: Differentially expressed genes in adr-1(-);adr-2(-) animals grown on OP40 for 7 hours compared to wildtype animals grown on OP50 for 7 hours (Related to Figure 4). DESeq2 was used to identify differentially expressed genes in adr-1(-);adr-2(-) double mutant animals compared to wildtype animals. p-value<0.05 were taken as targets misregulated in absence of ad-1 and adr-2. Differentially expressed genes are listed as Wormbase annotations in Column A, Gene name in Column B. The log2fold change values, raw p values and the adjusted p values as obtained from the DESeq2 analysis are shown in columns C, D, and E, respectively. Read counts from the DESeq2 analysis for the three biological replicates of wildtype are listed in columns F, G and H. Read counts from the DESeq2 analysis for the three biological replicates of adr-1(-);adr-2(-) animals are listed in columns I, J and K . "	"Supplemental Table 4: Differentially expressed genes in adr-1(-);adr-2(-) animals grown on OP40 for 7 hours compared to wildtype animals grown on OP50 for 7 hours (Related to Figure 4). DESeq2 was used to identify differentially expressed genes in adr-1(-);adr-2(-) double mutant animals compared to wildtype animals. p-value<0.05 were taken as targets misregulated in absence of ad-1 and adr-2. Differentially expressed genes are listed as Wormbase annotations in Column A, Gene name in Column B. The log2fold change values, raw p values and the adjusted p values as obtained from the DESeq2 analysis are shown in columns C, D, and E, respectively. Read counts from the DESeq2 analysis for the three biological replicates of wildtype are listed in columns F, G and H. Read counts from the DESeq2 analysis for the three biological replicates of adr-1(-);adr-2(-) animals are listed in columns I, J and K . "	"Supplemental Table 4: Differentially expressed genes in adr-1(-);adr-2(-) animals grown on OP40 for 7 hours compared to wildtype animals grown on OP50 for 7 hours (Related to Figure 4). DESeq2 was used to identify differentially expressed genes in adr-1(-);adr-2(-) double mutant animals compared to wildtype animals. p-value<0.05 were taken as targets misregulated in absence of ad-1 and adr-2. Differentially expressed genes are listed as Wormbase annotations in Column A, Gene name in Column B. The log2fold change values, raw p values and the adjusted p values as obtained from the DESeq2 analysis are shown in columns C, D, and E, respectively. Read counts from the DESeq2 analysis for the three biological replicates of wildtype are listed in columns F, G and H. Read counts from the DESeq2 analysis for the three biological replicates of adr-1(-);adr-2(-) animals are listed in columns I, J and K . "	"Supplemental Table 4: Differentially expressed genes in adr-1(-);adr-2(-) animals grown on OP40 for 7 hours compared to wildtype animals grown on OP50 for 7 hours (Related to Figure 4). DESeq2 was used to identify differentially expressed genes in adr-1(-);adr-2(-) double mutant animals compared to wildtype animals. p-value<0.05 were taken as targets misregulated in absence of ad-1 and adr-2. Differentially expressed genes are listed as Wormbase annotations in Column A, Gene name in Column B. The log2fold change values, raw p values and the adjusted p values as obtained from the DESeq2 analysis are shown in columns C, D, and E, respectively. Read counts from the DESeq2 analysis for the three biological replicates of wildtype are listed in columns F, G and H. Read counts from the DESeq2 analysis for the three biological replicates of adr-1(-);adr-2(-) animals are listed in columns I, J and K . "	"Supplemental Table 4: Differentially expressed genes in adr-1(-);adr-2(-) animals grown on OP40 for 7 hours compared to wildtype animals grown on OP50 for 7 hours (Related to Figure 4). DESeq2 was used to identify differentially expressed genes in adr-1(-);adr-2(-) double mutant animals compared to wildtype animals. p-value<0.05 were taken as targets misregulated in absence of ad-1 and adr-2. Differentially expressed genes are listed as Wormbase annotations in Column A, Gene name in Column B. The log2fold change values, raw p values and the adjusted p values as obtained from the DESeq2 analysis are shown in columns C, D, and E, respectively. Read counts from the DESeq2 analysis for the three biological replicates of wildtype are listed in columns F, G and H. Read counts from the DESeq2 analysis for the three biological replicates of adr-1(-);adr-2(-) animals are listed in columns I, J and K . "	"Supplemental Table 4: Differentially expressed genes in adr-1(-);adr-2(-) animals grown on OP40 for 7 hours compared to wildtype animals grown on OP50 for 7 hours (Related to Figure 4). DESeq2 was used to identify differentially expressed genes in adr-1(-);adr-2(-) double mutant animals compared to wildtype animals. p-value<0.05 were taken as targets misregulated in absence of ad-1 and adr-2. Differentially expressed genes are listed as Wormbase annotations in Column A, Gene name in Column B. The log2fold change values, raw p values and the adjusted p values as obtained from the DESeq2 analysis are shown in columns C, D, and E, respectively. Read counts from the DESeq2 analysis for the three biological replicates of wildtype are listed in columns F, G and H. Read counts from the DESeq2 analysis for the three biological replicates of adr-1(-);adr-2(-) animals are listed in columns I, J and K . "	"Supplemental Table 4: Differentially expressed genes in adr-1(-);adr-2(-) animals grown on OP40 for 7 hours compared to wildtype animals grown on OP50 for 7 hours (Related to Figure 4). DESeq2 was used to identify differentially expressed genes in adr-1(-);adr-2(-) double mutant animals compared to wildtype animals. p-value<0.05 were taken as targets misregulated in absence of ad-1 and adr-2. Differentially expressed genes are listed as Wormbase annotations in Column A, Gene name in Column B. The log2fold change values, raw p values and the adjusted p values as obtained from the DESeq2 analysis are shown in columns C, D, and E, respectively. Read counts from the DESeq2 analysis for the three biological replicates of wildtype are listed in columns F, G and H. Read counts from the DESeq2 analysis for the three biological replicates of adr-1(-);adr-2(-) animals are listed in columns I, J and K . "	"Supplemental Table 4: Differentially expressed genes in adr-1(-);adr-2(-) animals grown on OP40 for 7 hours compared to wildtype animals grown on OP50 for 7 hours (Related to Figure 4). DESeq2 was used to identify differentially expressed genes in adr-1(-);adr-2(-) double mutant animals compared to wildtype animals. p-value<0.05 were taken as targets misregulated in absence of ad-1 and adr-2. Differentially expressed genes are listed as Wormbase annotations in Column A, Gene name in Column B. The log2fold change values, raw p values and the adjusted p values as obtained from the DESeq2 analysis are shown in columns C, D, and E, respectively. Read counts from the DESeq2 analysis for the three biological replicates of wildtype are listed in columns F, G and H. Read counts from the DESeq2 analysis for the three biological replicates of adr-1(-);adr-2(-) animals are listed in columns I, J and K . "	"Supplemental Table 4: Differentially expressed genes in adr-1(-);adr-2(-) animals grown on OP40 for 7 hours compared to wildtype animals grown on OP50 for 7 hours (Related to Figure 4). DESeq2 was used to identify differentially expressed genes in adr-1(-);adr-2(-) double mutant animals compared to wildtype animals. p-value<0.05 were taken as targets misregulated in absence of ad-1 and adr-2. Differentially expressed genes are listed as Wormbase annotations in Column A, Gene name in Column B. The log2fold change values, raw p values and the adjusted p values as obtained from the DESeq2 analysis are shown in columns C, D, and E, respectively. Read counts from the DESeq2 analysis for the three biological replicates of wildtype are listed in columns F, G and H. Read counts from the DESeq2 analysis for the three biological replicates of adr-1(-);adr-2(-) animals are listed in columns I, J and K . "	"Supplemental Table 4: Differentially expressed genes in adr-1(-);adr-2(-) animals grown on OP40 for 7 hours compared to wildtype animals grown on OP50 for 7 hours (Related to Figure 4). DESeq2 was used to identify differentially expressed genes in adr-1(-);adr-2(-) double mutant animals compared to wildtype animals. p-value<0.05 were taken as targets misregulated in absence of ad-1 and adr-2. Differentially expressed genes are listed as Wormbase annotations in Column A, Gene name in Column B. The log2fold change values, raw p values and the adjusted p values as obtained from the DESeq2 analysis are shown in columns C, D, and E, respectively. Read counts from the DESeq2 analysis for the three biological replicates of wildtype are listed in columns F, G and H. Read counts from the DESeq2 analysis for the three biological replicates of adr-1(-);adr-2(-) animals are listed in columns I, J and K . "	"Supplemental Table 4: Differentially expressed genes in adr-1(-);adr-2(-) animals grown on OP40 for 7 hours compared to wildtype animals grown on OP50 for 7 hours (Related to Figure 4). DESeq2 was used to identify differentially expressed genes in adr-1(-);adr-2(-) double mutant animals compared to wildtype animals. p-value<0.05 were taken as targets misregulated in absence of ad-1 and adr-2. Differentially expressed genes are listed as Wormbase annotations in Column A, Gene name in Column B. The log2fold change values, raw p values and the adjusted p values as obtained from the DESeq2 analysis are shown in columns C, D, and E, respectively. Read counts from the DESeq2 analysis for the three biological replicates of wildtype are listed in columns F, G and H. Read counts from the DESeq2 analysis for the three biological replicates of adr-1(-);adr-2(-) animals are listed in columns I, J and K . "	"Supplemental Table 4: Differentially expressed genes in adr-1(-);adr-2(-) animals grown on OP40 for 7 hours compared to wildtype animals grown on OP50 for 7 hours (Related to Figure 4). DESeq2 was used to identify differentially expressed genes in adr-1(-);adr-2(-) double mutant animals compared to wildtype animals. p-value<0.05 were taken as targets misregulated in absence of ad-1 and adr-2. Differentially expressed genes are listed as Wormbase annotations in Column A, Gene name in Column B. The log2fold change values, raw p values and the adjusted p values as obtained from the DESeq2 analysis are shown in columns C, D, and E, respectively. Read counts from the DESeq2 analysis for the three biological replicates of wildtype are listed in columns F, G and H. Read counts from the DESeq2 analysis for the three biological replicates of adr-1(-);adr-2(-) animals are listed in columns I, J and K . "	"Supplemental Table 4: Differentially expressed genes in adr-1(-);adr-2(-) animals grown on OP40 for 7 hours compared to wildtype animals grown on OP50 for 7 hours (Related to Figure 4). DESeq2 was used to identify differentially expressed genes in adr-1(-);adr-2(-) double mutant animals compared to wildtype animals. p-value<0.05 were taken as targets misregulated in absence of ad-1 and adr-2. Differentially expressed genes are listed as Wormbase annotations in Column A, Gene name in Column B. The log2fold change values, raw p values and the adjusted p values as obtained from the DESeq2 analysis are shown in columns C, D, and E, respectively. Read counts from the DESeq2 analysis for the three biological replicates of wildtype are listed in columns F, G and H. Read counts from the DESeq2 analysis for the three biological replicates of adr-1(-);adr-2(-) animals are listed in columns I, J and K . "	"Supplemental Table 4: Differentially expressed genes in adr-1(-);adr-2(-) animals grown on OP40 for 7 hours compared to wildtype animals grown on OP50 for 7 hours (Related to Figure 4). DESeq2 was used to identify differentially expressed genes in adr-1(-);adr-2(-) double mutant animals compared to wildtype animals. p-value<0.05 were taken as targets misregulated in absence of ad-1 and adr-2. Differentially expressed genes are listed as Wormbase annotations in Column A, Gene name in Column B. The log2fold change values, raw p values and the adjusted p values as obtained from the DESeq2 analysis are shown in columns C, D, and E, respectively. Read counts from the DESeq2 analysis for the three biological replicates of wildtype are listed in columns F, G and H. Read counts from the DESeq2 analysis for the three biological replicates of adr-1(-);adr-2(-) animals are listed in columns I, J and K . "	"Supplemental Table 4: Differentially expressed genes in adr-1(-);adr-2(-) animals grown on OP40 for 7 hours compared to wildtype animals grown on OP50 for 7 hours (Related to Figure 4). DESeq2 was used to identify differentially expressed genes in adr-1(-);adr-2(-) double mutant animals compared to wildtype animals. p-value<0.05 were taken as targets misregulated in absence of ad-1 and adr-2. Differentially expressed genes are listed as Wormbase annotations in Column A, Gene name in Column B. The log2fold change values, raw p values and the adjusted p values as obtained from the DESeq2 analysis are shown in columns C, D, and E, respectively. Read counts from the DESeq2 analysis for the three biological replicates of wildtype are listed in columns F, G and H. Read counts from the DESeq2 analysis for the three biological replicates of adr-1(-);adr-2(-) animals are listed in columns I, J and K . "	"Supplemental Table 4: Differentially expressed genes in adr-1(-);adr-2(-) animals grown on OP40 for 7 hours compared to wildtype animals grown on OP50 for 7 hours (Related to Figure 4). DESeq2 was used to identify differentially expressed genes in adr-1(-);adr-2(-) double mutant animals compared to wildtype animals. p-value<0.05 were taken as targets misregulated in absence of ad-1 and adr-2. Differentially expressed genes are listed as Wormbase annotations in Column A, Gene name in Column B. The log2fold change values, raw p values and the adjusted p values as obtained from the DESeq2 analysis are shown in columns C, D, and E, respectively. Read counts from the DESeq2 analysis for the three biological replicates of wildtype are listed in columns F, G and H. Read counts from the DESeq2 analysis for the three biological replicates of adr-1(-);adr-2(-) animals are listed in columns I, J and K . "	"Supplemental Table 4: Differentially expressed genes in adr-1(-);adr-2(-) animals grown on OP40 for 7 hours compared to wildtype animals grown on OP50 for 7 hours (Related to Figure 4). DESeq2 was used to identify differentially expressed genes in adr-1(-);adr-2(-) double mutant animals compared to wildtype animals. p-value<0.05 were taken as targets misregulated in absence of ad-1 and adr-2. Differentially expressed genes are listed as Wormbase annotations in Column A, Gene name in Column B. The log2fold change values, raw p values and the adjusted p values as obtained from the DESeq2 analysis are shown in columns C, D, and E, respectively. Read counts from the DESeq2 analysis for the three biological replicates of wildtype are listed in columns F, G and H. Read counts from the DESeq2 analysis for the three biological replicates of adr-1(-);adr-2(-) animals are listed in columns I, J and K . "	"Supplemental Table 4: Differentially expressed genes in adr-1(-);adr-2(-) animals grown on OP40 for 7 hours compared to wildtype animals grown on OP50 for 7 hours (Related to Figure 4). DESeq2 was used to identify differentially expressed genes in adr-1(-);adr-2(-) double mutant animals compared to wildtype animals. p-value<0.05 were taken as targets misregulated in absence of ad-1 and adr-2. Differentially expressed genes are listed as Wormbase annotations in Column A, Gene name in Column B. The log2fold change values, raw p values and the adjusted p values as obtained from the DESeq2 analysis are shown in columns C, D, and E, respectively. Read counts from the DESeq2 analysis for the three biological replicates of wildtype are listed in columns F, G and H. Read counts from the DESeq2 analysis for the three biological replicates of adr-1(-);adr-2(-) animals are listed in columns I, J and K . "	"Supplemental Table 4: Differentially expressed genes in adr-1(-);adr-2(-) animals grown on OP40 for 7 hours compared to wildtype animals grown on OP50 for 7 hours (Related to Figure 4). DESeq2 was used to identify differentially expressed genes in adr-1(-);adr-2(-) double mutant animals compared to wildtype animals. p-value<0.05 were taken as targets misregulated in absence of ad-1 and adr-2. Differentially expressed genes are listed as Wormbase annotations in Column A, Gene name in Column B. The log2fold change values, raw p values and the adjusted p values as obtained from the DESeq2 analysis are shown in columns C, D, and E, respectively. Read counts from the DESeq2 analysis for the three biological replicates of wildtype are listed in columns F, G and H. Read counts from the DESeq2 analysis for the three biological replicates of adr-1(-);adr-2(-) animals are listed in columns I, J and K . "	"Supplemental Table 4: Differentially expressed genes in adr-1(-);adr-2(-) animals grown on OP40 for 7 hours compared to wildtype animals grown on OP50 for 7 hours (Related to Figure 4). DESeq2 was used to identify differentially expressed genes in adr-1(-);adr-2(-) double mutant animals compared to wildtype animals. p-value<0.05 were taken as targets misregulated in absence of ad-1 and adr-2. Differentially expressed genes are listed as Wormbase annotations in Column A, Gene name in Column B. The log2fold change values, raw p values and the adjusted p values as obtained from the DESeq2 analysis are shown in columns C, D, and E, respectively. Read counts from the DESeq2 analysis for the three biological replicates of wildtype are listed in columns F, G and H. Read counts from the DESeq2 analysis for the three biological replicates of adr-1(-);adr-2(-) animals are listed in columns I, J and K . "	"Supplemental Table 4: Differentially expressed genes in adr-1(-);adr-2(-) animals grown on OP40 for 7 hours compared to wildtype animals grown on OP50 for 7 hours (Related to Figure 4). DESeq2 was used to identify differentially expressed genes in adr-1(-);adr-2(-) double mutant animals compared to wildtype animals. p-value<0.05 were taken as targets misregulated in absence of ad-1 and adr-2. Differentially expressed genes are listed as Wormbase annotations in Column A, Gene name in Column B. The log2fold change values, raw p values and the adjusted p values as obtained from the DESeq2 analysis are shown in columns C, D, and E, respectively. Read counts from the DESeq2 analysis for the three biological replicates of wildtype are listed in columns F, G and H. Read counts from the DESeq2 analysis for the three biological replicates of adr-1(-);adr-2(-) animals are listed in columns I, J and K . "	"Supplemental Table 4: Differentially expressed genes in adr-1(-);adr-2(-) animals grown on OP40 for 7 hours compared to wildtype animals grown on OP50 for 7 hours (Related to Figure 4). DESeq2 was used to identify differentially expressed genes in adr-1(-);adr-2(-) double mutant animals compared to wildtype animals. p-value<0.05 were taken as targets misregulated in absence of ad-1 and adr-2. Differentially expressed genes are listed as Wormbase annotations in Column A, Gene name in Column B. The log2fold change values, raw p values and the adjusted p values as obtained from the DESeq2 analysis are shown in columns C, D, and E, respectively. Read counts from the DESeq2 analysis for the three biological replicates of wildtype are listed in columns F, G and H. Read counts from the DESeq2 analysis for the three biological replicates of adr-1(-);adr-2(-) animals are listed in columns I, J and K . "	"Supplemental Table 4: Differentially expressed genes in adr-1(-);adr-2(-) animals grown on OP40 for 7 hours compared to wildtype animals grown on OP50 for 7 hours (Related to Figure 4). DESeq2 was used to identify differentially expressed genes in adr-1(-);adr-2(-) double mutant animals compared to wildtype animals. p-value<0.05 were taken as targets misregulated in absence of ad-1 and adr-2. Differentially expressed genes are listed as Wormbase annotations in Column A, Gene name in Column B. The log2fold change values, raw p values and the adjusted p values as obtained from the DESeq2 analysis are shown in columns C, D, and E, respectively. Read counts from the DESeq2 analysis for the three biological replicates of wildtype are listed in columns F, G and H. Read counts from the DESeq2 analysis for the three biological replicates of adr-1(-);adr-2(-) animals are listed in columns I, J and K . "	"Supplemental Table 4: Differentially expressed genes in adr-1(-);adr-2(-) animals grown on OP40 for 7 hours compared to wildtype animals grown on OP50 for 7 hours (Related to Figure 4). DESeq2 was used to identify differentially expressed genes in adr-1(-);adr-2(-) double mutant animals compared to wildtype animals. p-value<0.05 were taken as targets misregulated in absence of ad-1 and adr-2. Differentially expressed genes are listed as Wormbase annotations in Column A, Gene name in Column B. The log2fold change values, raw p values and the adjusted p values as obtained from the DESeq2 analysis are shown in columns C, D, and E, respectively. Read counts from the DESeq2 analysis for the three biological replicates of wildtype are listed in columns F, G and H. Read counts from the DESeq2 analysis for the three biological replicates of adr-1(-);adr-2(-) animals are listed in columns I, J and K . "	"Supplemental Table 4: Differentially expressed genes in adr-1(-);adr-2(-) animals grown on OP40 for 7 hours compared to wildtype animals grown on OP50 for 7 hours (Related to Figure 4). DESeq2 was used to identify differentially expressed genes in adr-1(-);adr-2(-) double mutant animals compared to wildtype animals. p-value<0.05 were taken as targets misregulated in absence of ad-1 and adr-2. Differentially expressed genes are listed as Wormbase annotations in Column A, Gene name in Column B. The log2fold change values, raw p values and the adjusted p values as obtained from the DESeq2 analysis are shown in columns C, D, and E, respectively. Read counts from the DESeq2 analysis for the three biological replicates of wildtype are listed in columns F, G and H. Read counts from the DESeq2 analysis for the three biological replicates of adr-1(-);adr-2(-) animals are listed in columns I, J and K . "	"Supplemental Table 4: Differentially expressed genes in adr-1(-);adr-2(-) animals grown on OP40 for 7 hours compared to wildtype animals grown on OP50 for 7 hours (Related to Figure 4). DESeq2 was used to identify differentially expressed genes in adr-1(-);adr-2(-) double mutant animals compared to wildtype animals. p-value<0.05 were taken as targets misregulated in absence of ad-1 and adr-2. Differentially expressed genes are listed as Wormbase annotations in Column A, Gene name in Column B. The log2fold change values, raw p values and the adjusted p values as obtained from the DESeq2 analysis are shown in columns C, D, and E, respectively. Read counts from the DESeq2 analysis for the three biological replicates of wildtype are listed in columns F, G and H. Read counts from the DESeq2 analysis for the three biological replicates of adr-1(-);adr-2(-) animals are listed in columns I, J and K . "	"Supplemental Table 4: Differentially expressed genes in adr-1(-);adr-2(-) animals grown on OP40 for 7 hours compared to wildtype animals grown on OP50 for 7 hours (Related to Figure 4). DESeq2 was used to identify differentially expressed genes in adr-1(-);adr-2(-) double mutant animals compared to wildtype animals. p-value<0.05 were taken as targets misregulated in absence of ad-1 and adr-2. Differentially expressed genes are listed as Wormbase annotations in Column A, Gene name in Column B. The log2fold change values, raw p values and the adjusted p values as obtained from the DESeq2 analysis are shown in columns C, D, and E, respectively. Read counts from the DESeq2 analysis for the three biological replicates of wildtype are listed in columns F, G and H. Read counts from the DESeq2 analysis for the three biological replicates of adr-1(-);adr-2(-) animals are listed in columns I, J and K . "	"Supplemental Table 4: Differentially expressed genes in adr-1(-);adr-2(-) animals grown on OP40 for 7 hours compared to wildtype animals grown on OP50 for 7 hours (Related to Figure 4). DESeq2 was used to identify differentially expressed genes in adr-1(-);adr-2(-) double mutant animals compared to wildtype animals. p-value<0.05 were taken as targets misregulated in absence of ad-1 and adr-2. Differentially expressed genes are listed as Wormbase annotations in Column A, Gene name in Column B. The log2fold change values, raw p values and the adjusted p values as obtained from the DESeq2 analysis are shown in columns C, D, and E, respectively. Read counts from the DESeq2 analysis for the three biological replicates of wildtype are listed in columns F, G and H. Read counts from the DESeq2 analysis for the three biological replicates of adr-1(-);adr-2(-) animals are listed in columns I, J and K . "	"Supplemental Table 4: Differentially expressed genes in adr-1(-);adr-2(-) animals grown on OP40 for 7 hours compared to wildtype animals grown on OP50 for 7 hours (Related to Figure 4). DESeq2 was used to identify differentially expressed genes in adr-1(-);adr-2(-) double mutant animals compared to wildtype animals. p-value<0.05 were taken as targets misregulated in absence of ad-1 and adr-2. Differentially expressed genes are listed as Wormbase annotations in Column A, Gene name in Column B. The log2fold change values, raw p values and the adjusted p values as obtained from the DESeq2 analysis are shown in columns C, D, and E, respectively. Read counts from the DESeq2 analysis for the three biological replicates of wildtype are listed in columns F, G and H. Read counts from the DESeq2 analysis for the three biological replicates of adr-1(-);adr-2(-) animals are listed in columns I, J and K . "	"Supplemental Table 4: Differentially expressed genes in adr-1(-);adr-2(-) animals grown on OP40 for 7 hours compared to wildtype animals grown on OP50 for 7 hours (Related to Figure 4). DESeq2 was used to identify differentially expressed genes in adr-1(-);adr-2(-) double mutant animals compared to wildtype animals. p-value<0.05 were taken as targets misregulated in absence of ad-1 and adr-2. Differentially expressed genes are listed as Wormbase annotations in Column A, Gene name in Column B. The log2fold change values, raw p values and the adjusted p values as obtained from the DESeq2 analysis are shown in columns C, D, and E, respectively. Read counts from the DESeq2 analysis for the three biological replicates of wildtype are listed in columns F, G and H. Read counts from the DESeq2 analysis for the three biological replicates of adr-1(-);adr-2(-) animals are listed in columns I, J and K . "	"Supplemental Table 4: Differentially expressed genes in adr-1(-);adr-2(-) animals grown on OP40 for 7 hours compared to wildtype animals grown on OP50 for 7 hours (Related to Figure 4). DESeq2 was used to identify differentially expressed genes in adr-1(-);adr-2(-) double mutant animals compared to wildtype animals. p-value<0.05 were taken as targets misregulated in absence of ad-1 and adr-2. Differentially expressed genes are listed as Wormbase annotations in Column A, Gene name in Column B. The log2fold change values, raw p values and the adjusted p values as obtained from the DESeq2 analysis are shown in columns C, D, and E, respectively. Read counts from the DESeq2 analysis for the three biological replicates of wildtype are listed in columns F, G and H. Read counts from the DESeq2 analysis for the three biological replicates of adr-1(-);adr-2(-) animals are listed in columns I, J and K . "	"Supplemental Table 4: Differentially expressed genes in adr-1(-);adr-2(-) animals grown on OP40 for 7 hours compared to wildtype animals grown on OP50 for 7 hours (Related to Figure 4). DESeq2 was used to identify differentially expressed genes in adr-1(-);adr-2(-) double mutant animals compared to wildtype animals. p-value<0.05 were taken as targets misregulated in absence of ad-1 and adr-2. Differentially expressed genes are listed as Wormbase annotations in Column A, Gene name in Column B. The log2fold change values, raw p values and the adjusted p values as obtained from the DESeq2 analysis are shown in columns C, D, and E, respectively. Read counts from the DESeq2 analysis for the three biological replicates of wildtype are listed in columns F, G and H. Read counts from the DESeq2 analysis for the three biological replicates of adr-1(-);adr-2(-) animals are listed in columns I, J and K . "	"Supplemental Table 4: Differentially expressed genes in adr-1(-);adr-2(-) animals grown on OP40 for 7 hours compared to wildtype animals grown on OP50 for 7 hours (Related to Figure 4). DESeq2 was used to identify differentially expressed genes in adr-1(-);adr-2(-) double mutant animals compared to wildtype animals. p-value<0.05 were taken as targets misregulated in absence of ad-1 and adr-2. Differentially expressed genes are listed as Wormbase annotations in Column A, Gene name in Column B. The log2fold change values, raw p values and the adjusted p values as obtained from the DESeq2 analysis are shown in columns C, D, and E, respectively. Read counts from the DESeq2 analysis for the three biological replicates of wildtype are listed in columns F, G and H. Read counts from the DESeq2 analysis for the three biological replicates of adr-1(-);adr-2(-) animals are listed in columns I, J and K . "	"Supplemental Table 4: Differentially expressed genes in adr-1(-);adr-2(-) animals grown on OP40 for 7 hours compared to wildtype animals grown on OP50 for 7 hours (Related to Figure 4). DESeq2 was used to identify differentially expressed genes in adr-1(-);adr-2(-) double mutant animals compared to wildtype animals. p-value<0.05 were taken as targets misregulated in absence of ad-1 and adr-2. Differentially expressed genes are listed as Wormbase annotations in Column A, Gene name in Column B. The log2fold change values, raw p values and the adjusted p values as obtained from the DESeq2 analysis are shown in columns C, D, and E, respectively. Read counts from the DESeq2 analysis for the three biological replicates of wildtype are listed in columns F, G and H. Read counts from the DESeq2 analysis for the three biological replicates of adr-1(-);adr-2(-) animals are listed in columns I, J and K . "	"Supplemental Table 4: Differentially expressed genes in adr-1(-);adr-2(-) animals grown on OP40 for 7 hours compared to wildtype animals grown on OP50 for 7 hours (Related to Figure 4). DESeq2 was used to identify differentially expressed genes in adr-1(-);adr-2(-) double mutant animals compared to wildtype animals. p-value<0.05 were taken as targets misregulated in absence of ad-1 and adr-2. Differentially expressed genes are listed as Wormbase annotations in Column A, Gene name in Column B. The log2fold change values, raw p values and the adjusted p values as obtained from the DESeq2 analysis are shown in columns C, D, and E, respectively. Read counts from the DESeq2 analysis for the three biological replicates of wildtype are listed in columns F, G and H. Read counts from the DESeq2 analysis for the three biological replicates of adr-1(-);adr-2(-) animals are listed in columns I, J and K . "	"Supplemental Table 4: Differentially expressed genes in adr-1(-);adr-2(-) animals grown on OP40 for 7 hours compared to wildtype animals grown on OP50 for 7 hours (Related to Figure 4). DESeq2 was used to identify differentially expressed genes in adr-1(-);adr-2(-) double mutant animals compared to wildtype animals. p-value<0.05 were taken as targets misregulated in absence of ad-1 and adr-2. Differentially expressed genes are listed as Wormbase annotations in Column A, Gene name in Column B. The log2fold change values, raw p values and the adjusted p values as obtained from the DESeq2 analysis are shown in columns C, D, and E, respectively. Read counts from the DESeq2 analysis for the three biological replicates of wildtype are listed in columns F, G and H. Read counts from the DESeq2 analysis for the three biological replicates of adr-1(-);adr-2(-) animals are listed in columns I, J and K . "	"Supplemental Table 4: Differentially expressed genes in adr-1(-);adr-2(-) animals grown on OP40 for 7 hours compared to wildtype animals grown on OP50 for 7 hours (Related to Figure 4). DESeq2 was used to identify differentially expressed genes in adr-1(-);adr-2(-) double mutant animals compared to wildtype animals. p-value<0.05 were taken as targets misregulated in absence of ad-1 and adr-2. Differentially expressed genes are listed as Wormbase annotations in Column A, Gene name in Column B. The log2fold change values, raw p values and the adjusted p values as obtained from the DESeq2 analysis are shown in columns C, D, and E, respectively. Read counts from the DESeq2 analysis for the three biological replicates of wildtype are listed in columns F, G and H. Read counts from the DESeq2 analysis for the three biological replicates of adr-1(-);adr-2(-) animals are listed in columns I, J and K . "	"Supplemental Table 4: Differentially expressed genes in adr-1(-);adr-2(-) animals grown on OP40 for 7 hours compared to wildtype animals grown on OP50 for 7 hours (Related to Figure 4). DESeq2 was used to identify differentially expressed genes in adr-1(-);adr-2(-) double mutant animals compared to wildtype animals. p-value<0.05 were taken as targets misregulated in absence of ad-1 and adr-2. Differentially expressed genes are listed as Wormbase annotations in Column A, Gene name in Column B. The log2fold change values, raw p values and the adjusted p values as obtained from the DESeq2 analysis are shown in columns C, D, and E, respectively. Read counts from the DESeq2 analysis for the three biological replicates of wildtype are listed in columns F, G and H. Read counts from the DESeq2 analysis for the three biological replicates of adr-1(-);adr-2(-) animals are listed in columns I, J and K . "	"Supplemental Table 4: Differentially expressed genes in adr-1(-);adr-2(-) animals grown on OP40 for 7 hours compared to wildtype animals grown on OP50 for 7 hours (Related to Figure 4). DESeq2 was used to identify differentially expressed genes in adr-1(-);adr-2(-) double mutant animals compared to wildtype animals. p-value<0.05 were taken as targets misregulated in absence of ad-1 and adr-2. Differentially expressed genes are listed as Wormbase annotations in Column A, Gene name in Column B. The log2fold change values, raw p values and the adjusted p values as obtained from the DESeq2 analysis are shown in columns C, D, and E, respectively. Read counts from the DESeq2 analysis for the three biological replicates of wildtype are listed in columns F, G and H. Read counts from the DESeq2 analysis for the three biological replicates of adr-1(-);adr-2(-) animals are listed in columns I, J and K . "	"Supplemental Table 4: Differentially expressed genes in adr-1(-);adr-2(-) animals grown on OP40 for 7 hours compared to wildtype animals grown on OP50 for 7 hours (Related to Figure 4). DESeq2 was used to identify differentially expressed genes in adr-1(-);adr-2(-) double mutant animals compared to wildtype animals. p-value<0.05 were taken as targets misregulated in absence of ad-1 and adr-2. Differentially expressed genes are listed as Wormbase annotations in Column A, Gene name in Column B. The log2fold change values, raw p values and the adjusted p values as obtained from the DESeq2 analysis are shown in columns C, D, and E, respectively. Read counts from the DESeq2 analysis for the three biological replicates of wildtype are listed in columns F, G and H. Read counts from the DESeq2 analysis for the three biological replicates of adr-1(-);adr-2(-) animals are listed in columns I, J and K . "	"Supplemental Table 4: Differentially expressed genes in adr-1(-);adr-2(-) animals grown on OP40 for 7 hours compared to wildtype animals grown on OP50 for 7 hours (Related to Figure 4). DESeq2 was used to identify differentially expressed genes in adr-1(-);adr-2(-) double mutant animals compared to wildtype animals. p-value<0.05 were taken as targets misregulated in absence of ad-1 and adr-2. Differentially expressed genes are listed as Wormbase annotations in Column A, Gene name in Column B. The log2fold change values, raw p values and the adjusted p values as obtained from the DESeq2 analysis are shown in columns C, D, and E, respectively. Read counts from the DESeq2 analysis for the three biological replicates of wildtype are listed in columns F, G and H. Read counts from the DESeq2 analysis for the three biological replicates of adr-1(-);adr-2(-) animals are listed in columns I, J and K . "	"Supplemental Table 4: Differentially expressed genes in adr-1(-);adr-2(-) animals grown on OP40 for 7 hours compared to wildtype animals grown on OP50 for 7 hours (Related to Figure 4). DESeq2 was used to identify differentially expressed genes in adr-1(-);adr-2(-) double mutant animals compared to wildtype animals. p-value<0.05 were taken as targets misregulated in absence of ad-1 and adr-2. Differentially expressed genes are listed as Wormbase annotations in Column A, Gene name in Column B. The log2fold change values, raw p values and the adjusted p values as obtained from the DESeq2 analysis are shown in columns C, D, and E, respectively. Read counts from the DESeq2 analysis for the three biological replicates of wildtype are listed in columns F, G and H. Read counts from the DESeq2 analysis for the three biological replicates of adr-1(-);adr-2(-) animals are listed in columns I, J and K . "	"Supplemental Table 4: Differentially expressed genes in adr-1(-);adr-2(-) animals grown on OP40 for 7 hours compared to wildtype animals grown on OP50 for 7 hours (Related to Figure 4). DESeq2 was used to identify differentially expressed genes in adr-1(-);adr-2(-) double mutant animals compared to wildtype animals. p-value<0.05 were taken as targets misregulated in absence of ad-1 and adr-2. Differentially expressed genes are listed as Wormbase annotations in Column A, Gene name in Column B. The log2fold change values, raw p values and the adjusted p values as obtained from the DESeq2 analysis are shown in columns C, D, and E, respectively. Read counts from the DESeq2 analysis for the three biological replicates of wildtype are listed in columns F, G and H. Read counts from the DESeq2 analysis for the three biological replicates of adr-1(-);adr-2(-) animals are listed in columns I, J and K . "	"Supplemental Table 4: Differentially expressed genes in adr-1(-);adr-2(-) animals grown on OP40 for 7 hours compared to wildtype animals grown on OP50 for 7 hours (Related to Figure 4). DESeq2 was used to identify differentially expressed genes in adr-1(-);adr-2(-) double mutant animals compared to wildtype animals. p-value<0.05 were taken as targets misregulated in absence of ad-1 and adr-2. Differentially expressed genes are listed as Wormbase annotations in Column A, Gene name in Column B. The log2fold change values, raw p values and the adjusted p values as obtained from the DESeq2 analysis are shown in columns C, D, and E, respectively. Read counts from the DESeq2 analysis for the three biological replicates of wildtype are listed in columns F, G and H. Read counts from the DESeq2 analysis for the three biological replicates of adr-1(-);adr-2(-) animals are listed in columns I, J and K . "	"Supplemental Table 4: Differentially expressed genes in adr-1(-);adr-2(-) animals grown on OP40 for 7 hours compared to wildtype animals grown on OP50 for 7 hours (Related to Figure 4). DESeq2 was used to identify differentially expressed genes in adr-1(-);adr-2(-) double mutant animals compared to wildtype animals. p-value<0.05 were taken as targets misregulated in absence of ad-1 and adr-2. Differentially expressed genes are listed as Wormbase annotations in Column A, Gene name in Column B. The log2fold change values, raw p values and the adjusted p values as obtained from the DESeq2 analysis are shown in columns C, D, and E, respectively. Read counts from the DESeq2 analysis for the three biological replicates of wildtype are listed in columns F, G and H. Read counts from the DESeq2 analysis for the three biological replicates of adr-1(-);adr-2(-) animals are listed in columns I, J and K . "	"Supplemental Table 4: Differentially expressed genes in adr-1(-);adr-2(-) animals grown on OP40 for 7 hours compared to wildtype animals grown on OP50 for 7 hours (Related to Figure 4). DESeq2 was used to identify differentially expressed genes in adr-1(-);adr-2(-) double mutant animals compared to wildtype animals. p-value<0.05 were taken as targets misregulated in absence of ad-1 and adr-2. Differentially expressed genes are listed as Wormbase annotations in Column A, Gene name in Column B. The log2fold change values, raw p values and the adjusted p values as obtained from the DESeq2 analysis are shown in columns C, D, and E, respectively. Read counts from the DESeq2 analysis for the three biological replicates of wildtype are listed in columns F, G and H. Read counts from the DESeq2 analysis for the three biological replicates of adr-1(-);adr-2(-) animals are listed in columns I, J and K . "	"Supplemental Table 4: Differentially expressed genes in adr-1(-);adr-2(-) animals grown on OP40 for 7 hours compared to wildtype animals grown on OP50 for 7 hours (Related to Figure 4). DESeq2 was used to identify differentially expressed genes in adr-1(-);adr-2(-) double mutant animals compared to wildtype animals. p-value<0.05 were taken as targets misregulated in absence of ad-1 and adr-2. Differentially expressed genes are listed as Wormbase annotations in Column A, Gene name in Column B. The log2fold change values, raw p values and the adjusted p values as obtained from the DESeq2 analysis are shown in columns C, D, and E, respectively. Read counts from the DESeq2 analysis for the three biological replicates of wildtype are listed in columns F, G and H. Read counts from the DESeq2 analysis for the three biological replicates of adr-1(-);adr-2(-) animals are listed in columns I, J and K . "	"Supplemental Table 4: Differentially expressed genes in adr-1(-);adr-2(-) animals grown on OP40 for 7 hours compared to wildtype animals grown on OP50 for 7 hours (Related to Figure 4). DESeq2 was used to identify differentially expressed genes in adr-1(-);adr-2(-) double mutant animals compared to wildtype animals. p-value<0.05 were taken as targets misregulated in absence of ad-1 and adr-2. Differentially expressed genes are listed as Wormbase annotations in Column A, Gene name in Column B. The log2fold change values, raw p values and the adjusted p values as obtained from the DESeq2 analysis are shown in columns C, D, and E, respectively. Read counts from the DESeq2 analysis for the three biological replicates of wildtype are listed in columns F, G and H. Read counts from the DESeq2 analysis for the three biological replicates of adr-1(-);adr-2(-) animals are listed in columns I, J and K . "	"Supplemental Table 4: Differentially expressed genes in adr-1(-);adr-2(-) animals grown on OP40 for 7 hours compared to wildtype animals grown on OP50 for 7 hours (Related to Figure 4). DESeq2 was used to identify differentially expressed genes in adr-1(-);adr-2(-) double mutant animals compared to wildtype animals. p-value<0.05 were taken as targets misregulated in absence of ad-1 and adr-2. Differentially expressed genes are listed as Wormbase annotations in Column A, Gene name in Column B. The log2fold change values, raw p values and the adjusted p values as obtained from the DESeq2 analysis are shown in columns C, D, and E, respectively. Read counts from the DESeq2 analysis for the three biological replicates of wildtype are listed in columns F, G and H. Read counts from the DESeq2 analysis for the three biological replicates of adr-1(-);adr-2(-) animals are listed in columns I, J and K . "	"Supplemental Table 4: Differentially expressed genes in adr-1(-);adr-2(-) animals grown on OP40 for 7 hours compared to wildtype animals grown on OP50 for 7 hours (Related to Figure 4). DESeq2 was used to identify differentially expressed genes in adr-1(-);adr-2(-) double mutant animals compared to wildtype animals. p-value<0.05 were taken as targets misregulated in absence of ad-1 and adr-2. Differentially expressed genes are listed as Wormbase annotations in Column A, Gene name in Column B. The log2fold change values, raw p values and the adjusted p values as obtained from the DESeq2 analysis are shown in columns C, D, and E, respectively. Read counts from the DESeq2 analysis for the three biological replicates of wildtype are listed in columns F, G and H. Read counts from the DESeq2 analysis for the three biological replicates of adr-1(-);adr-2(-) animals are listed in columns I, J and K . "	"Supplemental Table 4: Differentially expressed genes in adr-1(-);adr-2(-) animals grown on OP40 for 7 hours compared to wildtype animals grown on OP50 for 7 hours (Related to Figure 4). DESeq2 was used to identify differentially expressed genes in adr-1(-);adr-2(-) double mutant animals compared to wildtype animals. p-value<0.05 were taken as targets misregulated in absence of ad-1 and adr-2. Differentially expressed genes are listed as Wormbase annotations in Column A, Gene name in Column B. The log2fold change values, raw p values and the adjusted p values as obtained from the DESeq2 analysis are shown in columns C, D, and E, respectively. Read counts from the DESeq2 analysis for the three biological replicates of wildtype are listed in columns F, G and H. Read counts from the DESeq2 analysis for the three biological replicates of adr-1(-);adr-2(-) animals are listed in columns I, J and K . "	"Supplemental Table 4: Differentially expressed genes in adr-1(-);adr-2(-) animals grown on OP40 for 7 hours compared to wildtype animals grown on OP50 for 7 hours (Related to Figure 4). DESeq2 was used to identify differentially expressed genes in adr-1(-);adr-2(-) double mutant animals compared to wildtype animals. p-value<0.05 were taken as targets misregulated in absence of ad-1 and adr-2. Differentially expressed genes are listed as Wormbase annotations in Column A, Gene name in Column B. The log2fold change values, raw p values and the adjusted p values as obtained from the DESeq2 analysis are shown in columns C, D, and E, respectively. Read counts from the DESeq2 analysis for the three biological replicates of wildtype are listed in columns F, G and H. Read counts from the DESeq2 analysis for the three biological replicates of adr-1(-);adr-2(-) animals are listed in columns I, J and K . "	"Supplemental Table 4: Differentially expressed genes in adr-1(-);adr-2(-) animals grown on OP40 for 7 hours compared to wildtype animals grown on OP50 for 7 hours (Related to Figure 4). DESeq2 was used to identify differentially expressed genes in adr-1(-);adr-2(-) double mutant animals compared to wildtype animals. p-value<0.05 were taken as targets misregulated in absence of ad-1 and adr-2. Differentially expressed genes are listed as Wormbase annotations in Column A, Gene name in Column B. The log2fold change values, raw p values and the adjusted p values as obtained from the DESeq2 analysis are shown in columns C, D, and E, respectively. Read counts from the DESeq2 analysis for the three biological replicates of wildtype are listed in columns F, G and H. Read counts from the DESeq2 analysis for the three biological replicates of adr-1(-);adr-2(-) animals are listed in columns I, J and K . "	"Supplemental Table 4: Differentially expressed genes in adr-1(-);adr-2(-) animals grown on OP40 for 7 hours compared to wildtype animals grown on OP50 for 7 hours (Related to Figure 4). DESeq2 was used to identify differentially expressed genes in adr-1(-);adr-2(-) double mutant animals compared to wildtype animals. p-value<0.05 were taken as targets misregulated in absence of ad-1 and adr-2. Differentially expressed genes are listed as Wormbase annotations in Column A, Gene name in Column B. The log2fold change values, raw p values and the adjusted p values as obtained from the DESeq2 analysis are shown in columns C, D, and E, respectively. Read counts from the DESeq2 analysis for the three biological replicates of wildtype are listed in columns F, G and H. Read counts from the DESeq2 analysis for the three biological replicates of adr-1(-);adr-2(-) animals are listed in columns I, J and K . "	"Supplemental Table 4: Differentially expressed genes in adr-1(-);adr-2(-) animals grown on OP40 for 7 hours compared to wildtype animals grown on OP50 for 7 hours (Related to Figure 4). DESeq2 was used to identify differentially expressed genes in adr-1(-);adr-2(-) double mutant animals compared to wildtype animals. p-value<0.05 were taken as targets misregulated in absence of ad-1 and adr-2. Differentially expressed genes are listed as Wormbase annotations in Column A, Gene name in Column B. The log2fold change values, raw p values and the adjusted p values as obtained from the DESeq2 analysis are shown in columns C, D, and E, respectively. Read counts from the DESeq2 analysis for the three biological replicates of wildtype are listed in columns F, G and H. Read counts from the DESeq2 analysis for the three biological replicates of adr-1(-);adr-2(-) animals are listed in columns I, J and K . "	"Supplemental Table 4: Differentially expressed genes in adr-1(-);adr-2(-) animals grown on OP40 for 7 hours compared to wildtype animals grown on OP50 for 7 hours (Related to Figure 4). DESeq2 was used to identify differentially expressed genes in adr-1(-);adr-2(-) double mutant animals compared to wildtype animals. p-value<0.05 were taken as targets misregulated in absence of ad-1 and adr-2. Differentially expressed genes are listed as Wormbase annotations in Column A, Gene name in Column B. The log2fold change values, raw p values and the adjusted p values as obtained from the DESeq2 analysis are shown in columns C, D, and E, respectively. Read counts from the DESeq2 analysis for the three biological replicates of wildtype are listed in columns F, G and H. Read counts from the DESeq2 analysis for the three biological replicates of adr-1(-);adr-2(-) animals are listed in columns I, J and K . "	"Supplemental Table 4: Differentially expressed genes in adr-1(-);adr-2(-) animals grown on OP40 for 7 hours compared to wildtype animals grown on OP50 for 7 hours (Related to Figure 4). DESeq2 was used to identify differentially expressed genes in adr-1(-);adr-2(-) double mutant animals compared to wildtype animals. p-value<0.05 were taken as targets misregulated in absence of ad-1 and adr-2. Differentially expressed genes are listed as Wormbase annotations in Column A, Gene name in Column B. The log2fold change values, raw p values and the adjusted p values as obtained from the DESeq2 analysis are shown in columns C, D, and E, respectively. Read counts from the DESeq2 analysis for the three biological replicates of wildtype are listed in columns F, G and H. Read counts from the DESeq2 analysis for the three biological replicates of adr-1(-);adr-2(-) animals are listed in columns I, J and K . "	"Supplemental Table 4: Differentially expressed genes in adr-1(-);adr-2(-) animals grown on OP40 for 7 hours compared to wildtype animals grown on OP50 for 7 hours (Related to Figure 4). DESeq2 was used to identify differentially expressed genes in adr-1(-);adr-2(-) double mutant animals compared to wildtype animals. p-value<0.05 were taken as targets misregulated in absence of ad-1 and adr-2. Differentially expressed genes are listed as Wormbase annotations in Column A, Gene name in Column B. The log2fold change values, raw p values and the adjusted p values as obtained from the DESeq2 analysis are shown in columns C, D, and E, respectively. Read counts from the DESeq2 analysis for the three biological replicates of wildtype are listed in columns F, G and H. Read counts from the DESeq2 analysis for the three biological replicates of adr-1(-);adr-2(-) animals are listed in columns I, J and K . "	"Supplemental Table 4: Differentially expressed genes in adr-1(-);adr-2(-) animals grown on OP40 for 7 hours compared to wildtype animals grown on OP50 for 7 hours (Related to Figure 4). DESeq2 was used to identify differentially expressed genes in adr-1(-);adr-2(-) double mutant animals compared to wildtype animals. p-value<0.05 were taken as targets misregulated in absence of ad-1 and adr-2. Differentially expressed genes are listed as Wormbase annotations in Column A, Gene name in Column B. The log2fold change values, raw p values and the adjusted p values as obtained from the DESeq2 analysis are shown in columns C, D, and E, respectively. Read counts from the DESeq2 analysis for the three biological replicates of wildtype are listed in columns F, G and H. Read counts from the DESeq2 analysis for the three biological replicates of adr-1(-);adr-2(-) animals are listed in columns I, J and K . "	"Supplemental Table 4: Differentially expressed genes in adr-1(-);adr-2(-) animals grown on OP40 for 7 hours compared to wildtype animals grown on OP50 for 7 hours (Related to Figure 4). DESeq2 was used to identify differentially expressed genes in adr-1(-);adr-2(-) double mutant animals compared to wildtype animals. p-value<0.05 were taken as targets misregulated in absence of ad-1 and adr-2. Differentially expressed genes are listed as Wormbase annotations in Column A, Gene name in Column B. The log2fold change values, raw p values and the adjusted p values as obtained from the DESeq2 analysis are shown in columns C, D, and E, respectively. Read counts from the DESeq2 analysis for the three biological replicates of wildtype are listed in columns F, G and H. Read counts from the DESeq2 analysis for the three biological replicates of adr-1(-);adr-2(-) animals are listed in columns I, J and K . "	"Supplemental Table 4: Differentially expressed genes in adr-1(-);adr-2(-) animals grown on OP40 for 7 hours compared to wildtype animals grown on OP50 for 7 hours (Related to Figure 4). DESeq2 was used to identify differentially expressed genes in adr-1(-);adr-2(-) double mutant animals compared to wildtype animals. p-value<0.05 were taken as targets misregulated in absence of ad-1 and adr-2. Differentially expressed genes are listed as Wormbase annotations in Column A, Gene name in Column B. The log2fold change values, raw p values and the adjusted p values as obtained from the DESeq2 analysis are shown in columns C, D, and E, respectively. Read counts from the DESeq2 analysis for the three biological replicates of wildtype are listed in columns F, G and H. Read counts from the DESeq2 analysis for the three biological replicates of adr-1(-);adr-2(-) animals are listed in columns I, J and K . "	"Supplemental Table 4: Differentially expressed genes in adr-1(-);adr-2(-) animals grown on OP40 for 7 hours compared to wildtype animals grown on OP50 for 7 hours (Related to Figure 4). DESeq2 was used to identify differentially expressed genes in adr-1(-);adr-2(-) double mutant animals compared to wildtype animals. p-value<0.05 were taken as targets misregulated in absence of ad-1 and adr-2. Differentially expressed genes are listed as Wormbase annotations in Column A, Gene name in Column B. The log2fold change values, raw p values and the adjusted p values as obtained from the DESeq2 analysis are shown in columns C, D, and E, respectively. Read counts from the DESeq2 analysis for the three biological replicates of wildtype are listed in columns F, G and H. Read counts from the DESeq2 analysis for the three biological replicates of adr-1(-);adr-2(-) animals are listed in columns I, J and K . "	"Supplemental Table 4: Differentially expressed genes in adr-1(-);adr-2(-) animals grown on OP40 for 7 hours compared to wildtype animals grown on OP50 for 7 hours (Related to Figure 4). DESeq2 was used to identify differentially expressed genes in adr-1(-);adr-2(-) double mutant animals compared to wildtype animals. p-value<0.05 were taken as targets misregulated in absence of ad-1 and adr-2. Differentially expressed genes are listed as Wormbase annotations in Column A, Gene name in Column B. The log2fold change values, raw p values and the adjusted p values as obtained from the DESeq2 analysis are shown in columns C, D, and E, respectively. Read counts from the DESeq2 analysis for the three biological replicates of wildtype are listed in columns F, G and H. Read counts from the DESeq2 analysis for the three biological replicates of adr-1(-);adr-2(-) animals are listed in columns I, J and K . "	"Supplemental Table 4: Differentially expressed genes in adr-1(-);adr-2(-) animals grown on OP40 for 7 hours compared to wildtype animals grown on OP50 for 7 hours (Related to Figure 4). DESeq2 was used to identify differentially expressed genes in adr-1(-);adr-2(-) double mutant animals compared to wildtype animals. p-value<0.05 were taken as targets misregulated in absence of ad-1 and adr-2. Differentially expressed genes are listed as Wormbase annotations in Column A, Gene name in Column B. The log2fold change values, raw p values and the adjusted p values as obtained from the DESeq2 analysis are shown in columns C, D, and E, respectively. Read counts from the DESeq2 analysis for the three biological replicates of wildtype are listed in columns F, G and H. Read counts from the DESeq2 analysis for the three biological replicates of adr-1(-);adr-2(-) animals are listed in columns I, J and K . "	"Supplemental Table 4: Differentially expressed genes in adr-1(-);adr-2(-) animals grown on OP40 for 7 hours compared to wildtype animals grown on OP50 for 7 hours (Related to Figure 4). DESeq2 was used to identify differentially expressed genes in adr-1(-);adr-2(-) double mutant animals compared to wildtype animals. p-value<0.05 were taken as targets misregulated in absence of ad-1 and adr-2. Differentially expressed genes are listed as Wormbase annotations in Column A, Gene name in Column B. The log2fold change values, raw p values and the adjusted p values as obtained from the DESeq2 analysis are shown in columns C, D, and E, respectively. Read counts from the DESeq2 analysis for the three biological replicates of wildtype are listed in columns F, G and H. Read counts from the DESeq2 analysis for the three biological replicates of adr-1(-);adr-2(-) animals are listed in columns I, J and K . "	"Supplemental Table 4: Differentially expressed genes in adr-1(-);adr-2(-) animals grown on OP40 for 7 hours compared to wildtype animals grown on OP50 for 7 hours (Related to Figure 4). DESeq2 was used to identify differentially expressed genes in adr-1(-);adr-2(-) double mutant animals compared to wildtype animals. p-value<0.05 were taken as targets misregulated in absence of ad-1 and adr-2. Differentially expressed genes are listed as Wormbase annotations in Column A, Gene name in Column B. The log2fold change values, raw p values and the adjusted p values as obtained from the DESeq2 analysis are shown in columns C, D, and E, respectively. Read counts from the DESeq2 analysis for the three biological replicates of wildtype are listed in columns F, G and H. Read counts from the DESeq2 analysis for the three biological replicates of adr-1(-);adr-2(-) animals are listed in columns I, J and K . "	"Supplemental Table 4: Differentially expressed genes in adr-1(-);adr-2(-) animals grown on OP40 for 7 hours compared to wildtype animals grown on OP50 for 7 hours (Related to Figure 4). DESeq2 was used to identify differentially expressed genes in adr-1(-);adr-2(-) double mutant animals compared to wildtype animals. p-value<0.05 were taken as targets misregulated in absence of ad-1 and adr-2. Differentially expressed genes are listed as Wormbase annotations in Column A, Gene name in Column B. The log2fold change values, raw p values and the adjusted p values as obtained from the DESeq2 analysis are shown in columns C, D, and E, respectively. Read counts from the DESeq2 analysis for the three biological replicates of wildtype are listed in columns F, G and H. Read counts from the DESeq2 analysis for the three biological replicates of adr-1(-);adr-2(-) animals are listed in columns I, J and K . "	"Supplemental Table 4: Differentially expressed genes in adr-1(-);adr-2(-) animals grown on OP40 for 7 hours compared to wildtype animals grown on OP50 for 7 hours (Related to Figure 4). DESeq2 was used to identify differentially expressed genes in adr-1(-);adr-2(-) double mutant animals compared to wildtype animals. p-value<0.05 were taken as targets misregulated in absence of ad-1 and adr-2. Differentially expressed genes are listed as Wormbase annotations in Column A, Gene name in Column B. The log2fold change values, raw p values and the adjusted p values as obtained from the DESeq2 analysis are shown in columns C, D, and E, respectively. Read counts from the DESeq2 analysis for the three biological replicates of wildtype are listed in columns F, G and H. Read counts from the DESeq2 analysis for the three biological replicates of adr-1(-);adr-2(-) animals are listed in columns I, J and K . "	"Supplemental Table 4: Differentially expressed genes in adr-1(-);adr-2(-) animals grown on OP40 for 7 hours compared to wildtype animals grown on OP50 for 7 hours (Related to Figure 4). DESeq2 was used to identify differentially expressed genes in adr-1(-);adr-2(-) double mutant animals compared to wildtype animals. p-value<0.05 were taken as targets misregulated in absence of ad-1 and adr-2. Differentially expressed genes are listed as Wormbase annotations in Column A, Gene name in Column B. The log2fold change values, raw p values and the adjusted p values as obtained from the DESeq2 analysis are shown in columns C, D, and E, respectively. Read counts from the DESeq2 analysis for the three biological replicates of wildtype are listed in columns F, G and H. Read counts from the DESeq2 analysis for the three biological replicates of adr-1(-);adr-2(-) animals are listed in columns I, J and K . "	"Supplemental Table 4: Differentially expressed genes in adr-1(-);adr-2(-) animals grown on OP40 for 7 hours compared to wildtype animals grown on OP50 for 7 hours (Related to Figure 4). DESeq2 was used to identify differentially expressed genes in adr-1(-);adr-2(-) double mutant animals compared to wildtype animals. p-value<0.05 were taken as targets misregulated in absence of ad-1 and adr-2. Differentially expressed genes are listed as Wormbase annotations in Column A, Gene name in Column B. The log2fold change values, raw p values and the adjusted p values as obtained from the DESeq2 analysis are shown in columns C, D, and E, respectively. Read counts from the DESeq2 analysis for the three biological replicates of wildtype are listed in columns F, G and H. Read counts from the DESeq2 analysis for the three biological replicates of adr-1(-);adr-2(-) animals are listed in columns I, J and K . "	"Supplemental Table 4: Differentially expressed genes in adr-1(-);adr-2(-) animals grown on OP40 for 7 hours compared to wildtype animals grown on OP50 for 7 hours (Related to Figure 4). DESeq2 was used to identify differentially expressed genes in adr-1(-);adr-2(-) double mutant animals compared to wildtype animals. p-value<0.05 were taken as targets misregulated in absence of ad-1 and adr-2. Differentially expressed genes are listed as Wormbase annotations in Column A, Gene name in Column B. The log2fold change values, raw p values and the adjusted p values as obtained from the DESeq2 analysis are shown in columns C, D, and E, respectively. Read counts from the DESeq2 analysis for the three biological replicates of wildtype are listed in columns F, G and H. Read counts from the DESeq2 analysis for the three biological replicates of adr-1(-);adr-2(-) animals are listed in columns I, J and K . "	"Supplemental Table 4: Differentially expressed genes in adr-1(-);adr-2(-) animals grown on OP40 for 7 hours compared to wildtype animals grown on OP50 for 7 hours (Related to Figure 4). DESeq2 was used to identify differentially expressed genes in adr-1(-);adr-2(-) double mutant animals compared to wildtype animals. p-value<0.05 were taken as targets misregulated in absence of ad-1 and adr-2. Differentially expressed genes are listed as Wormbase annotations in Column A, Gene name in Column B. The log2fold change values, raw p values and the adjusted p values as obtained from the DESeq2 analysis are shown in columns C, D, and E, respectively. Read counts from the DESeq2 analysis for the three biological replicates of wildtype are listed in columns F, G and H. Read counts from the DESeq2 analysis for the three biological replicates of adr-1(-);adr-2(-) animals are listed in columns I, J and K . "	"Supplemental Table 4: Differentially expressed genes in adr-1(-);adr-2(-) animals grown on OP40 for 7 hours compared to wildtype animals grown on OP50 for 7 hours (Related to Figure 4). DESeq2 was used to identify differentially expressed genes in adr-1(-);adr-2(-) double mutant animals compared to wildtype animals. p-value<0.05 were taken as targets misregulated in absence of ad-1 and adr-2. Differentially expressed genes are listed as Wormbase annotations in Column A, Gene name in Column B. The log2fold change values, raw p values and the adjusted p values as obtained from the DESeq2 analysis are shown in columns C, D, and E, respectively. Read counts from the DESeq2 analysis for the three biological replicates of wildtype are listed in columns F, G and H. Read counts from the DESeq2 analysis for the three biological replicates of adr-1(-);adr-2(-) animals are listed in columns I, J and K . "	"Supplemental Table 4: Differentially expressed genes in adr-1(-);adr-2(-) animals grown on OP40 for 7 hours compared to wildtype animals grown on OP50 for 7 hours (Related to Figure 4). DESeq2 was used to identify differentially expressed genes in adr-1(-);adr-2(-) double mutant animals compared to wildtype animals. p-value<0.05 were taken as targets misregulated in absence of ad-1 and adr-2. Differentially expressed genes are listed as Wormbase annotations in Column A, Gene name in Column B. The log2fold change values, raw p values and the adjusted p values as obtained from the DESeq2 analysis are shown in columns C, D, and E, respectively. Read counts from the DESeq2 analysis for the three biological replicates of wildtype are listed in columns F, G and H. Read counts from the DESeq2 analysis for the three biological replicates of adr-1(-);adr-2(-) animals are listed in columns I, J and K . "	"Supplemental Table 4: Differentially expressed genes in adr-1(-);adr-2(-) animals grown on OP40 for 7 hours compared to wildtype animals grown on OP50 for 7 hours (Related to Figure 4). DESeq2 was used to identify differentially expressed genes in adr-1(-);adr-2(-) double mutant animals compared to wildtype animals. p-value<0.05 were taken as targets misregulated in absence of ad-1 and adr-2. Differentially expressed genes are listed as Wormbase annotations in Column A, Gene name in Column B. The log2fold change values, raw p values and the adjusted p values as obtained from the DESeq2 analysis are shown in columns C, D, and E, respectively. Read counts from the DESeq2 analysis for the three biological replicates of wildtype are listed in columns F, G and H. Read counts from the DESeq2 analysis for the three biological replicates of adr-1(-);adr-2(-) animals are listed in columns I, J and K . "	"Supplemental Table 4: Differentially expressed genes in adr-1(-);adr-2(-) animals grown on OP40 for 7 hours compared to wildtype animals grown on OP50 for 7 hours (Related to Figure 4). DESeq2 was used to identify differentially expressed genes in adr-1(-);adr-2(-) double mutant animals compared to wildtype animals. p-value<0.05 were taken as targets misregulated in absence of ad-1 and adr-2. Differentially expressed genes are listed as Wormbase annotations in Column A, Gene name in Column B. The log2fold change values, raw p values and the adjusted p values as obtained from the DESeq2 analysis are shown in columns C, D, and E, respectively. Read counts from the DESeq2 analysis for the three biological replicates of wildtype are listed in columns F, G and H. Read counts from the DESeq2 analysis for the three biological replicates of adr-1(-);adr-2(-) animals are listed in columns I, J and K . "	"Supplemental Table 4: Differentially expressed genes in adr-1(-);adr-2(-) animals grown on OP40 for 7 hours compared to wildtype animals grown on OP50 for 7 hours (Related to Figure 4). DESeq2 was used to identify differentially expressed genes in adr-1(-);adr-2(-) double mutant animals compared to wildtype animals. p-value<0.05 were taken as targets misregulated in absence of ad-1 and adr-2. Differentially expressed genes are listed as Wormbase annotations in Column A, Gene name in Column B. The log2fold change values, raw p values and the adjusted p values as obtained from the DESeq2 analysis are shown in columns C, D, and E, respectively. Read counts from the DESeq2 analysis for the three biological replicates of wildtype are listed in columns F, G and H. Read counts from the DESeq2 analysis for the three biological replicates of adr-1(-);adr-2(-) animals are listed in columns I, J and K . "	"Supplemental Table 4: Differentially expressed genes in adr-1(-);adr-2(-) animals grown on OP40 for 7 hours compared to wildtype animals grown on OP50 for 7 hours (Related to Figure 4). DESeq2 was used to identify differentially expressed genes in adr-1(-);adr-2(-) double mutant animals compared to wildtype animals. p-value<0.05 were taken as targets misregulated in absence of ad-1 and adr-2. Differentially expressed genes are listed as Wormbase annotations in Column A, Gene name in Column B. The log2fold change values, raw p values and the adjusted p values as obtained from the DESeq2 analysis are shown in columns C, D, and E, respectively. Read counts from the DESeq2 analysis for the three biological replicates of wildtype are listed in columns F, G and H. Read counts from the DESeq2 analysis for the three biological replicates of adr-1(-);adr-2(-) animals are listed in columns I, J and K . "	"Supplemental Table 4: Differentially expressed genes in adr-1(-);adr-2(-) animals grown on OP40 for 7 hours compared to wildtype animals grown on OP50 for 7 hours (Related to Figure 4). DESeq2 was used to identify differentially expressed genes in adr-1(-);adr-2(-) double mutant animals compared to wildtype animals. p-value<0.05 were taken as targets misregulated in absence of ad-1 and adr-2. Differentially expressed genes are listed as Wormbase annotations in Column A, Gene name in Column B. The log2fold change values, raw p values and the adjusted p values as obtained from the DESeq2 analysis are shown in columns C, D, and E, respectively. Read counts from the DESeq2 analysis for the three biological replicates of wildtype are listed in columns F, G and H. Read counts from the DESeq2 analysis for the three biological replicates of adr-1(-);adr-2(-) animals are listed in columns I, J and K . "	"Supplemental Table 4: Differentially expressed genes in adr-1(-);adr-2(-) animals grown on OP40 for 7 hours compared to wildtype animals grown on OP50 for 7 hours (Related to Figure 4). DESeq2 was used to identify differentially expressed genes in adr-1(-);adr-2(-) double mutant animals compared to wildtype animals. p-value<0.05 were taken as targets misregulated in absence of ad-1 and adr-2. Differentially expressed genes are listed as Wormbase annotations in Column A, Gene name in Column B. The log2fold change values, raw p values and the adjusted p values as obtained from the DESeq2 analysis are shown in columns C, D, and E, respectively. Read counts from the DESeq2 analysis for the three biological replicates of wildtype are listed in columns F, G and H. Read counts from the DESeq2 analysis for the three biological replicates of adr-1(-);adr-2(-) animals are listed in columns I, J and K . "	"Supplemental Table 4: Differentially expressed genes in adr-1(-);adr-2(-) animals grown on OP40 for 7 hours compared to wildtype animals grown on OP50 for 7 hours (Related to Figure 4). DESeq2 was used to identify differentially expressed genes in adr-1(-);adr-2(-) double mutant animals compared to wildtype animals. p-value<0.05 were taken as targets misregulated in absence of ad-1 and adr-2. Differentially expressed genes are listed as Wormbase annotations in Column A, Gene name in Column B. The log2fold change values, raw p values and the adjusted p values as obtained from the DESeq2 analysis are shown in columns C, D, and E, respectively. Read counts from the DESeq2 analysis for the three biological replicates of wildtype are listed in columns F, G and H. Read counts from the DESeq2 analysis for the three biological replicates of adr-1(-);adr-2(-) animals are listed in columns I, J and K . "	"Supplemental Table 4: Differentially expressed genes in adr-1(-);adr-2(-) animals grown on OP40 for 7 hours compared to wildtype animals grown on OP50 for 7 hours (Related to Figure 4). DESeq2 was used to identify differentially expressed genes in adr-1(-);adr-2(-) double mutant animals compared to wildtype animals. p-value<0.05 were taken as targets misregulated in absence of ad-1 and adr-2. Differentially expressed genes are listed as Wormbase annotations in Column A, Gene name in Column B. The log2fold change values, raw p values and the adjusted p values as obtained from the DESeq2 analysis are shown in columns C, D, and E, respectively. Read counts from the DESeq2 analysis for the three biological replicates of wildtype are listed in columns F, G and H. Read counts from the DESeq2 analysis for the three biological replicates of adr-1(-);adr-2(-) animals are listed in columns I, J and K . "	"Supplemental Table 4: Differentially expressed genes in adr-1(-);adr-2(-) animals grown on OP40 for 7 hours compared to wildtype animals grown on OP50 for 7 hours (Related to Figure 4). DESeq2 was used to identify differentially expressed genes in adr-1(-);adr-2(-) double mutant animals compared to wildtype animals. p-value<0.05 were taken as targets misregulated in absence of ad-1 and adr-2. Differentially expressed genes are listed as Wormbase annotations in Column A, Gene name in Column B. The log2fold change values, raw p values and the adjusted p values as obtained from the DESeq2 analysis are shown in columns C, D, and E, respectively. Read counts from the DESeq2 analysis for the three biological replicates of wildtype are listed in columns F, G and H. Read counts from the DESeq2 analysis for the three biological replicates of adr-1(-);adr-2(-) animals are listed in columns I, J and K . "	"Supplemental Table 4: Differentially expressed genes in adr-1(-);adr-2(-) animals grown on OP40 for 7 hours compared to wildtype animals grown on OP50 for 7 hours (Related to Figure 4). DESeq2 was used to identify differentially expressed genes in adr-1(-);adr-2(-) double mutant animals compared to wildtype animals. p-value<0.05 were taken as targets misregulated in absence of ad-1 and adr-2. Differentially expressed genes are listed as Wormbase annotations in Column A, Gene name in Column B. The log2fold change values, raw p values and the adjusted p values as obtained from the DESeq2 analysis are shown in columns C, D, and E, respectively. Read counts from the DESeq2 analysis for the three biological replicates of wildtype are listed in columns F, G and H. Read counts from the DESeq2 analysis for the three biological replicates of adr-1(-);adr-2(-) animals are listed in columns I, J and K . "	"Supplemental Table 4: Differentially expressed genes in adr-1(-);adr-2(-) animals grown on OP40 for 7 hours compared to wildtype animals grown on OP50 for 7 hours (Related to Figure 4). DESeq2 was used to identify differentially expressed genes in adr-1(-);adr-2(-) double mutant animals compared to wildtype animals. p-value<0.05 were taken as targets misregulated in absence of ad-1 and adr-2. Differentially expressed genes are listed as Wormbase annotations in Column A, Gene name in Column B. The log2fold change values, raw p values and the adjusted p values as obtained from the DESeq2 analysis are shown in columns C, D, and E, respectively. Read counts from the DESeq2 analysis for the three biological replicates of wildtype are listed in columns F, G and H. Read counts from the DESeq2 analysis for the three biological replicates of adr-1(-);adr-2(-) animals are listed in columns I, J and K . "	"Supplemental Table 4: Differentially expressed genes in adr-1(-);adr-2(-) animals grown on OP40 for 7 hours compared to wildtype animals grown on OP50 for 7 hours (Related to Figure 4). DESeq2 was used to identify differentially expressed genes in adr-1(-);adr-2(-) double mutant animals compared to wildtype animals. p-value<0.05 were taken as targets misregulated in absence of ad-1 and adr-2. Differentially expressed genes are listed as Wormbase annotations in Column A, Gene name in Column B. The log2fold change values, raw p values and the adjusted p values as obtained from the DESeq2 analysis are shown in columns C, D, and E, respectively. Read counts from the DESeq2 analysis for the three biological replicates of wildtype are listed in columns F, G and H. Read counts from the DESeq2 analysis for the three biological replicates of adr-1(-);adr-2(-) animals are listed in columns I, J and K . "	"Supplemental Table 4: Differentially expressed genes in adr-1(-);adr-2(-) animals grown on OP40 for 7 hours compared to wildtype animals grown on OP50 for 7 hours (Related to Figure 4). DESeq2 was used to identify differentially expressed genes in adr-1(-);adr-2(-) double mutant animals compared to wildtype animals. p-value<0.05 were taken as targets misregulated in absence of ad-1 and adr-2. Differentially expressed genes are listed as Wormbase annotations in Column A, Gene name in Column B. The log2fold change values, raw p values and the adjusted p values as obtained from the DESeq2 analysis are shown in columns C, D, and E, respectively. Read counts from the DESeq2 analysis for the three biological replicates of wildtype are listed in columns F, G and H. Read counts from the DESeq2 analysis for the three biological re
