## Additional File 9 for "ADARs regulate cuticle collagen expression to promote survival to pathogen infection": Additional File 9.pdf

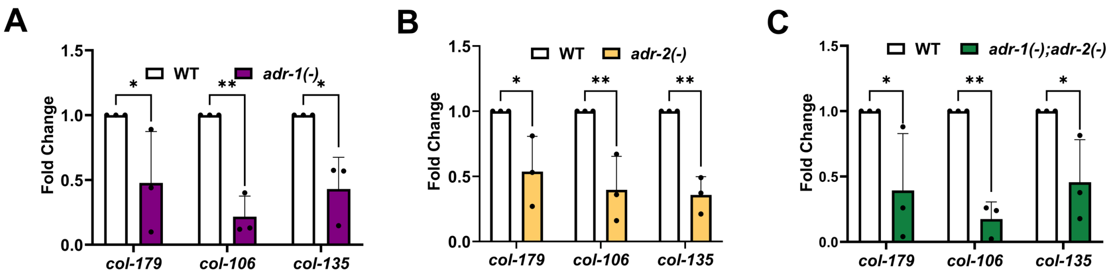


**Additional File 9: Fig. S9. qRT-PCR validation of downregulated collagen genes from RNA-seq analysis.** qRT-PCR quantification of the level of the indicated genes relative to *gpd-3* and normalized to the ratios obtained for wildtype (WT). The mean of three biological replicates for each strain was plotted; however, it should be noted the wildtype values used in A, B and C are the same biological replicates. Error bars represent SEM. Statistical significance was determined using a two-way ANOVA Sidak's multiple comparisons test. *$p \leq 0.05$, **$p \leq 0.005$.
